# Identification of potential and novel target genes in pituitary prolactinoma by bioinformatics analysis

**DOI:** 10.1101/2020.12.21.423732

**Authors:** Vikrant Ghatnatti, Basavaraj Vastrad, Swetha Patil, Chanabasayya Vastrad, Iranna Kotturshetti

## Abstract

Pituitary prolactinoma is one of the most complicated and fatally pathogenic pituitary adenomas. Therefore, there is an urgent need to improve our understanding of the underlying molecular mechanism that drives the initiation, progression, and metastasis of pituitary prolactinoma. The aim of the present study was to identify the key genes and signaling pathways associated with pituitary prolactinoma using bioinformatics analysis. Transcriptome microarray dataset GSE119063 was acquired from Gene Expression Omnibus datasets, which included 5 pituitary prolactinoma samples and 4 normal pituitaries samples. We screened differentially expressed genes (DEGs) with limma and investigated their biological function by pathway and Gene Ontology (GO) enrichment analysis. A protein-protein interaction (PPI) network of the up and down DEGs were constructed and analyzed by HIPPIE and Cytoscape software. Module analyses were performed. In addition, a target gene - miRNA network and target gene - TF network of the up and down DEGs were constructed by NetworkAnalyst and Cytoscape software. The set of DEGs exhibited an intersection consisting of 989 genes (461 up-regulated and 528 down-regulated), which may be associated with pituitary prolactinoma. Pathway enrichment analysis showed that the 989 DEGs were significantly enriched in the retinoate biosynthesis II, signaling pathways regulating pluripotency of stem cells, ALK2 signaling events, vitamin D3 biosynthesis, cell cycle and aurora B signaling. Gene Ontology (GO) enrichment analysis also showed that sensory organ morphogenesis, extracellular matrix, hormone activity, nuclear division, condensed chromosome and microtubule binding. In the PPI network and modules, SOX2, PRSS45, CLTC, PLK1, B4GALT6, RUNX1 and GTSE1 were considered as hub genes. In the target gene miRNA network and target gene - TF network, LINC00598, SOX4, IRX1 and UNC13A were considered as hub genes. Using integrated bioinformatics analysis, we identified candidate genes in pituitary prolactinoma, which may improve our understanding of the mechanisms of the pathogenesis and integration; genes may be therapeutic targets and prognostic markers for pituitary prolactinoma.

## Introduction

Prolactinoma is named as prolactin secreting pituitary adenoma seen more frequently in women and is characterized by irregular menstrual, erectile dysfunction, eye problems and loss of sexual function [1]. Its typical signs and symptoms include amenorrhoea, glactorrhoea, headache, anaemia and hypertension [2]. Prolactinoma involves 20.7% among other pituitary adenomas [3]. Although treatment methods, including surgery [4], chemotherapy [5] and radiotherapy [6] are improving, overall survival rate remains lower. Consequently, elucidating the molecular mechanism associated in the proliferation of pituitary prolactinoma is essential for the improvement of efficacious diagnosis and treatment strategies.

Presently, a wealth of previous studies has been enforced to advance a better understanding of the molecular mechanisms of pituitary prolactinoma. One study showed that allelic loss of DRD2 is responsible for development of pituitary prolactinoma [7]. Elevated expression of HMGA1 and HMGA2 are diagnosed with pituitary prolactinoma [8–9]. BMP4 is associated with SMAD4 and ER in development of pituitary prolactinoma [10]. FGF4 is responsible for improvement of pituitary prolactinoma through invasion and cell proliferation [11]. Alteration in oncogene GNAS is important for improvement of pituitary prolactinoma [12]. Stimulation of Raf/MEK/ERK and PI3K/Akt/mTOR signaling pathway is liable for development of prolactinoma [13]. Abnormal expression of CTNNB1 is diagnosed with prolactinoma [14]. Epigenetic inactivation of CDKN2A is responsible for advancement of prolactinoma [15]. AIP play key role in pathogenesis of pituitary prolactinoma [16]. Hence, searching for specific and sensitive molecular marker as well as some core genes or proteins as therapeutic target will benefit the diagnosis and treatment of pituitary prolactinoma.

At current study, microarray analysis has been applied as a very critical tool for medical research [17]. In this analysis, we chose GSE119063 dataset from Gene Expression Omnibus (GEO) (http:// www.ncbi.nlm.nih.gov/geo/) and used limma bioconductor package to find the differentially expressed genes (DEGs). Subsequently, we made pathway enrichment and gene ontology (biological process (BP), molecular function (MF), cellular component (CC) analyses were performed. In addition, we made PPI network of the DEGs and selected core genes with a high degree of connectivity, high betweenness centrality, high stress centrality, high closeness centrality and low Clustering coefficient, and modules analysis were performed. Moreover, miRNA-target gene regulatory network and TF-target gene regulatory network were constructed and analyzed. Briefly, this study would provide novel targets for diagnosis and treatment of pituitary prolactinoma.

## Results

### Identification of DEGs

After data preprocessing, the raw data of nine samples is proved to be eligible (Fig. 1). The GSE119063 expression profile data from GEO was investigated to screen for DEGs between the experimental and control groups. Under the threshold of FDR <0.05, and fold change ≥ 0.93 for up regulated gene and fold change ≥ -0.29 for down regulated gene. Comparison of prolactinoma with normal pituitaries identified total of 989 DEGs, including 461 up regulated genes and 528 down regulated genes, were revealed (Table 1). A corresponding heat map is shown in Fig. 2 and Fig 3. All the DEGs were presented by volcano plot in the study (Fig. 4).

**Fig. 1.**
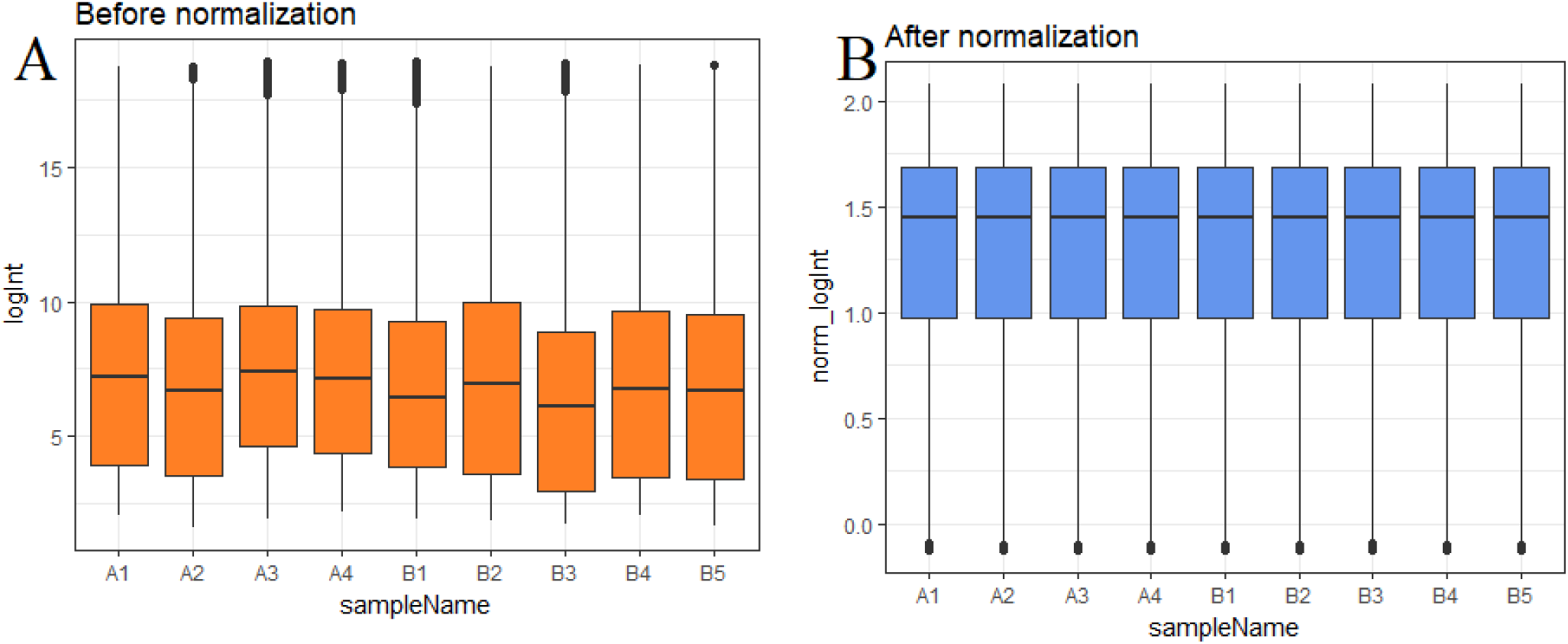
Box plots of the gene expression data before and after normalization. Horizontal axis represents the sample symbol and the vertical axis represents the gene expression values. The black line in the box plot represents the median value of gene expression. (A1, A2, A3, A4 = normal pituitaries samples; B1, B2, B3, B4, B5 = 5 pituitary prolactinoma)

**Fig. 2.**
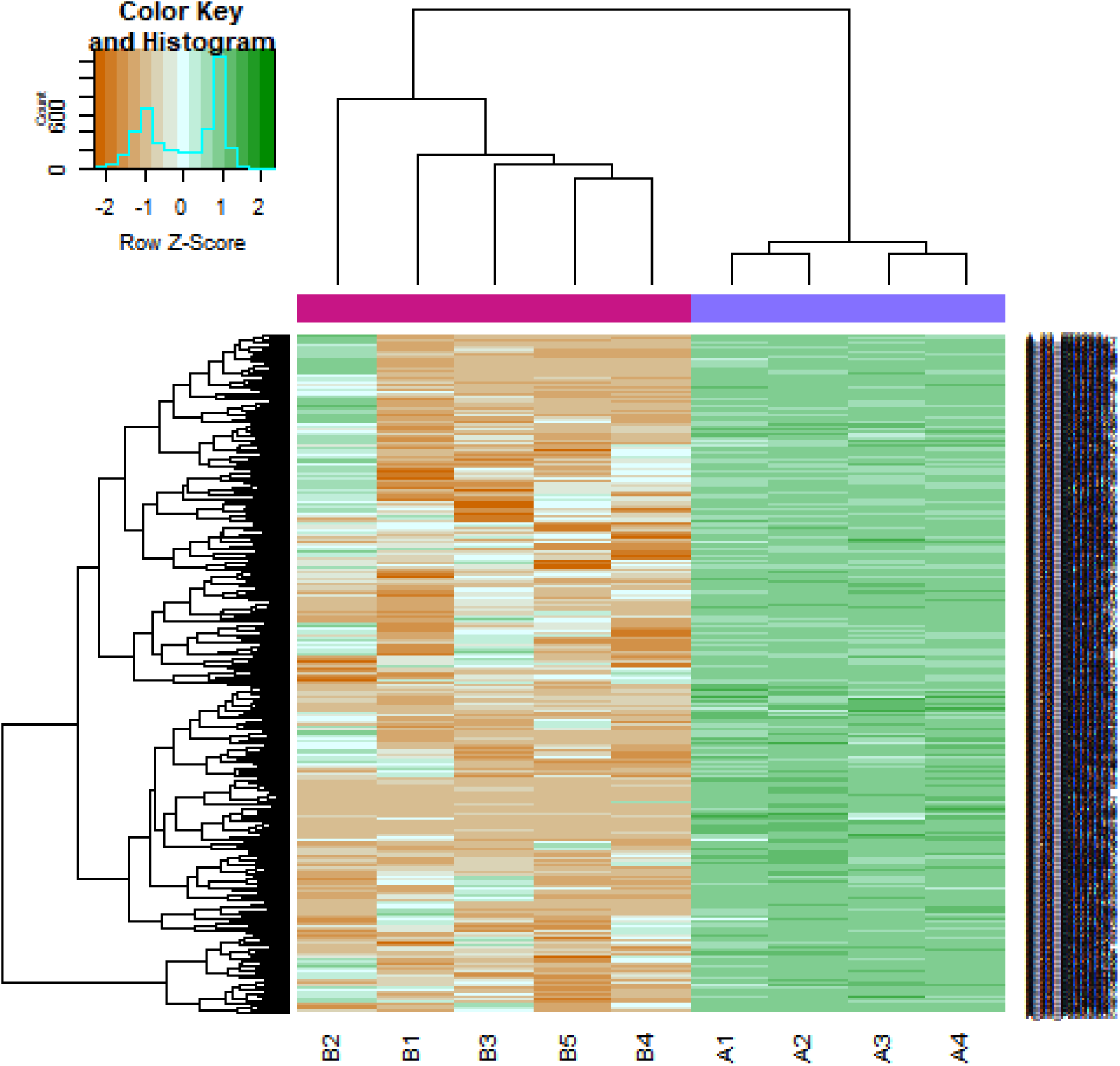
Heat map of up regulated differentially expressed genes. Legend on the top left indicate log fold change of genes. (A1, A2, A3, A4 = normal pituitaries samples; B1, B2, B3, B4, B5 = 5 pituitary prolactinoma)

**Fig. 3.**
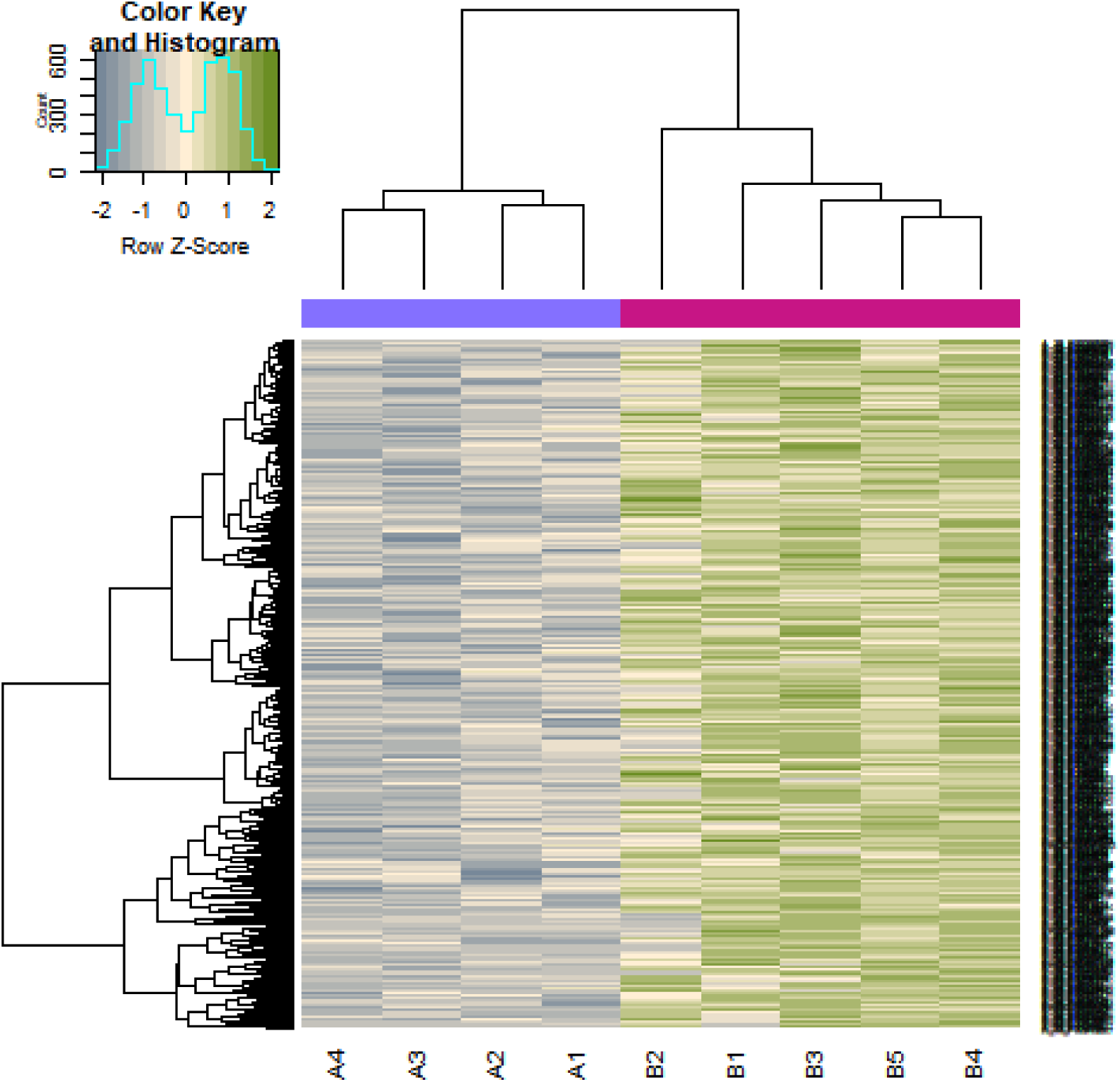
Heat map of down regulated differentially expressed genes. Legend on the top left indicate log fold change of genes. (A1, A2, A3, A4 = normal pituitaries samples; B1, B2, B3, B4, B5 = 5 pituitary prolactinoma)

**Fig. 4.**
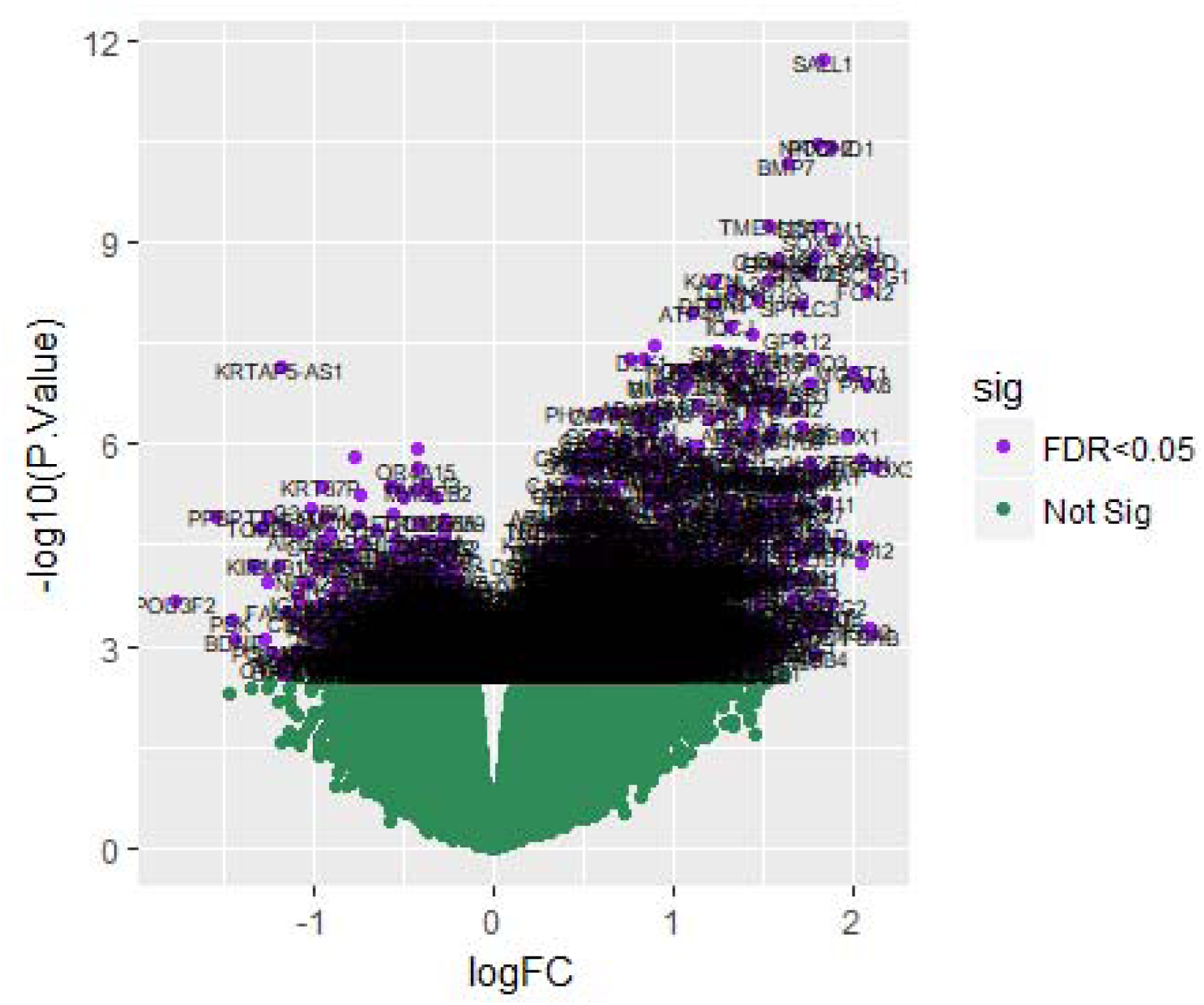
Volcano plot of differentially expressed genes. Genes with a significant change of more than two-fold were selected.

**Table 1.**
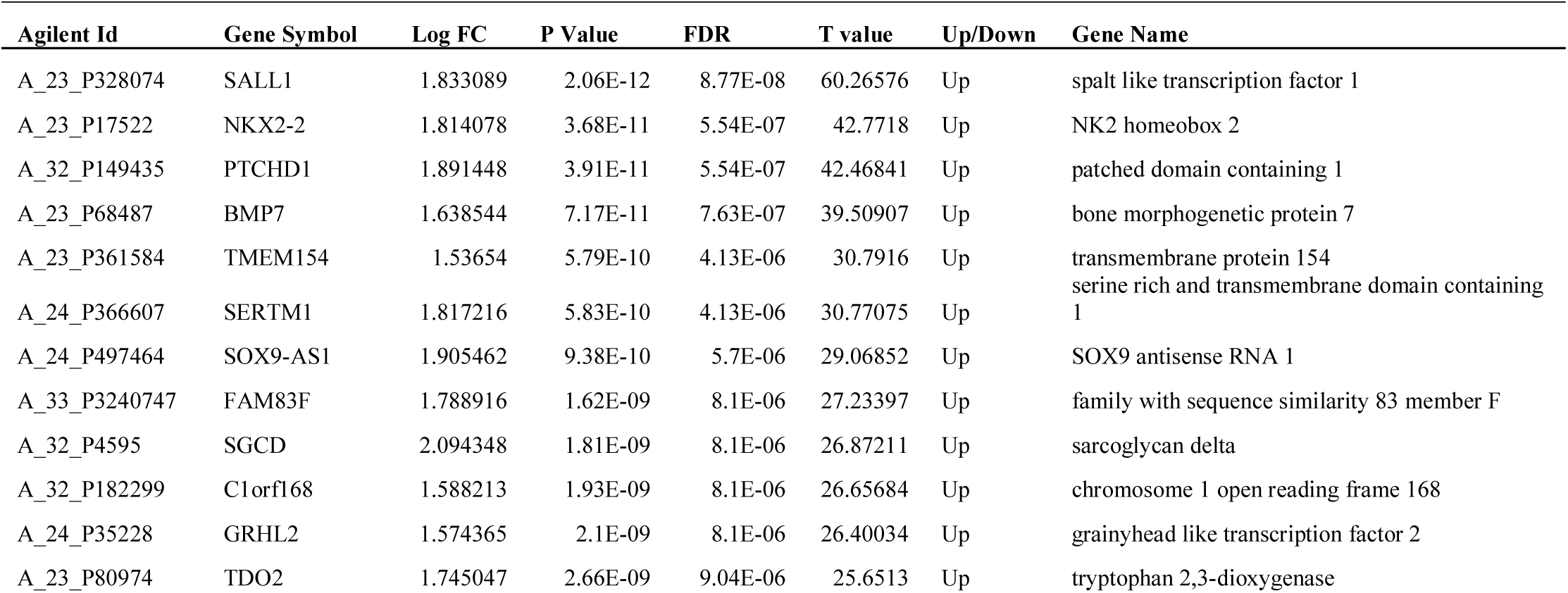
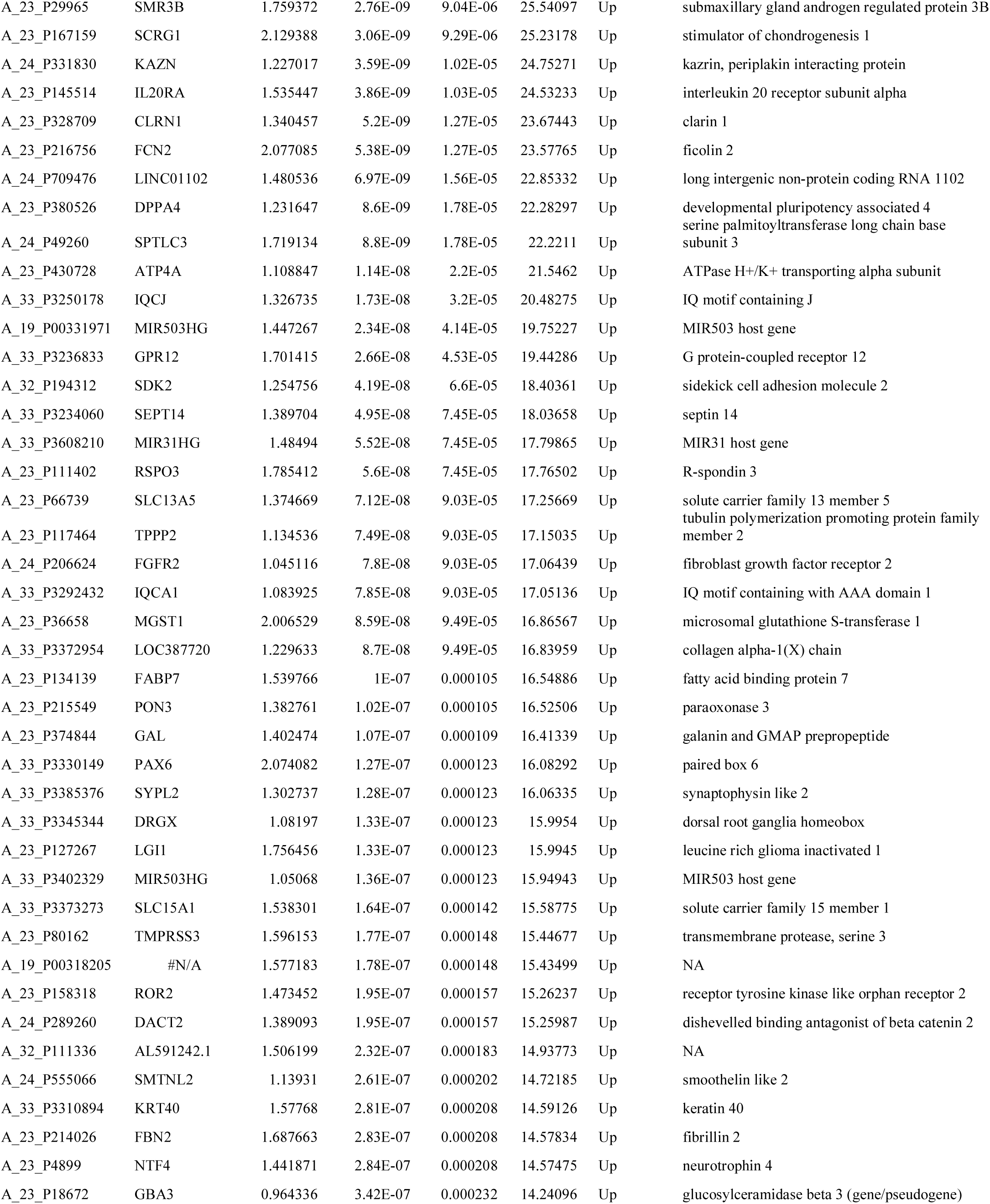
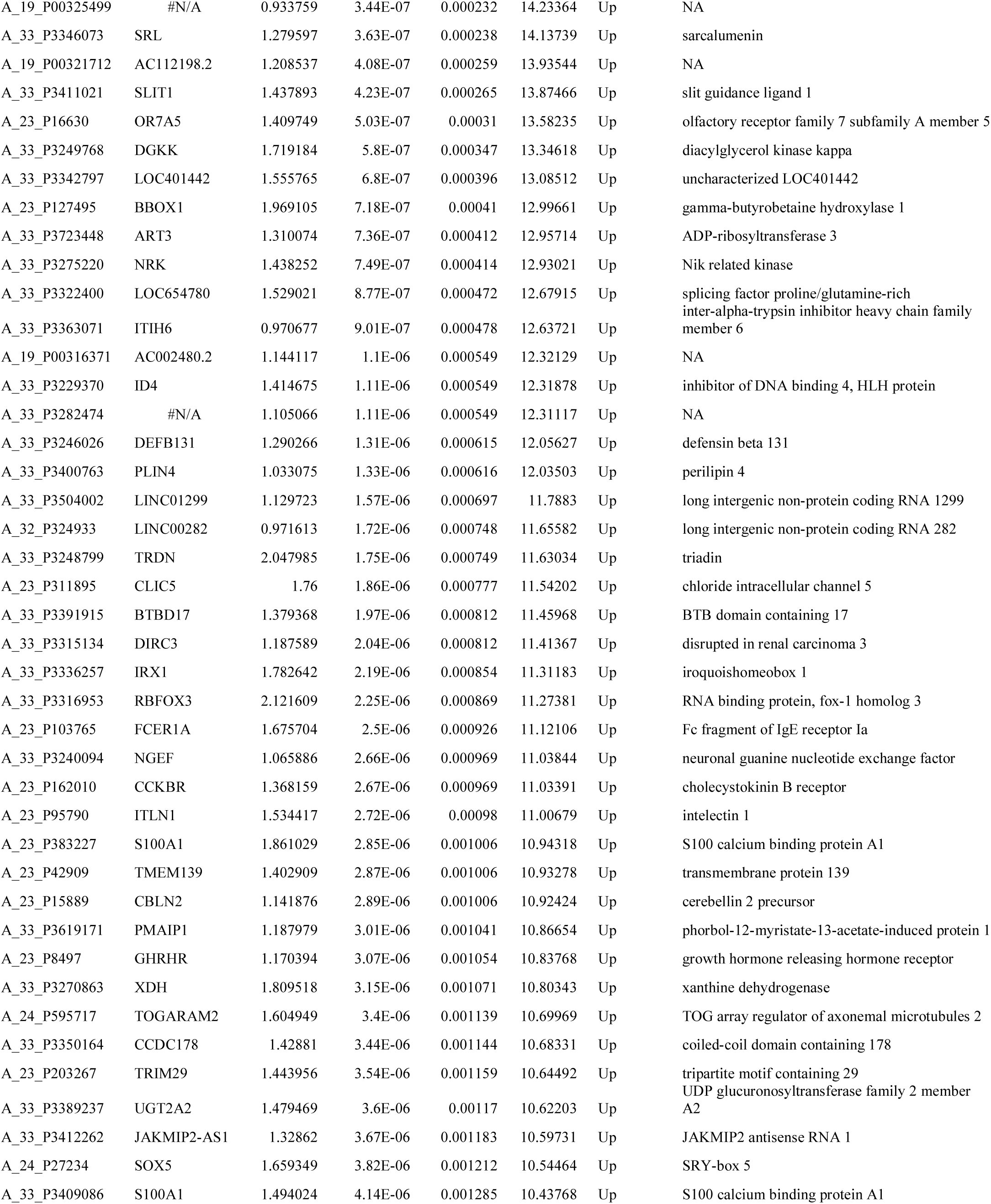
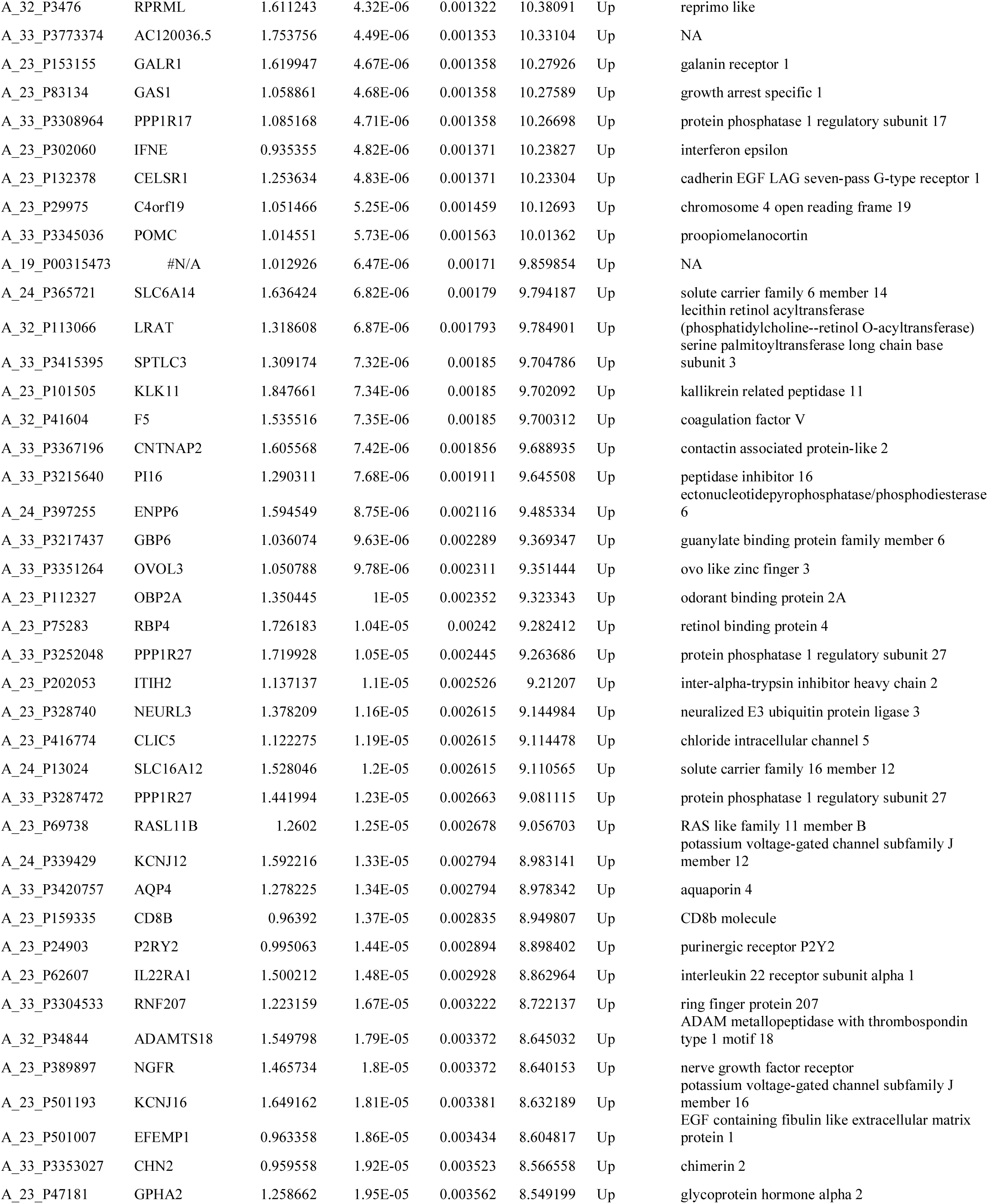
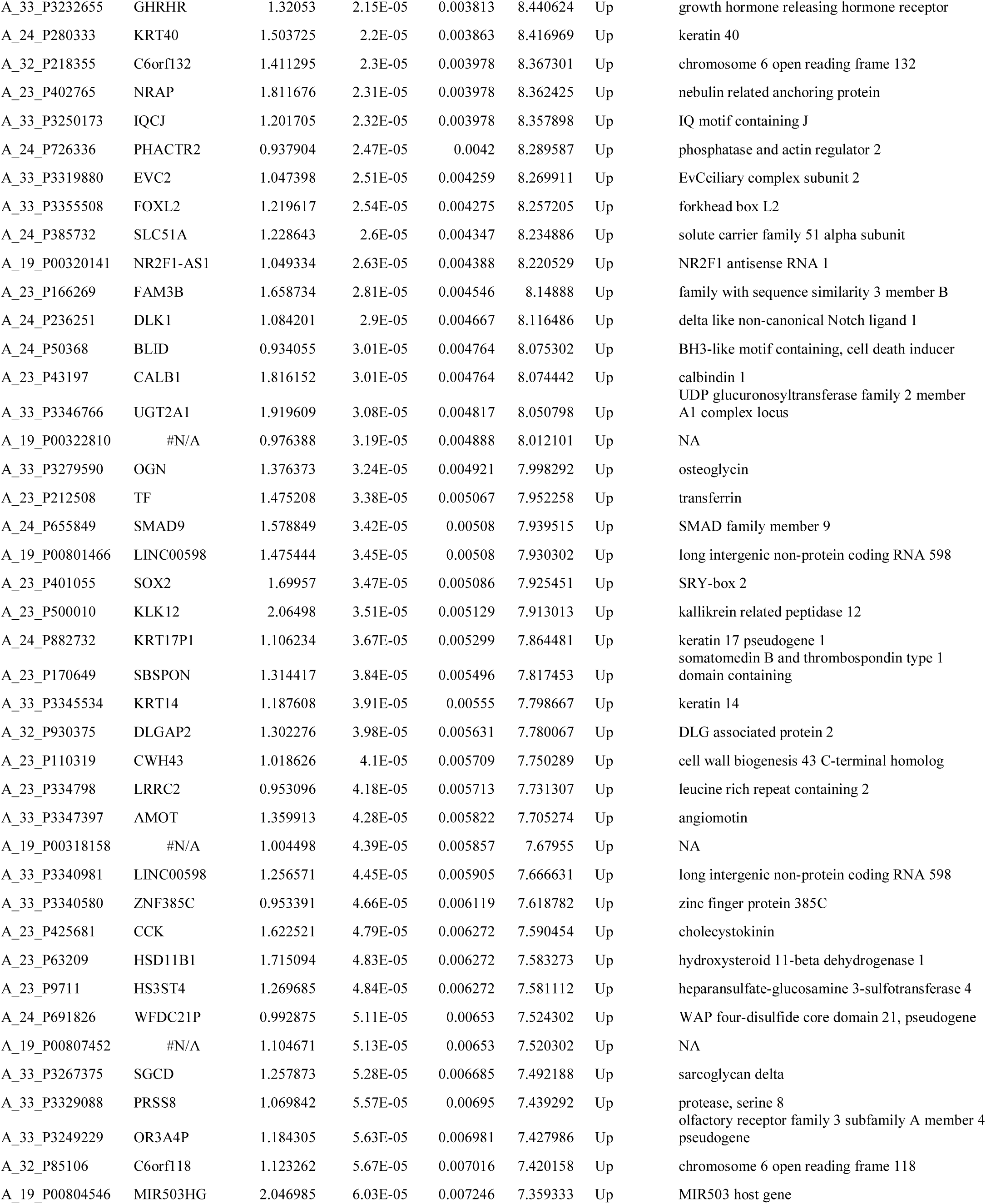
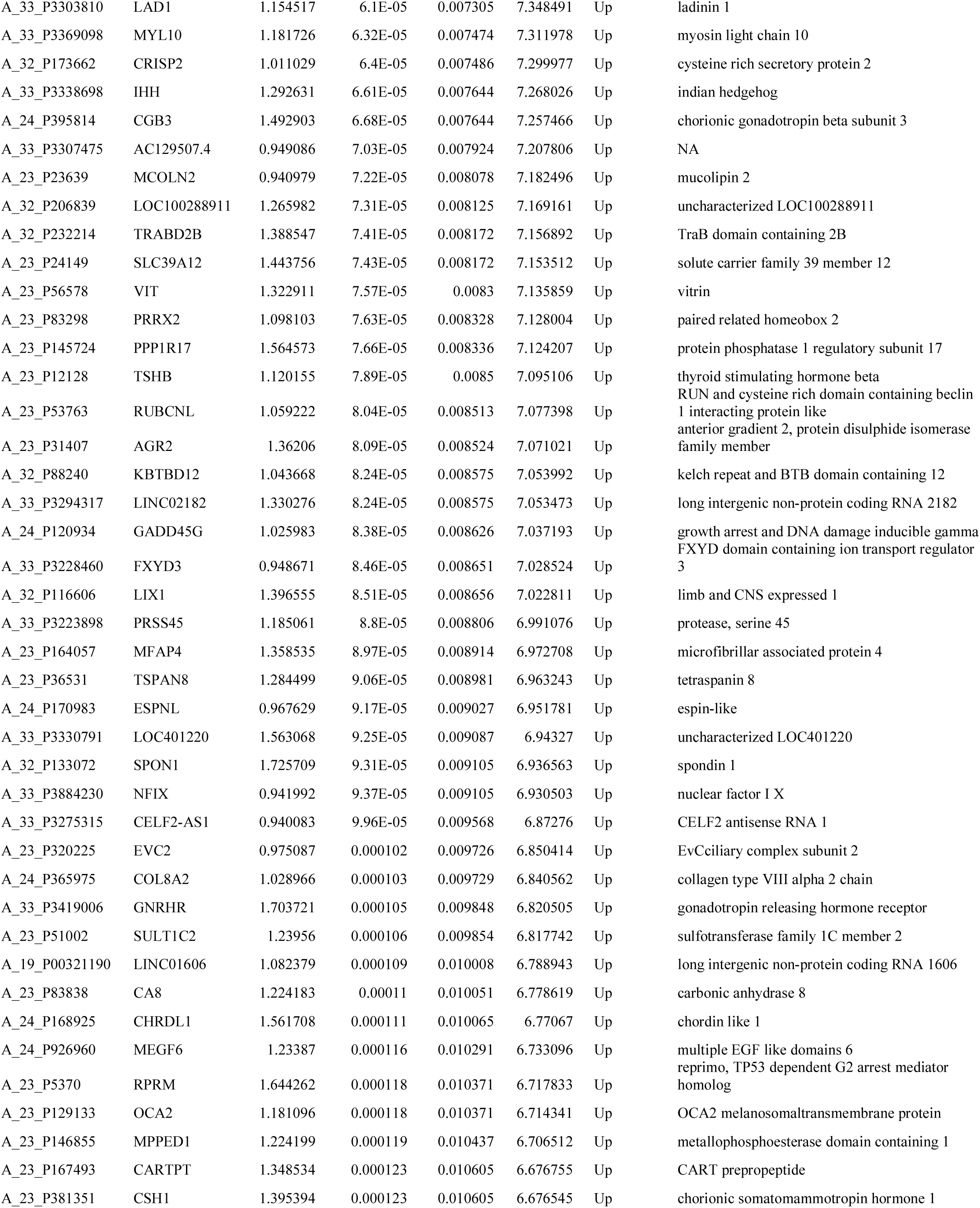
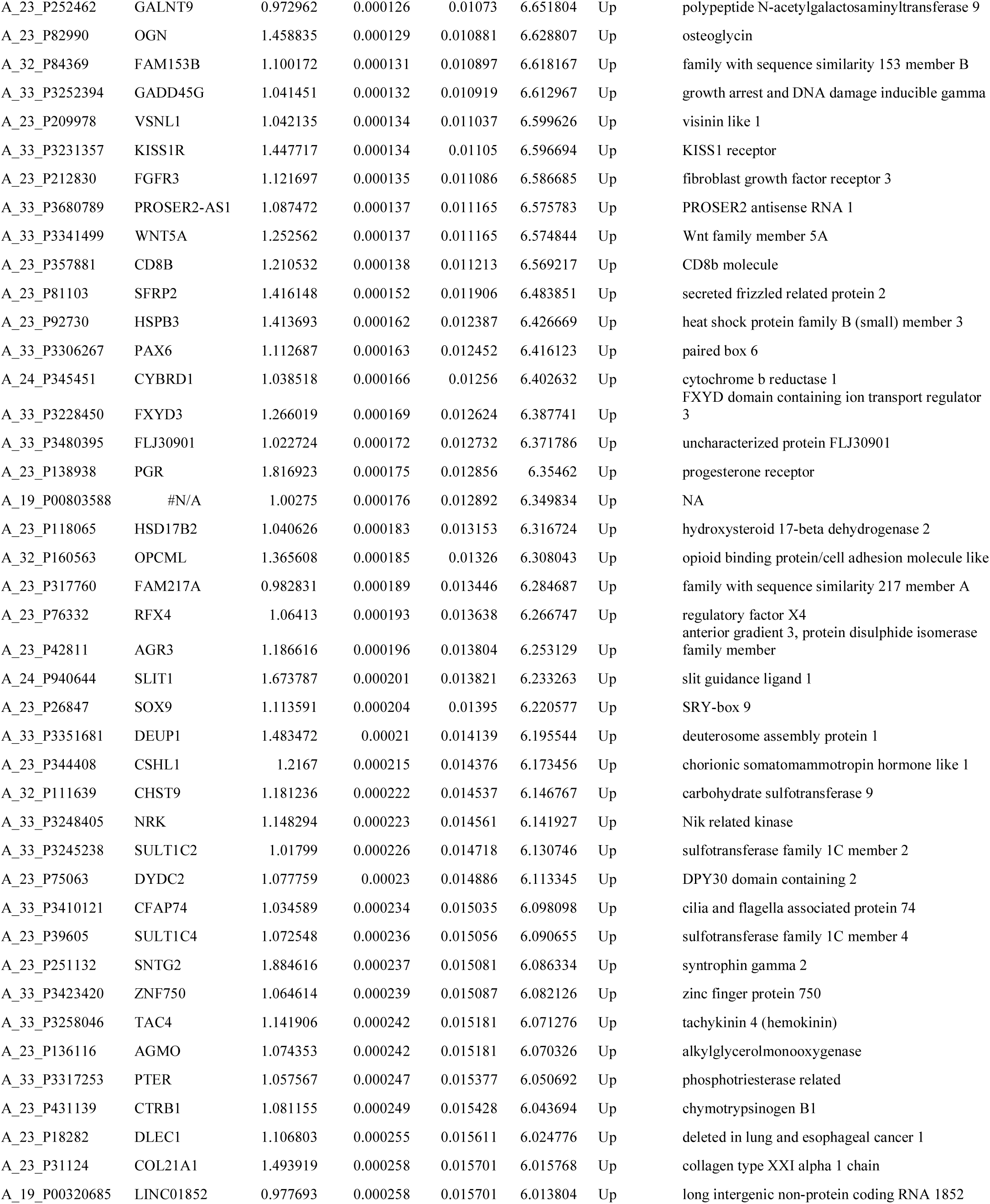
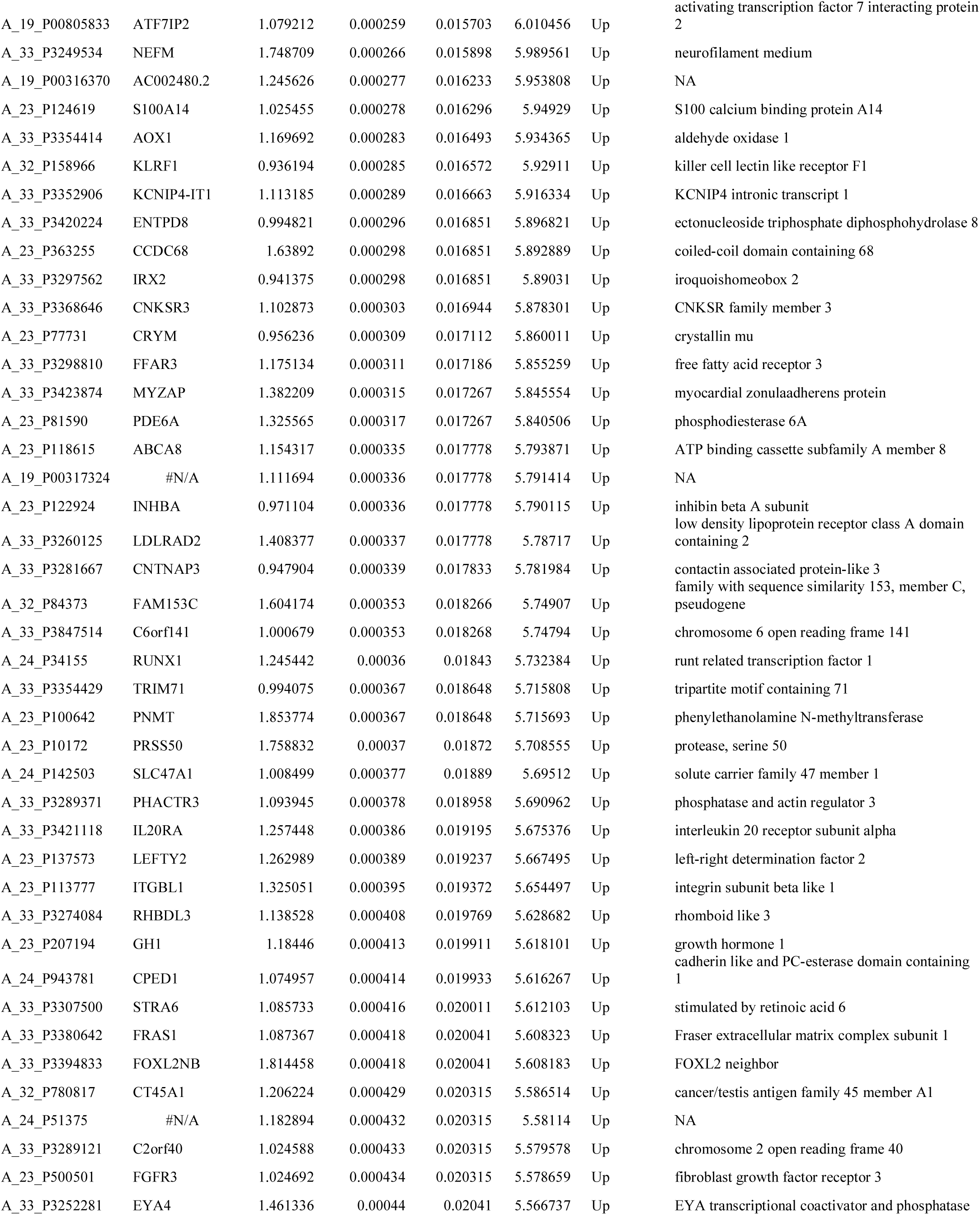
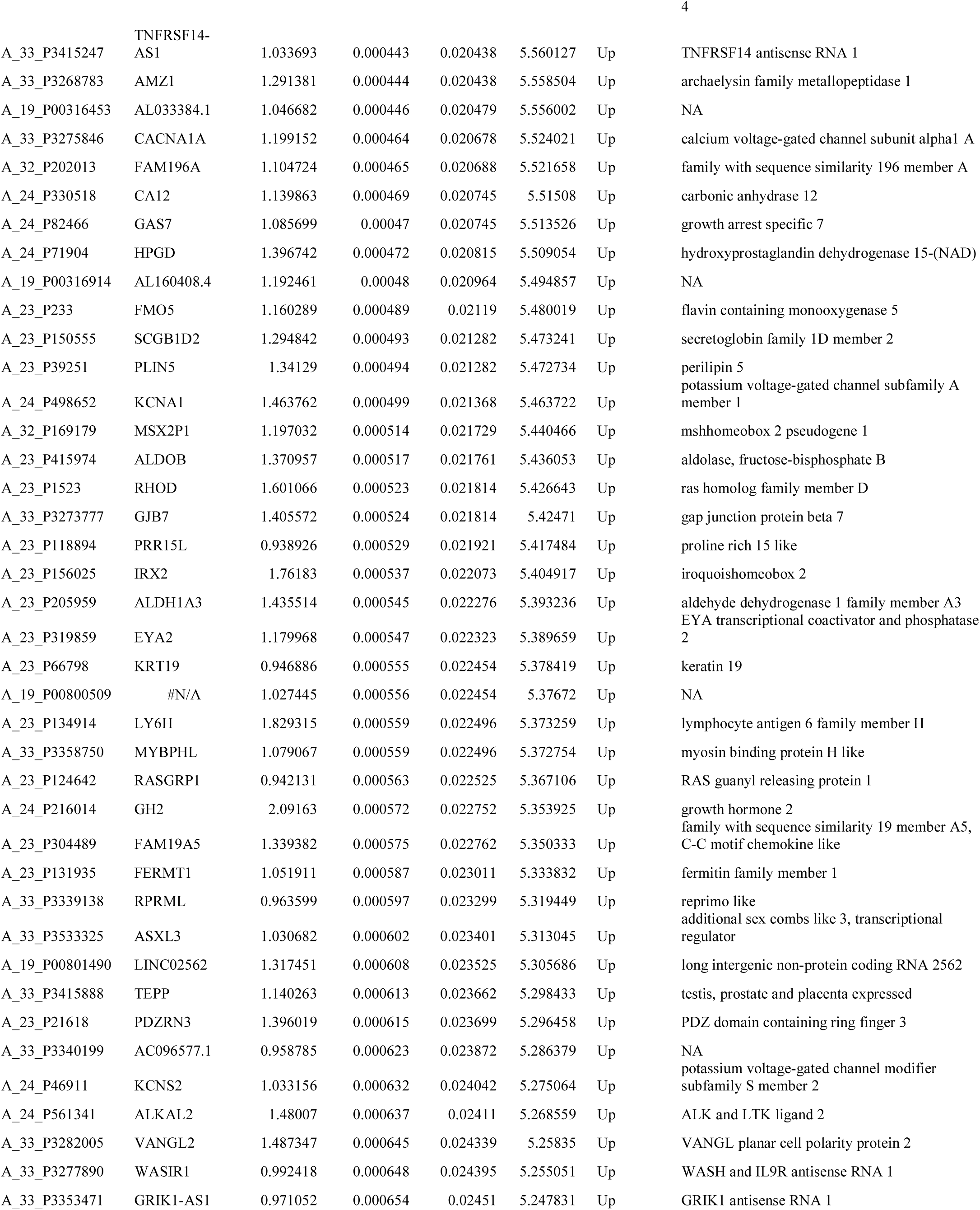
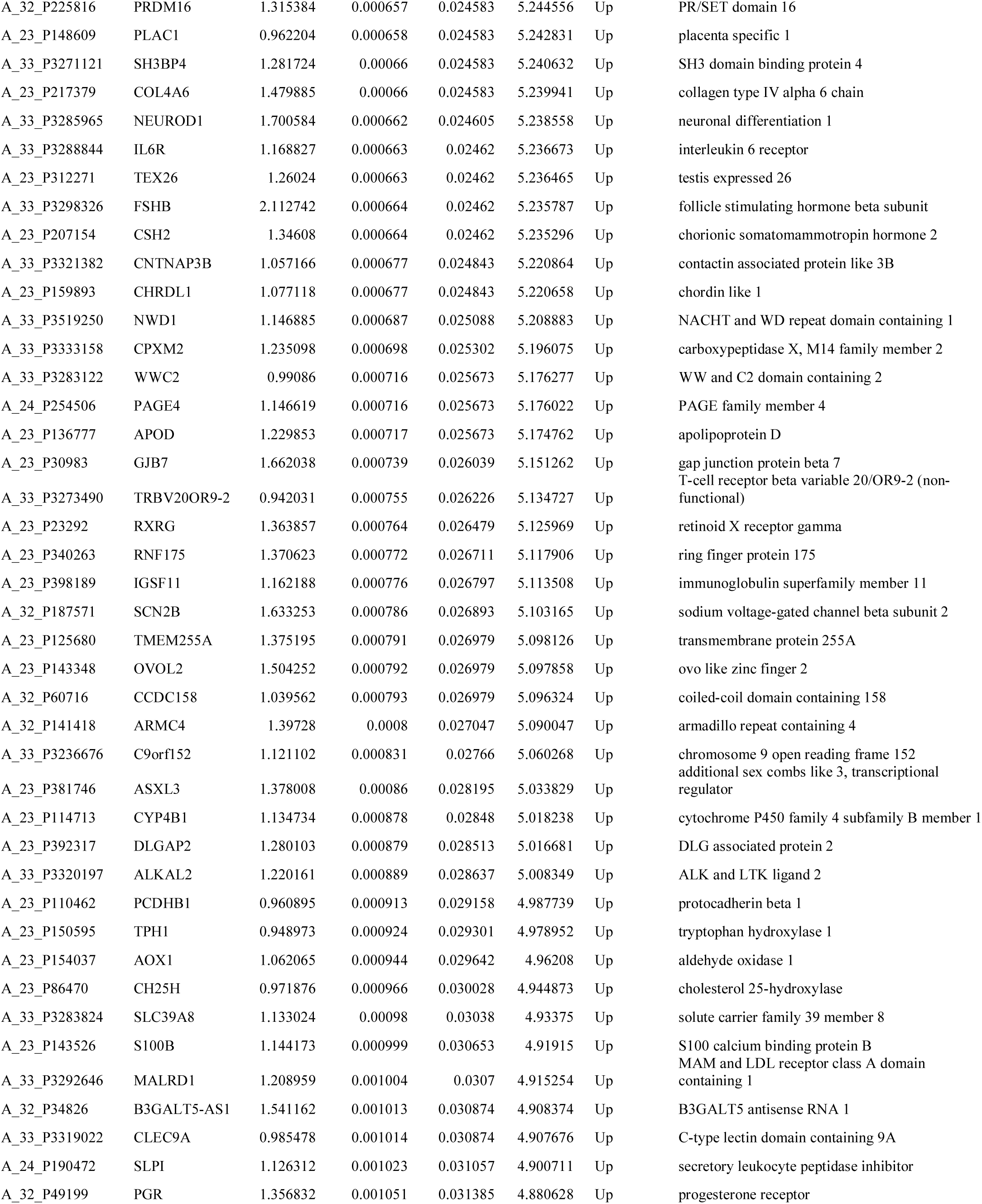
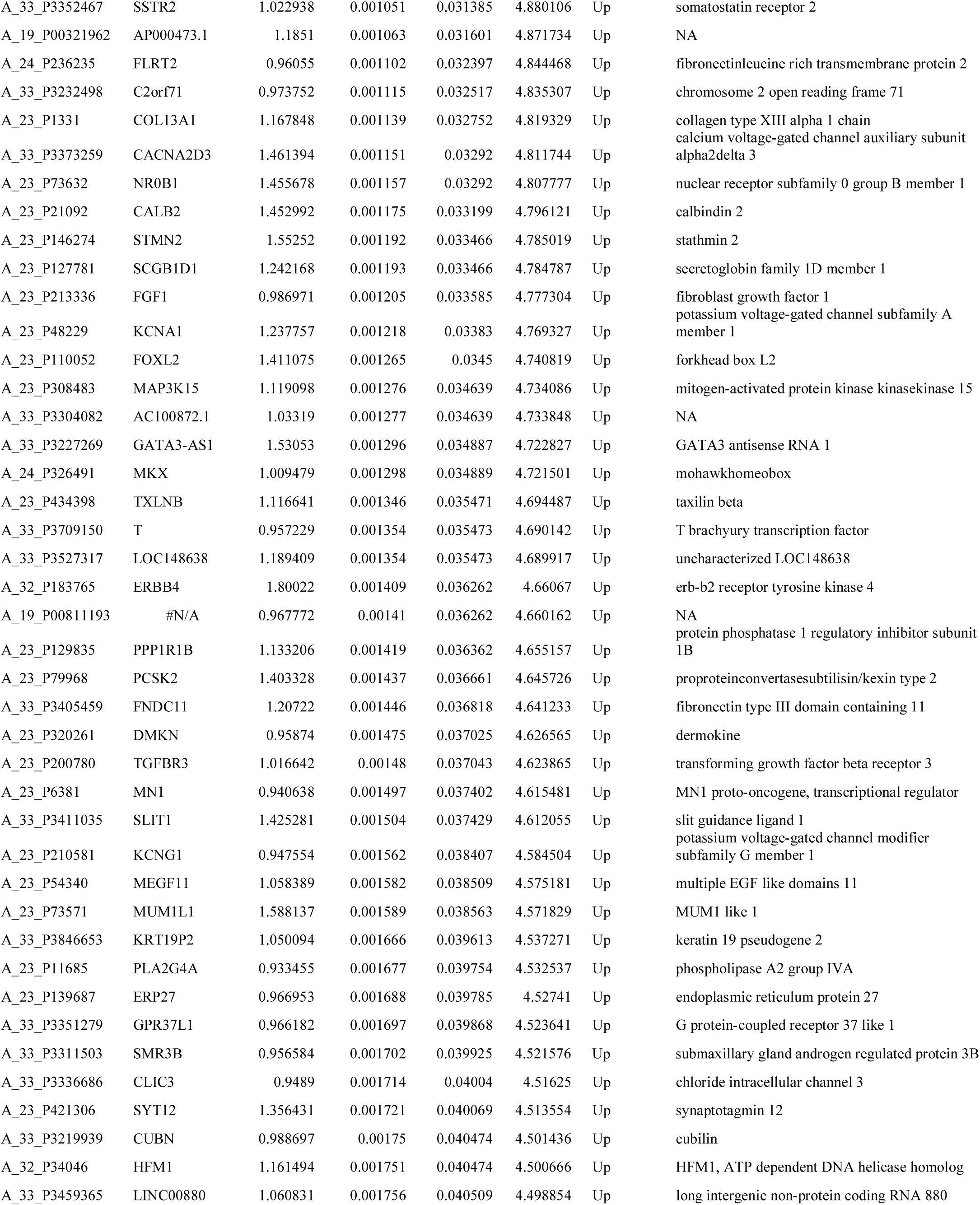
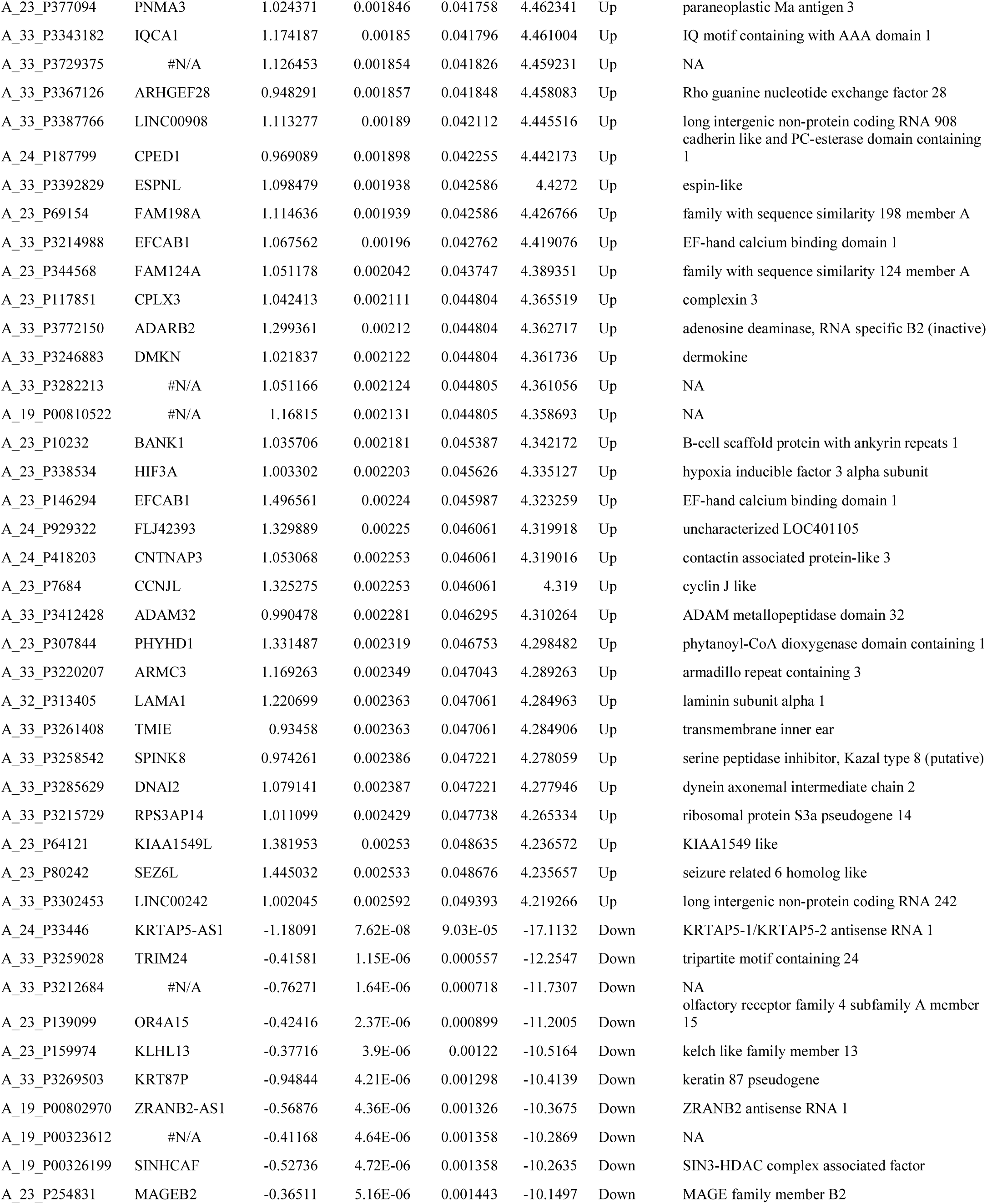
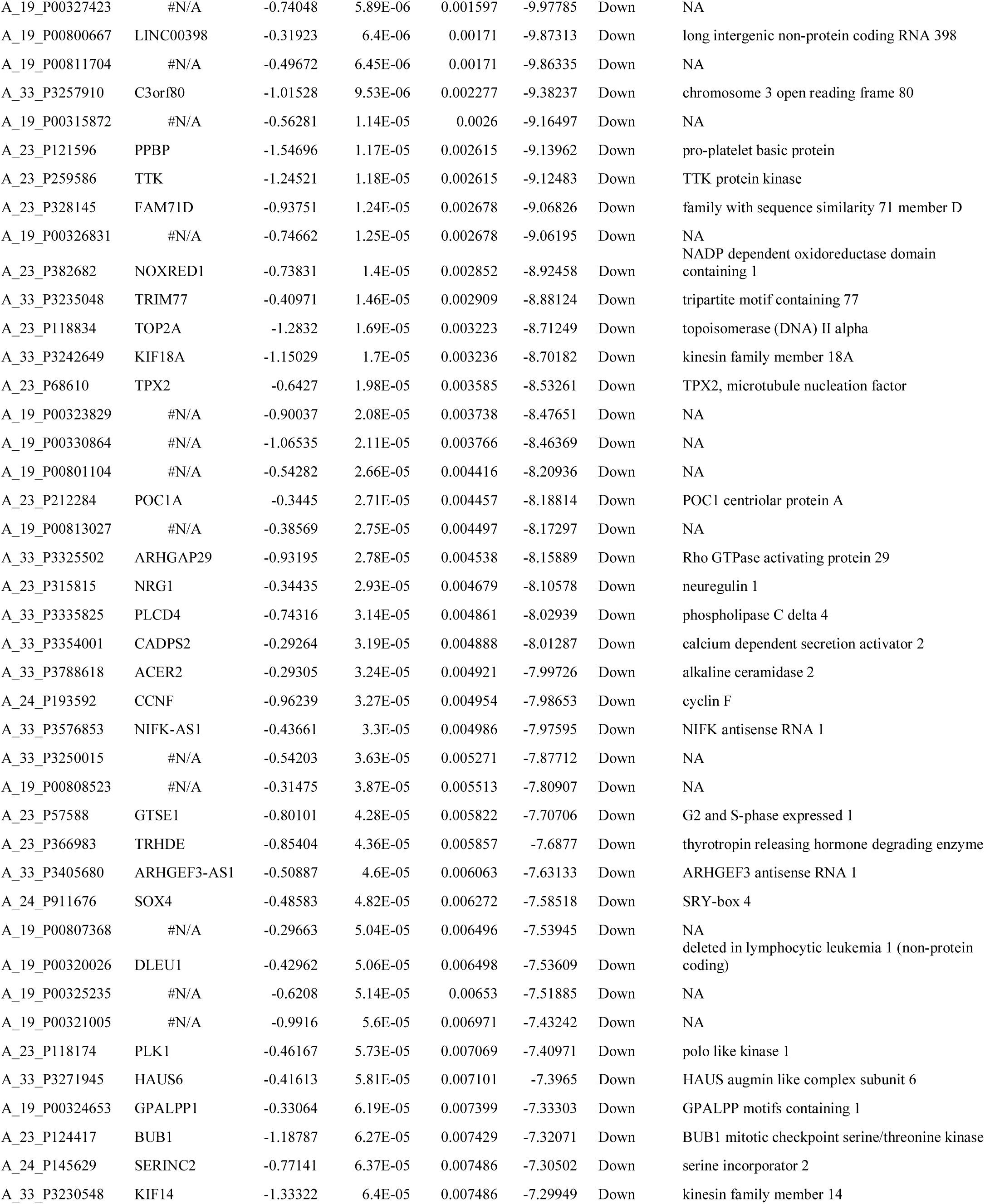
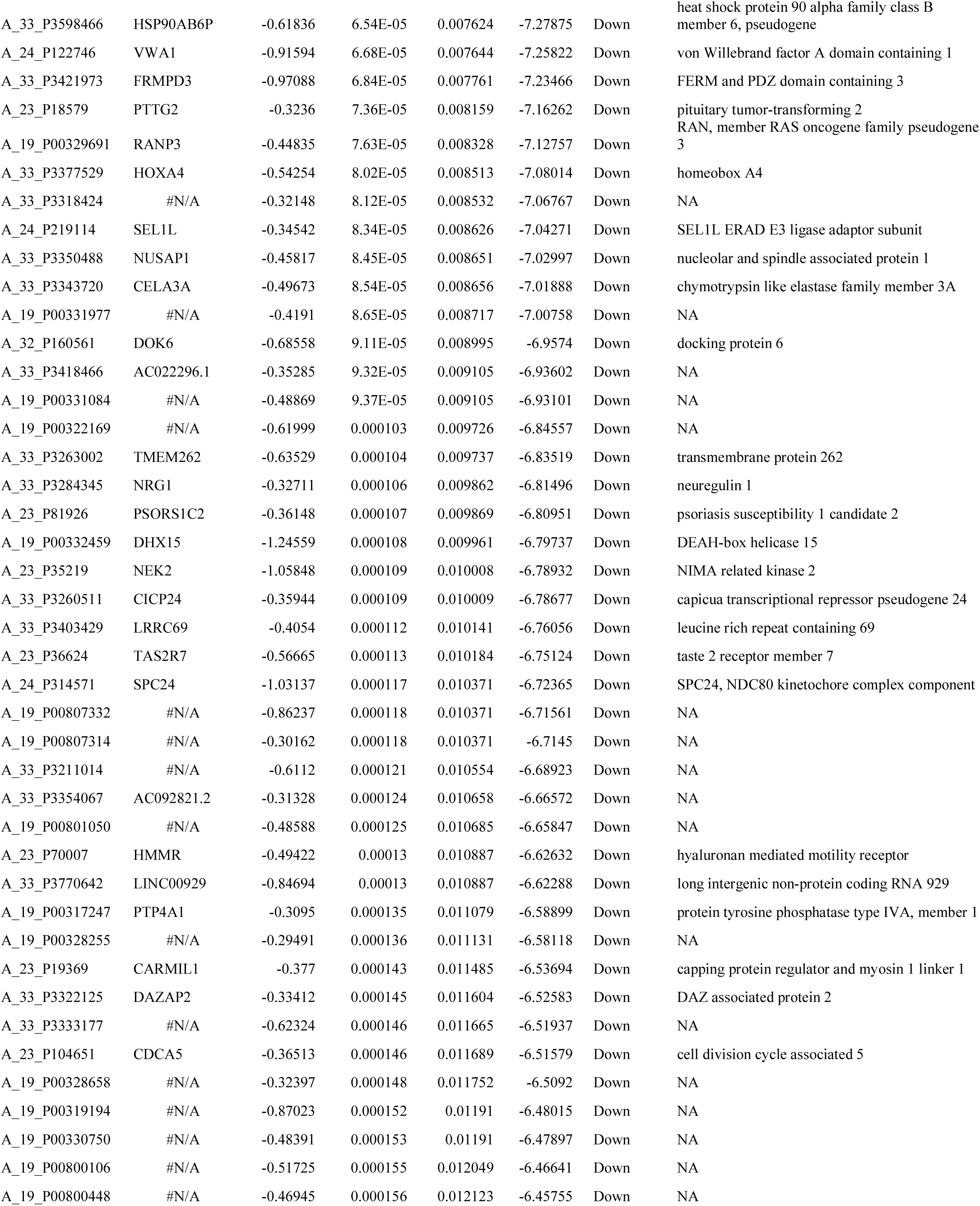
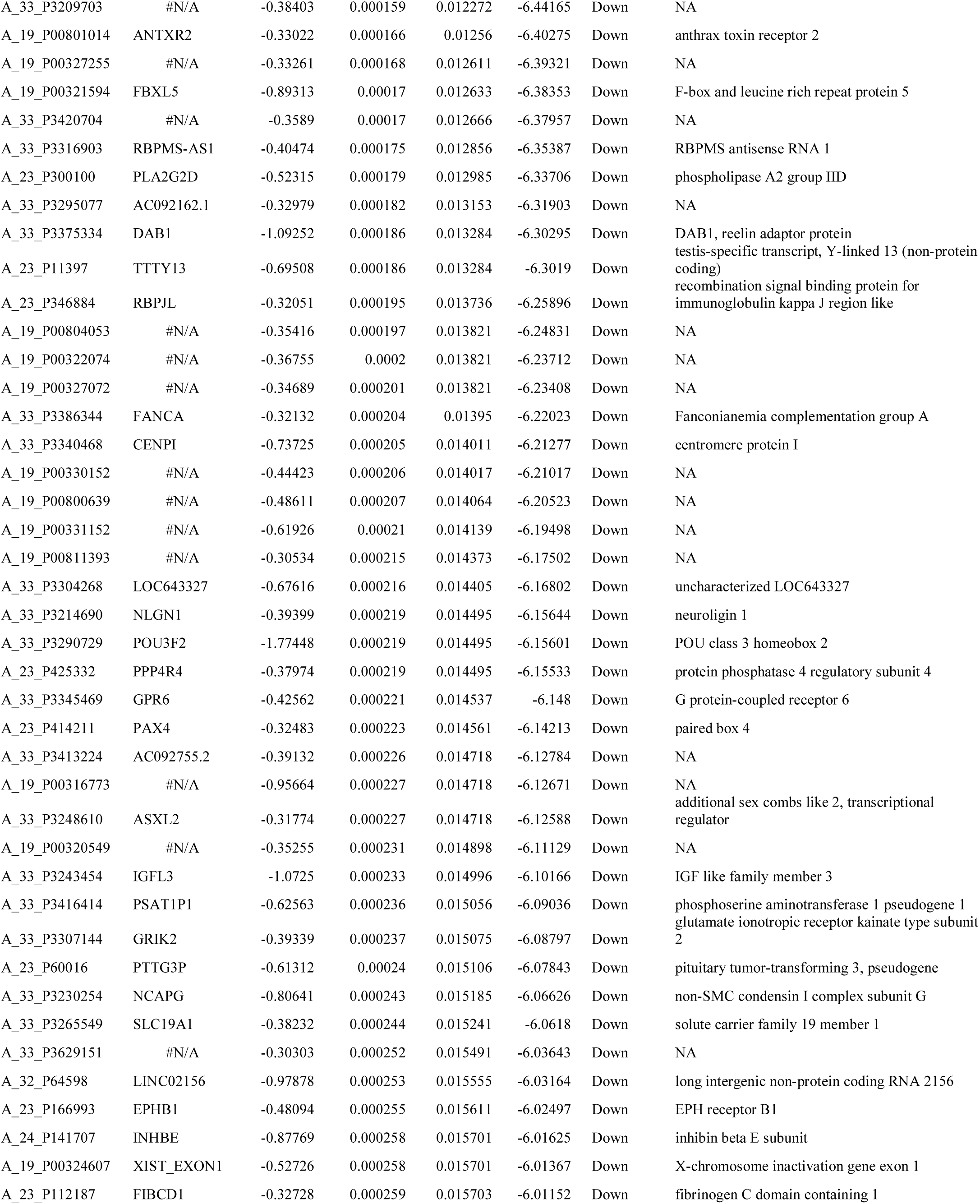
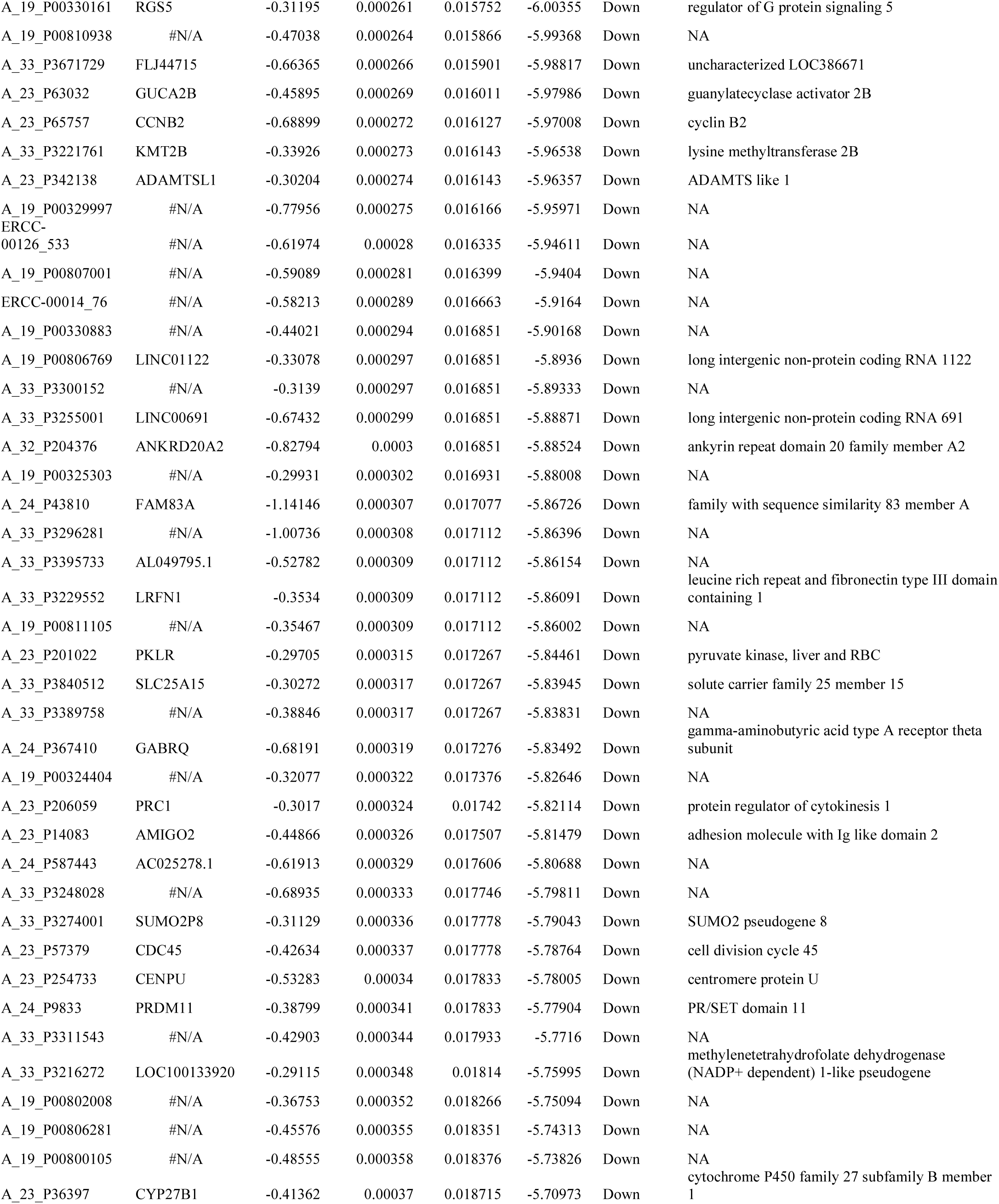
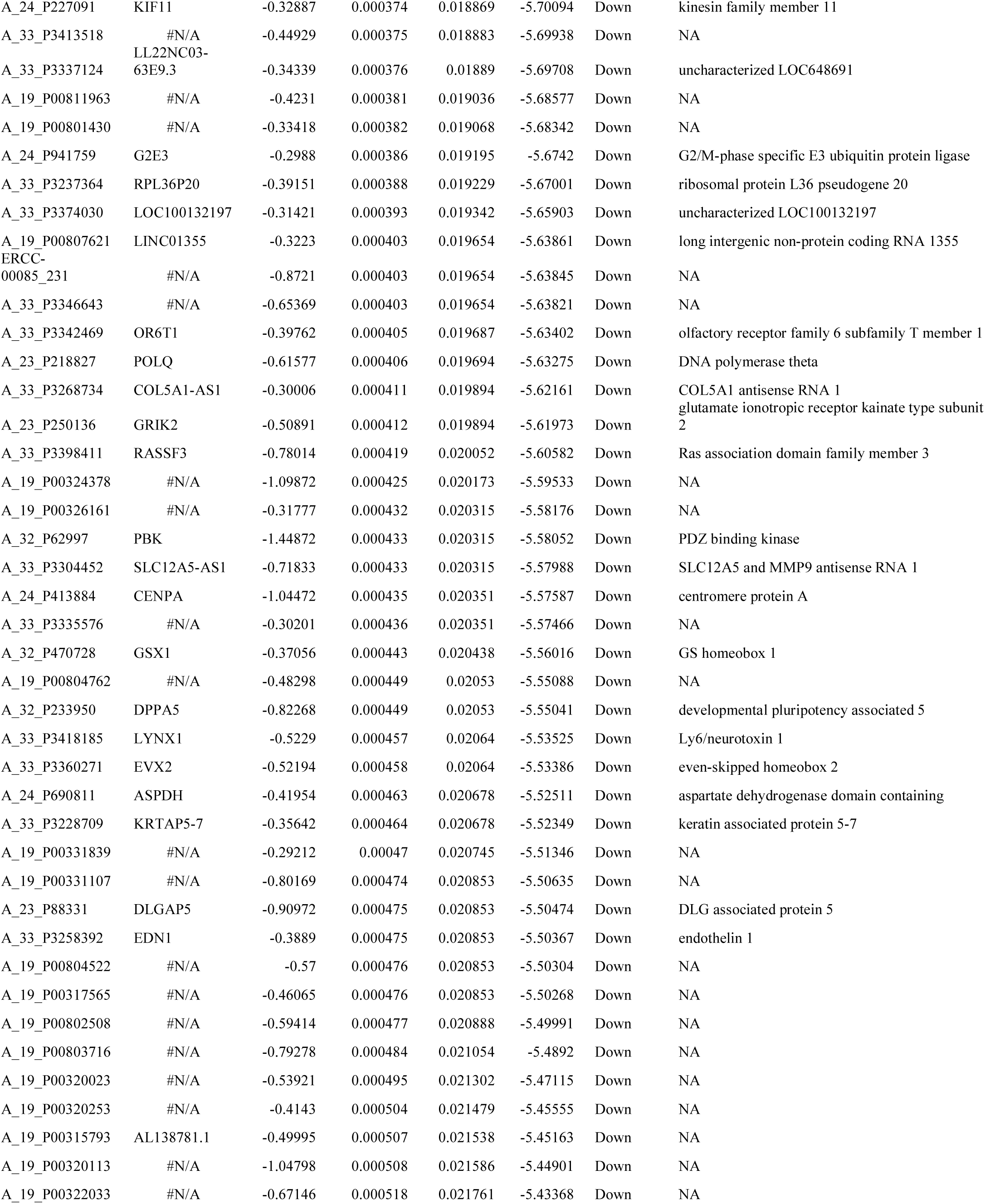
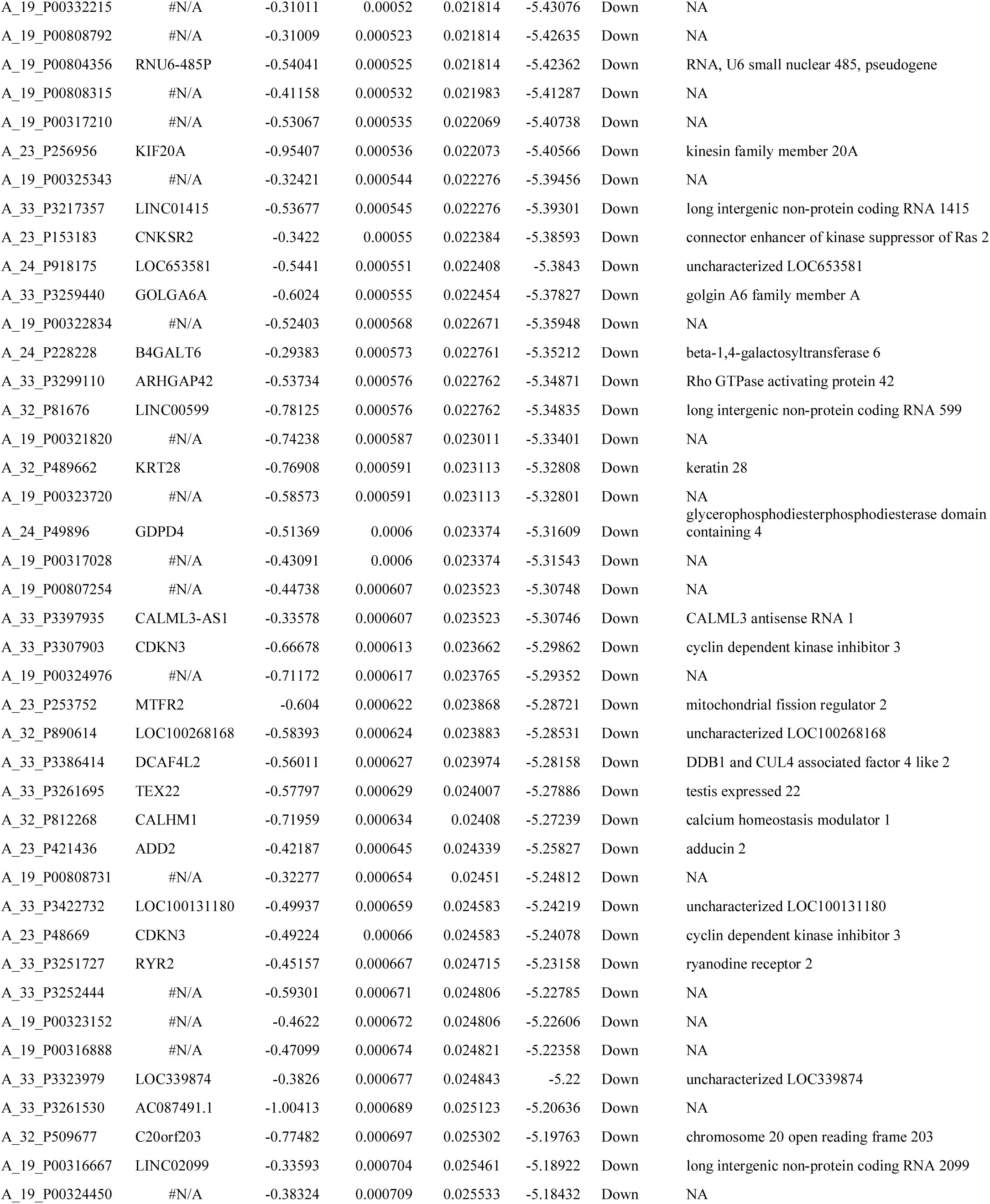
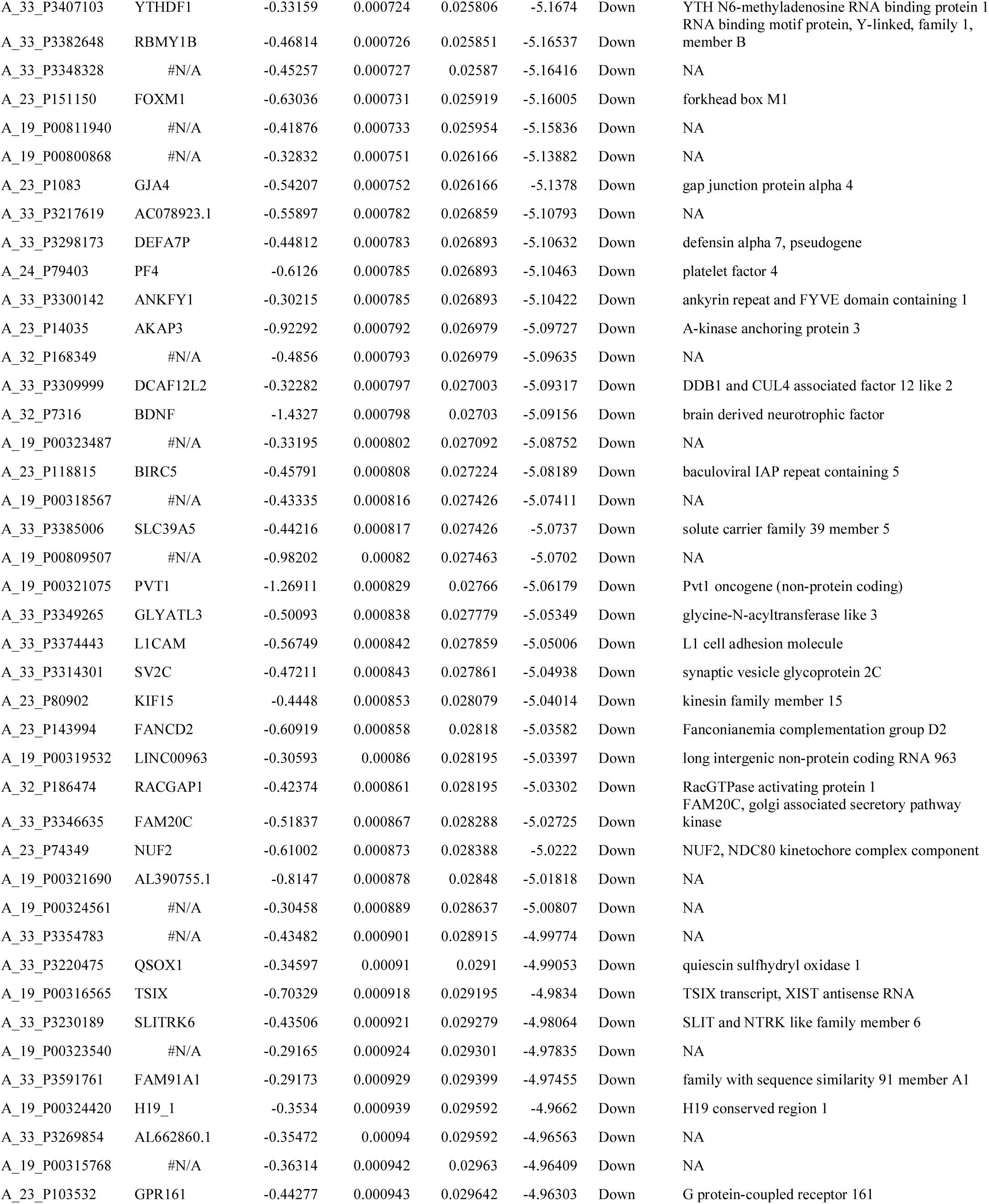
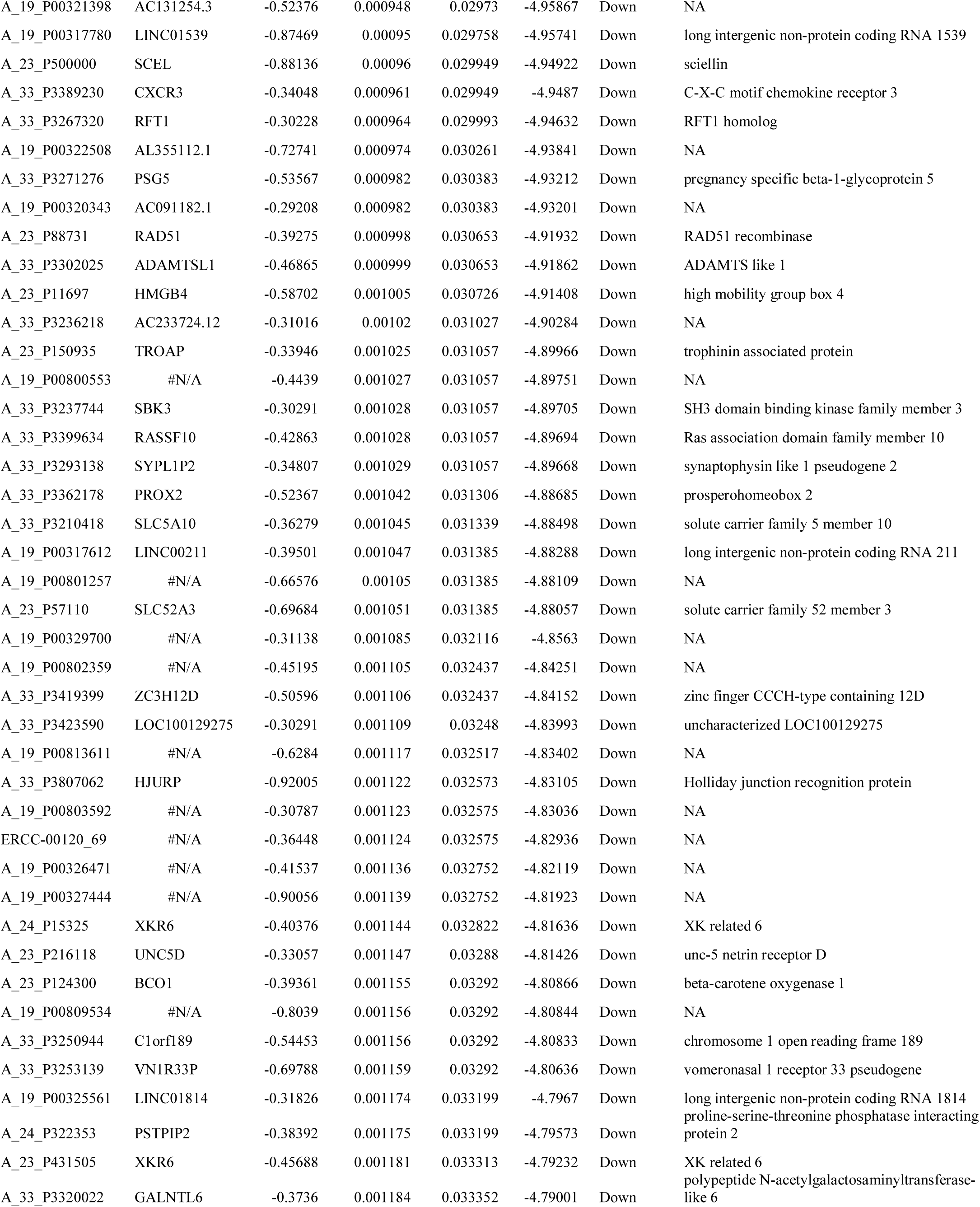
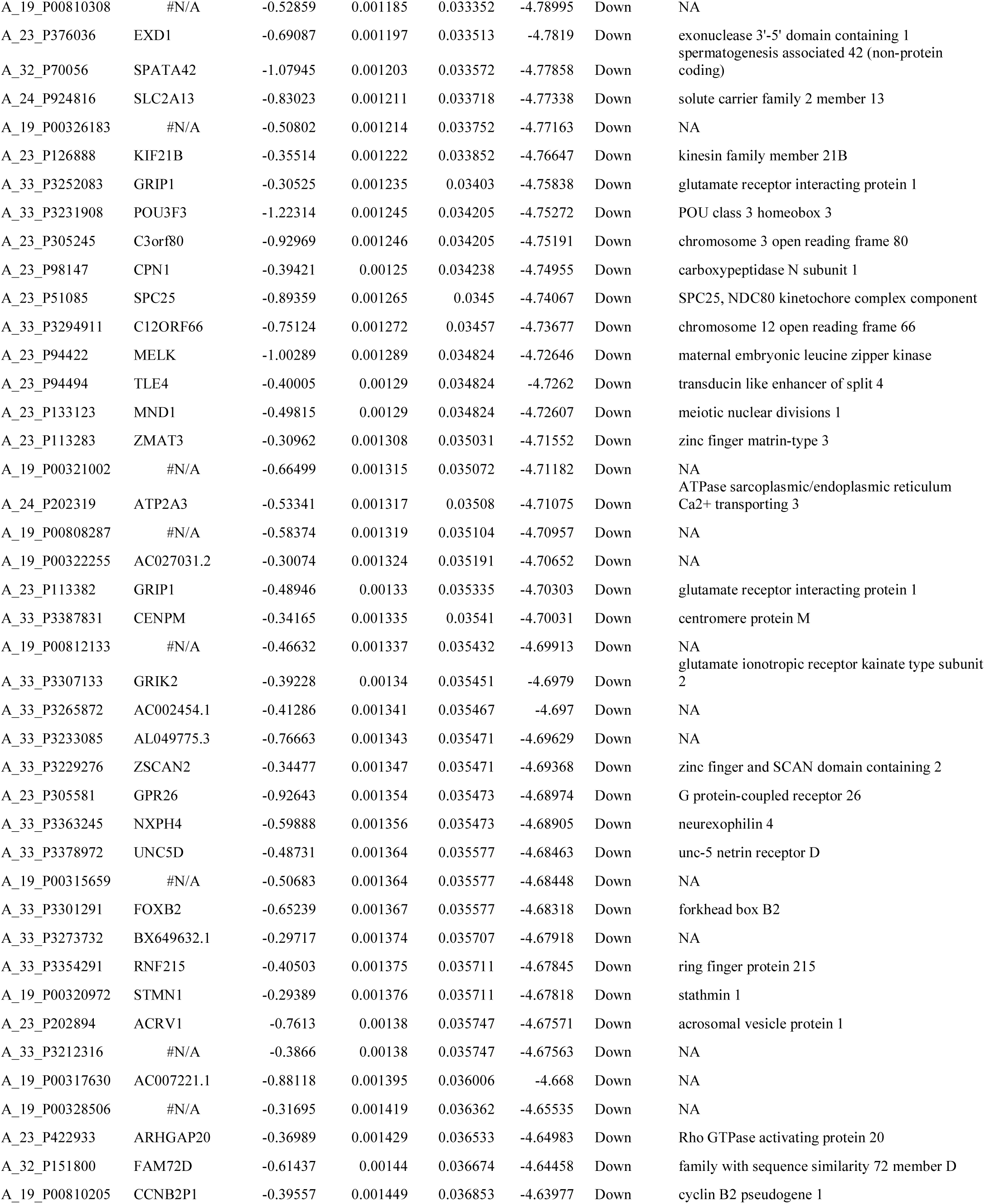
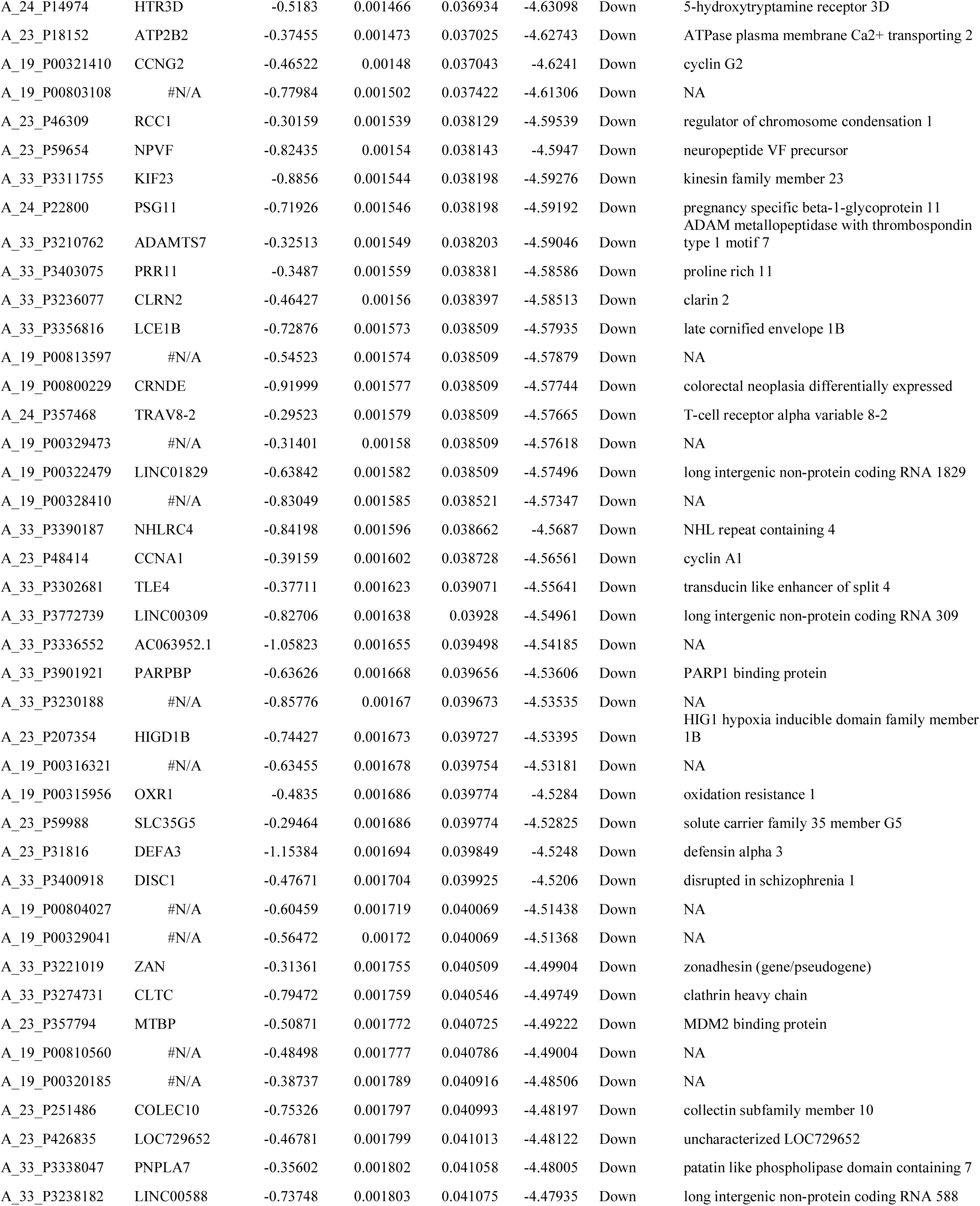
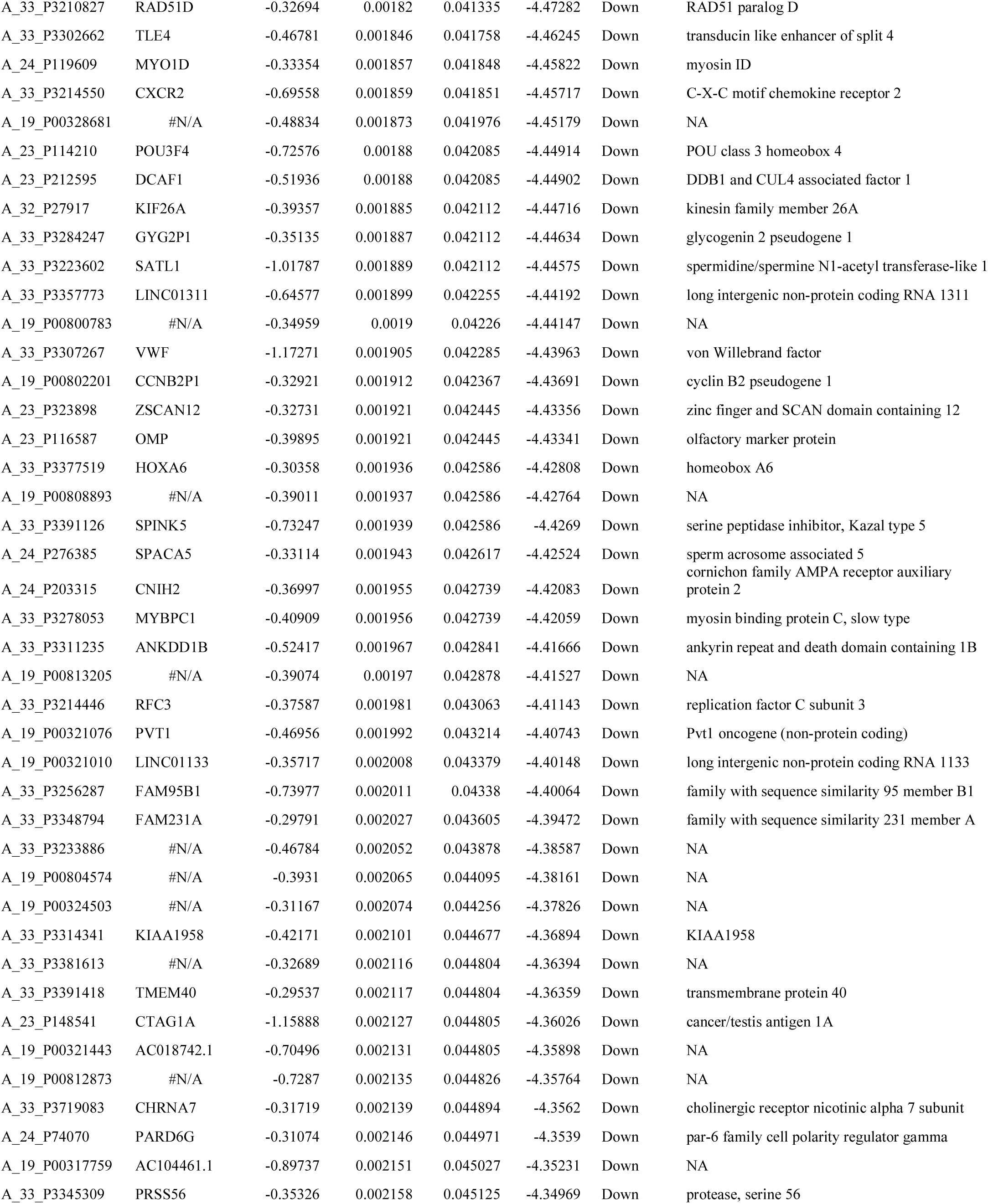
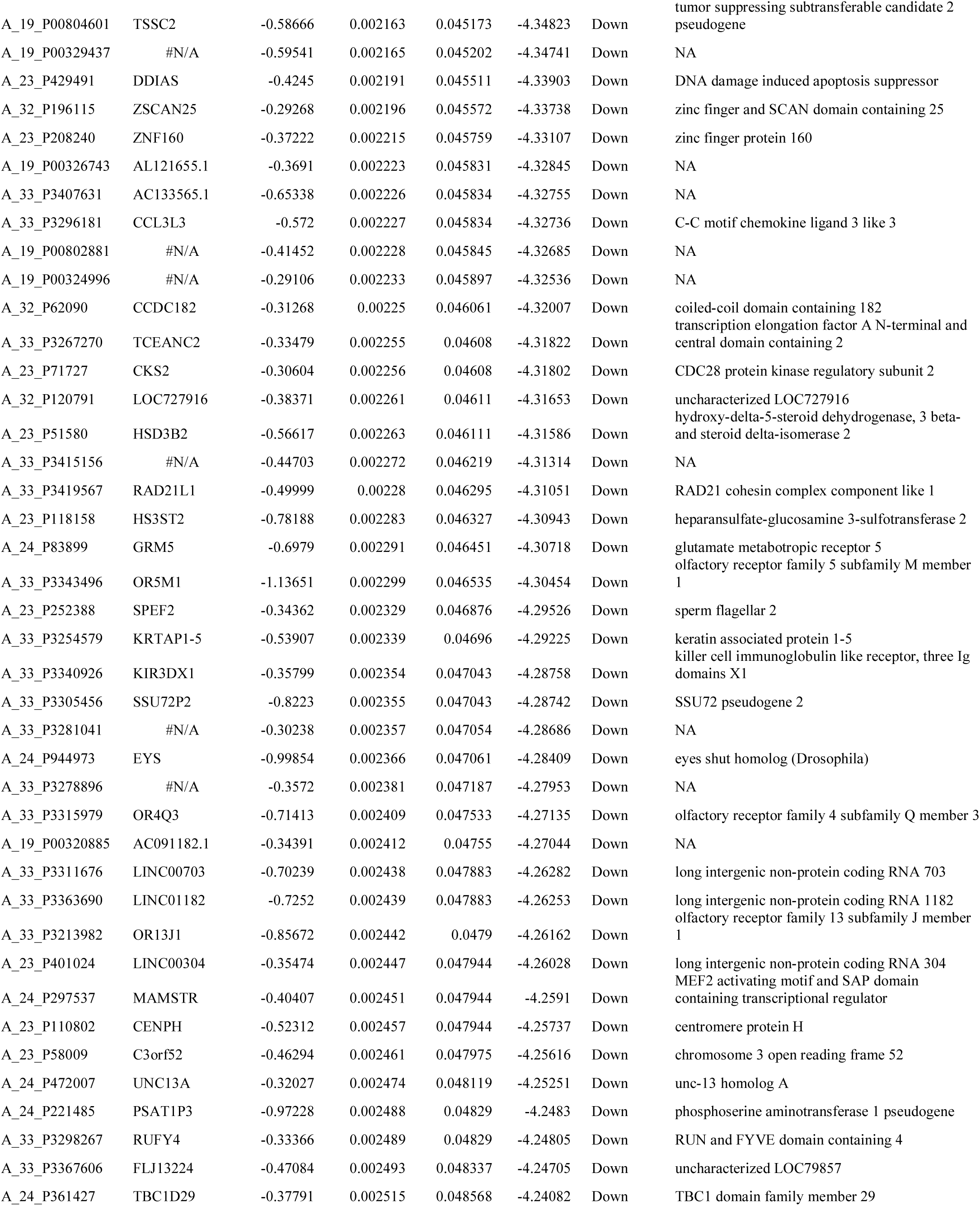
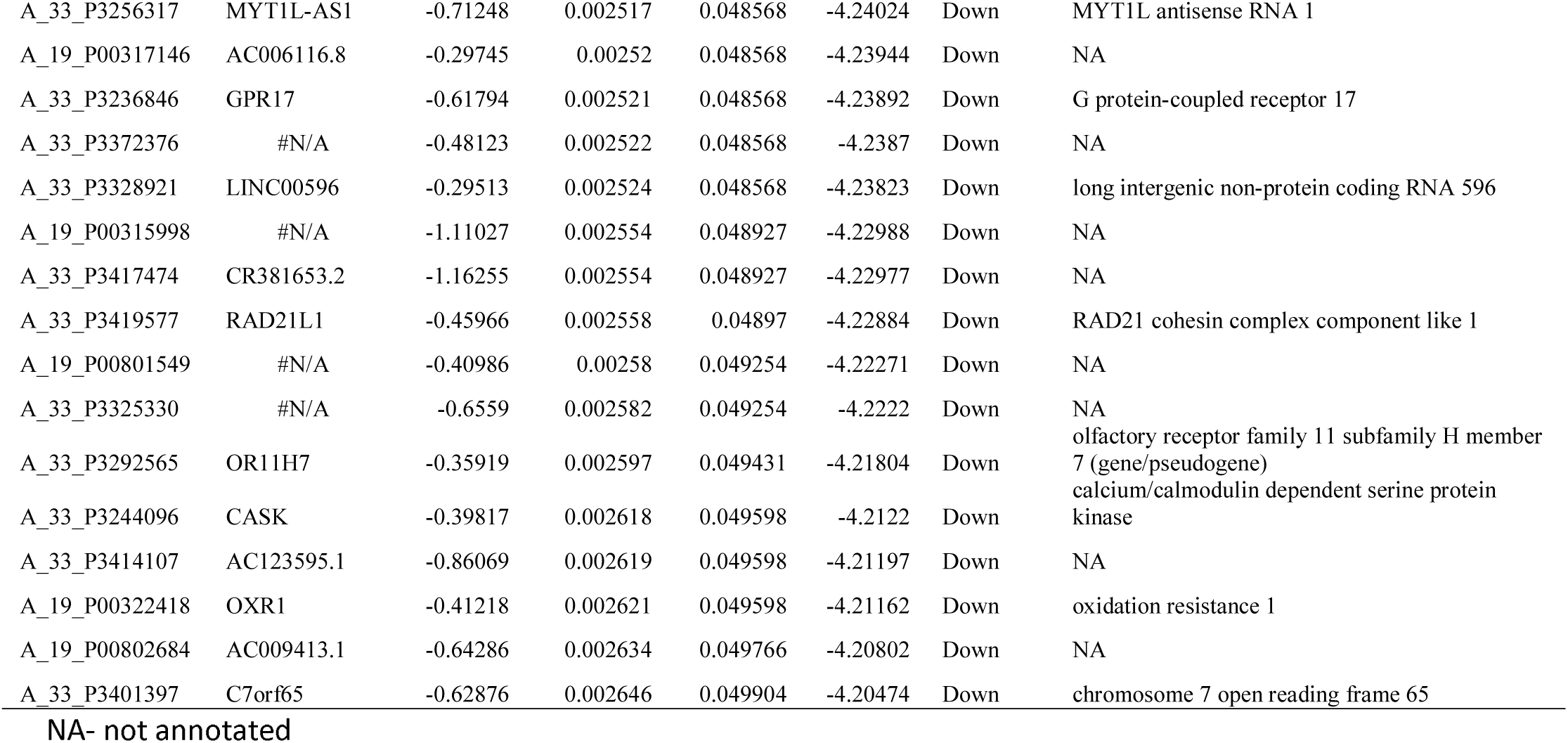
The statistical metrics for key differentially expressed genes (DEGs)

### Pathway enrichment analysis of DEGs

Several significant enriched pathways are acquired through BIOCYC, KEGG, PID, REACTOME, GenMAPP, MSigDB C2 BIOCARTA, PantherDB, Pathway Ontology and SMPDB pathway enrichment analysis (Table 2 and Table 3). The top enriched pathways for up regulated genes included retinoate biosynthesis II, retinoate biosynthesis I, signaling pathways regulating pluripotency of stem cells, neuroactive ligand-receptor interaction, ALK2 signaling events, BMP receptor signaling, peptide hormone biosynthesis, glycoprotein hormones, tyrosine metabolism, androgen and estrogen metabolism, ensemble of genes encoding extracellular matrix and extracellular matrix-associated proteins, genes encoding secreted soluble factors, adenine and hypoxanthine salvage pathway, 5-Hydroxytryptamine biosynthesis, melanocortin system, androgen and estrogen metabolic, tryptophan metabolism and xanthine dehydrogenase deficiency (Xanthinuria). Meanwhile, down regulated DEGs strikingly enriched in vitamin D3 biosynthesis, cell cycle, pancreatic secretion, aurora B signaling, FOXM1 transcription factor network, mitotic prometaphase, resolution of sister chromatid cohesion, role of ran in mitotic spindle regulation, Eph kinases and ephrins support platelet aggregation, inflammation mediated by chemokine and cytokine signaling pathway, O-Glycans biosynthetic, ganglioside biosynthetic, eptifibatide pathway and ticlopidine pathway.

**Table 2.**
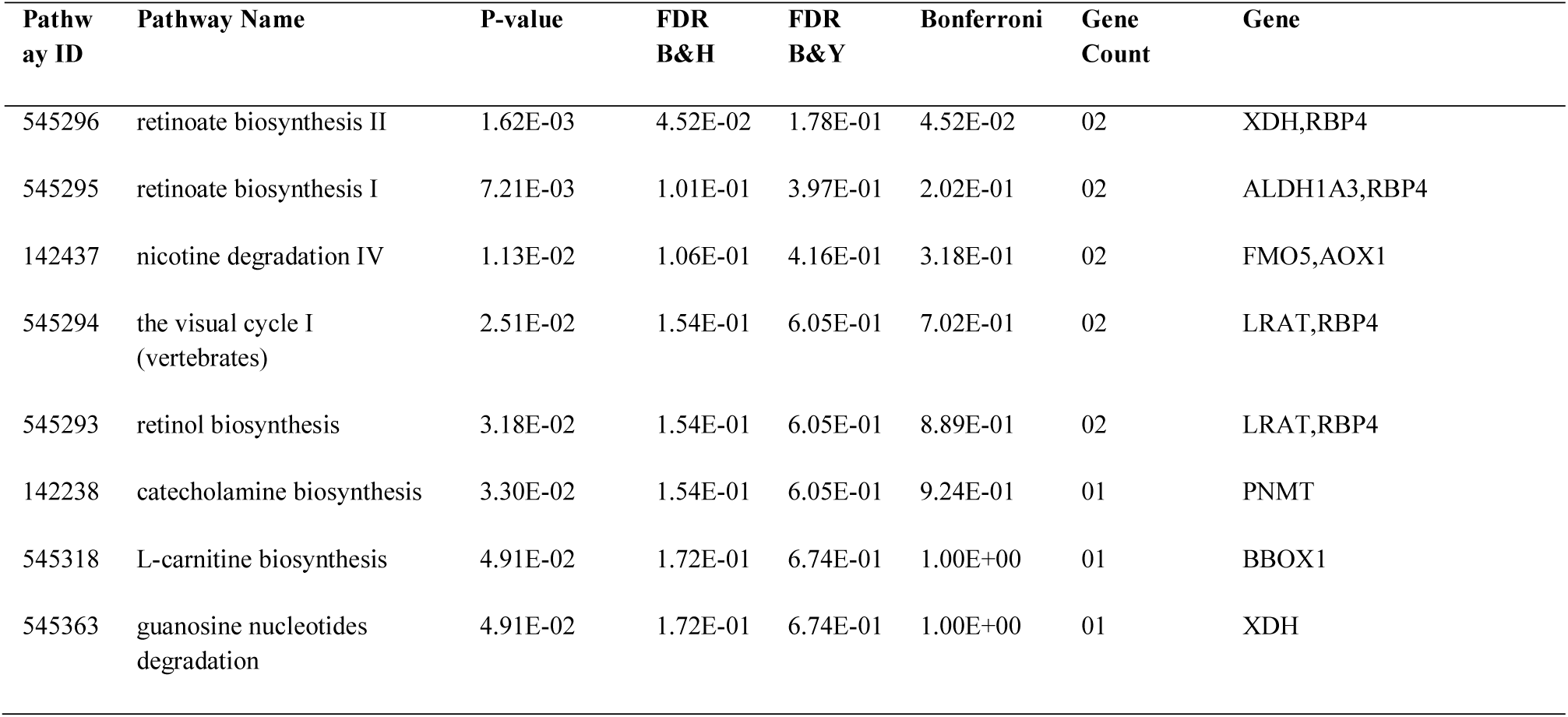
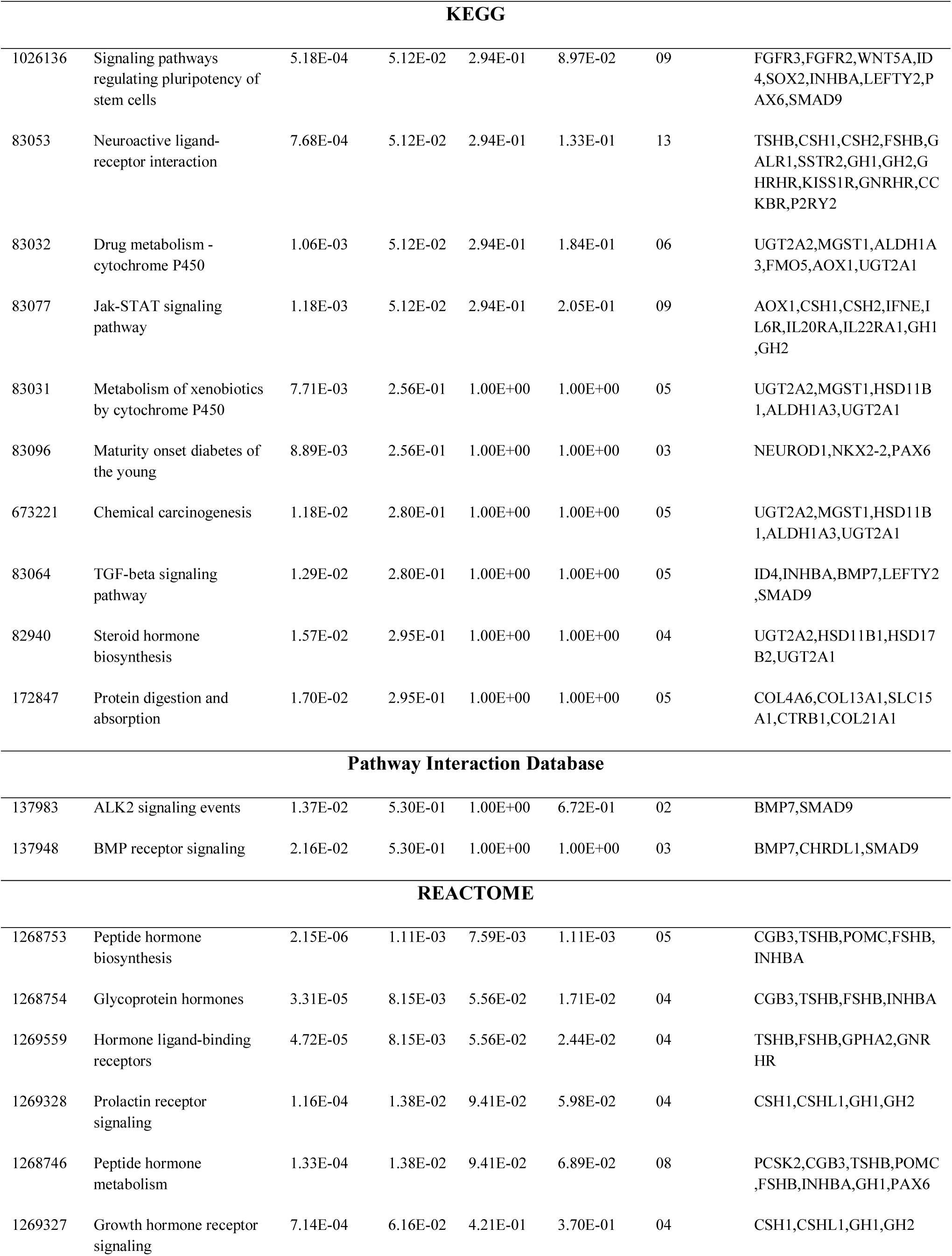
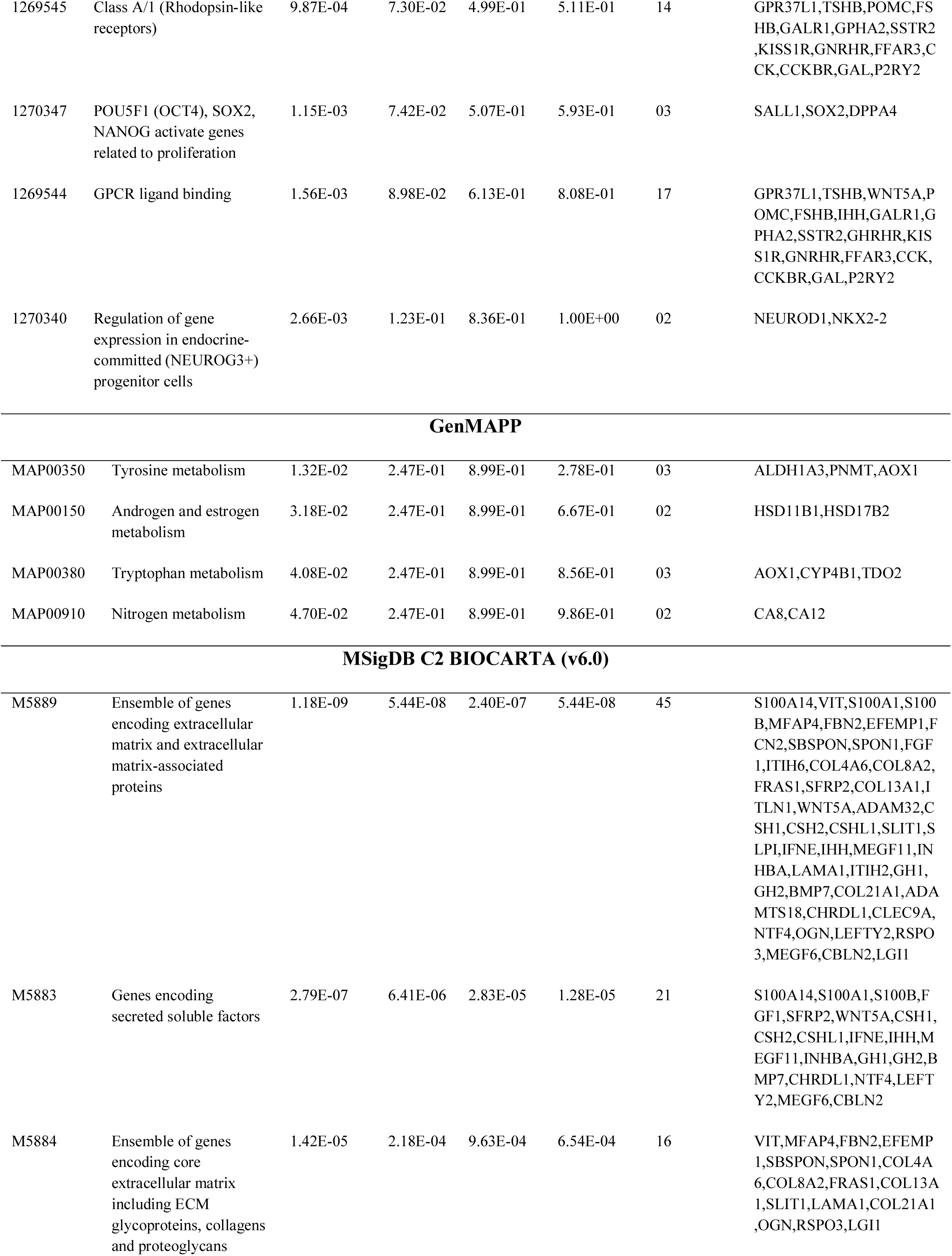
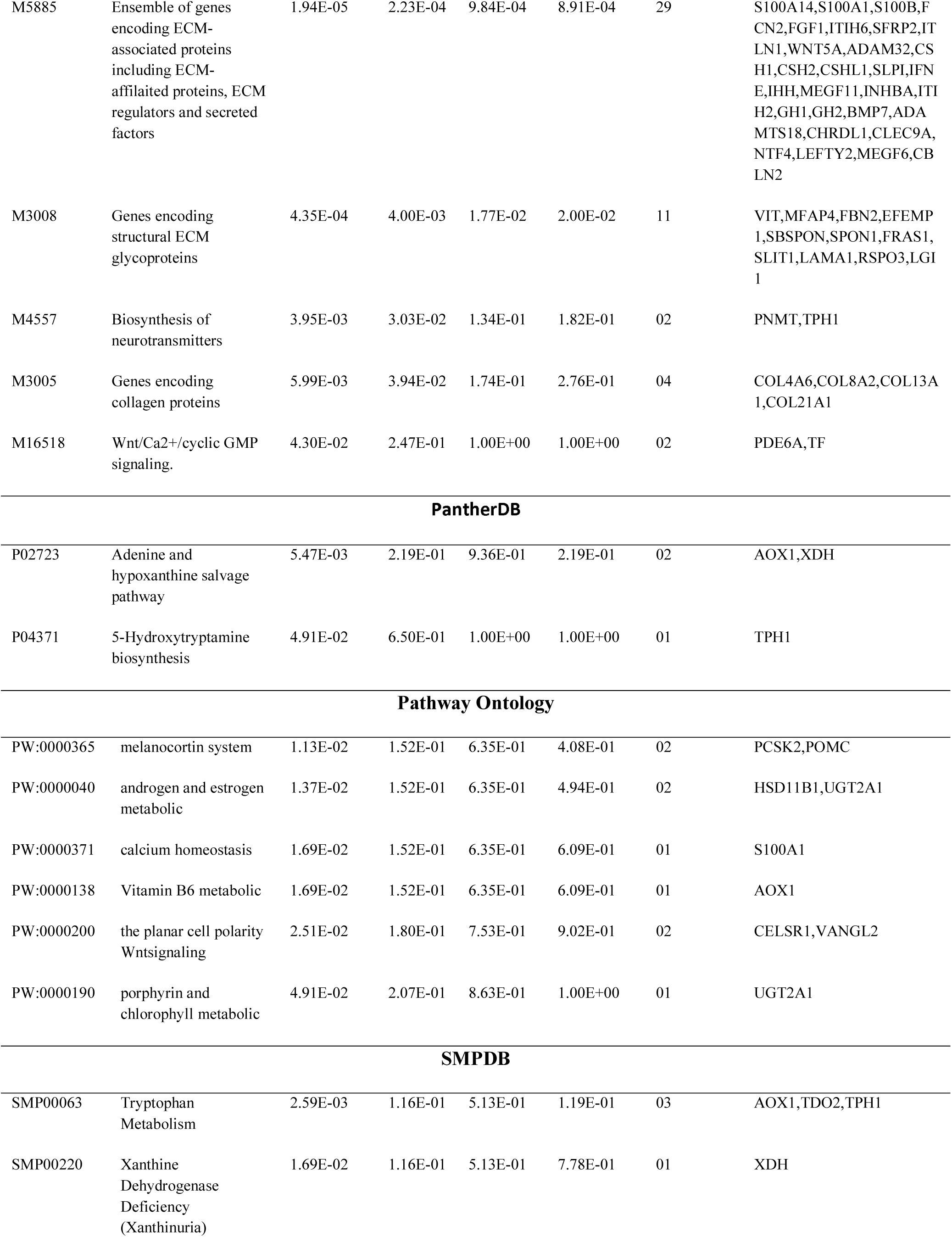
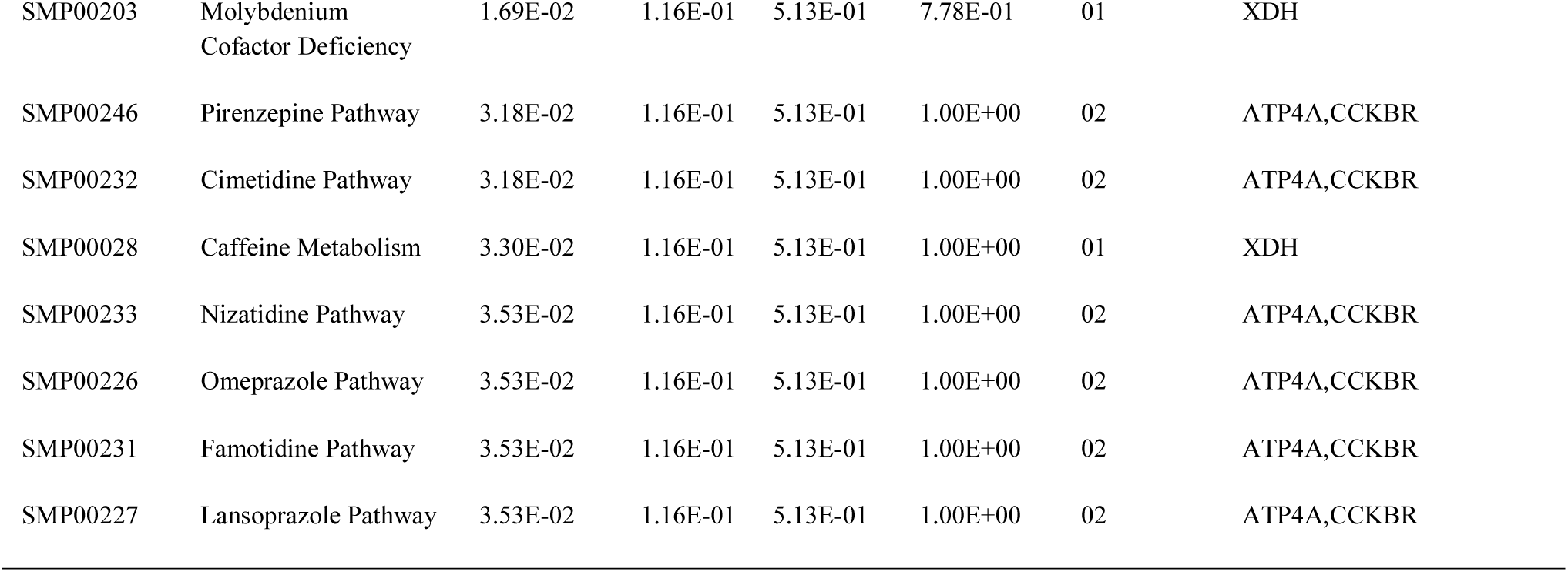
The enriched pathway terms of the up-regulated differentially expressed genes

**Table 3.**
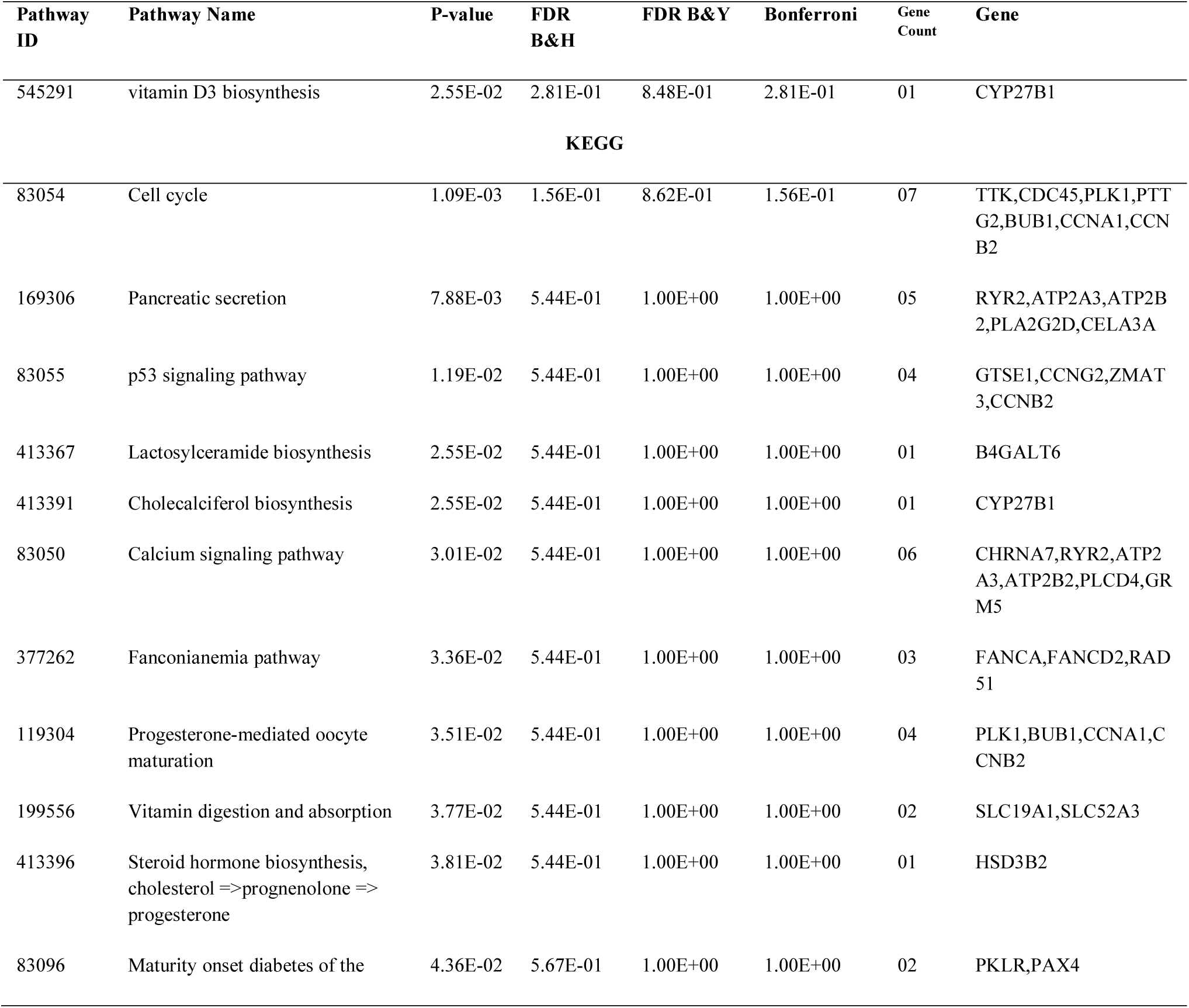
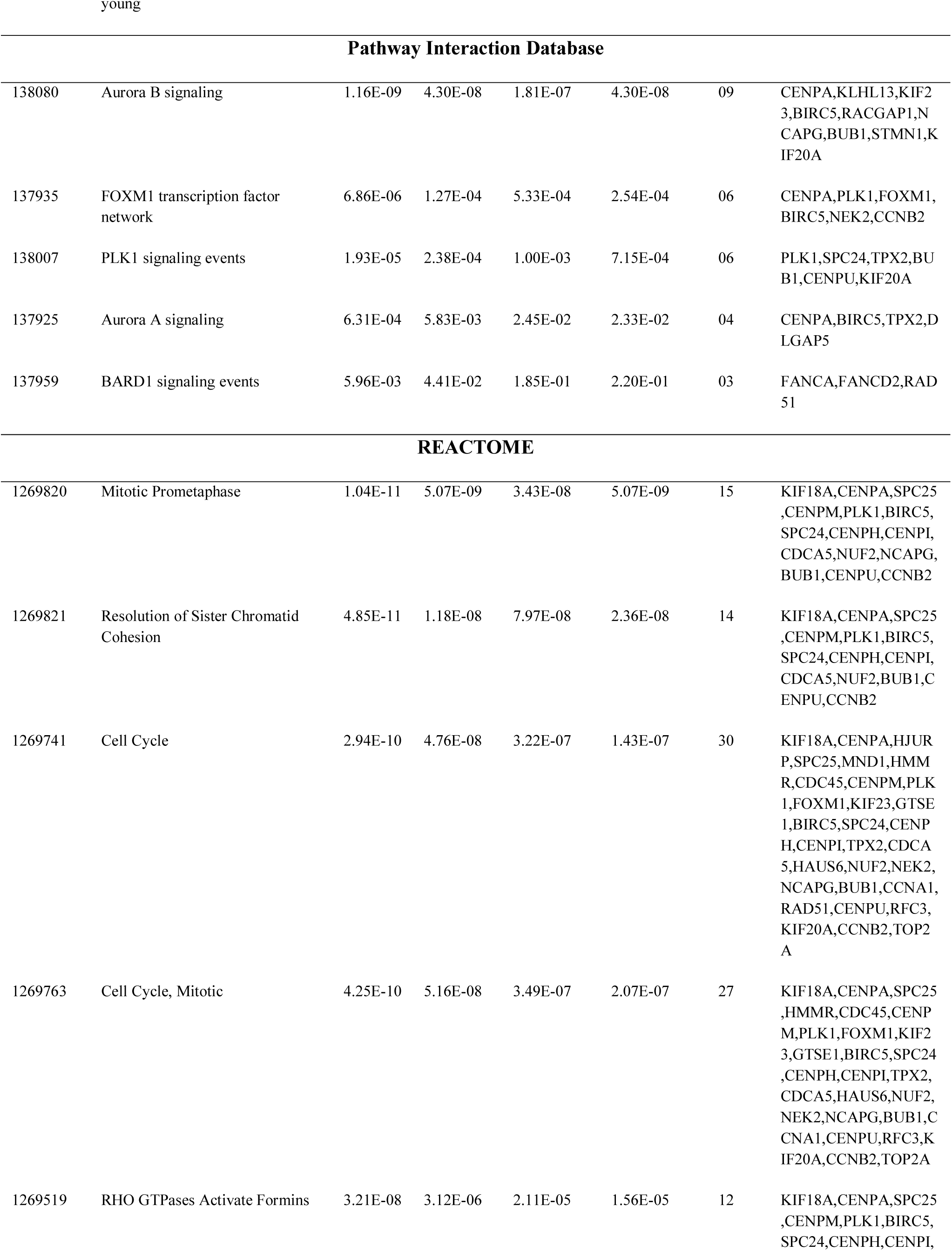
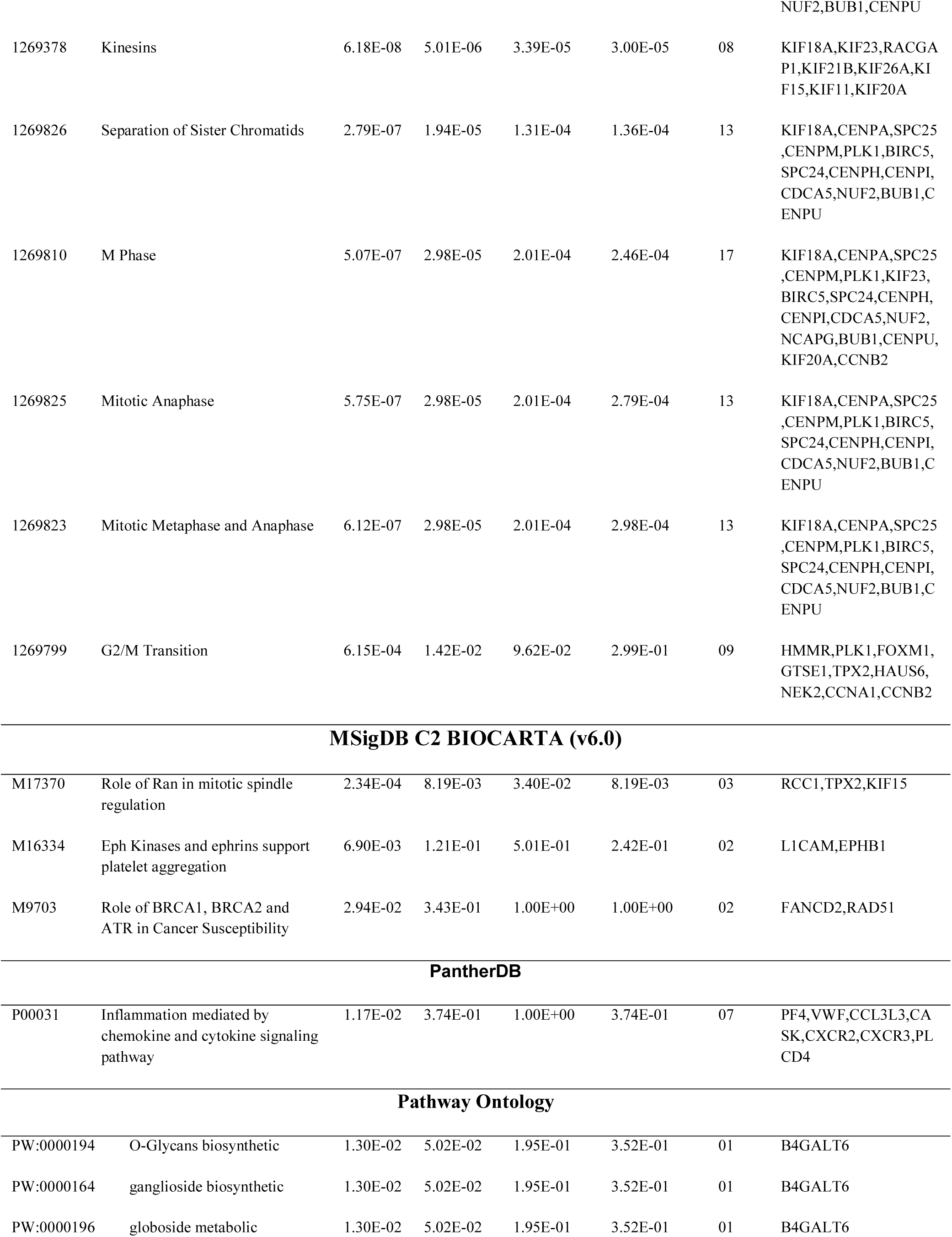
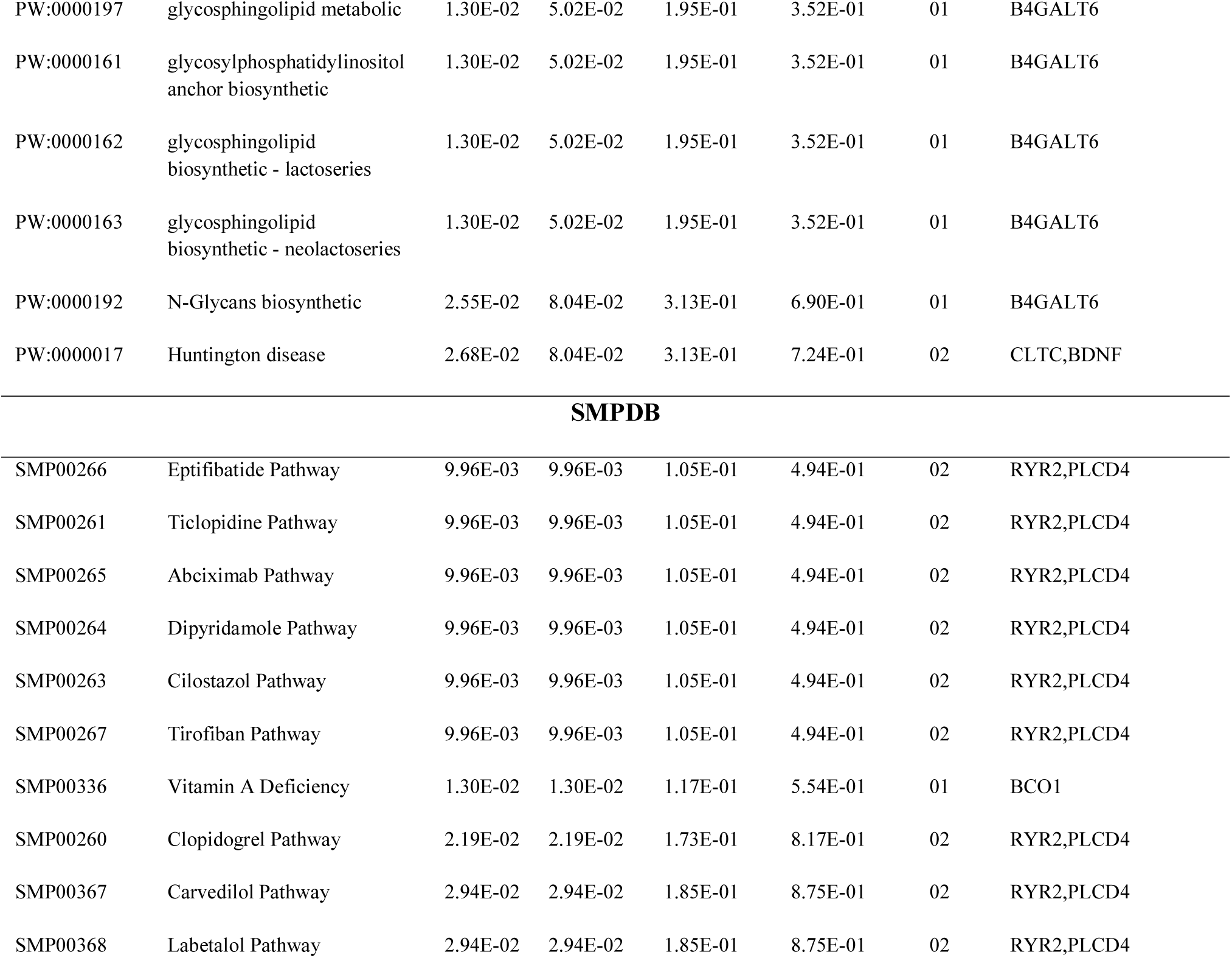
The enriched pathway terms of the down-regulated differentially expressed genes

### Gene Ontology (GO) enrichment analysis of DEGs

All significant DEGs were divided into up regulated genes and down regulated genes. GO categories analyses are conducted for these 2 lists of genes, respectively. Results of GO categories are presented by 3 functional groups, which are group BP, CC, and MF (Table 4 and Table 5). In group BP, up and down regulated DEGs are significantly enriched in sensory organ morphogenesis, embryonic organ morphogenesis, nuclear division and organelle fission. For group CC, up and down regulated DEGs mainly enriched in extracellular matrix and extracellular space, condensed chromosome and kinetochore. In addition, GO results of group MF showed that up and down regulated DEGs mainly enriched in hormone activity, signaling receptor binding, microtubule binding and microtubule motor activity.

**Table 4.**
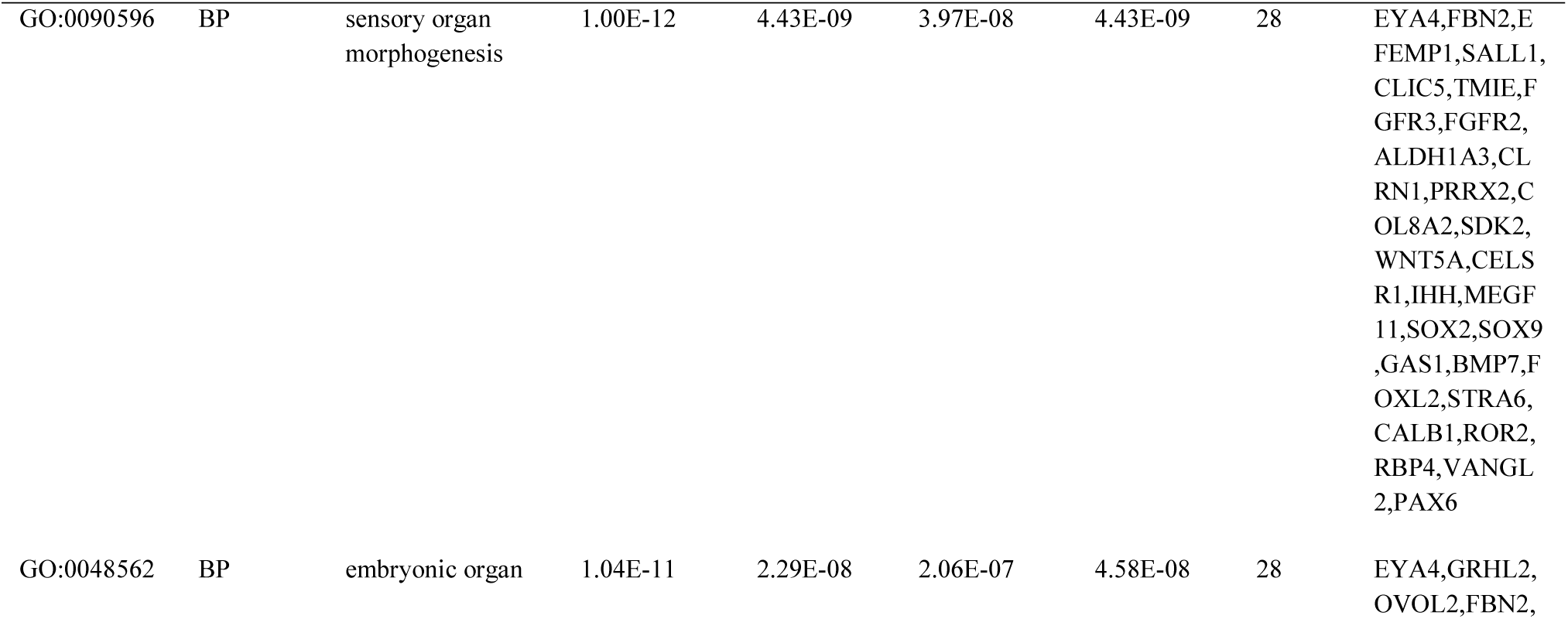
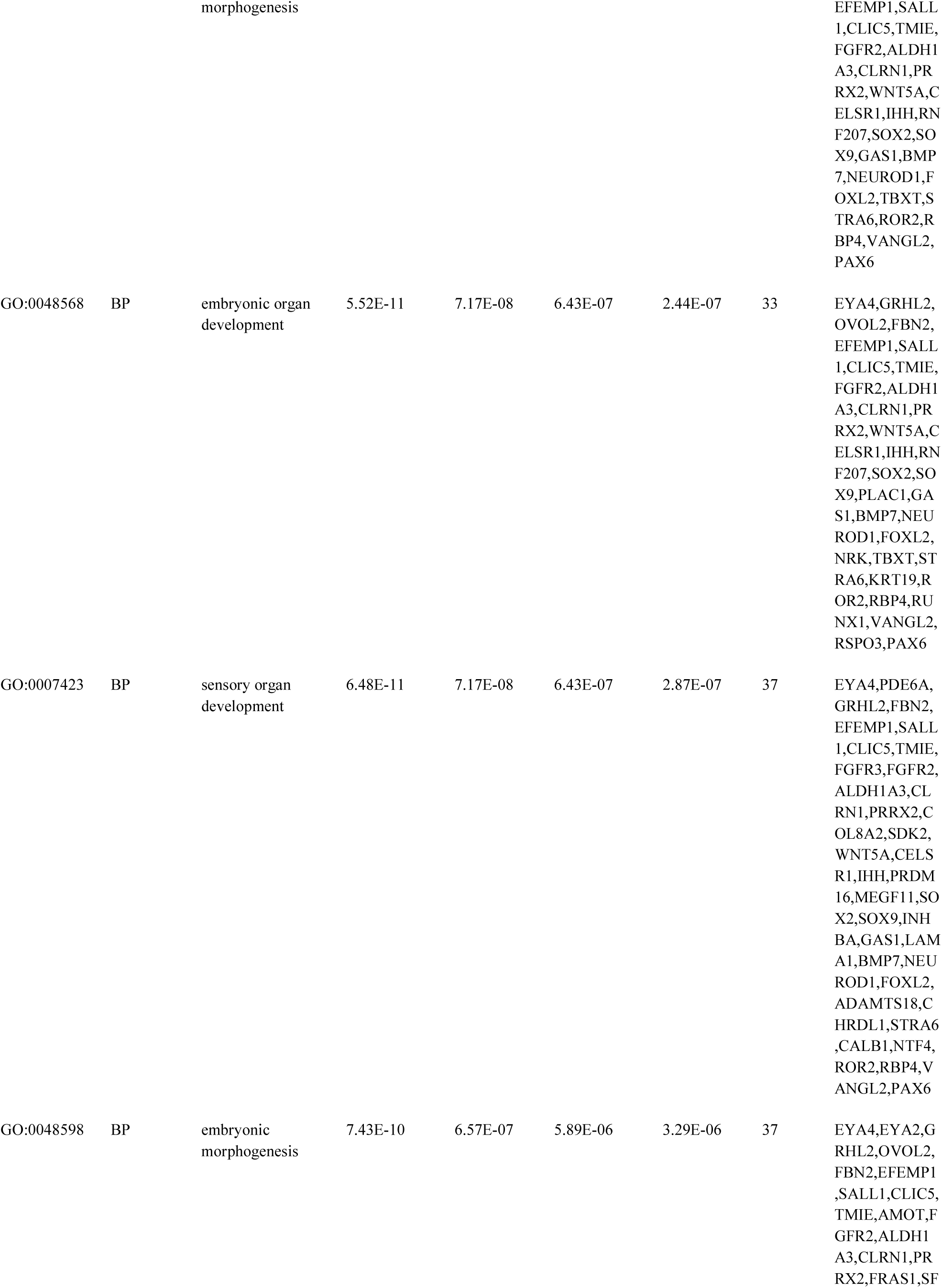
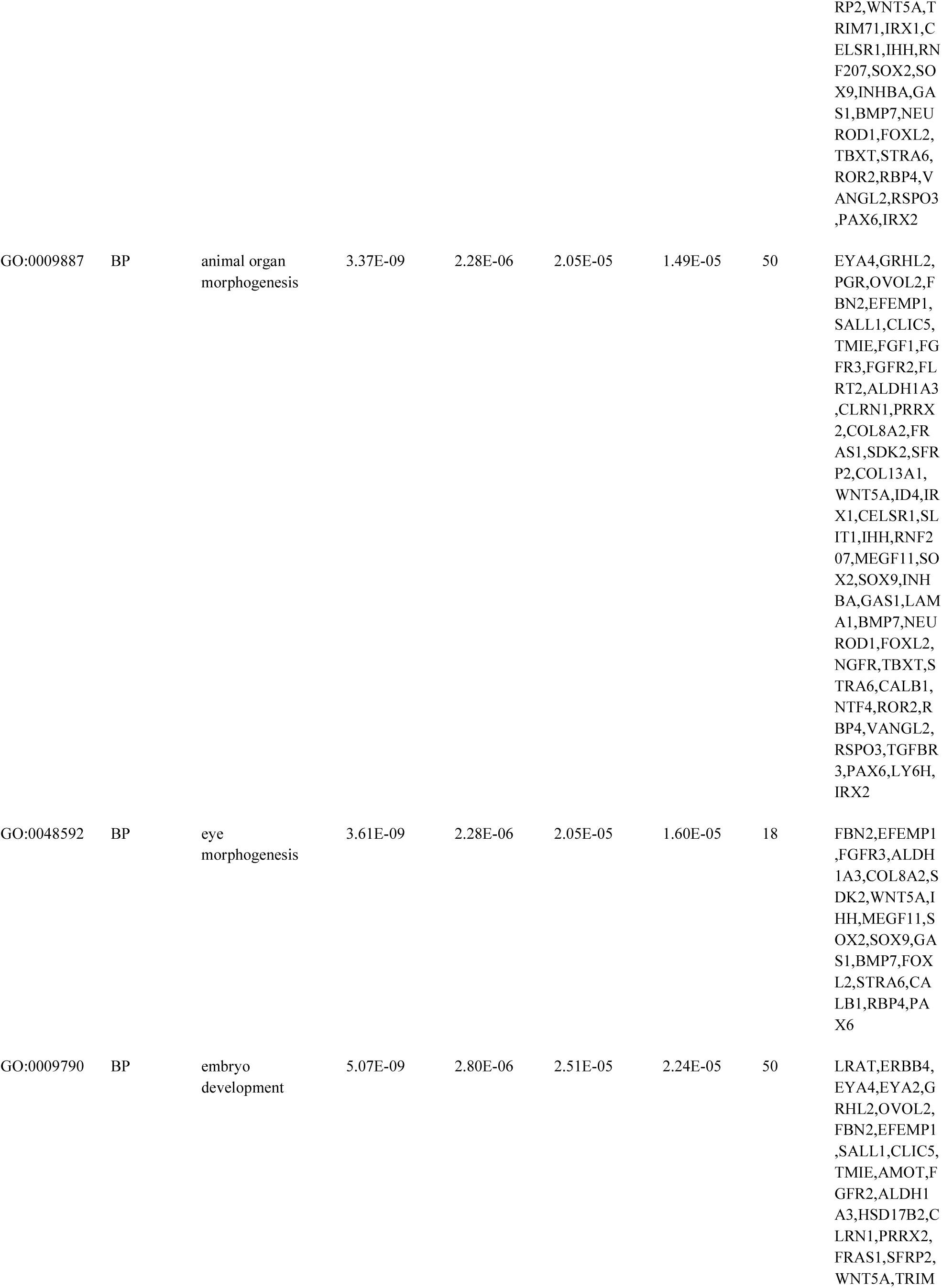
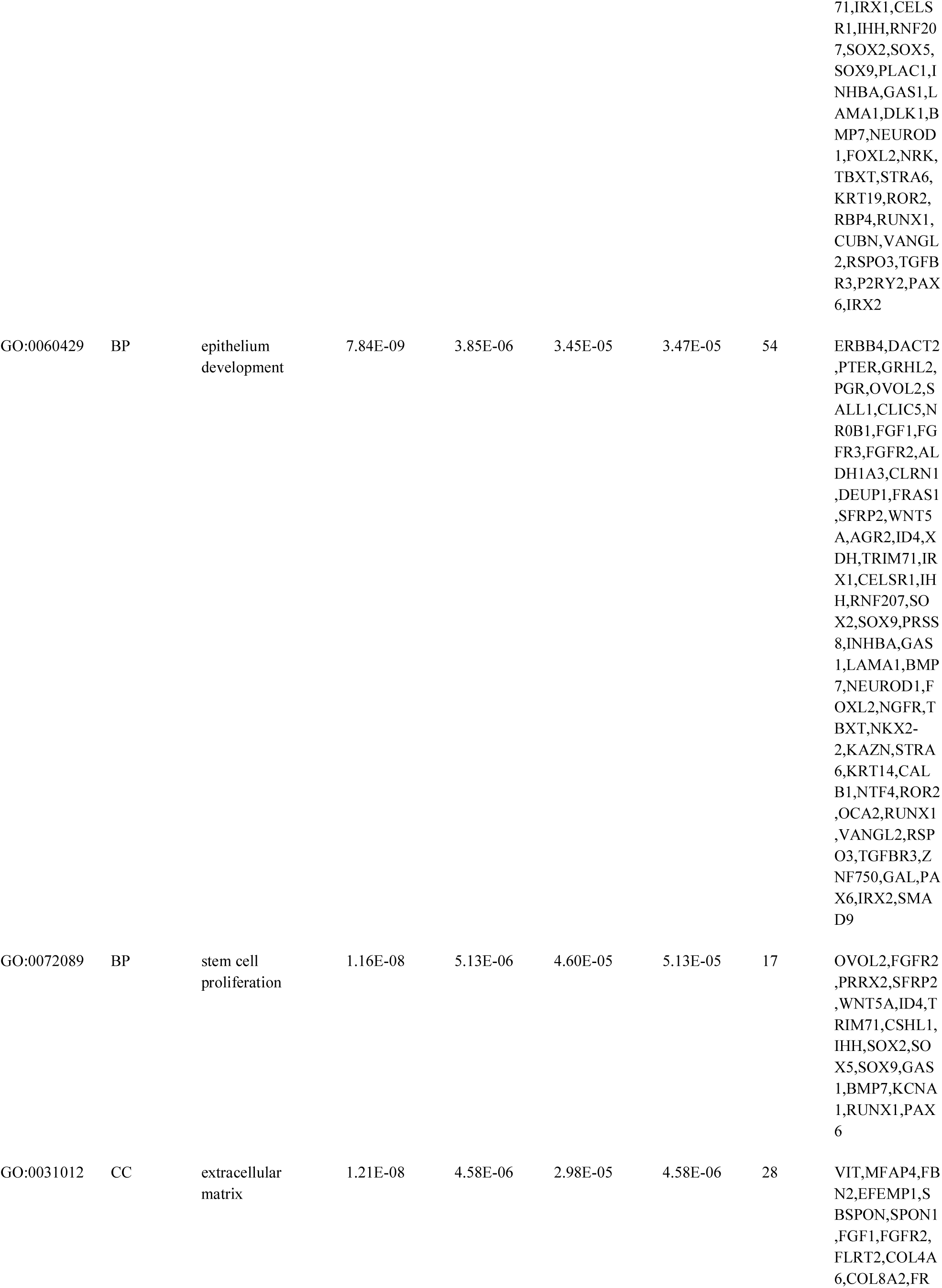
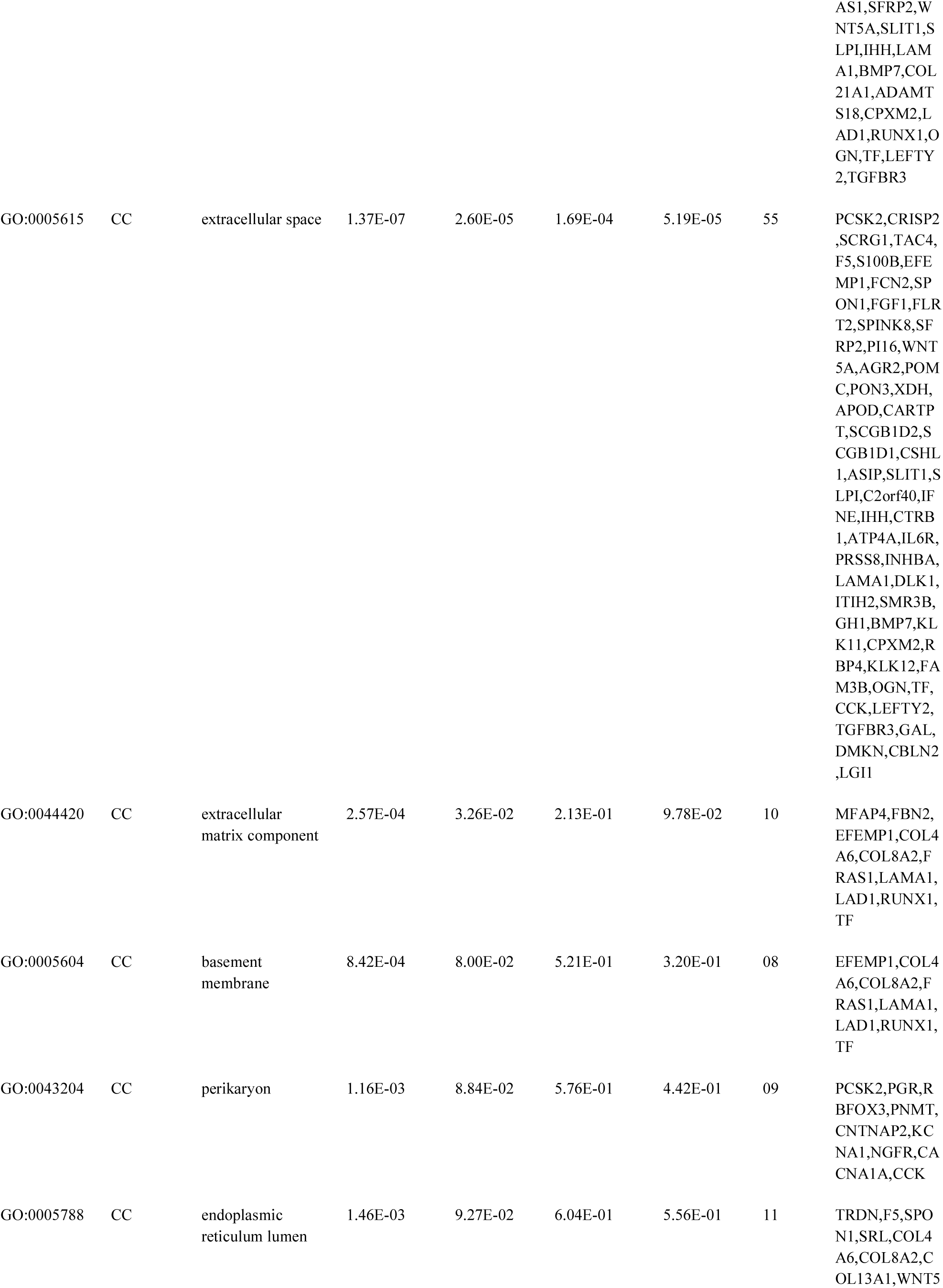
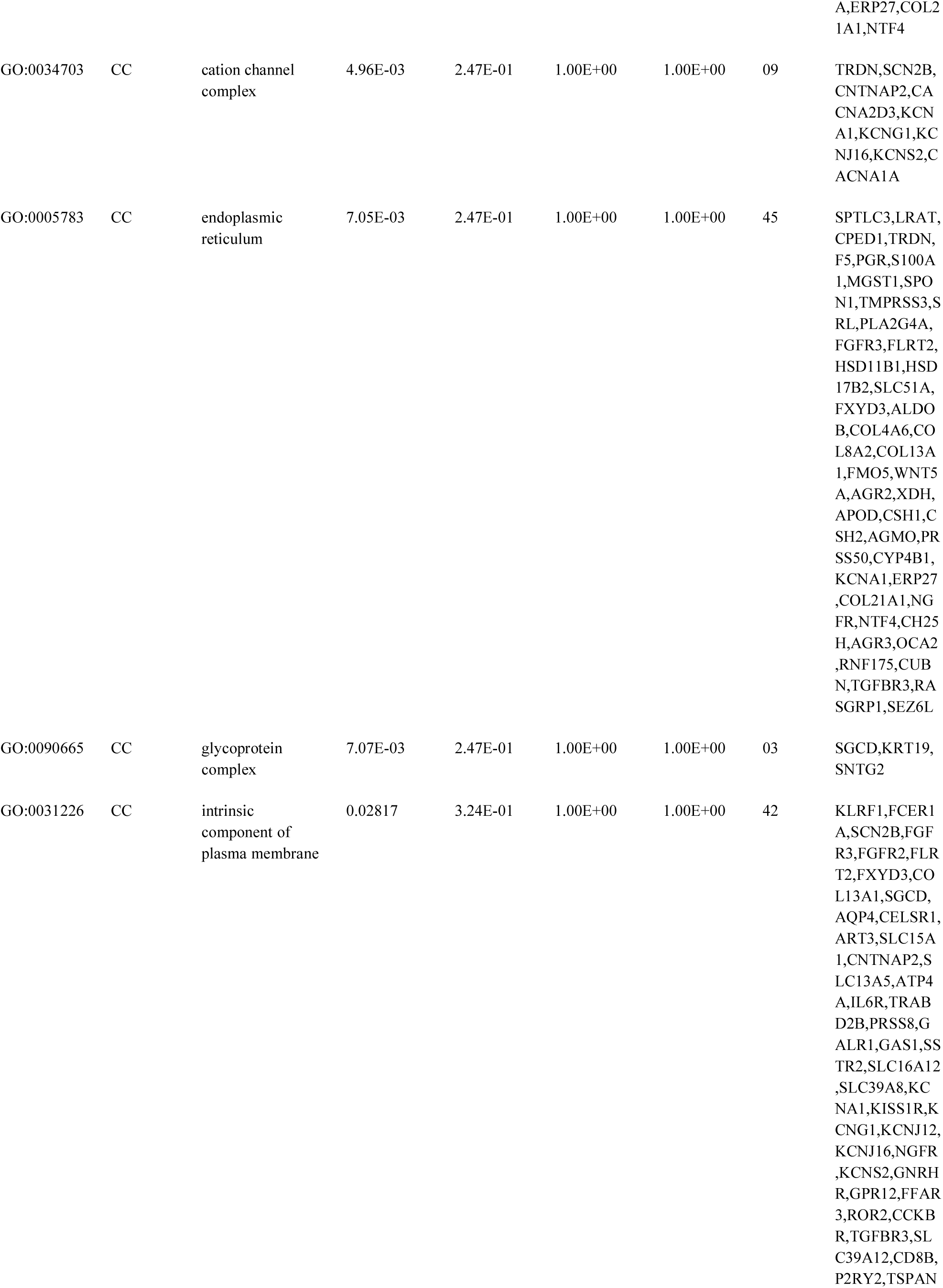
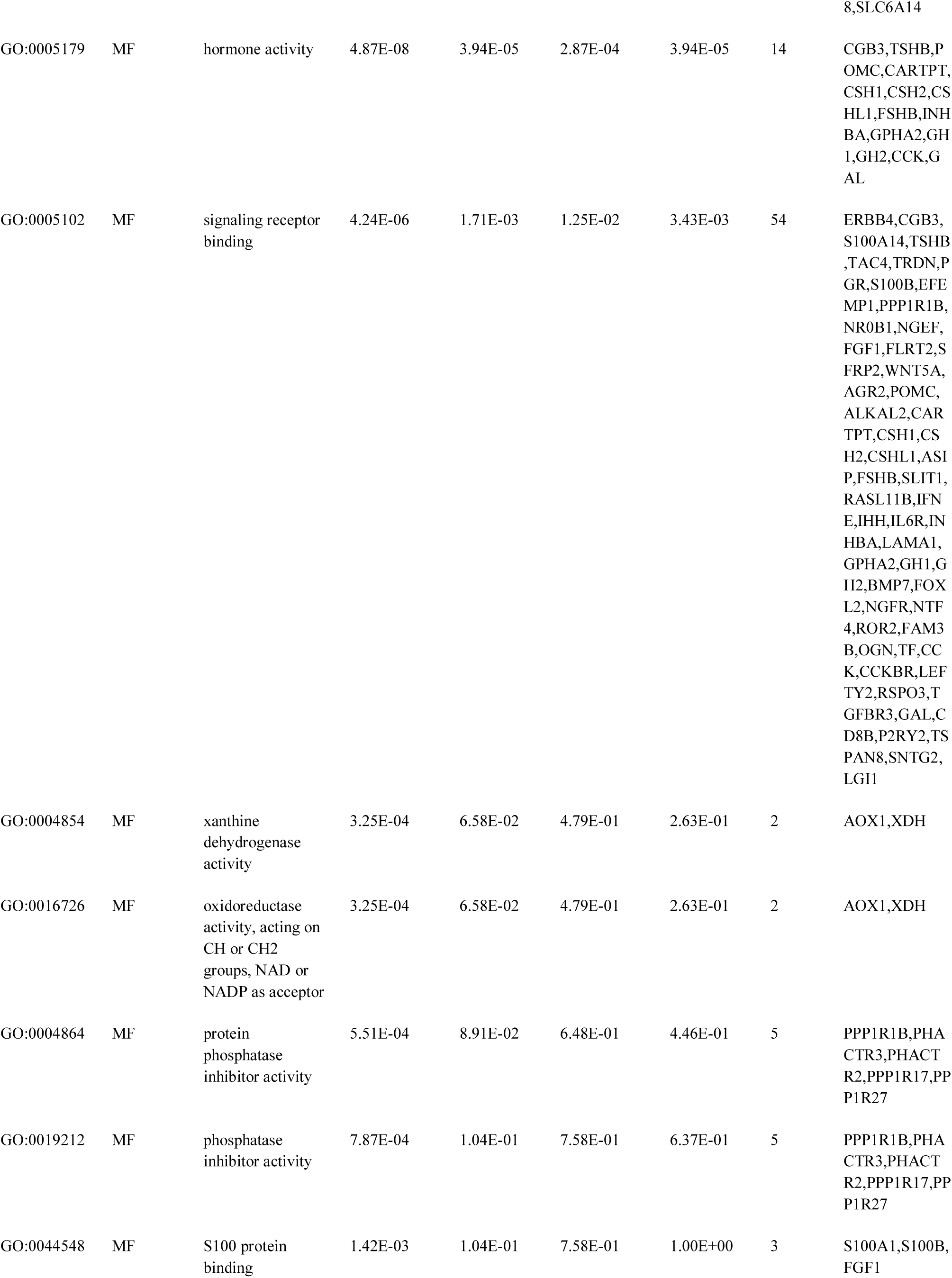
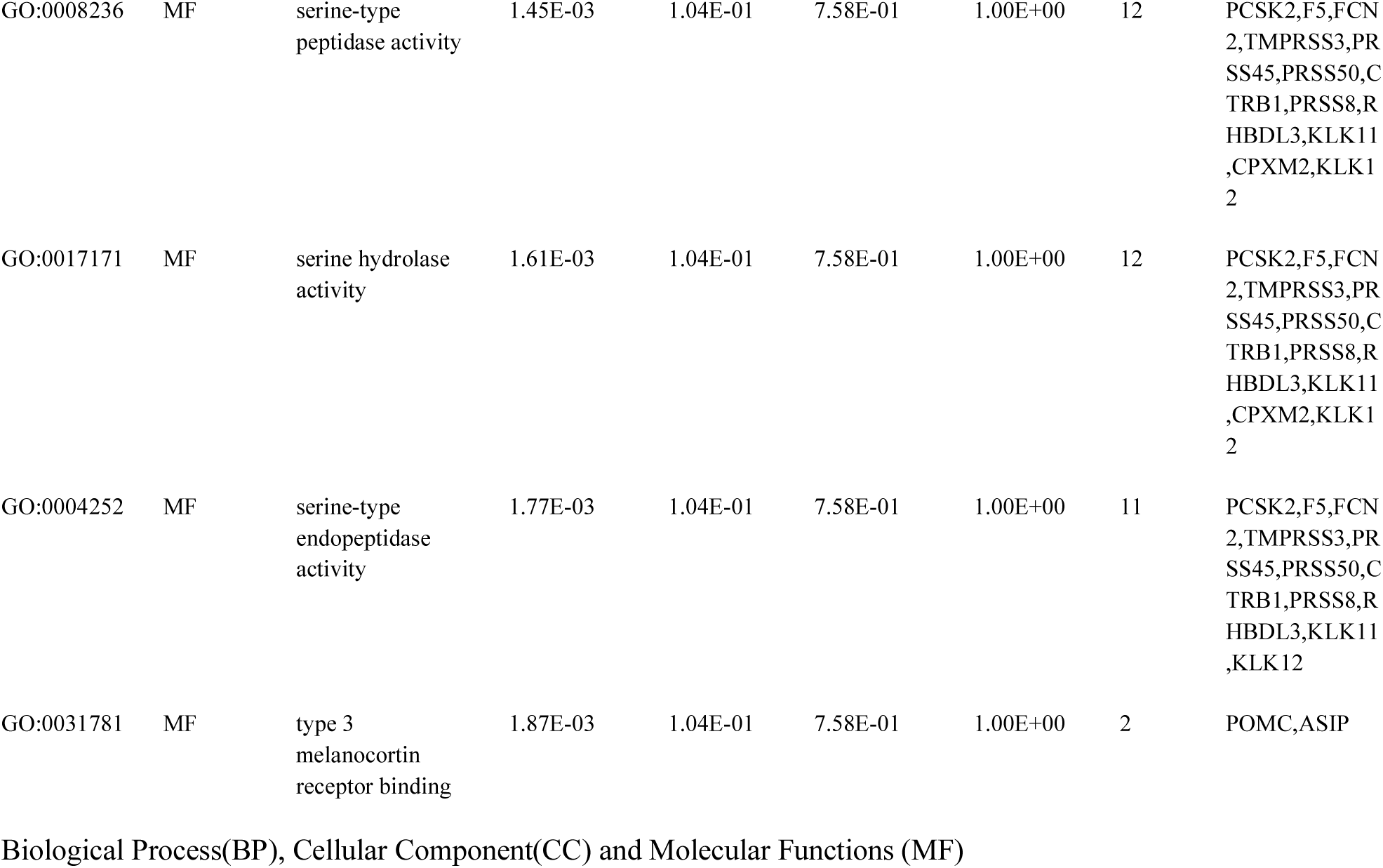
The enriched GO terms of the up-regulated differentially expressed genes

**Table 5.**
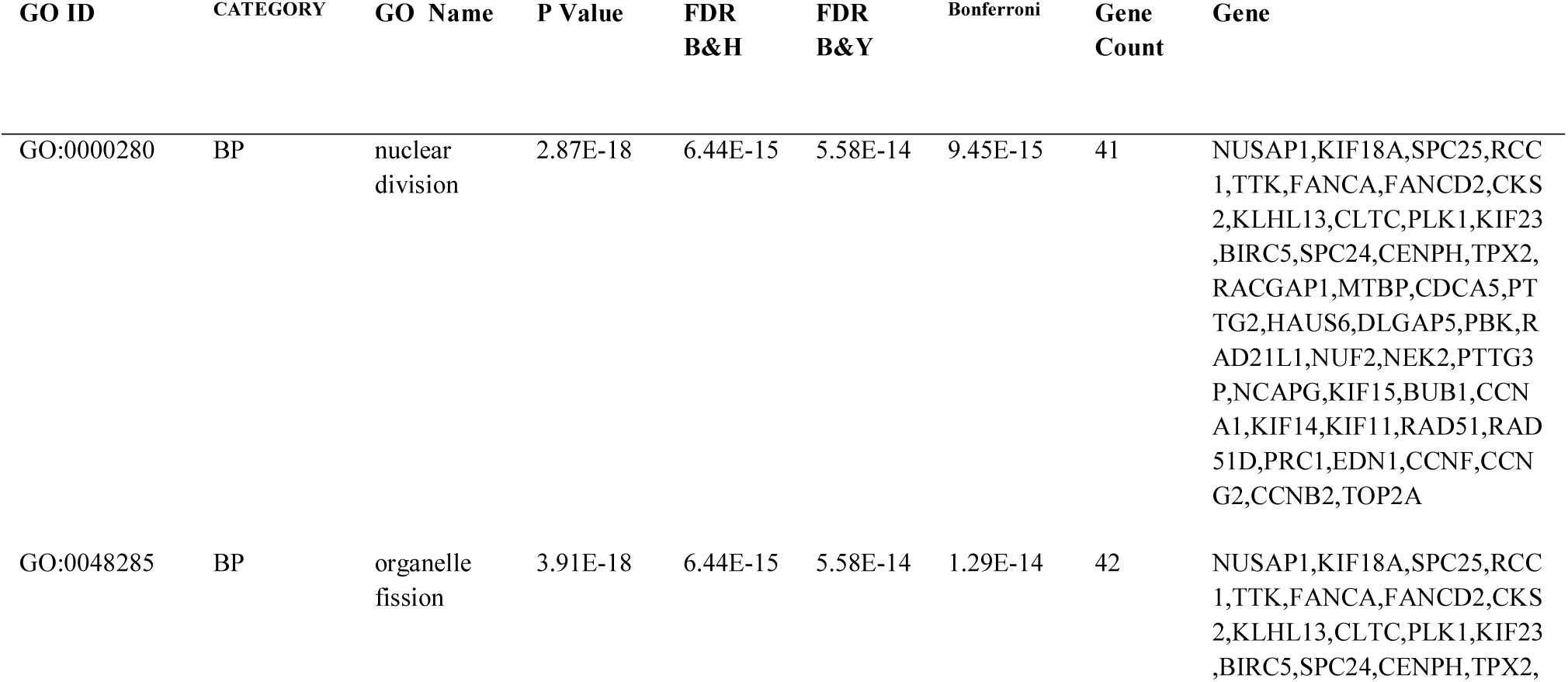
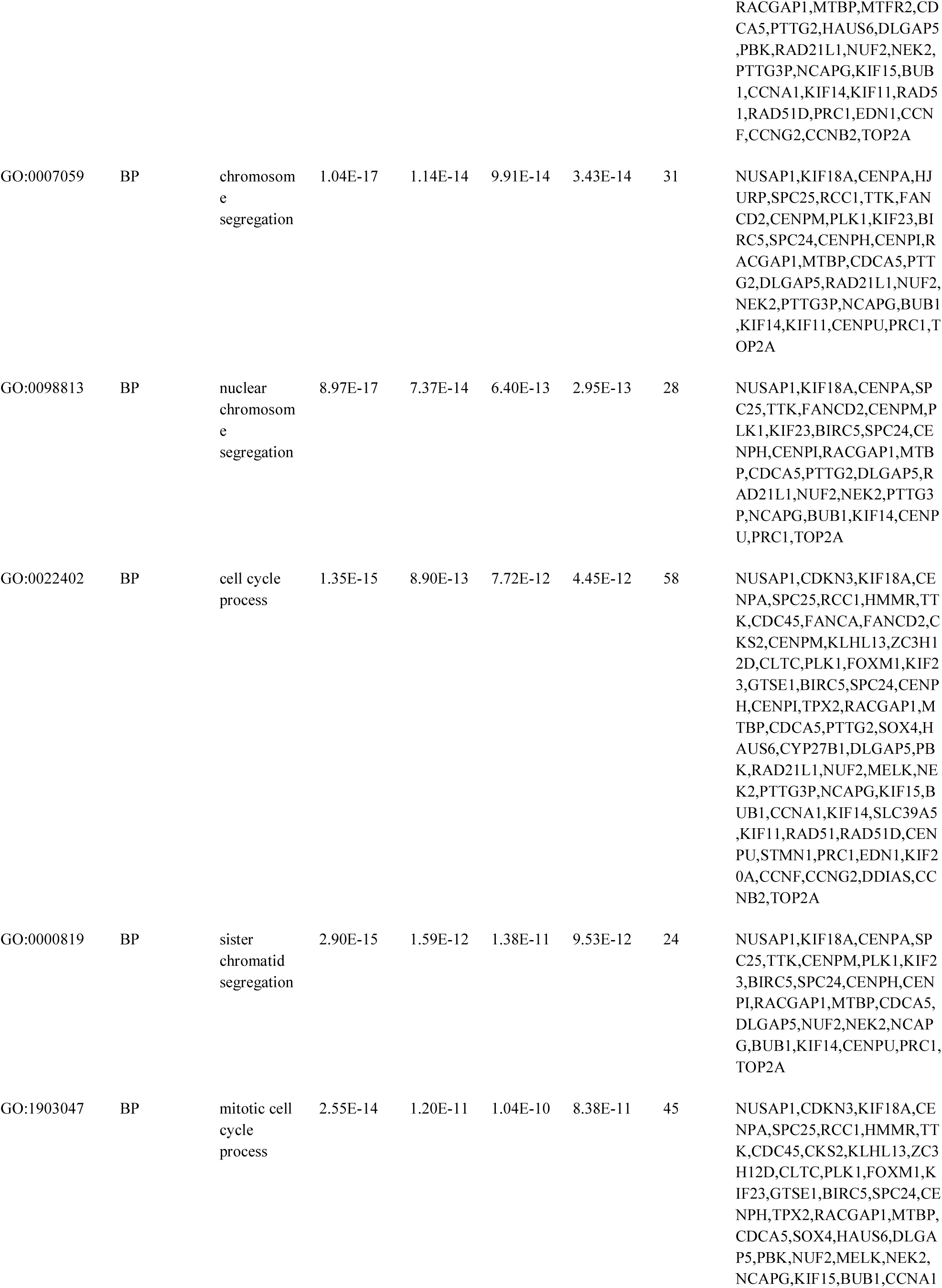
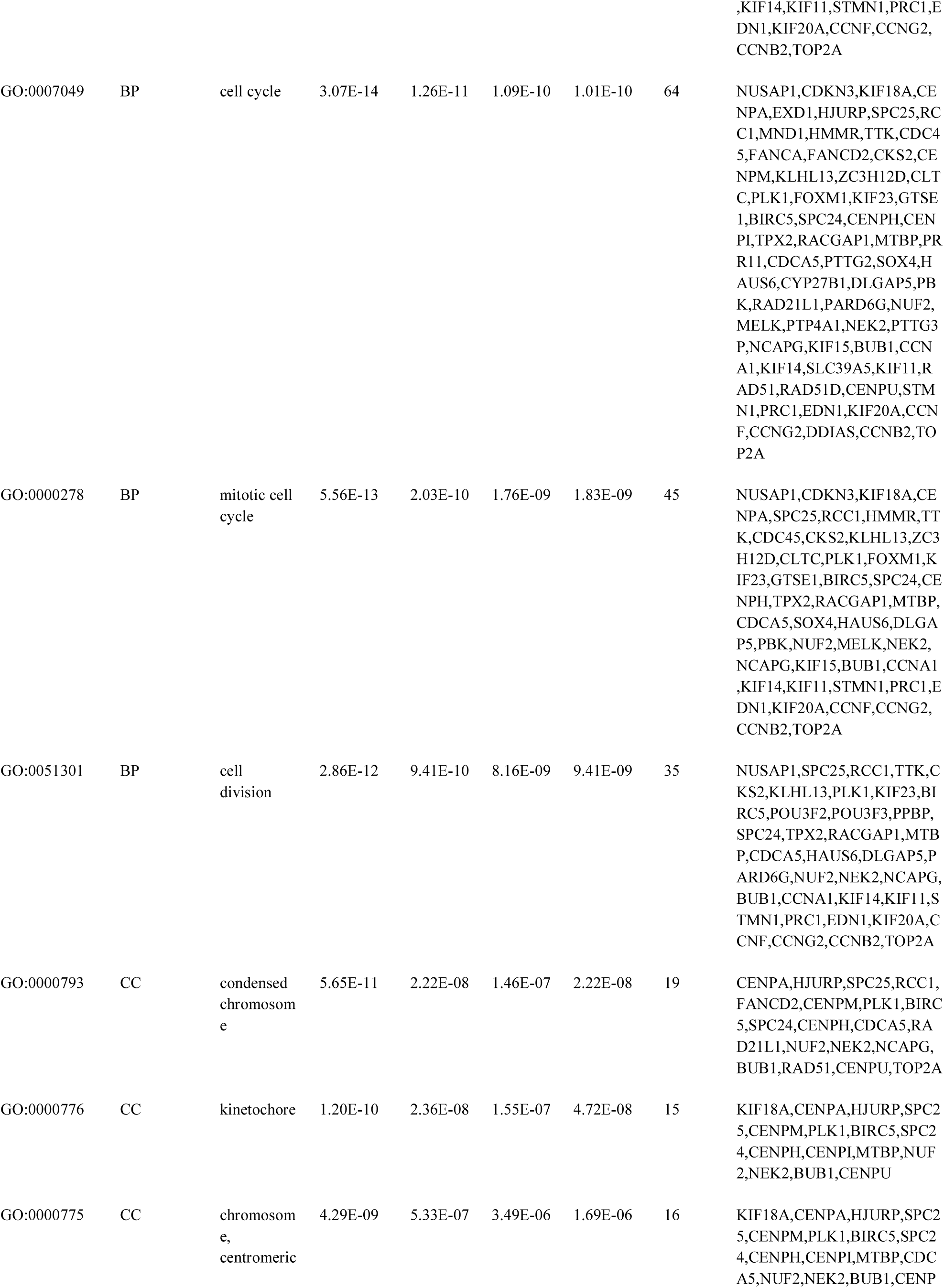
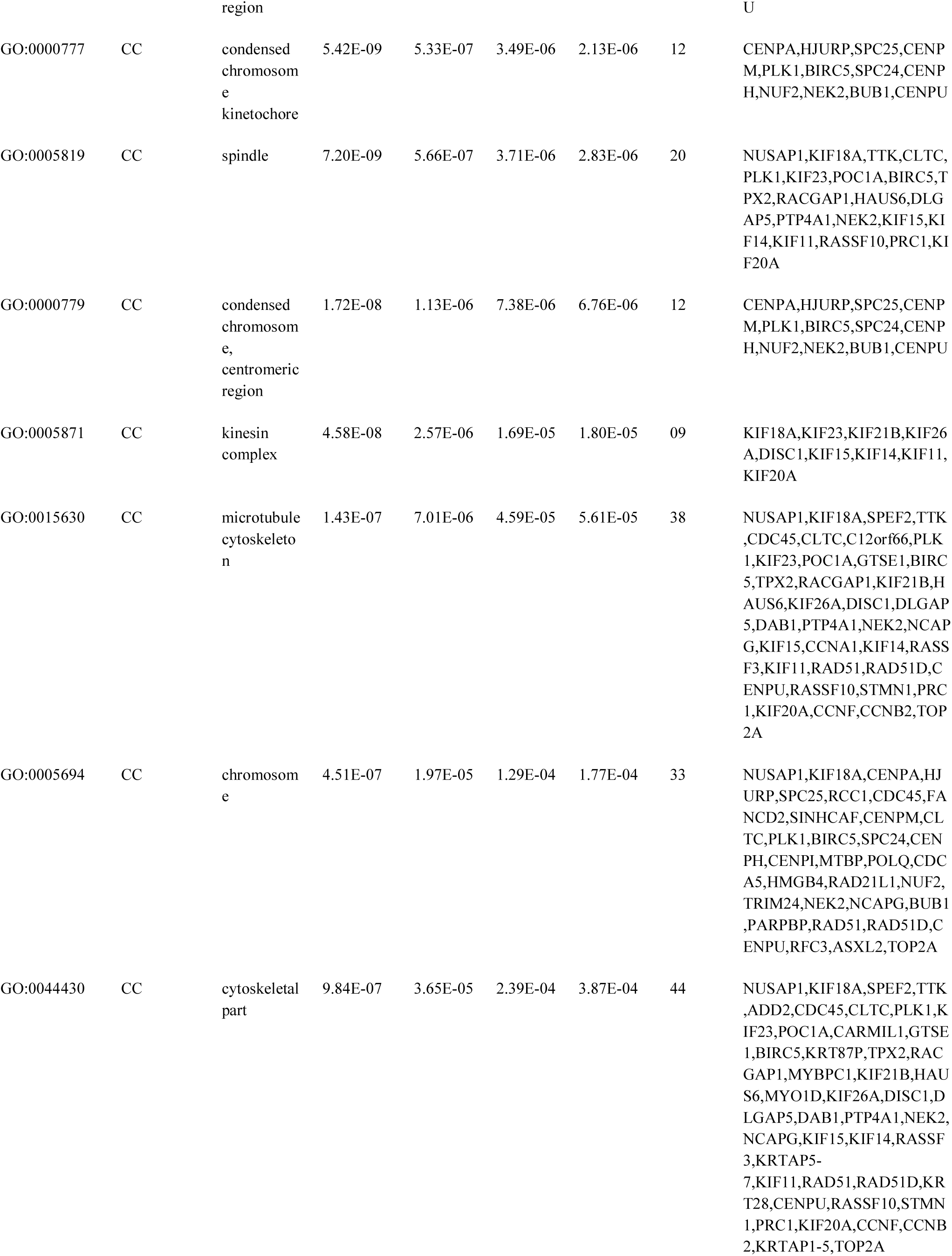
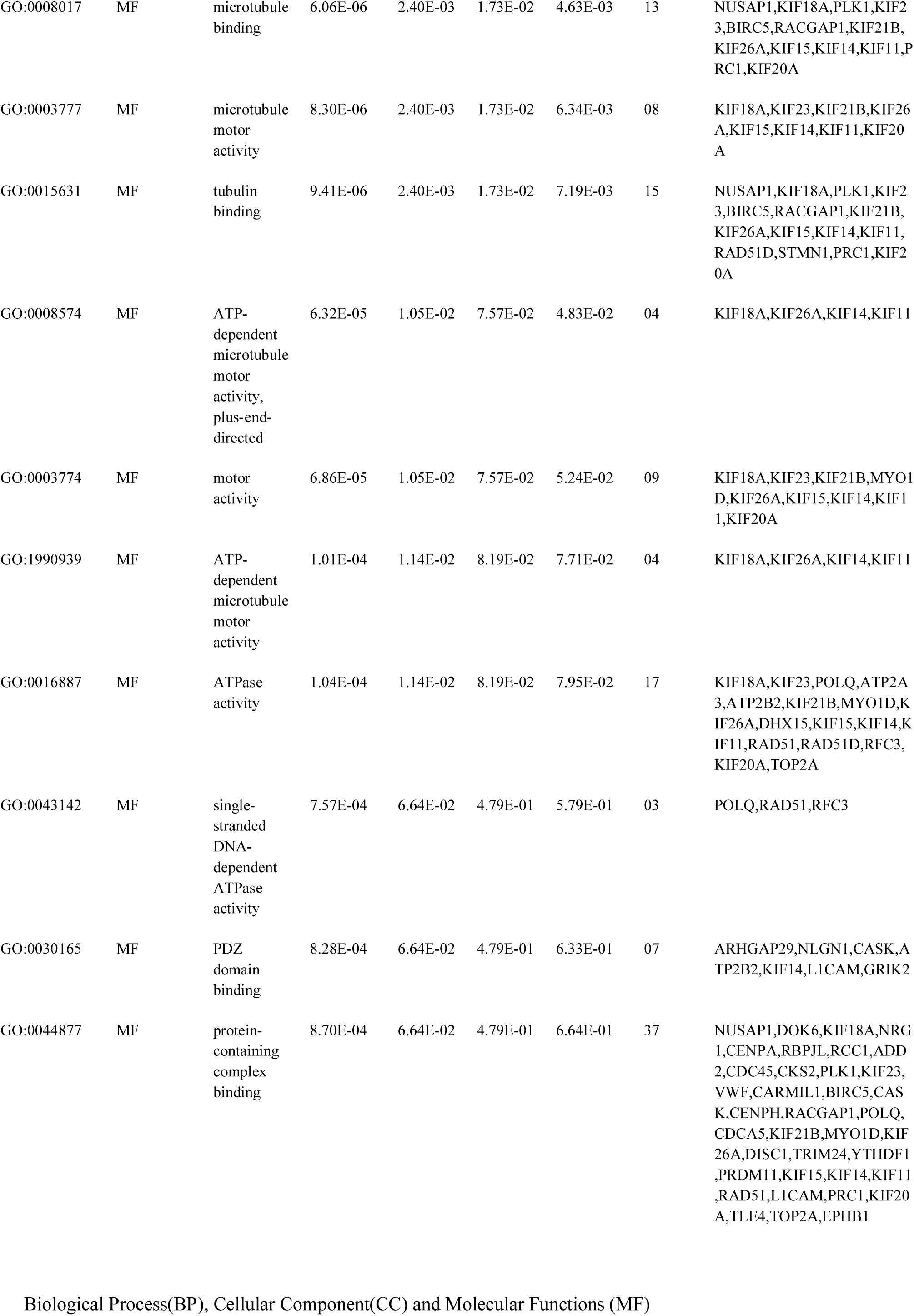
The enriched GO terms of the down-regulated differentially expressed genes

### PPI network construction and topology analysis

PPI networks were constructed on the basis of HIPPIE online tool. We also analyzed the network properties such as node degree, betweenness centrality, stress centrality, closeness centrality and cluster coefficient. The PPI network for up regulated DEGs is shown in Fig. 5, which has 4005 nodes and 5562 interactions. The top 5 nodes with greater degrees are listed in Table 6, including SOX2 (degree = 355), KRT40 (degree = 313), SMAD9 (degree = 114), AMOT (degree = 111) and DPPA4 (degree = 105). R square and correlation coefficient are 0.776 and 0.966, respectively (Fig. 7A). Top 5 up regulated genes with high betweenness centrality are SOX2 (betweeness = 0.20715558), KRT40 (betweeness = 0.15884752), SMAD9 (betweeness = 0.07865032), AMOT (betweeness = 0.06814966) and TRIM29 (betweeness = 0.0613444) shown in Table 6. R square and correlation coefficient are 0.474 and 0.098, respectively (Fig. 8A). Top 5 up regulated high stress genes are SOX2 (stress = 37524600), KRT40 (stress = 22565808), AMOT (stress = 13889668), C6orf141 (stress = 9207512) and TRIM29 (stress = 9041322) shown in Table 6. R square and correlation coefficient are 0.088 and 0.072, respectively (Fig. 8B). Top 5 up regulated gene with high closeness centrality are SOX2 (closeness = 0.31817447), KRT40 (closeness = 0.30482897), SMAD9 (closeness = 0.30309326), TRIM29 (closeness = 0.2975525) and FGFR2 (closeness = 0.29117386) shown in Table 6. R square and correlation coefficient are 0.036 and 0.082, respectively (Fig. 8C). Top 5 up regulated gene with low clustering coefficient are PRSS45 (clustering coefficient = 0), CARTPT (clustering coefficient = 0), TAC4 (clustering coefficient = 0), CT45A1 (clustering coefficient = 0) and PON3 (clustering coefficient = 0) shown in Table 6. R square and correlation coefficient are 0.616 and 0.882, respectively (Fig. 8D).

**Fig. 5.**
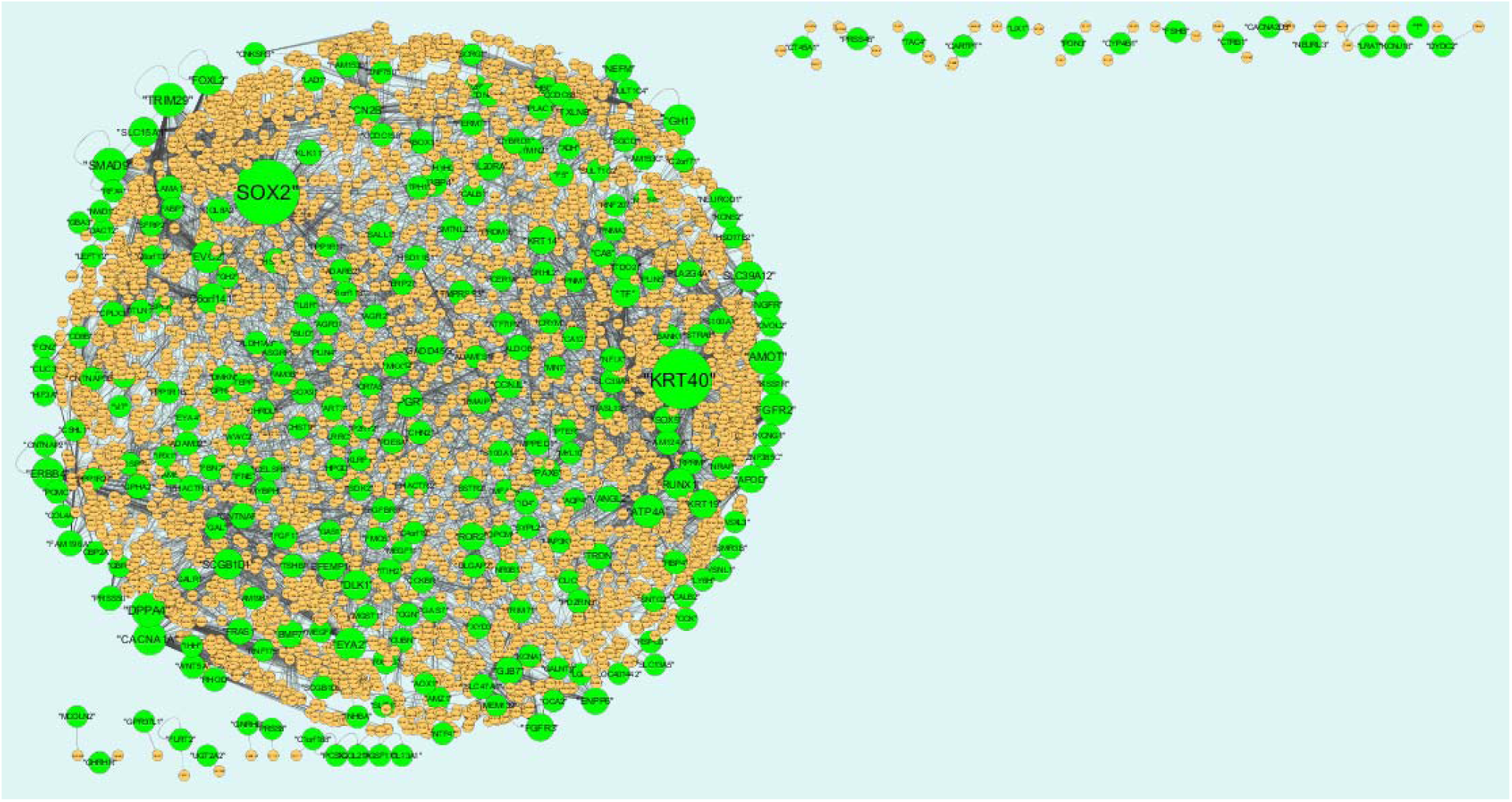
Protein–protein interaction network of differentially expressed genes (DEGs). Green nodes denotes up regulated genes.

**Fig. 7.**
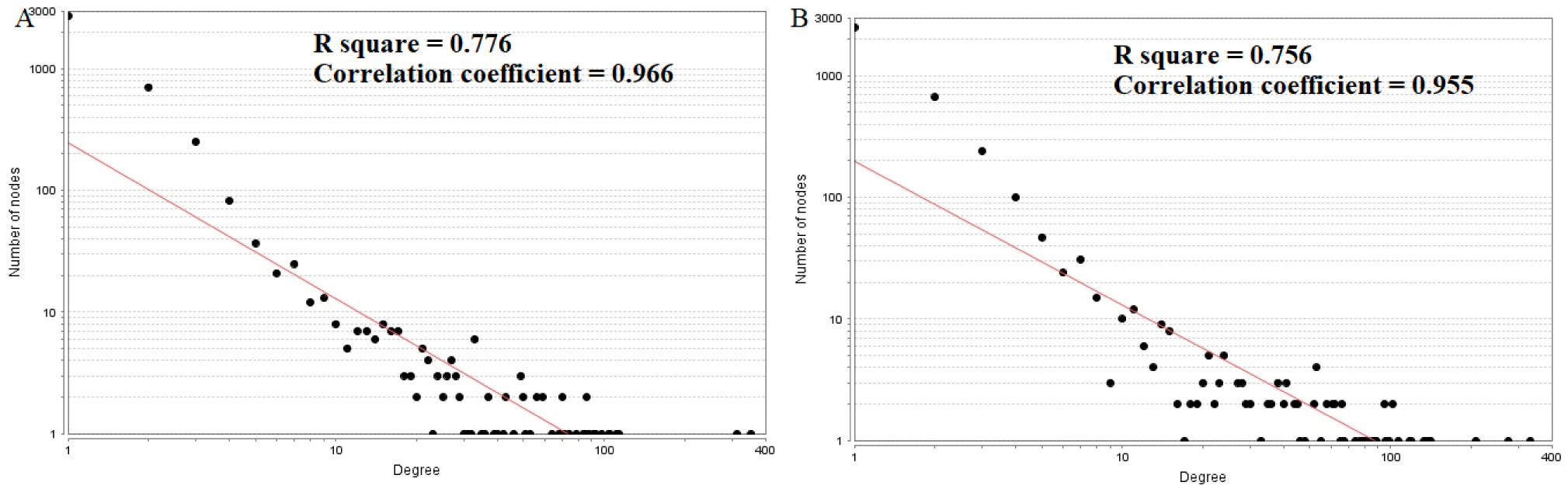
Node degree distribution (A-Up regulated genes; B- - Down regulated genes)

**Fig. 8.**
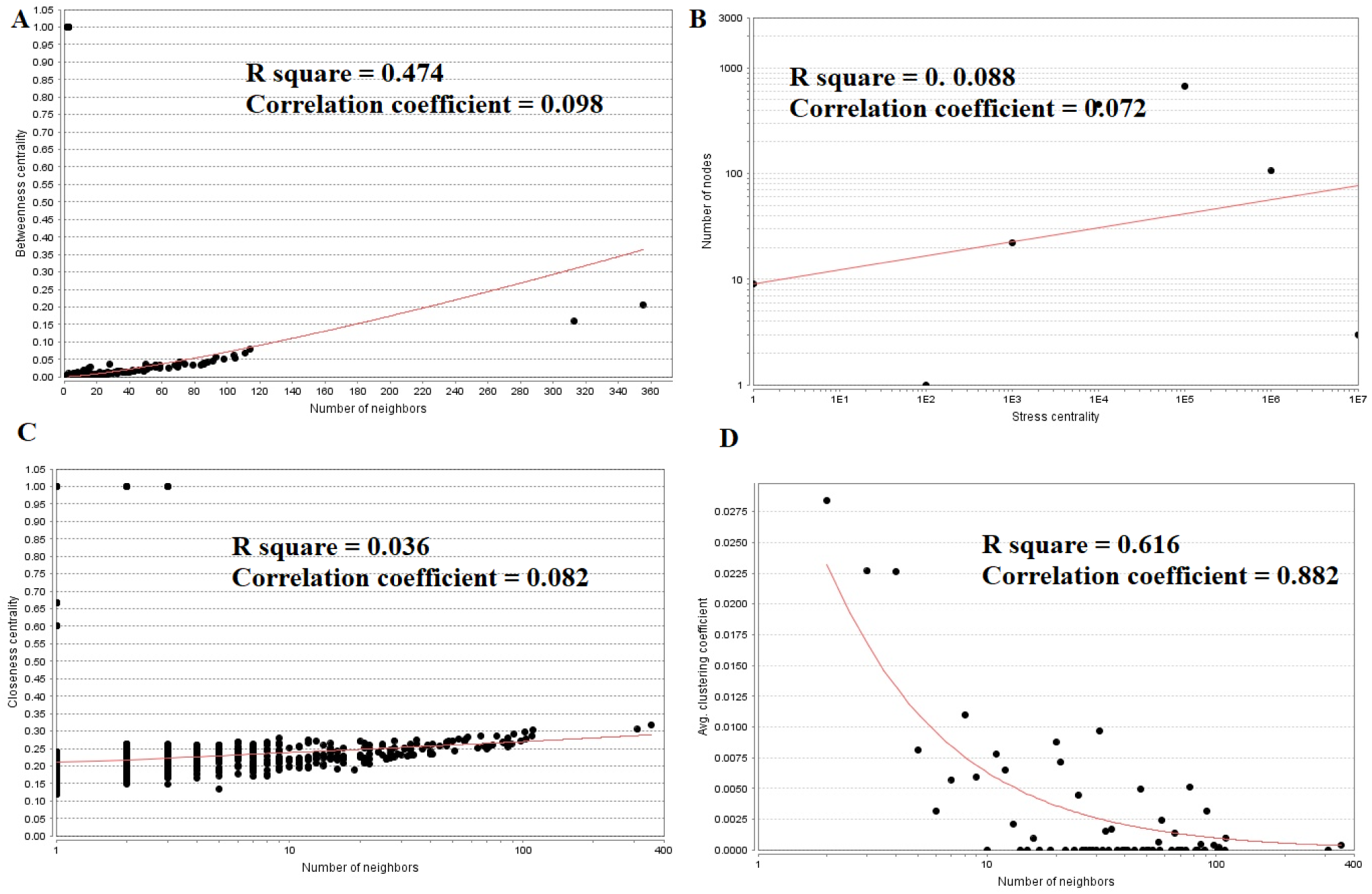
Regression diagrams for Up regulated genes (A- Betweenness centrality; B- Stress centrality; C- Closeness centrality; D- Clustering coefficient)

**Table 6.** Topology table for up and down regulated genes

| Category | Node | Degree | Betweenness | Stress | Closeness | Clustering Coefficient |
| --- | --- | --- | --- | --- | --- | --- |
| Up | SALL1 | 20 | 6.73E-03 | 599040 | 0.270852 | 0.026316 |
| Up | BMP7 | 42 | 1.85E-02 | 2547692 | 0.241805 | 0 |
| Up | FAM83F | 2 | 5.08E-04 | 43304 | 0.20099 | 0 |
| Up | SGCD | 8 | 3.55E-03 | 808402 | 0.171417 | 0 |
| Up | C1orf168 | 1 | 0.00E+00 | 0 | 1 | 0 |
| Up | GRHL2 | 7 | 1.70E-03 | 324668 | 0.243088 | 0 |
| Up | TDO2 | 18 | 3.51E-03 | 741118 | 0.221367 | 0 |
| Up | SMR3B | 11 | 2.91E-03 | 241848 | 0.209856 | 0 |
| Up | SCRG1 | 1 | 0.00E+00 | 0 | 0.171291 | 0 |
| Up | IL20RA | 36 | 1.30E-02 | 3240444 | 0.233049 | 0 |
| Up | FCN2 | 7 | 2.56E-03 | 197754 | 0.189402 | 0 |
| Up | DPPA4 | 105 | 5.28E-02 | 8404518 | 0.278355 | 1.90E-04 |
| Up | ATP4A | 91 | 4.54E-02 | 4727962 | 0.263514 | 0 |
| Up | GPR12 | 2 | 5.02E-04 | 50910 | 0.215258 | 0 |
| Up | SDK2 | 11 | 1.32E-03 | 347148 | 0.206891 | 0 |
| Up | SEPT14 | 9 | 3.56E-03 | 546322 | 0.189457 | 0 |
| Up | RSPO3 | 5 | 2.03E-03 | 178044 | 0.172718 | 0 |
| Up | SLC13A5 | 2 | 1.07E-04 | 16672 | 0.203829 | 0 |
| Up | FGFR2 | 93 | 5.79E-02 | 6781948 | 0.291174 | 0.003175 |
| Up | MGST1 | 7 | 1.66E-03 | 259370 | 0.209421 | 0 |
| Up | FABP7 | 10 | 3.53E-03 | 348144 | 0.198368 | 0 |
| Up | PON3 | 2 | 1.00E+00 | 2 | 1 | 0 |
| Up | GAL | 9 | 2.89E-03 | 341670 | 0.219065 | 0 |
| Up | PAX6 | 49 | 2.00E-02 | 1487900 | 0.266617 | 0.002775 |
| Up | SYPL2 | 12 | 3.97E-03 | 648624 | 0.216559 | 0 |
| Up | LGI1 | 2 | 5.08E-04 | 71602 | 0.185225 | 0 |
| Up | SLC15A1 | 64 | 2.63E-02 | 6462618 | 0.251196 | 0 |
| Up | TMPRSS3 | 33 | 1.58E-02 | 3314720 | 0.24797 | 0 |
| Up | ROR2 | 49 | 1.54E-02 | 2469916 | 0.257031 | 0 |
| Up | DACT2 | 4 | 1.06E-03 | 191246 | 0.177576 | 0 |
| Up | SMTNL2 | 16 | 4.38E-03 | 927394 | 0.241701 | 0 |
| Up | KRT40 | 313 | 1.59E-01 | 22565808 | 0.304829 | 0 |
| Up | FBN2 | 24 | 6.88E-03 | 1381044 | 0.220957 | 0 |
| Up | NTF4 | 5 | 2.03E-03 | 147964 | 0.13475 | 0 |
| Up | GBA3 | 2 | 5.08E-04 | 52696 | 0.148844 | 0 |
| Up | SLIT1 | 2 | 5.08E-04 | 49644 | 0.176676 | 0 |
| Up | OR7A5 | 1 | 0.00E+00 | 0 | 0.214321 | 0 |
| Up | BBOX1 | 19 | 6.49E-03 | 440000 | 0.226003 | 0 |
| Up | ART3 | 24 | 8.13E-03 | 976682 | 0.239993 | 0 |
| Up | ID4 | 13 | 3.35E-03 | 782844 | 0.231447 | 0 |
| Up | PLIN4 | 3 | 7.56E-05 | 37378 | 0.213461 | 0 |
| Up | TRDN | 37 | 1.31E-02 | 2095642 | 0.247674 | 0 |
| Up | CLIC5 | 8 | 2.39E-03 | 360044 | 0.208391 | 0 |
| Up | IRX1 | 1 | 0.00E+00 | 0 | 0.200969 | 0 |
| Up | FCER1A | 3 | 3.26E-05 | 5636 | 0.185557 | 0 |
| Up | NGEF | 21 | 7.48E-03 | 1282594 | 0.208005 | 0 |
| Up | CCKBR | 17 | 6.42E-03 | 1313620 | 0.225189 | 0 |
| Up | ITLN1 | 11 | 3.22E-03 | 584478 | 0.223274 | 0 |
| Up | S100A1 | 23 | 5.90E-03 | 824942 | 0.251533 | 0 |
| Up | TMEM139 | 3 | 3.55E-03 | 258886 | 0.184289 | 0 |
| Up | PMAIP1 | 14 | 4.74E-03 | 615246 | 0.230621 | 0 |
| Up | GHRHR | 1 | 0.00E+00 | 0 | 1 | 0 |
| Up | XDH | 2 | 5.08E-04 | 44858 | 0.200601 | 0 |
| Up | TRIM29 | 104 | 6.13E-02 | 9041322 | 0.297553 | 0 |
| Up | UGT2A2 | 1 | 0.00E+00 | 0 | 1 | 0 |
| Up | SOX5 | 37 | 1.43E-02 | 1508376 | 0.275378 | 0 |
| Up | S100A1 | 23 | 5.90E-03 | 824942 | 0.251533 | 0 |
| Up | GALR1 | 1 | 0.00E+00 | 0 | 0.179707 | 0 |
| Up | GAS1 | 3 | 1.01E-03 | 43314 | 0.226575 | 0 |
| Up | PPP1R17 | 1 | 0.00E+00 | 0 | 0.214321 | 0 |
| Up | IFNE | 10 | 2.80E-03 | 416368 | 0.217901 | 0 |
| Up | CELSR1 | 17 | 4.71E-03 | 588542 | 0.223388 | 0 |
| Up | C4orf19 | 7 | 5.94E-04 | 75814 | 0.20404 | 0 |
| Up | POMC | 13 | 5.46E-03 | 738136 | 0.221941 | 0 |
| Up | LRAT | 3 | 0.00E+00 | 0 | 1 | 0 |
| Up | KLK11 | 27 | 1.36E-02 | 2408872 | 0.236335 | 0 |
| Up | F5 | 15 | 4.61E-03 | 881738 | 0.220265 | 0 |
| Up | CNTNAP2 | 8 | 2.06E-03 | 345082 | 0.217348 | 0 |
| Up | ENPP6 | 40 | 1.39E-02 | 3610524 | 0.233408 | 0 |
| Up | GBP6 | 3 | 1.02E-03 | 53086 | 0.167077 | 0 |
| Up | OBP2A | 7 | 3.04E-03 | 662994 | 0.170129 | 0 |
| Up | RBP4 | 15 | 4.72E-03 | 550068 | 0.226992 | 0 |
| Up | PPP1R27 | 1 | 0.00E+00 | 0 | 0.166294 | 0 |
| Up | ITIH2 | 16 | 5.09E-03 | 851246 | 0.219884 | 0 |
| Up | NEURL3 | 1 | 0.00E+00 | 0 | 1 | 0 |
| Up | CLIC5 | 8 | 2.39E-03 | 360044 | 0.208391 | 0 |
| Up | PPP1R27 | 1 | 0.00E+00 | 0 | 0.166294 | 0 |
| Up | RASL11B | 6 | 1.08E-03 | 251748 | 0.233312 | 0 |
| Up | KCNJ12 | 17 | 4.14E-03 | 1228964 | 0.207786 | 0 |
| Up | AQP4 | 3 | 5.64E-04 | 54624 | 0.198508 | 0 |
| Up | CD8B | 2 | 5.08E-04 | 43304 | 0.20099 | 0 |
| Up | P2RY2 | 4 | 9.10E-04 | 137648 | 0.233077 | 0 |
| Up | IL22RA1 | 7 | 2.15E-03 | 456438 | 0.199181 | 0 |
| Up | RNF207 | 4 | 6.33E-04 | 99124 | 0.213496 | 0 |
| Up | ADAMTS18 | 5 | 7.80E-04 | 175548 | 0.216858 | 0 |
| Up | NGFR | 46 | 1.93E-02 | 2626710 | 0.258024 | 0 |
| Up | KCNJ16 | 1 | 0.00E+00 | 0 | 1 | 0 |
| Up | EFEMP1 | 56 | 3.53E-02 | 3383126 | 0.271749 | 0 |
| Up | CHN2 | 13 | 3.42E-03 | 578204 | 0.234506 | 0 |
| Up | GPHA2 | 21 | 6.38E-03 | 1083752 | 0.214718 | 0 |
| Up | GHRHR | 1 | 0.00E+00 | 0 | 1 | 0 |
| Up | KRT40 | 313 | 1.59E-01 | 22565808 | 0.304829 | 0 |
| Up | C6orf132 | 1 | 0.00E+00 | 0 | 0.195135 | 0 |
| Up | NRAP | 7 | 2.54E-03 | 723962 | 0.182437 | 0 |
| Up | PHACTR2 | 16 | 5.85E-03 | 1042524 | 0.242609 | 0 |
| Up | EVC2 | 86 | 4.01E-02 | 3002590 | 0.269961 | 0 |
| Up | FOXL2 | 70 | 2.86E-02 | 8356260 | 0.257889 | 0 |
| Up | FAM3B | 9 | 5.47E-03 | 400338 | 0.24077 | 0 |
| Up | DLK1 | 69 | 3.09E-02 | 5339202 | 0.257485 | 0 |
| Up | BLID | 12 | 2.22E-03 | 427182 | 0.219345 | 0 |
| Up | CALB1 | 8 | 1.87E-03 | 281870 | 0.227635 | 0 |
| Up | OGN | 10 | 3.36E-03 | 588086 | 0.221218 | 0 |
| Up | TF | 59 | 3.30E-02 | 5546776 | 0.282609 | 0 |
| Up | SMAD9 | 114 | 7.87E-02 | 8103704 | 0.303093 | 0 |
| Up | SOX2 | 355 | 2.07E-01 | 37524600 | 0.318174 | 0 |
| Up | KRT14 | 53 | 2.76E-02 | 6537242 | 0.277277 | 0 |
| Up | DLGAP2 | 14 | 8.62E-03 | 607018 | 0.259128 | 0 |
| Up | LRRC2 | 2 | 1.61E-05 | 2042 | 0.164853 | 0 |
| Up | AMOT | 111 | 6.81E-02 | 13889668 | 0.286702 | 0 |
| Up | ZNF385C | 7 | 2.12E-03 | 396460 | 0.198308 | 0 |
| Up | CCK | 2 | 5.08E-04 | 109288 | 0.183825 | 0 |
| Up | HSD11B1 | 16 | 5.27E-03 | 800492 | 0.218251 | 0 |
| Up | SGCD | 8 | 3.55E-03 | 808402 | 0.171417 | 0 |
| Up | PRSS8 | 1 | 0.00E+00 | 0 | 1 | 0 |
| Up | C6orf118 | 5 | 1.11E-03 | 182260 | 0.220228 | 0 |
| Up | LAD1 | 7 | 2.00E-03 | 282468 | 0.222492 | 0 |
| Up | MYL10 | 3 | 3.70E-04 | 73004 | 0.215777 | 0 |
| Up | CRISP2 | 5 | 2.02E-03 | 232114 | 0.218082 | 0 |
| Up | IHH | 2 | 5.08E-04 | 70496 | 0.159532 | 0 |
| Up | MCOLN2 | 1 | 0.00E+00 | 0 | 1 | 0 |
| Up | SLC39A12 | 70 | 3.22E-02 | 4986186 | 0.249178 | 0 |
| Up | VIT | 2 | 5.08E-04 | 69608 | 0.209376 | 0 |
| Up | PPP1R17 | 1 | 0.00E+00 | 0 | 0.214321 | 0 |
| Up | TSHB | 15 | 6.40E-03 | 1264174 | 0.237848 | 0 |
| Up | GADD45G | 56 | 2.74E-02 | 5419566 | 0.271843 | 0 |
| Up | FXYD3 | 7 | 1.80E-03 | 219094 | 0.20662 | 0 |
| Up | LIX1 | 2 | 1.00E+00 | 2 | 1 | 0 |
| Up | PRSS45 | 3 | 1.00E+00 | 6 | 1 | 0 |
| Up | MFAP4 | 6 | 1.38E-03 | 246964 | 0.216631 | 0 |
| Up | SPON1 | 3 | 1.02E-03 | 109098 | 0.214367 | 0 |
| Up | NFIX | 22 | 5.02E-03 | 1428620 | 0.232403 | 0 |
| Up | COL8A2 | 18 | 6.18E-03 | 653142 | 0.221205 | 0 |
| Up | GNRHR | 1 | 0.00E+00 | 0 | 1 | 0 |
| Up | SULT1C2 | 10 | 4.91E-03 | 344354 | 0.215375 | 0 |
| Up | CA8 | 33 | 1.30E-02 | 1324240 | 0.262215 | 0 |
| Up | CHRD1 | 6 | 7.97E-04 | 108788 | 0.198089 | 0 |
| Up | MEGF6 | 6 | 1.43E-03 | 100130 | 0.228083 | 0 |
| Up | RPRM | 9 | 3.81E-03 | 39669 | 0.211331 | 0 |
| Up | OCA2 | 3 | 8.78E-05 | 7862 | 0.168788 | 0 |
| Up | MPPED1 | 26 | 8.03E-03 | 1622472 | 0.222492 | 0 |
| Up | CARTPT | 3 | 1.00E+00 | 6 | 1 | 0 |
| Up | CSH1 | 13 | 5.61E-03 | 475316 | 0.257014 | 0 |
| Up | GALNT9 | 1 | 0.00E+00 | 0 | 0.189969 | 0 |
| Up | OGN | 10 | 3.36E-03 | 588086 | 0.221218 | 0 |
| Up | FAM153B | 14 | 5.40E-03 | 935250 | 0.2049 | 0 |
| Up | GADD45G | 56 | 2.74E-02 | 5419566 | 0.271843 | 0 |
| Up | VSNL1 | 10 | 1.31E-03 | 162184 | 0.199151 | 0 |
| Up | KISS1R | 3 | 1.02E-03 | 144830 | 0.182319 | 0 |
| Up | FGFR3 | 49 | 2.12E-02 | 2766462 | 0.268434 | 0 |
| Up | WNT5A | 22 | 7.80E-03 | 881282 | 0.206349 | 0 |
| Up | CD8B | 2 | 5.08E-04 | 43304 | 0.20099 | 0 |
| Up | SFRP2 | 17 | 3.65E-03 | 988952 | 0.223769 | 0 |
| Up | HSPB3 | 24 | 9.14E-03 | 1066592 | 0.24811 | 0 |
| Up | PAX6 | 49 | 2.00E-02 | 1487900 | 0.266617 | 0.002775 |
| Up | CYBRD1 | 21 | 7.61E-03 | 1486198 | 0.221255 | 0 |
| Up | FXYD3 | 7 | 1.80E-03 | 219094 | 0.20662 | 0 |
| Up | PGR | 68 | 3.40E-02 | 5124700 | 0.28689 | 0 |
| Up | HSD17B2 | 2 | 5.08E-04 | 61020 | 0.152911 | 0 |
| Up | OPCML | 11 | 9.23E-03 | 415810 | 0.253949 | 0 |
| Up | RFX4 | 4 | 5.08E-04 | 68768 | 0.194519 | 0 |
| Up | AGR3 | 14 | 3.65E-03 | 622642 | 0.231135 | 0 |
| Up | SLIT1 | 2 | 5.08E-04 | 49644 | 0.176676 | 0 |
| Up | SOX9 | 19 | 4.66E-03 | 1135172 | 0.228918 | 0 |
| Up | CSHL1 | 3 | 1.02E-03 | 142558 | 0.195484 | 0 |
| Up | CHST9 | 4 | 4.30E-04 | 57912 | 0.2136 | 0 |
| Up | SULT1C2 | 10 | 4.91E-03 | 344354 | 0.215375 | 0 |
| Up | DYDC2 | 1 | 0.00E+00 | 0 | 1 | 0 |
| Up | SLIT1 | 4 | 1.52E-03 | 98748 | 0.177265 | 0 |
| Up | SNTG2 | 14 | 4.17E-03 | 686616 | 0.201051 | 0 |
| Up | ZNF750 | 1 | 0.00E+00 | 0 | 0.196381 | 0 |
| Up | TAC4 | 3 | 1.00E+00 | 6 | 1 | 0 |
| Up | PTER | 14 | 4.59E-03 | 989784 | 0.223477 | 0 |
| Up | CTRB1 | 2 | 1.00E+00 | 2 | 1 | 0 |
| Up | COL21A1 | 2 | 0.00E+00 | 0 | 0 | 0 |
| Up | ATF7IP2 | 4 | 7.03E-04 | 79612 | 0.215258 | 0 |
| Up | NEFM | 50 | 2.12E-02 | 4143078 | 0.272012 | 0 |
| Up | S100A14 | 21 | 8.84E-03 | 986808 | 0.265288 | 0 |
| Up | AOX1 | 6 | 1.58E-03 | 130420 | 0.202895 | 0 |
| Up | KLRF1 | 5 | 6.70E-04 | 77804 | 0.192503 | 0 |
| Up | CCDC68 | 13 | 5.14E-03 | 436104 | 0.208269 | 0 |
| Up | CNKSR3 | 9 | 2.74E-03 | 263976 | 0.196244 | 0 |
| Up | CRYM | 4 | 5.65E-04 | 44096 | 0.209666 | 0 |
| Up | MYZAP | 1 | 0.00E+00 | 0 | 0.213484 | 0 |
| Up | PDE6A | 4 | 1.43E-04 | 8434 | 0.238597 | 0 |
| Up | INHBA | 8 | 2.57E-03 | 734598 | 0.176407 | 0 |
| Up | CNTNAP3 | 33 | 1.34E-02 | 1433542 | 0.235939 | 0 |
| Up | FAM153C | 2 | 5.08E-04 | 85808 | 0.170078 | 0 |
| Up | C6orf141 | 74 | 3.72E-02 | 9207512 | 0.260292 | 0 |
| Up | RUNX1 | 79 | 3.50E-02 | 4622022 | 0.286015 | 0 |
| Up | TRIM71 | 17 | 6.52E-03 | 713150 | 0.220623 | 0 |
| Up | PNMT | 3 | 5.88E-04 | 55878 | 0.215493 | 0 |
| Up | PRSS50 | 25 | 9.88E-03 | 1981826 | 0.256663 | 0 |
| Up | SLC47A1 | 27 | 1.28E-02 | 765214 | 0.222832 | 0 |
| Up | PHACTR3 | 29 | 1.17E-02 | 1887264 | 0.234506 | 0 |
| Up | IL20RA | 36 | 1.30E-02 | 3240444 | 0.233049 | 0 |
| Up | LEFTY2 | 3 | 1.02E-03 | 95398 | 0.147528 | 0 |
| Up | RHBDL3 | 4 | 1.52E-03 | 323400 | 0.165874 | 0 |
| Up | GH1 | 85 | 3.79E-02 | 5815610 | 0.26716 | 0 |
| Up | STRA6 | 2 | 5.08E-04 | 57684 | 0.184652 | 0 |
| Up | FRAS1 | 33 | 1.76E-02 | 1896950 | 0.244978 | 0 |
| Up | CT45A1 | 3 | 1.00E+00 | 6 | 1 | 0 |
| Up | FGFR3 | 49 | 2.12E-02 | 2766462 | 0.268434 | 0 |
| Up | EYA4 | 26 | 6.49E-03 | 1286994 | 0.239075 | 0 |
| Up | AMZ1 | 12 | 3.35E-03 | 644006 | 0.193934 | 0 |
| Up | CACNA1A | 98 | 5.07E-02 | 4392082 | 0.272841 | 0 |
| Up | CA12 | 13 | 3.67E-03 | 625692 | 0.218106 | 0 |
| Up | GAS7 | 28 | 9.64E-03 | 1499288 | 0.218894 | 0 |
| Up | HPGD | 1 | 0.00E+00 | 0 | 0.214321 | 0 |
| Up | FMO5 | 5 | 9.24E-04 | 141874 | 0.217516 | 0 |
| Up | SCGB1D2 | 9 | 2.67E-03 | 252442 | 0.209376 | 0 |
| Up | PLIN5 | 6 | 2.54E-03 | 488350 | 0.178558 | 0 |
| Up | KCNA1 | 7 | 2.68E-03 | 237802 | 0.195435 | 0 |
| Up | ALDOB | 28 | 1.11E-02 | 1644614 | 0.253084 | 0 |
| Up | RHOD | 19 | 7.80E-03 | 1084918 | 0.189786 | 0 |
| Up | GJB7 | 51 | 2.16E-02 | 2788870 | 0.243992 | 0 |
| Up | ALDH1A3 | 9 | 2.42E-03 | 378026 | 0.220327 | 0 |
| Up | EYA2 | 88 | 4.15E-02 | 6323538 | 0.281237 | 0 |
| Up | KRT19 | 59 | 2.60E-02 | 7305954 | 0.276926 | 0 |
| Up | LY6H | 3 | 5.57E-04 | 60178 | 0.176092 | 0 |
| Up | MYBPHL | 7 | 2.32E-03 | 246776 | 0.198308 | 0 |
| Up | RASGRP1 | 4 | 3.51E-04 | 53352 | 0.195019 | 0 |
| Up | GH2 | 2 | 8.82E-06 | 644 | 0.197325 | 0 |
| Up | FERMT1 | 15 | 4.58E-03 | 653358 | 0.227097 | 0 |
| Up | ASXL3 | 2 | 5.08E-04 | 44514 | 0.172627 | 0 |
| Up | TEPP | 1 | 0.00E+00 | 0 | 0.209354 | 0 |
| Up | PDZRN3 | 11 | 4.13E-03 | 858088 | 0.195387 | 0 |
| Up | KCNS2 | 1 | 0.00E+00 | 0 | 0.148569 | 0 |
| Up | VANGL2 | 33 | 1.46E-02 | 1934604 | 0.232321 | 0 |
| Up | PRDM16 | 6 | 1.44E-03 | 306010 | 0.223972 | 0 |
| Up | PLAC1 | 20 | 6.67E-03 | 1205836 | 0.224957 | 0 |
| Up | SH3BP4 | 43 | 1.67E-02 | 2668100 | 0.262618 | 0 |
| Up | COL4A6 | 9 | 4.29E-03 | 288964 | 0.222241 | 0 |
| Up | NEUROD1 | 15 | 3.84E-03 | 758734 | 0.219016 | 0 |
| Up | IL6R | 17 | 6.73E-03 | 876290 | 0.223845 | 0 |
| Up | FSHB | 2 | 1.00E+00 | 2 | 1 | 0 |
| Up | CSH2 | 13 | 5.61E-03 | 475316 | 0.257014 | 0 |
| Up | CNTNAP3B | 16 | 7.15E-03 | 582922 | 0.190741 | 0 |
| Up | CHRD1 | 6 | 7.97E-04 | 108788 | 0.198089 | 0 |
| Up | NWD1 | 1 | 0.00E+00 | 0 | 0.175339 | 0 |
| Up | WWC2 | 18 | 3.48E-03 | 975000 | 0.230041 | 0 |
| Up | APOD | 50 | 3.56E-02 | 2147634 | 0.271412 | 0 |
| Up | GJB7 | 51 | 2.16E-02 | 2788870 | 0.243992 | 0 |
| Up | RXRG | 32 | 1.14E-02 | 2426724 | 0.228268 | 0 |
| Up | RNF175 | 2 | 2.24E-05 | 1134 | 0.179355 | 0 |
| Up | IGSF11 | 2 | 0.00E+00 | 0 | 0 | 0 |
| Up | SCN2B | 86 | 3.56E-02 | 7895890 | 0.255896 | 0 |
| Up | OVOL2 | 1 | 0.00E+00 | 0 | 0.216108 | 0 |
| Up | CCDC158 | 12 | 2.91E-03 | 441576 | 0.225705 | 0 |
| Up | ASXL3 | 2 | 5.08E-04 | 44514 | 0.172627 | 0 |
| Up | CYP4B1 | 2 | 1.00E+00 | 2 | 1 | 0 |
| Up | DLGAP2 | 14 | 8.62E-03 | 607018 | 0.259128 | 0 |
| Up | TPH1 | 7 | 1.87E-03 | 396120 | 0.227635 | 0 |
| Up | AOX1 | 6 | 1.58E-03 | 130420 | 0.202895 | 0 |
| Up | SLC39A8 | 26 | 1.05E-02 | 1259100 | 0.220179 | 0 |
| Up | S100B | 33 | 1.07E-02 | 1384834 | 0.252922 | 0 |
| Up | SLPI | 7 | 4.10E-03 | 388202 | 0.209143 | 0 |
| Up | PGR | 68 | 3.40E-02 | 5124700 | 0.28689 | 0 |
| Up | SSTR2 | 7 | 2.14E-03 | 321652 | 0.229425 | 0 |
| Up | FLRT2 | 3 | 0.00E+00 | 0 | 1 | 0 |
| Up | C2orf71 | 5 | 2.03E-03 | 289652 | 0.182386 | 0 |
| Up | COL13A1 | 2 | 0.00E+00 | 0 | 0 | 0 |
| Up | CACNA2D3 | 1 | 0.00E+00 | 0 | 1 | 0 |
| Up | NR0B1 | 15 | 5.59E-03 | 309134 | 0.247425 | 0 |
| Up | CALB2 | 4 | 1.64E-05 | 1926 | 0.187706 | 0 |
| Up | STMN2 | 27 | 1.22E-02 | 1546160 | 0.247254 | 0 |
| Up | SCGB1D1 | 71 | 4.35E-02 | 6754916 | 0.254063 | 0 |
| Up | FGF1 | 27 | 1.23E-02 | 832698 | 0.262916 | 0 |
| Up | KCNA1 | 7 | 2.68E-03 | 237802 | 0.195435 | 0 |
| Up | FOXL2 | 70 | 2.86E-02 | 8356260 | 0.257889 | 0 |
| Up | MAP3K15 | 7 | 1.18E-03 | 196726 | 0.213172 | 0 |
| Up | MKX | 10 | 2.04E-03 | 492958 | 0.220302 | 0 |
| Up | TXLNB | 43 | 1.53E-02 | 4523946 | 0.233769 | 0 |
| Up | T | 1 | 0.00E+00 | 0 | 1 | 0 |
| Up | ERBB4 | 84 | 3.50E-02 | 8327424 | 0.269131 | 0 |
| Up | PPP1R1B | 8 | 1.69E-03 | 176268 | 0.187885 | 0 |
| Up | PCSK2 | 2 | 0.00E+00 | 0 | 0 | 0 |
| Up | DMKN | 7 | 2.21E-03 | 390378 | 0.236123 | 0 |
| Up | TGFBR3 | 22 | 9.08E-03 | 805594 | 0.23413 | 0 |
| Up | MN1 | 3 | 4.47E-05 | 9412 | 0.20941 | 0 |
| Up | SLIT1 | 2 | 5.08E-04 | 49644 | 0.176676 | 0 |
| Up | KCNG1 | 5 | 1.09E-03 | 200078 | 0.194567 | 0 |
| Up | MEGF11 | 2 | 5.16E-05 | 7530 | 0.192871 | 0 |
| Up | PLA2G4A | 35 | 1.68E-02 | 3125428 | 0.264079 | 0 |
| Up | ERP27 | 5 | 6.80E-04 | 96440 | 0.20941 | 0 |
| Up | GPR37L1 | 1 | 0.00E+00 | 0 | 1 | 0 |
| Up | SMR3B | 11 | 2.91E-03 | 241848 | 0.209856 | 0 |
| Up | CLIC3 | 9 | 3.25E-03 | 417244 | 0.21813 | 0 |
| Up | SYT12 | 16 | 5.27E-03 | 975528 | 0.226783 | 0 |
| Up | CUBN | 10 | 2.40E-03 | 351968 | 0.206793 | 0 |
| Up | PNMA3 | 5 | 7.32E-04 | 128516 | 0.189685 | 0 |
| Up | FAM198A | 3 | 9.84E-05 | 8632 | 0.182912 | 0 |
| Up | FAM124A | 29 | 1.00E-02 | 2057940 | 0.226796 | 0 |
| Up | CPLX3 | 17 | 7.35E-03 | 422424 | 0.250732 | 0 |
| Up | ADARB2 | 9 | 2.11E-03 | 368998 | 0.226171 | 0 |
| Up | DMKN | 7 | 2.21E-03 | 390378 | 0.236123 | 0 |
| Up | BANK1 | 4 | 1.46E-04 | 22828 | 0.217336 | 0 |
| Up | HIF3A | 3 | 6.24E-04 | 67108 | 0.19472 | 0 |
| Up | CNTNAP3 | 33 | 1.34E-02 | 1433542 | 0.235939 | 0 |
| Up | CCNJL | 39 | 1.50E-02 | 3448762 | 0.260533 | 0 |
| Up | ADAM32 | 21 | 6.74E-03 | 1279704 | 0.223185 | 0 |
| Up | PHYHD1 | 3 | 7.04E-04 | 88272 | 0.214905 | 0 |
| Up | LAMA1 | 31 | 1.32E-02 | 3129918 | 0.248298 | 0 |
| Up | DNAI2 | 12 | 3.01E-03 | 467486 | 0.2 | 0 |
| Down | TRIM24 | 66 | 2.39E-02 | 7566812 | 0.292625 | 0 |
| Down | KLHL13 | 30 | 8.55E-03 | 2607752 | 0.279499 | 0 |
| Down | MAGEB2 | 46 | 1.08E-02 | 4326748 | 0.268579 | 0 |
| Down | PPBP | 22 | 7.30E-03 | 2718118 | 0.22063 | 0 |
| Down | TTK | 77 | 3.23E-02 | 12267556 | 0.31332 | 7.21E-04 |
| Down | FAM71D | 3 | 6.39E-05 | 15292 | 0.197561 | 0 |
| Down | TOP2A | 102 | 4.15E-02 | 13126864 | 0.323325 | 0.002912 |
| Down | KIF18A | 27 | 9.22E-03 | 1047074 | 0.272947 | 0.002849 |
| Down | TPX2 | 45 | 1.33E-02 | 5212220 | 0.283324 | 0.00303 |
| Down | POC1A | 3 | 2.10E-04 | 38988 | 0.263713 | 0 |
| Down | ARHGAP29 | 14 | 2.99E-03 | 1435976 | 0.254092 | 0 |
| Down | NRG1 | 66 | 2.98E-02 | 8652224 | 0.277873 | 0 |
| Down | PLCD4 | 3 | 1.07E-03 | 235394 | 0.224878 | 0 |
| Down | CADPS2 | 15 | 4.01E-03 | 1430040 | 0.22412 | 0 |
| Down | CCNF | 13 | 2.36E-03 | 321772 | 0.266733 | 0 |
| Down | GTSE1 | 141 | 5.29E-02 | 7085030 | 0.337622 | 0.008713 |
| Down | TRHDE | 15 | 4.61E-03 | 1973410 | 0.23257 | 0 |
| Down | SOX4 | 27 | 1.24E-02 | 7096428 | 0.258212 | 0 |
| Down | DLEU1 | 21 | 6.78E-03 | 2289500 | 0.253469 | 0 |
| Down | PLK1 | 276 | 1.64E-01 | 39488508 | 0.35244 | 0.001016 |
| Down | HAUS6 | 53 | 1.32E-02 | 7866150 | 0.288256 | 0 |
| Down | BUB1 | 95 | 4.02E-02 | 4659062 | 0.318162 | 0.008649 |
| Down | SERINC2 | 8 | 2.18E-03 | 1257316 | 0.206187 | 0 |
| Down | KIF14 | 62 | 1.75E-02 | 10376292 | 0.284688 | 0 |
| Down | VWA1 | 14 | 4.26E-03 | 366250 | 0.21001 | 0 |
| Down | PTTG2 | 1 | 0.00E+00 | 0 | 0.224824 | 0 |
| Down | SEL1L | 58 | 2.60E-02 | 9557720 | 0.270569 | 0 |
| Down | NUSAP1 | 35 | 1.32E-02 | 2253376 | 0.300363 | 0.008913 |
| Down | CELA3A | 12 | 3.00E-03 | 434718 | 0.234812 | 0 |
| Down | DOK6 | 6 | 7.47E-04 | 300368 | 0.234664 | 0 |
| Down | NRG1 | 66 | 2.98E-02 | 8652224 | 0.277873 | 0 |
| Down | DHX15 | 208 | 9.39E-02 | 39966314 | 0.320847 | 0 |
| Down | NEK2 | 52 | 1.53E-02 | 5682386 | 0.277397 | 0 |
| Down | TAS2R7 | 14 | 5.78E-03 | 635150 | 0.247921 | 0.010989 |
| Down | SPC24 | 41 | 1.22E-02 | 1425412 | 0.291526 | 0.039024 |
| Down | HMMR | 44 | 1.23E-02 | 3764124 | 0.283002 | 0.003171 |
| Down | PTP4A1 | 53 | 2.21E-02 | 5500792 | 0.256435 | 0 |
| Down | DAZAP2 | 99 | 4.35E-02 | 9779248 | 0.28839 | 0.003651 |
| Down | CDCA5 | 55 | 1.97E-02 | 1719302 | 0.3 | 0.002903 |
| Down | ANTXR2 | 12 | 3.86E-03 | 876552 | 0.218842 | 0 |
| Down | FBXL5 | 41 | 1.34E-02 | 1287978 | 0.299253 | 0.006748 |
| Down | PLA2G2D | 1 | 0.00E+00 | 0 | 0.206084 | 0 |
| Down | DAB1 | 67 | 2.85E-02 | 5666990 | 0.279457 | 0.007689 |
| Down | RBPJL | 2 | 5.37E-04 | 117698 | 0.224851 | 0 |
| Down | FANCA | 86 | 2.87E-02 | 13664846 | 0.284167 | 0 |
| Down | NLGN1 | 11 | 3.82E-03 | 881670 | 0.215525 | 0 |
| Down | POU3F2 | 10 | 2.82E-03 | 567692 | 0.225258 | 0 |
| Down | PPP4R4 | 3 | 5.68E-04 | 157816 | 0.205041 | 0 |
| Down | GPR6 | 2 | 5.37E-04 | 60688 | 0.210938 | 0 |
| Down | PAX4 | 3 | 1.07E-03 | 234714 | 0.21372 | 0 |
| Down | ASXL2 | 18 | 6.05E-03 | 1686090 | 0.242263 | 0 |
| Down | IGFL3 | 1 | 0.00E+00 | 0 | 1 | 0 |
| Down | GRIK2 | 13 | 4.44E-03 | 430606 | 0.243976 | 0.025641 |
| Down | NCAPG | 52 | 1.62E-02 | 5737472 | 0.285517 | 0 |
| Down | SLC19A1 | 3 | 5.43E-04 | 132382 | 0.188974 | 0 |
| Down | EPHB1 | 14 | 3.21E-03 | 1135208 | 0.242942 | 0 |
| Down | INHBE | 15 | 7.30E-03 | 1600282 | 0.257445 | 0 |
| Down | FIBCD1 | 1 | 0.00E+00 | 0 | 1 | 0 |
| Down | RGS5 | 7 | 2.09E-03 | 462166 | 0.237446 | 0 |
| Down | CCNB2 | 36 | 8.70E-03 | 3365996 | 0.275673 | 0.009524 |
| Down | ADAMTSL1 | 7 | 2.82E-03 | 352088 | 0.208506 | 0 |
| Down | FAM83A | 7 | 1.40E-03 | 115518 | 0.279038 | 0.047619 |
| Down | LRFN1 | 20 | 9.73E-03 | 1618782 | 0.237416 | 0 |
| Down | PKLR | 19 | 4.89E-03 | 591642 | 0.267308 | 0.005848 |
| Down | SLC25A15 | 15 | 5.26E-03 | 1022014 | 0.258069 | 0 |
| Down | PRC1 | 29 | 6.84E-03 | 1142838 | 0.294452 | 0.031339 |
| Down | AMIGO2 | 6 | 1.08E-03 | 410596 | 0.178235 | 0 |
| Down | KIF11 | 134 | 5.97E-02 | 17181634 | 0.31282 | 0 |
| Down | G2E3 | 6 | 1.27E-03 | 455226 | 0.218061 | 0 |
| Down | OR6T1 | 1 | 0.00E+00 | 0 | 0.225613 | 0 |
| Down | POLQ | 1 | 0.00E+00 | 0 | 1 | 0 |
| Down | GRIK2 | 13 | 4.44E-03 | 430606 | 0.243976 | 0.025641 |
| Down | RASSF3 | 28 | 7.78E-03 | 3540288 | 0.258176 | 0 |
| Down | PBK | 107 | 4.20E-02 | 12464178 | 0.300775 | 0 |
| Down | CENPA | 88 | 3.07E-02 | 4765456 | 0.300799 | 0.006293 |
| Down | LYNX1 | 1 | 0.00E+00 | 0 | 0.189426 | 0 |
| Down | ASPDH | 10 | 3.69E-03 | 974500 | 0.235242 | 0 |
| Down | DLGAP5 | 23 | 7.53E-03 | 2047406 | 0.302705 | 0.019763 |
| Down | KIF20A | 38 | 9.99E-03 | 3761204 | 0.277108 | 0 |
| Down | CNKSR2 | 10 | 2.74E-03 | 698616 | 0.245083 | 0 |
| Down | GOLGA6A | 1 | 0.00E+00 | 0 | 0.194204 | 0 |
| Down | B4GALT6 | 4 | 1.00E+00 | 12 | 1 | 0 |
| Down | KRT28 | 1 | 0.00E+00 | 0 | 1 | 0 |
| Down | CDKN3 | 29 | 1.03E-02 | 1383840 | 0.274414 | 0.002849 |
| Down | DCAF4L2 | 7 | 1.14E-03 | 263466 | 0.229603 | 0 |
| Down | CALHM1 | 2 | 0.00E+00 | 0 | 0 | 0 |
| Down | ADD2 | 17 | 5.29E-03 | 1384972 | 0.251723 | 0 |
| Down | CDKN3 | 29 | 1.03E-02 | 1383840 | 0.274414 | 0.002849 |
| Down | RYR2 | 28 | 8.08E-03 | 2614302 | 0.258642 | 0 |
| Down | C20orf203 | 2 | 1.00E+00 | 2 | 1 | 0 |
| Down | YTHDF1 | 61 | 2.13E-02 | 7937228 | 0.298366 | 0 |
| Down | FOXM1 | 80 | 2.74E-02 | 8443400 | 0.298749 | 0.002331 |
| Down | GJA4 | 3 | 1.07E-03 | 514922 | 0.181296 | 0 |
| Down | PF4 | 11 | 5.89E-03 | 2016954 | 0.188248 | 0.027778 |
| Down | ANKFY1 | 41 | 1.24E-02 | 5747558 | 0.26362 | 0 |
| Down | AKAP3 | 10 | 3.76E-03 | 892188 | 0.240046 | 0 |
| Down | DCAF12L2 | 11 | 3.30E-03 | 771012 | 0.231472 | 0 |
| Down | BDNF | 11 | 4.67E-03 | 1264752 | 0.242373 | 0 |
| Down | BIRC5 | 61 | 2.10E-02 | 6990168 | 0.290549 | 0 |
| Down | SLC39A5 | 45 | 1.94E-02 | 2952184 | 0.228407 | 0 |
| Down | GLYATL3 | 1 | 0.00E+00 | 0 | 0.190715 | 0 |
| Down | L1CAM | 24 | 6.90E-03 | 2825930 | 0.267884 | 0 |
| Down | SV2C | 7 | 2.26E-03 | 545018 | 0.20922 | 0 |
| Down | KIF15 | 27 | 7.88E-03 | 1833408 | 0.267865 | 0 |
| Down | FANCD2 | 74 | 3.14E-02 | 6501308 | 0.312978 | 0.004813 |
| Down | RACGAP1 | 89 | 3.18E-02 | 7802394 | 0.304263 | 0.005107 |
| Down | FAM20C | 12 | 4.87E-03 | 828322 | 0.200852 | 0 |
| Down | NUF2 | 44 | 1.42E-02 | 2296298 | 0.301725 | 3.38E-02 |
| Down | QSOX1 | 15 | 4.90E-03 | 1223750 | 0.241087 | 0 |
| Down | SLITRK6 | 1 | 0.00E+00 | 0 | 0.206759 | 0 |
| Down | FAM91A1 | 23 | 7.07E-03 | 2542312 | 0.2639 | 0 |
| Down | GPR161 | 16 | 8.15E-03 | 864880 | 0.246363 | 0.008333 |
| Down | SCEL | 14 | 5.37E-03 | 1611186 | 0.249883 | 0 |
| Down | CXCR3 | 6 | 2.15E-03 | 733476 | 0.158492 | 0.066667 |
| Down | RFT1 | 21 | 6.86E-03 | 2273166 | 0.225517 | 0 |
| Down | PSG5 | 4 | 1.24E-03 | 315510 | 0.199775 | 0 |
| Down | RAD51 | 120 | 4.54E-02 | 8161594 | 0.307324 | 0.001883 |
| Down | ADAMTSL1 | 7 | 2.82E-03 | 352088 | 0.208506 | 0 |
| Down | HMGB4 | 3 | 6.06E-04 | 146328 | 0.226988 | 0 |
| Down | TROAP | 33 | 1.24E-02 | 1251972 | 0.293872 | 0.006452 |
| Down | RASSF10 | 38 | 1.21E-02 | 3884098 | 0.259851 | 0 |
| Down | ZC3H12D | 1 | 0.00E+00 | 0 | 1 | 0 |
| Down | HJURP | 21 | 4.31E-03 | 873270 | 0.271041 | 0.035088 |
| Down | XKR6 | 1 | 0.00E+00 | 0 | 0.225613 | 0 |
| Down | UNC5D | 1 | 0.00E+00 | 0 | 0.201809 | 0 |
| Down | C1orf189 | 2 | 1.00E+00 | 2 | 1 | 0 |
| Down | PSTPIP2 | 8 | 1.87E-03 | 472358 | 0.247427 | 0 |
| Down | XKR6 | 1 | 0.00E+00 | 0 | 0.225613 | 0 |
| Down | EXD1 | 1 | 0.00E+00 | 0 | 1 | 0 |
| Down | KIF21B | 30 | 9.70E-03 | 2431650 | 0.256842 | 0 |
| Down | GRIP1 | 53 | 1.84E-02 | 6233398 | 0.270372 | 0.001569 |
| Down | POU3F3 | 3 | 5.62E-04 | 84710 | 0.208378 | 0 |
| Down | CPN1 | 6 | 1.71E-03 | 328228 | 0.22172 | 0 |
| Down | SPC25 | 35 | 6.24E-03 | 630312 | 0.27207 | 0.052101 |
| Down | C12ORF66 | 4 | 1.61E-03 | 77742 | 0.201711 | 0 |
| Down | MELK | 23 | 6.63E-03 | 1668988 | 0.250673 | 0 |
| Down | TLE4 | 28 | 9.56E-03 | 2438214 | 0.252507 | 0 |
| Down | MND1 | 7 | 2.70E-03 | 788252 | 0.224986 | 0 |
| Down | ZMAT3 | 24 | 7.69E-03 | 2968830 | 0.271931 | 0 |
| Down | ATP2A3 | 53 | 2.39E-02 | 6619182 | 0.256171 | 0 |
| Down | GRIP1 | 53 | 1.84E-02 | 6233398 | 0.270372 | 0.001569 |
| Down | CENPM | 15 | 2.31E-03 | 174586 | 0.25524 | 0.114286 |
| Down | GRIK2 | 13 | 4.44E-03 | 430606 | 0.243976 | 0.025641 |
| Down | ZSCAN2 | 1 | 0.00E+00 | 0 | 1 | 0 |
| Down | GPR26 | 1 | 0.00E+00 | 0 | 1 | 0 |
| Down | NXPH4 | 4 | 1.61E-03 | 1133406 | 0.17246 | 0 |
| Down | UNC5D | 1 | 0.00E+00 | 0 | 0.201809 | 0 |
| Down | RNF215 | 1 | 0.00E+00 | 0 | 1 | 0 |
| Down | STMN1 | 65 | 2.51E-02 | 7293058 | 0.286065 | 4.81E-04 |
| Down | ARHGAP20 | 2 | 1.00E-05 | 3872 | 0.219564 | 0 |
| Down | FAM72D | 1 | 0.00E+00 | 0 | 0.225613 | 0 |
| Down | HTR3D | 4 | 7.06E-04 | 153078 | 0.180646 | 0 |
| Down | ATP2B2 | 118 | 5.57E-02 | 10270314 | 0.275124 | 2.90E-04 |
| Down | CCNG2 | 11 | 3.40E-03 | 744714 | 0.210806 | 0 |
| Down | RCC1 | 102 | 3.87E-02 | 10546556 | 0.314165 | 0.001941 |
| Down | NPVF | 3 | 3.46E-05 | 17228 | 0.217653 | 0 |
| Down | KIF23 | 62 | 2.33E-02 | 4491558 | 0.306515 | 0.015819 |
| Down | PSG11 | 4 | 1.12E-03 | 136444 | 0.188324 | 0 |
| Down | PRR11 | 95 | 3.70E-02 | 16341014 | 0.259923 | 0 |
| Down | LCE1B | 22 | 1.04E-02 | 1365886 | 0.209408 | 0 |
| Down | CCNA1 | 83 | 2.81E-02 | 8701494 | 0.2972 | 2.47E-03 |
| Down | TLE4 | 28 | 9.56E-03 | 2438214 | 0.252507 | 0 |
| Down | HIGD1B | 3 | 1.00E+00 | 6 | 1 | 0 |
| Down | OXR1 | 10 | 1.57E-03 | 396854 | 0.225914 | 0 |
| Down | DEFA3 | 1 | 0.00E+00 | 0 | 0.184876 | 0 |
| Down | DISC1 | 138 | 5.45E-02 | 23062588 | 0.288949 | 0 |
| Down | CLTC | 333 | 1.73E-01 | 59969246 | 0.345833 | 0.001483 |
| Down | MTBP | 7 | 9.00E-04 | 308238 | 0.25191 | 0 |
| Down | COLEC10 | 14 | 5.78E-03 | 496564 | 0.230569 | 0 |
| Down | PNPLA7 | 1 | 0.00E+00 | 0 | 1 | 0 |
| Down | TLE4 | 28 | 9.56E-03 | 2438214 | 0.252507 | 0 |
| Down | MYO1D | 58 | 2.01E-02 | 6943106 | 0.297723 | 0 |
| Down | CXCR2 | 9 | 2.70E-03 | 571596 | 0.206862 | 0 |
| Down | POU3F4 | 3 | 3.10E-04 | 10712 | 0.214335 | 0 |
| Down | KIF26A | 3 | 5.58E-04 | 216568 | 0.213757 | 0 |
| Down | VWF | 20 | 6.90E-03 | 3379006 | 0.226877 | 0 |
| Down | ZSCAN12 | 40 | 1.72E-02 | 4511560 | 0.249014 | 0 |
| Down | OMP | 1 | 0.00E+00 | 0 | 1 | 0 |
| Down | SPINK5 | 1 | 0.00E+00 | 0 | 0.200236 | 0 |
| Down | MYBPC1 | 24 | 1.02E-02 | 2208902 | 0.223047 | 0 |
| Down | RFC3 | 48 | 1.19E-02 | 6744340 | 0.286615 | 0 |
| Down | KIAA1958 | 14 | 4.06E-03 | 1021098 | 0.246641 | 0 |
| Down | TMEM40 | 1 | 0.00E+00 | 0 | 1 | 0 |
| Down | CTAG1A | 2 | 5.37E-04 | 160340 | 0.182031 | 0 |
| Down | CHRNA7 | 5 | 1.20E-03 | 315000 | 0.238785 | 0 |
| Down | PARD6G | 21 | 6.07E-03 | 2092448 | 0.251349 | 0 |
| Down | ZNF160 | 4 | 1.00E+00 | 12 | 1 | 0 |
| Down | CCL3L3 | 1 | 1.00E+00 | 6 | 1 | 0 |
| Down | CKS2 | 36 | 8.66E-03 | 1293372 | 0.279227 | 2.50E-02 |
| Down | HSD3B2 | 19 | 6.19E-03 | 434014 | 0.258678 | 0.005848 |
| Down | RAD21L1 | 1 | 0.00E+00 | 0 | 0.198223 | 0 |
| Down | HS3ST2 | 11 | 4.28E-03 | 1134682 | 0.226849 | 0 |
| Down | GRM5 | 10 | 2.07E-03 | 756604 | 0.225846 | 0 |
| Down | SPEF2 | 3 | 1.07E-03 | 311542 | 0.166704 | 0 |
| Down | KIR3DX1 | 3 | 5.37E-04 | 151612 | 0.229872 | 0 |
| Down | EYS | 1 | 0.00E+00 | 0 | 0.217653 | 0 |
| Down | CENPH | 38 | 1.29E-02 | 1191544 | 0.293478 | 0.022222 |
| Down | C3orf52 | 11 | 3.74E-03 | 897310 | 0.232454 | 0 |
| Down | UNC13A | 7 | 2.18E-03 | 1007070 | 0.215862 | 0 |
| Down | RUFY4 | 3 | 5.58E-04 | 112984 | 0.219927 | 0 |
| Down | GPR17 | 1 | 0.00E+00 | 0 | 1 | 0 |
| Down | RAD21L1 | 1 | 0.00E+00 | 0 | 0.198223 | 0 |
| Down | CASK | 79 | 3.54E-02 | 8491074 | 0.295926 | 0 |
| Down | OXR1 | 10 | 1.57E-03 | 396854 | 0.225914 | 0 |

The PPI network for down regulated DEGs is shown in Fig. 6, which has 5441 nodes and 9866 interactions. The top 5 nodes with grater degrees are listed in Table 6, including CLTC (degree = 333), PLK1 (degree = 276), DHX15 (degree = 208), GTSE1 (degree = 141) and DISC1 (degree = 138). R square and correlation coefficient are 0.756 and 0.955, respectively (Fig. 7B). Top 5 down regulated genes with high betweenness centrality are CLTC (betweeness = 0.17279707), PLK1 (betweeness = 0.16393694), DHX15 (betweeness = 0.09387272), KIF11 (betweeness = 0.05972209) and ATP2B2 (betweeness = 0.0557287) shown in Table 6. R square and correlation coefficient are 0.596 and 0.119, respectively (Fig. 9A). Top 5 down regulated genes with high stress genes are CLTC (stress = 59969246), DHX15 (stress = 39966314), PLK1 (stress = 39488508), ATP2B3 (stress = 23062588) and KIF11 (stress = 17181634) shown in Table 6. R square and correlation coefficient are 0.371 and 0.004, respectively (Fig. 9B). Top 5 down regulated genes with high closeness centrality are PLK1 (closeness = 0.35244041), CLTC (closeness = 0.34583256), ATP2B4 (closeness = 0.33762233), ATP2B8 (closeness = 0.32332523) and DHX15 (closeness = 0.32084733) shown in Table 6. R square and correlation coefficient are 0.081 and 0.144, respectively (Fig. 9C). Top 5 up regulated gene with low clustering coefficient are B4GALT6 (clustering coefficient = 0), ZNF160 (clustering coefficient = 0), HIGD1B (clustering coefficient = 0), CCL3L3 (clustering coefficient = 0) and C20orf203 (clustering coefficient = 0) shown in Table 6. R square and correlation coefficient are 0.569 and 0.860, respectively (Fig. 9D).

**Fig. 6.**
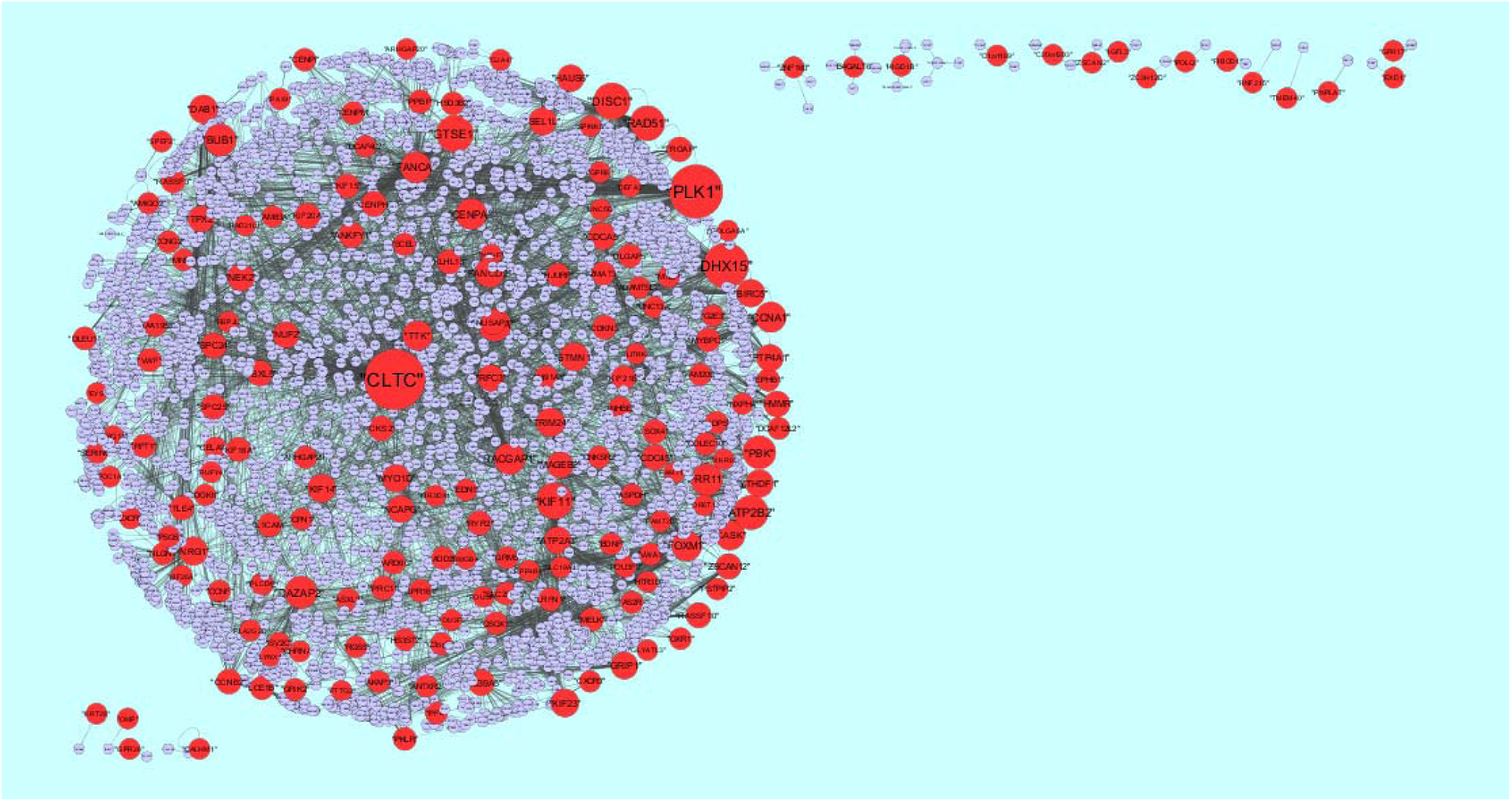
Protein–protein interaction network of differentially expressed genes (DEGs). Red nodes denotes up regulated genes.

**Fig. 9.**
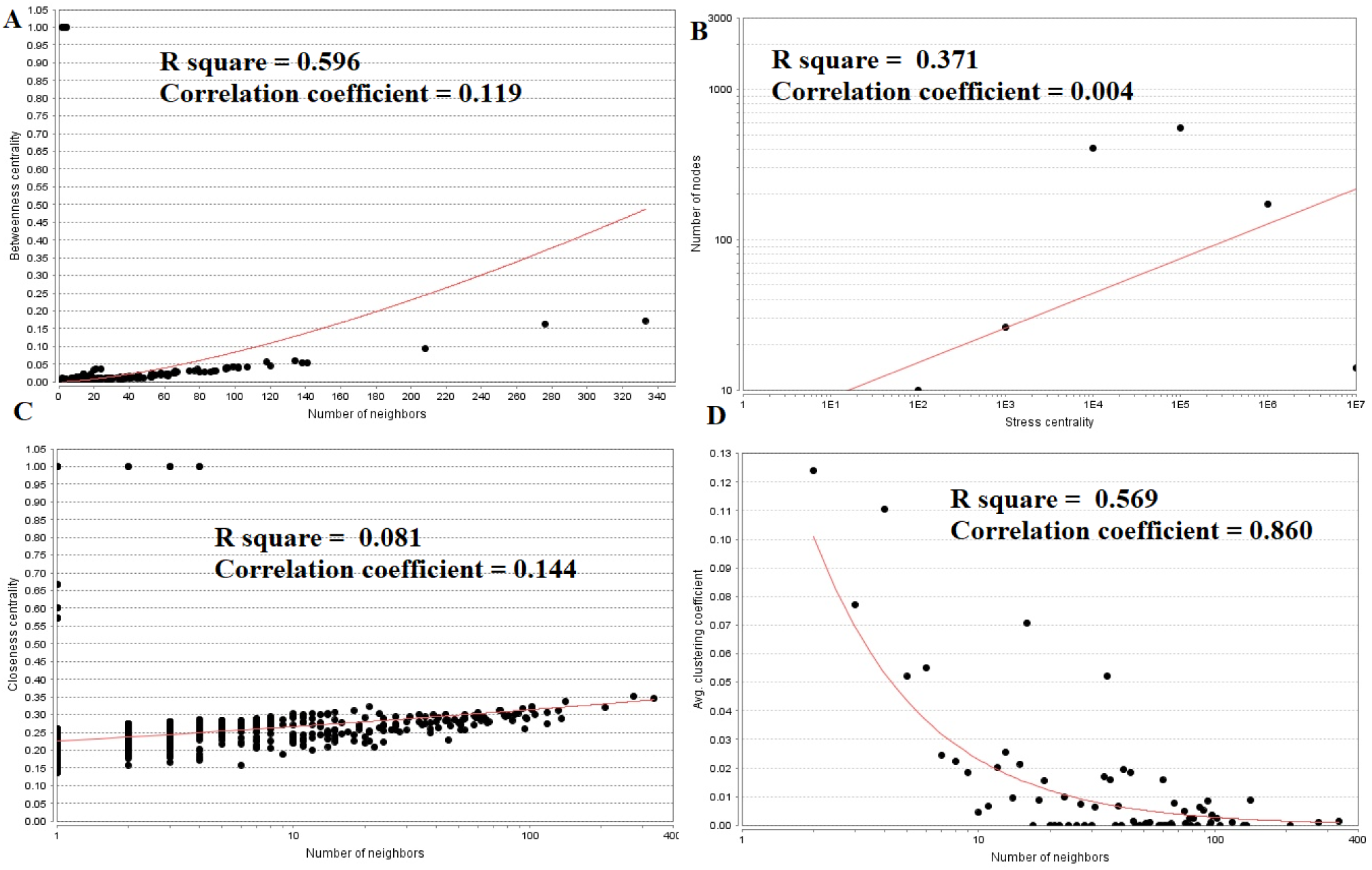
Regression diagrams for Down regulated genes (A- Betweenness centrality; B- Stress centrality; C- Closeness centrality; D- Clustering coefficient)

### Module analysis

A total of 332 modules are identified in up regulated PPI network, among which the best are module 1, module 2, module 3 and module 10 (Fig. 10). Module 1 is composed of 17 nodes and 33 edges. The hub proteins in this module such as RUNX1 (degree = 79) and SOX2 (degree = 355) are involved in module 1. Module 2 is composed of 11 nodes and 23 edges. The hub proteins in this module such as FGFR2 (degree = 93), FGF1 (degree = 27) and FGFR3 (degree = 49) are involved in module 2. Module 3 is composed of 11 nodes and 21 edges. The hub proteins in this module such as S100B (degree = 33) and S100A1 (degree = 23) are involved in module 3. Module 10 is composed of 5 nodes and 8 edges. The hub proteins in this module such as SMAD9 (degree = 114) and EVC2 (degree = 86) are involved in module 10.

**Fig. 10.**
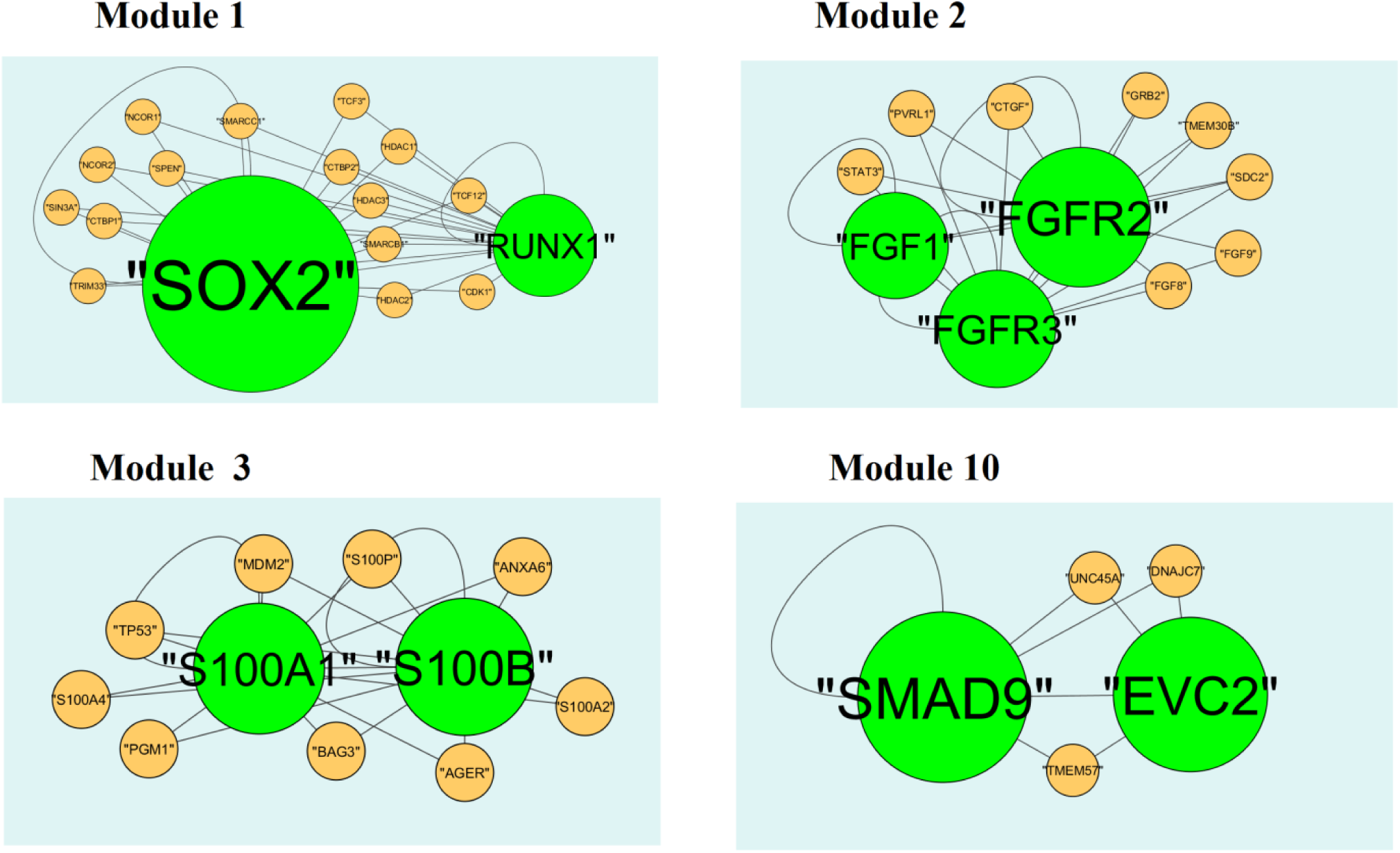
Modules in PPI network. The green nodes denote the up-regulated genes.

A total of 425 modules are identified in down regulated PPI network, among which the best are module 1, module 5, module 8 and module 12 (Fig.11). Module 1 is composed of 80 nodes and 157 edges. The hub proteins in this module such as CLTC (degree = 333) and GTSE1 (degree = 141) are involved in module 1. Module 5 is composed of 15 nodes and 43 edges. The hub proteins in this module such as NUF2 (degree = 44), BUB1 (degree = 95), SPC24 (degree = 41) and SPC25 (degree = 35) are involved in module 5. Module 8 is composed of 11 nodes and 23 edges. The hub proteins in this module such as CKS2 (degree = 36), CCNB2 (degree = 36) and CCNA1 (degree = 83) are involved in module 8. Module 12 is composed of 9 nodes and 17 edges. The hub proteins in this module such as KIF18A (Degree 27), FOXM1 (degree = 80) and PRC1 (degree = 29) are involved in module 12.

**Fig. 11.**
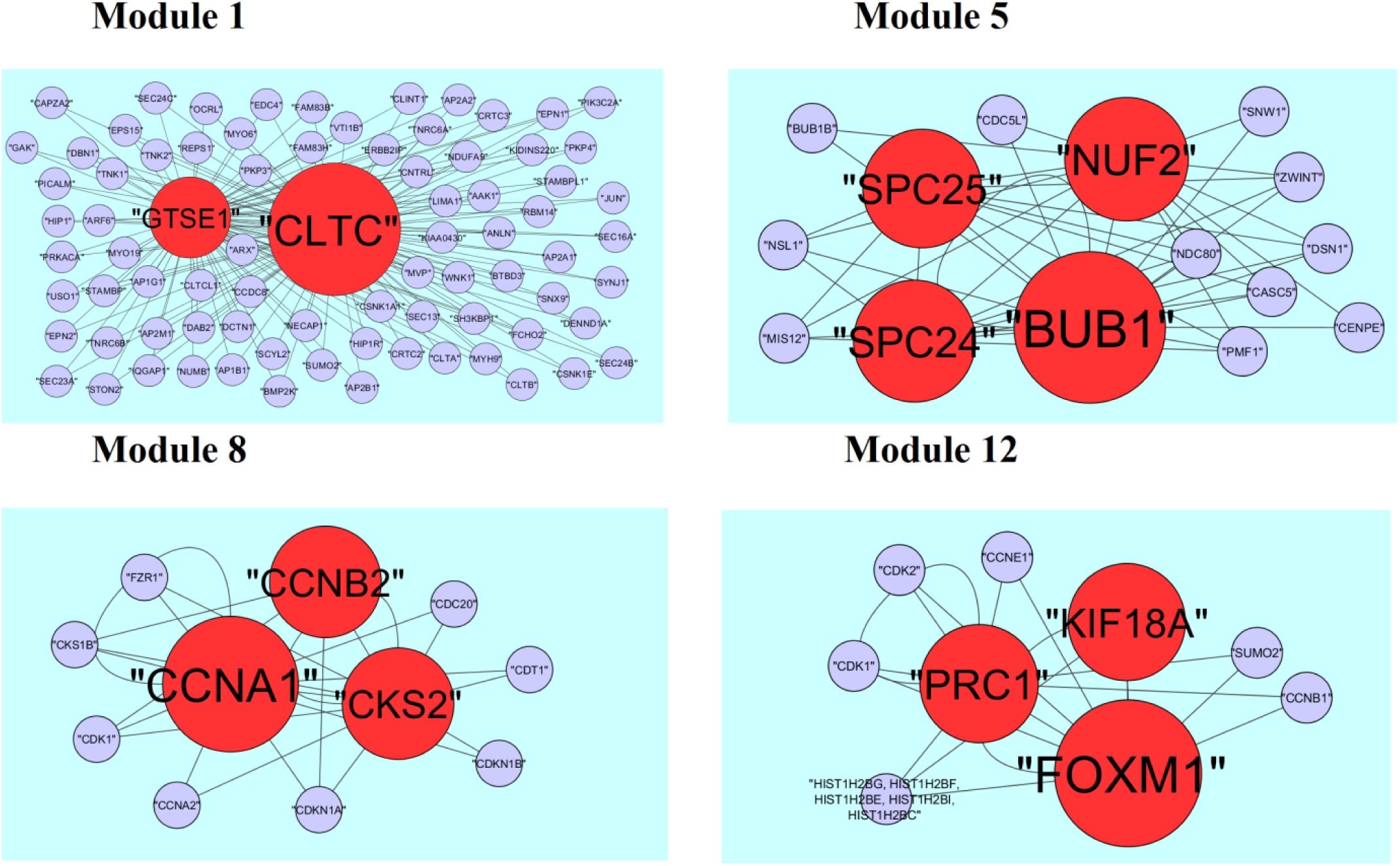
Modules in PPI network. The red nodes denote the down-regulated genes.

### Construction of the target gene **-** miRNA network

The miRNAs that may control the DEGs are diagnosed based on the up and down regulation expressions (Fig. 12 and Fig. 13). Top 5 up regulated targeted genes such as LINC00598 regulated by 209 miRNAs, CNKSR3 regulated by 138 miRNAs, PMAIP1 regulated by 128 miRNAs, TRIM71 regulated by 104 miRNAs and FAM83F regulated by 94 miRNAs are given in Table 7. Top 5 down regulated targeted genes such as SOX4 regulated by 160 miRNAs, ZMAT3 regulated by 145 miRNAs, PTP4A1 regulated by 132 miRNAs, RAD51 regulated by 113 miRNAs and DAZAP2 regulated by 109 miRNAs are given in Table 7.

**Fig. 12.**
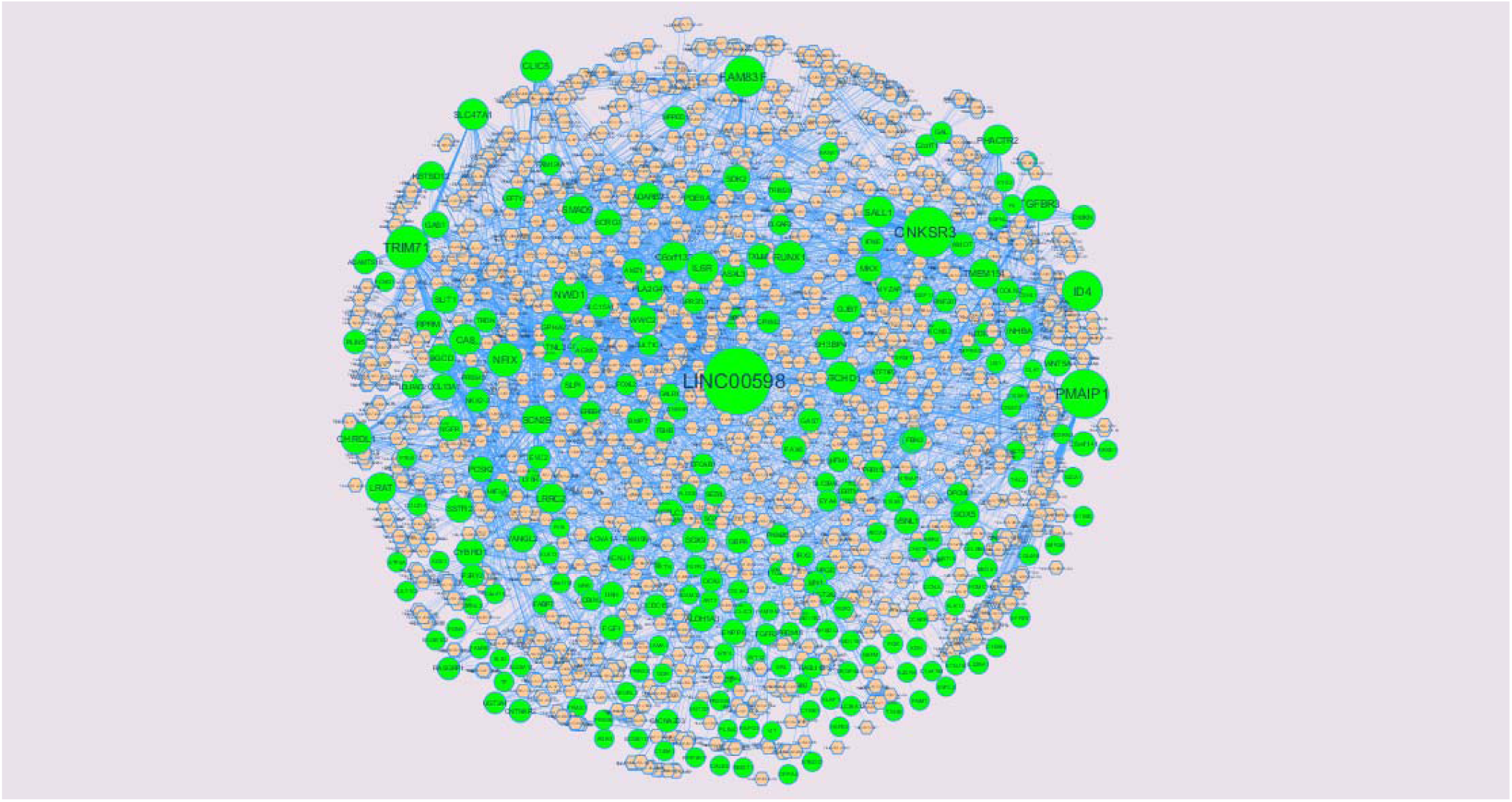
The network of up-regulated DEGs and their related miRNAs. The green circles nodes are the up regulated DEGs and pinkiamond nodes are the miRNAs

**Fig. 13.**
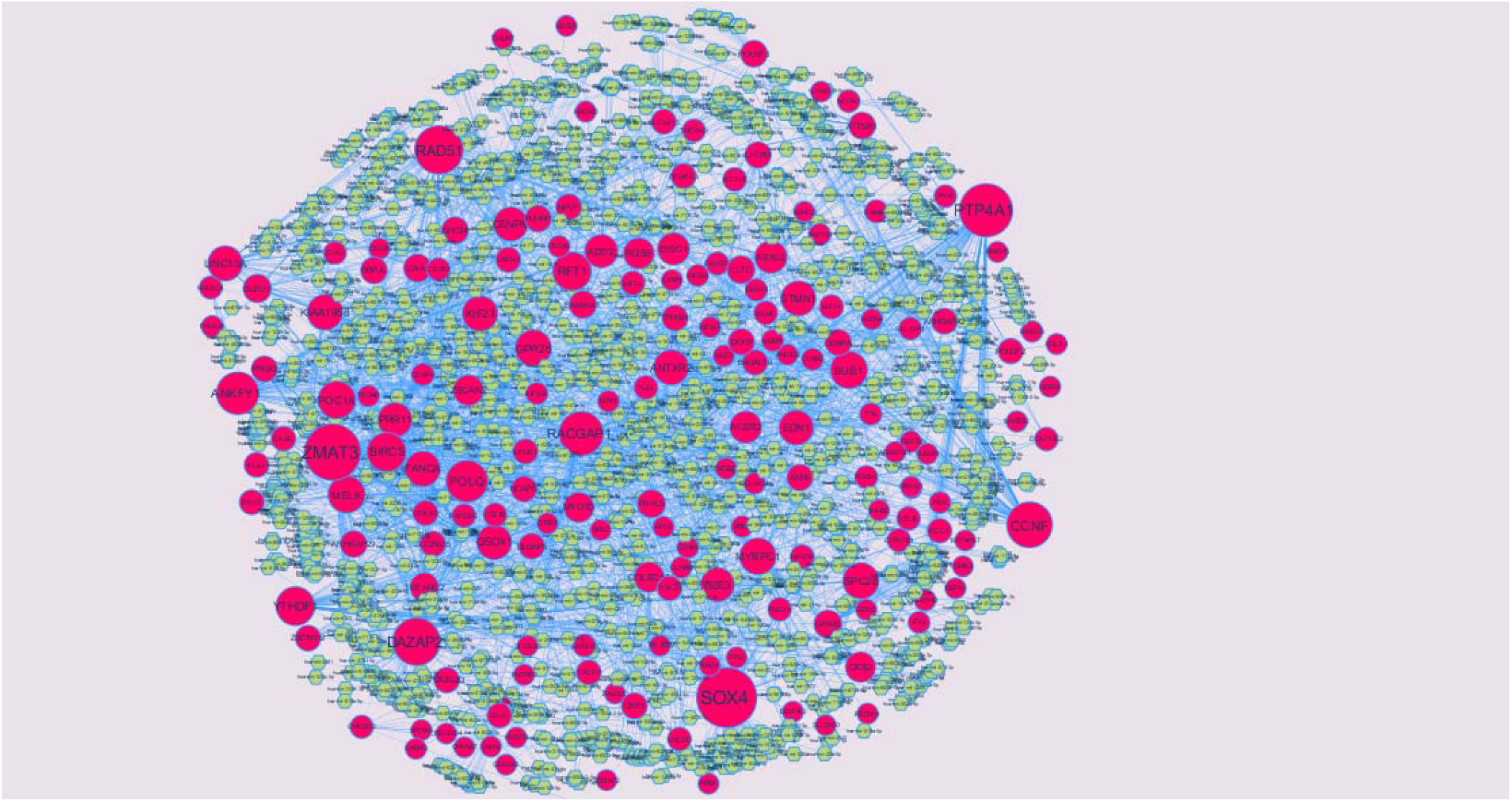
The network of up-regulated DEGs and their related miRNAs. The pink circles nodes are the up regulated DEGs and yellow diamond nodes are the miRNAs

**Table 7.**
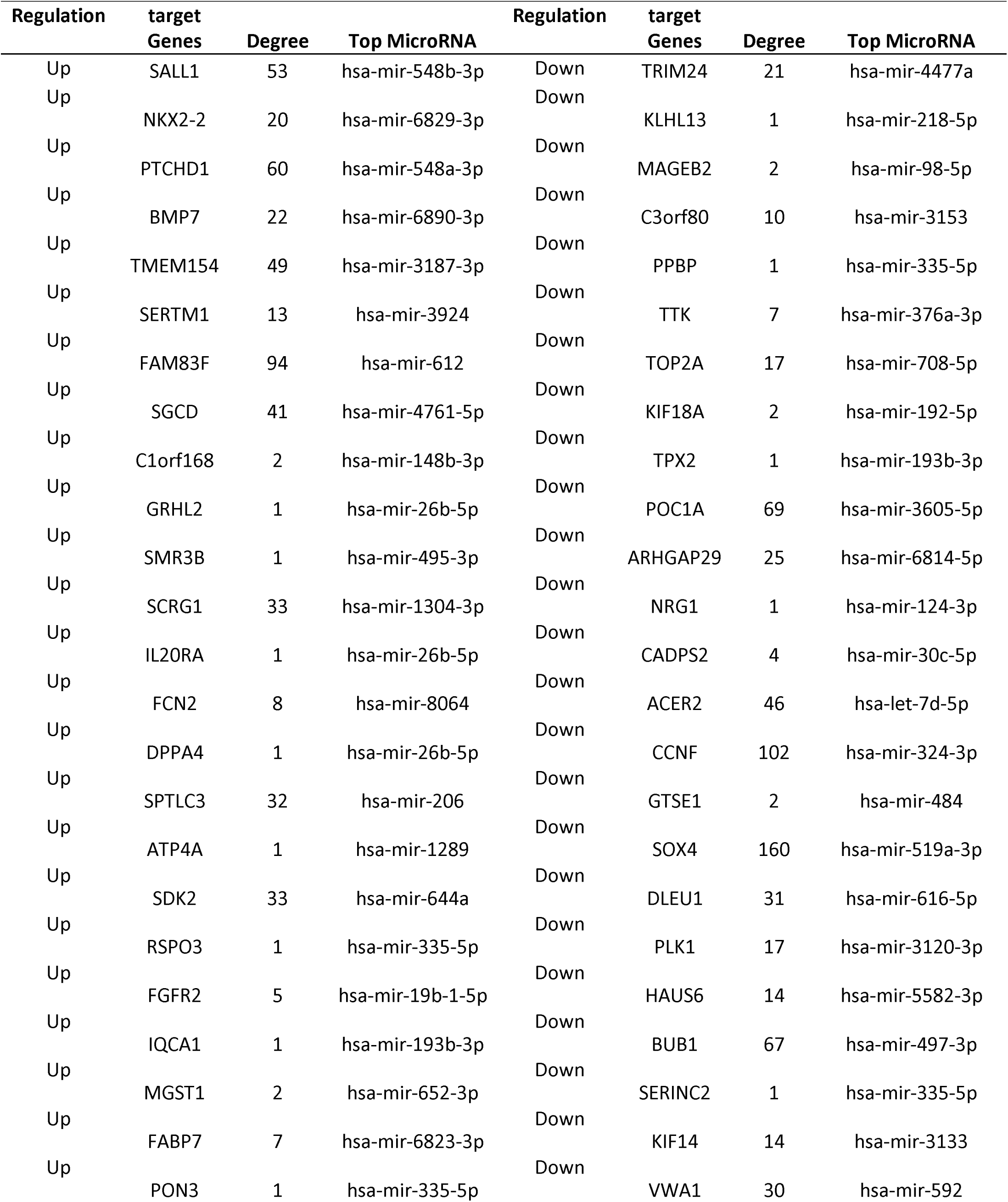
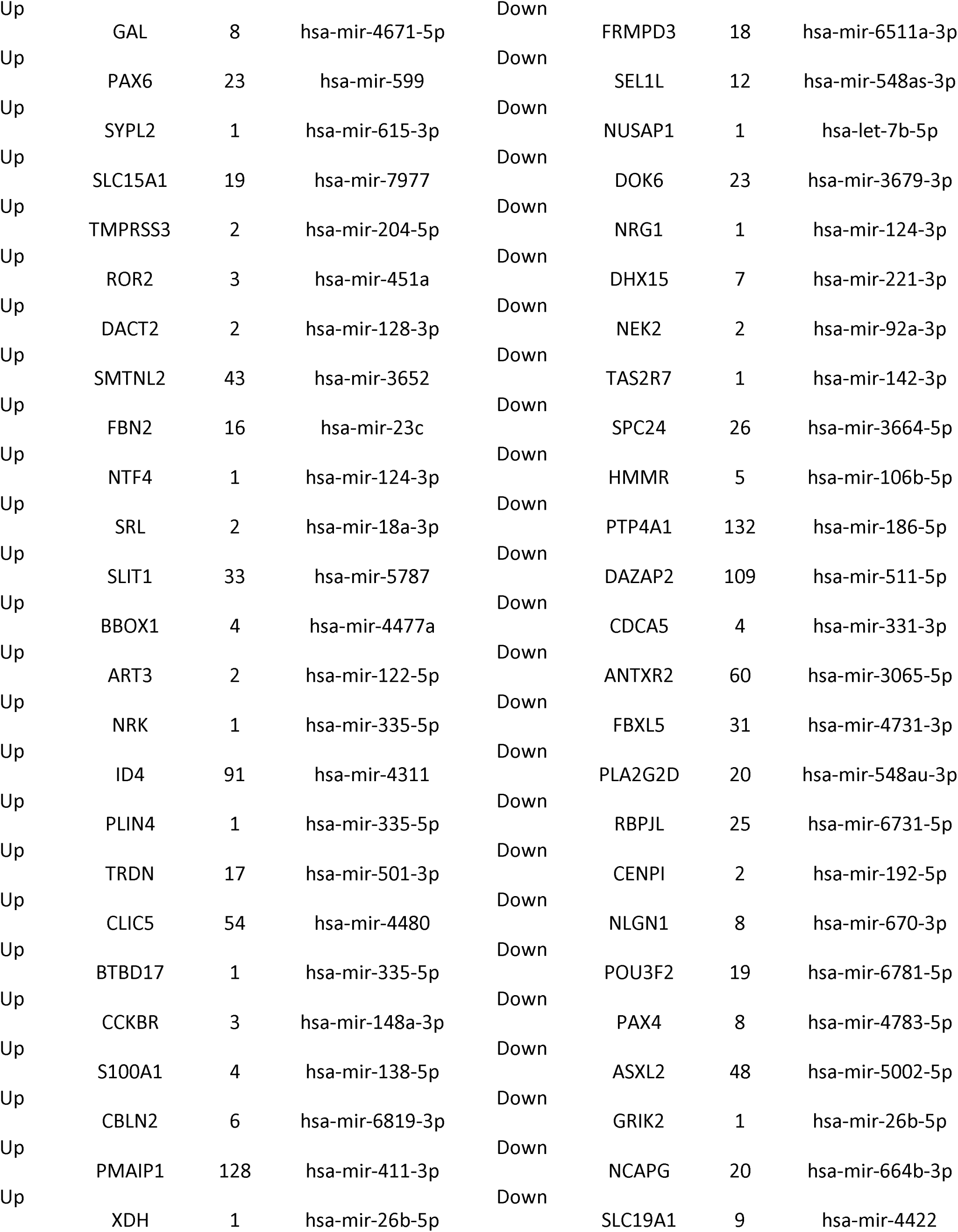
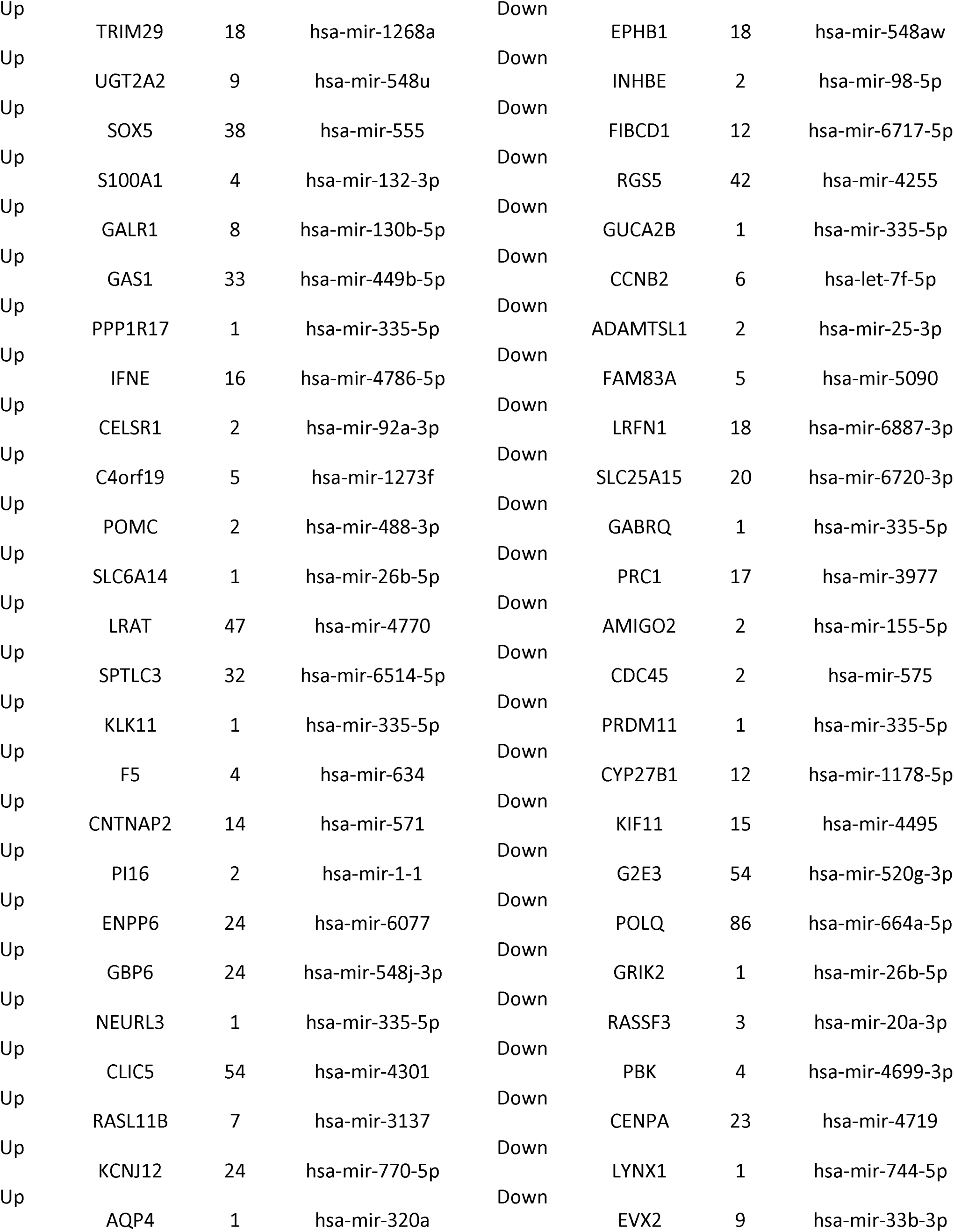
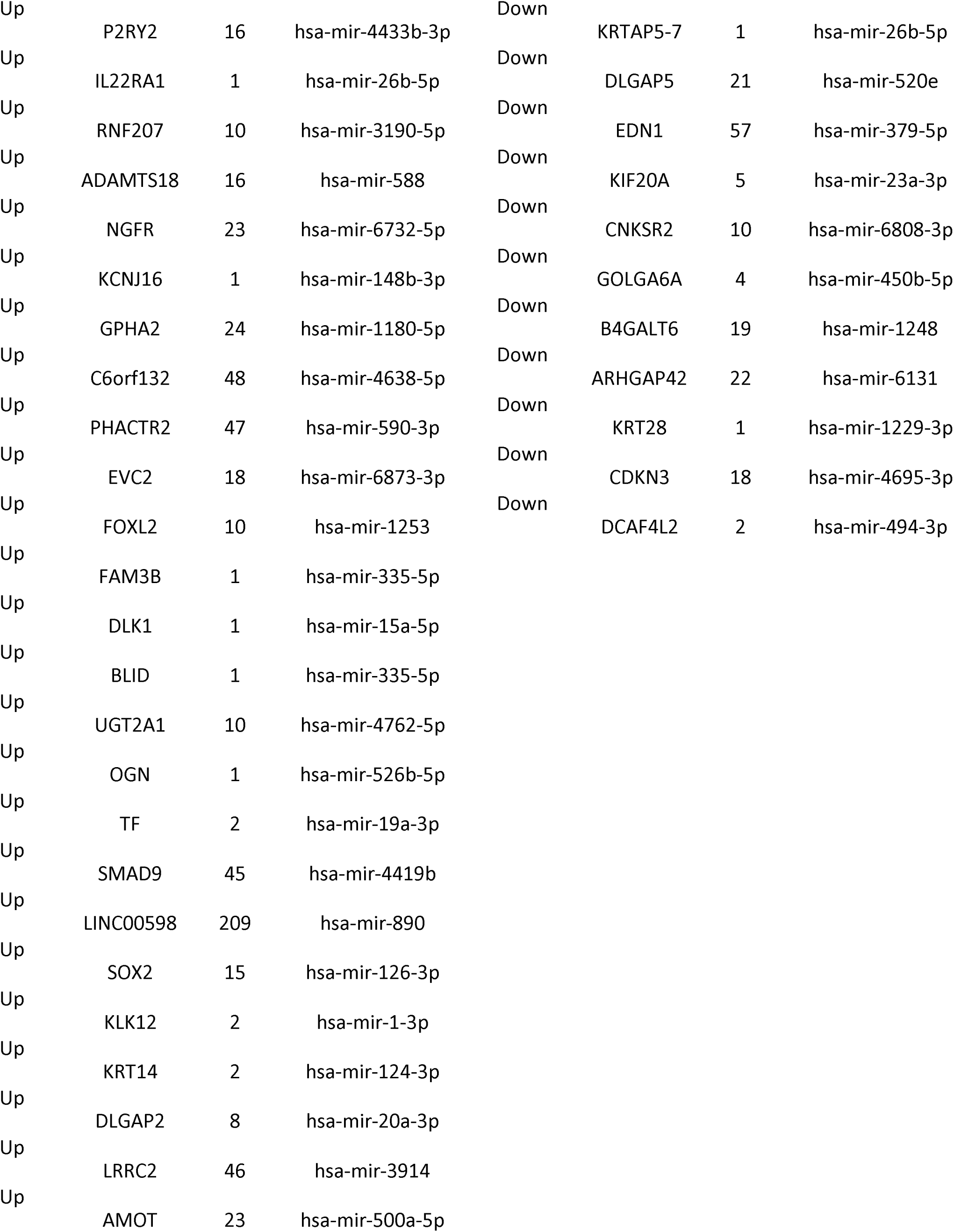
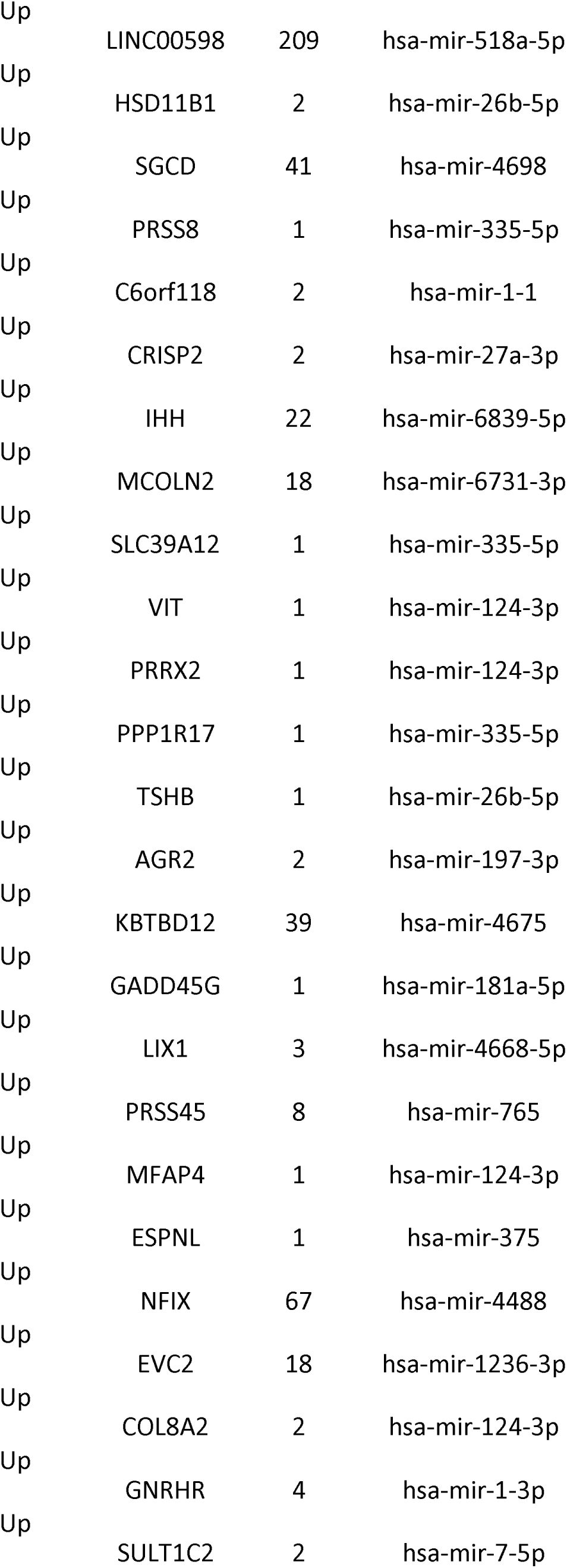
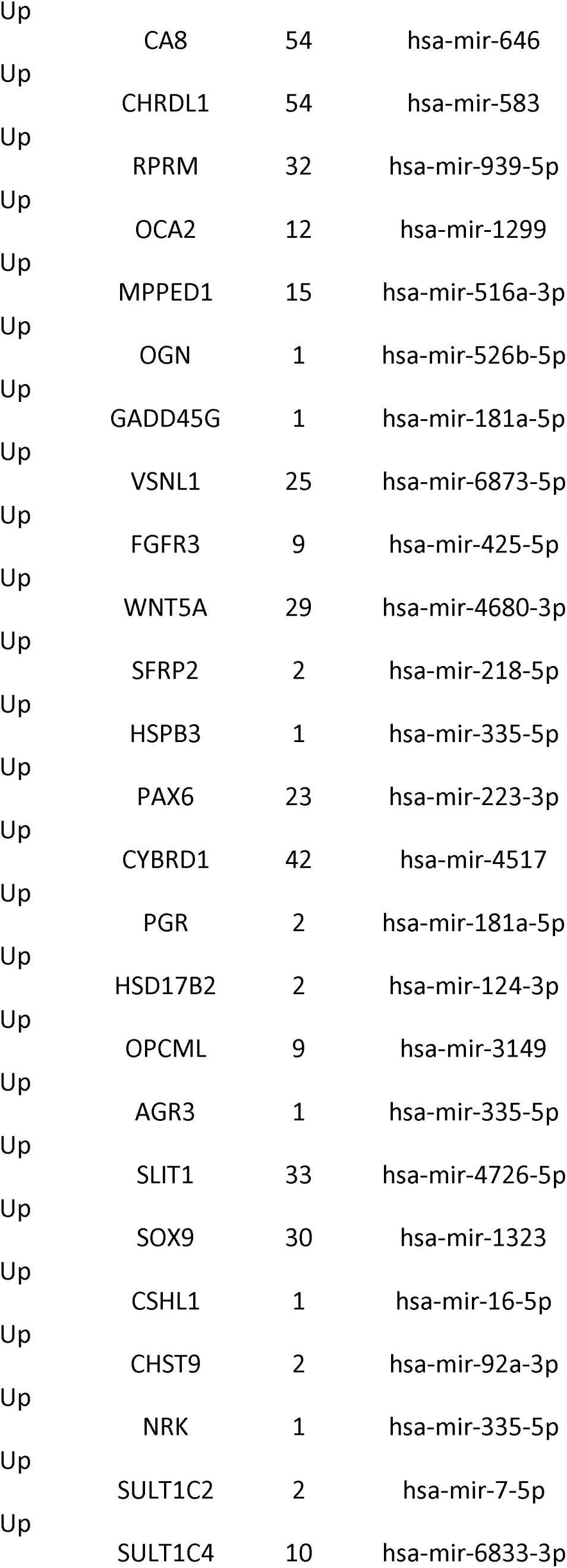
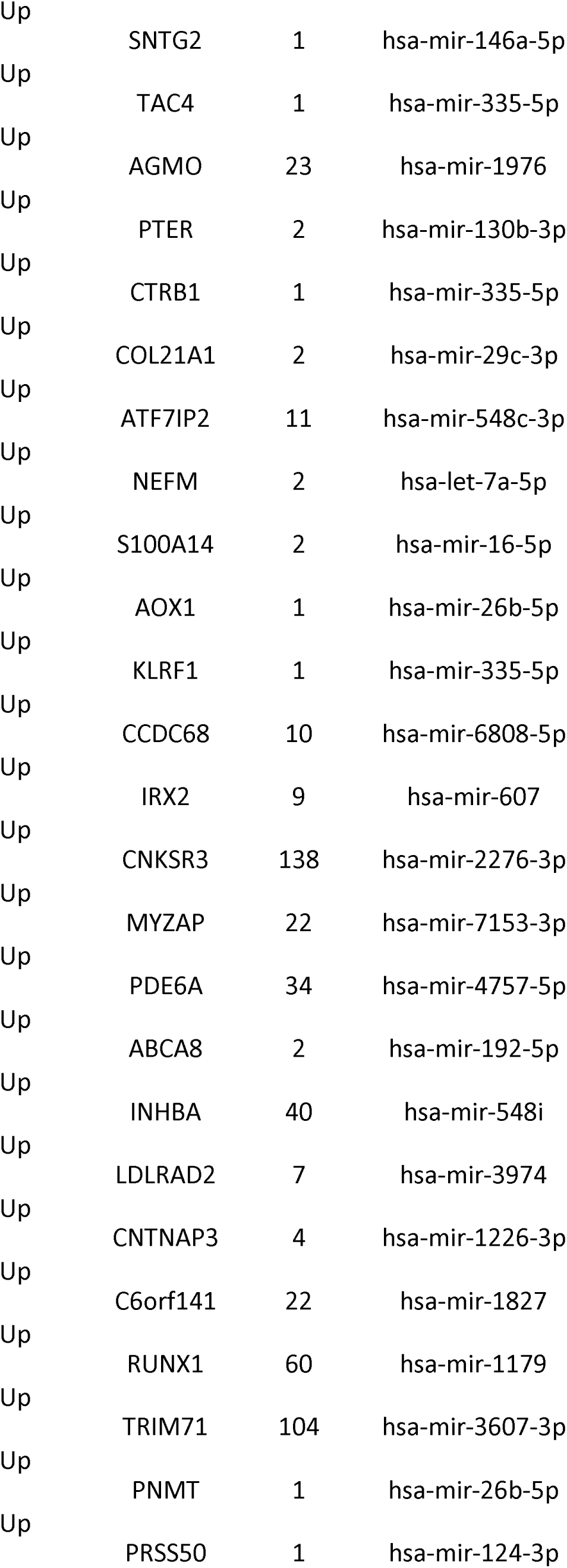
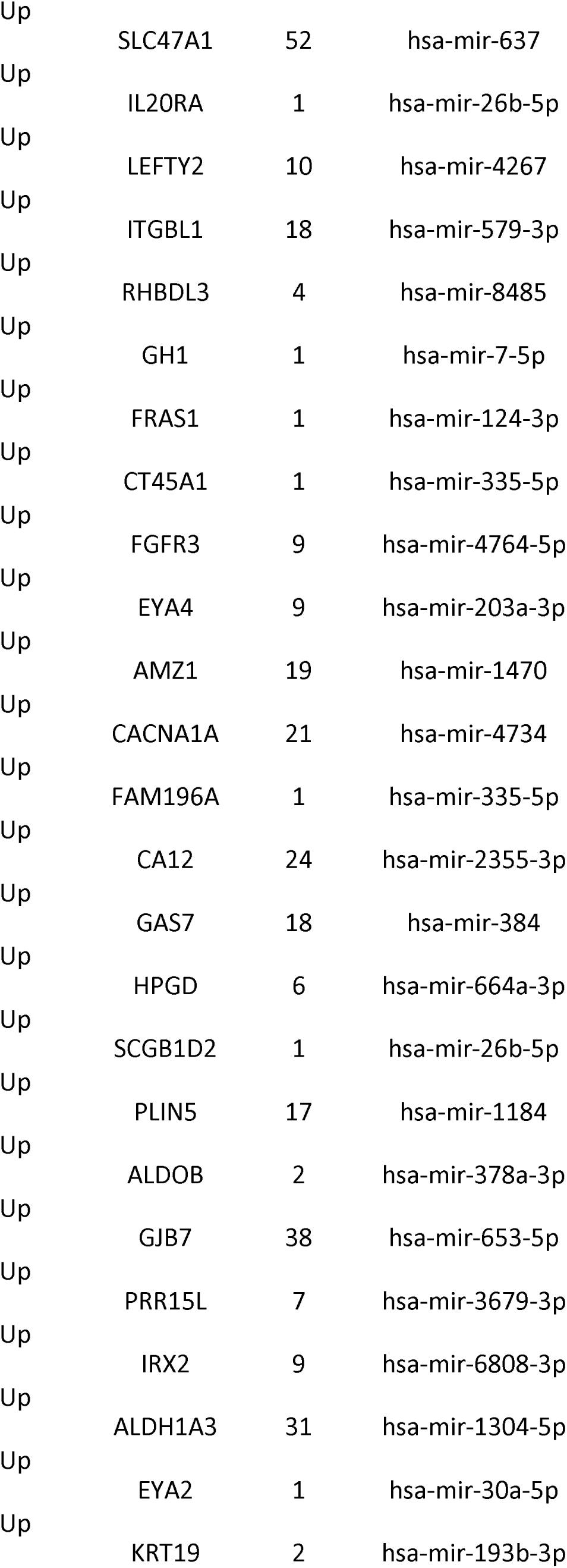
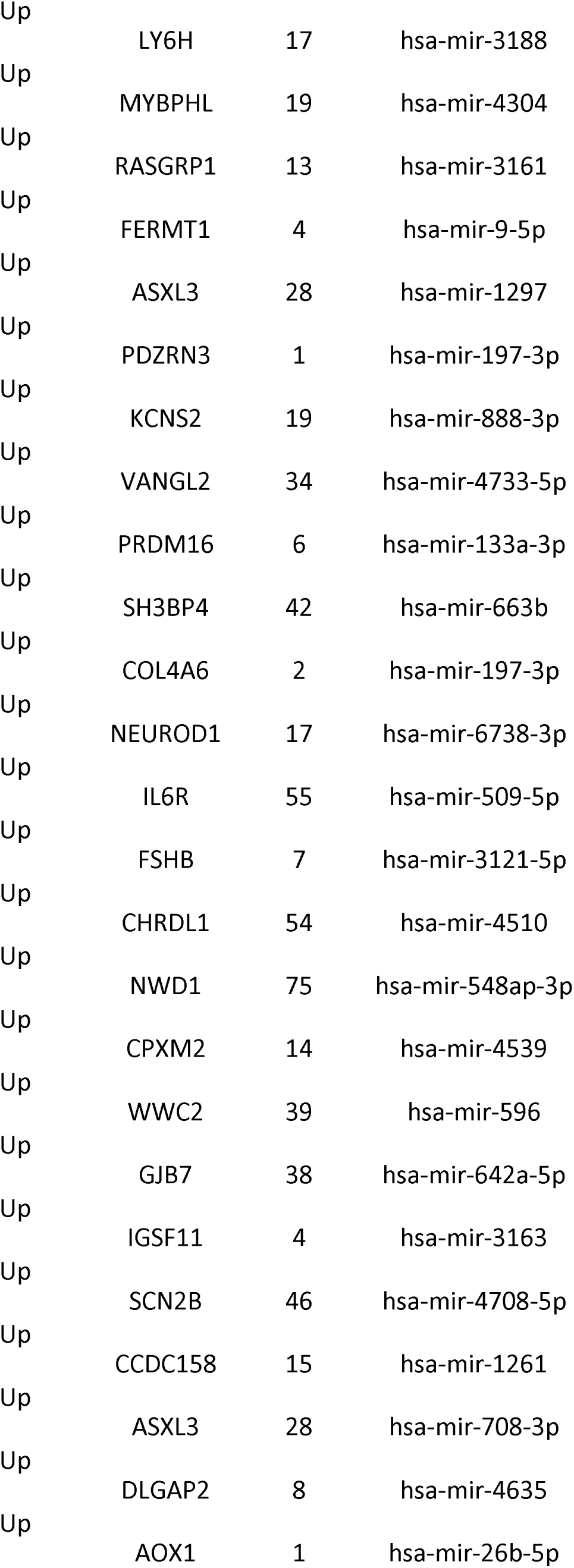
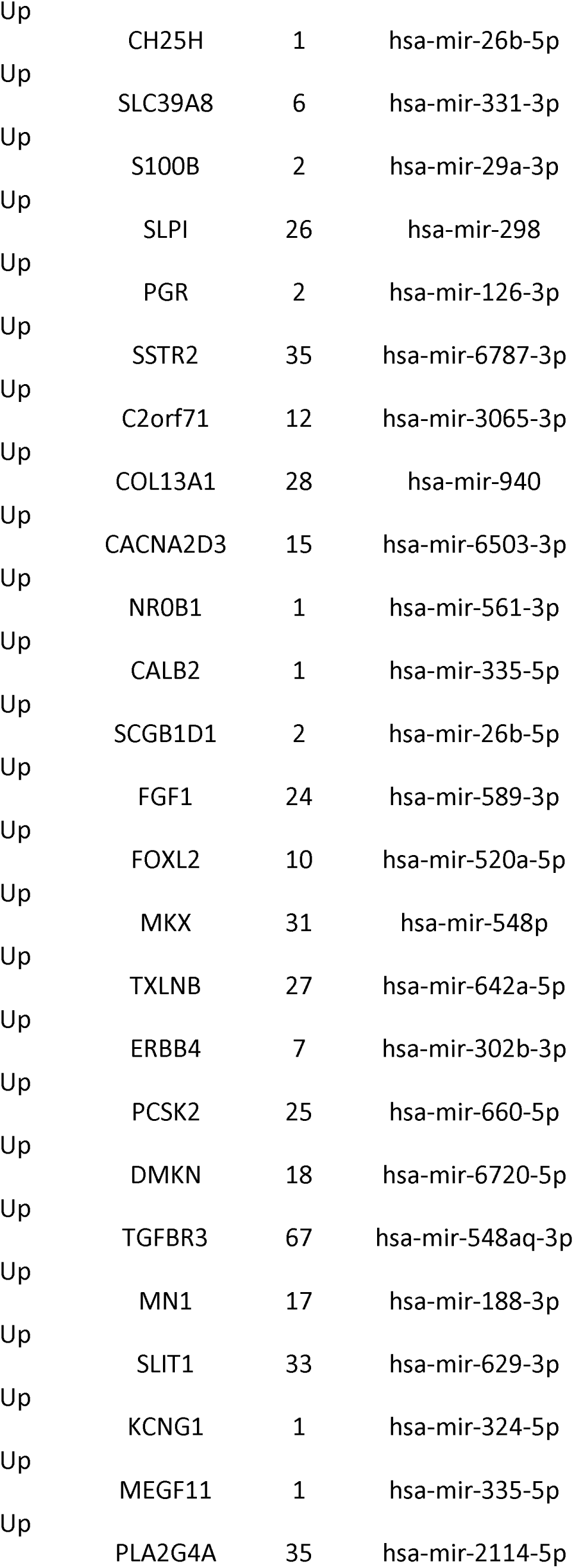
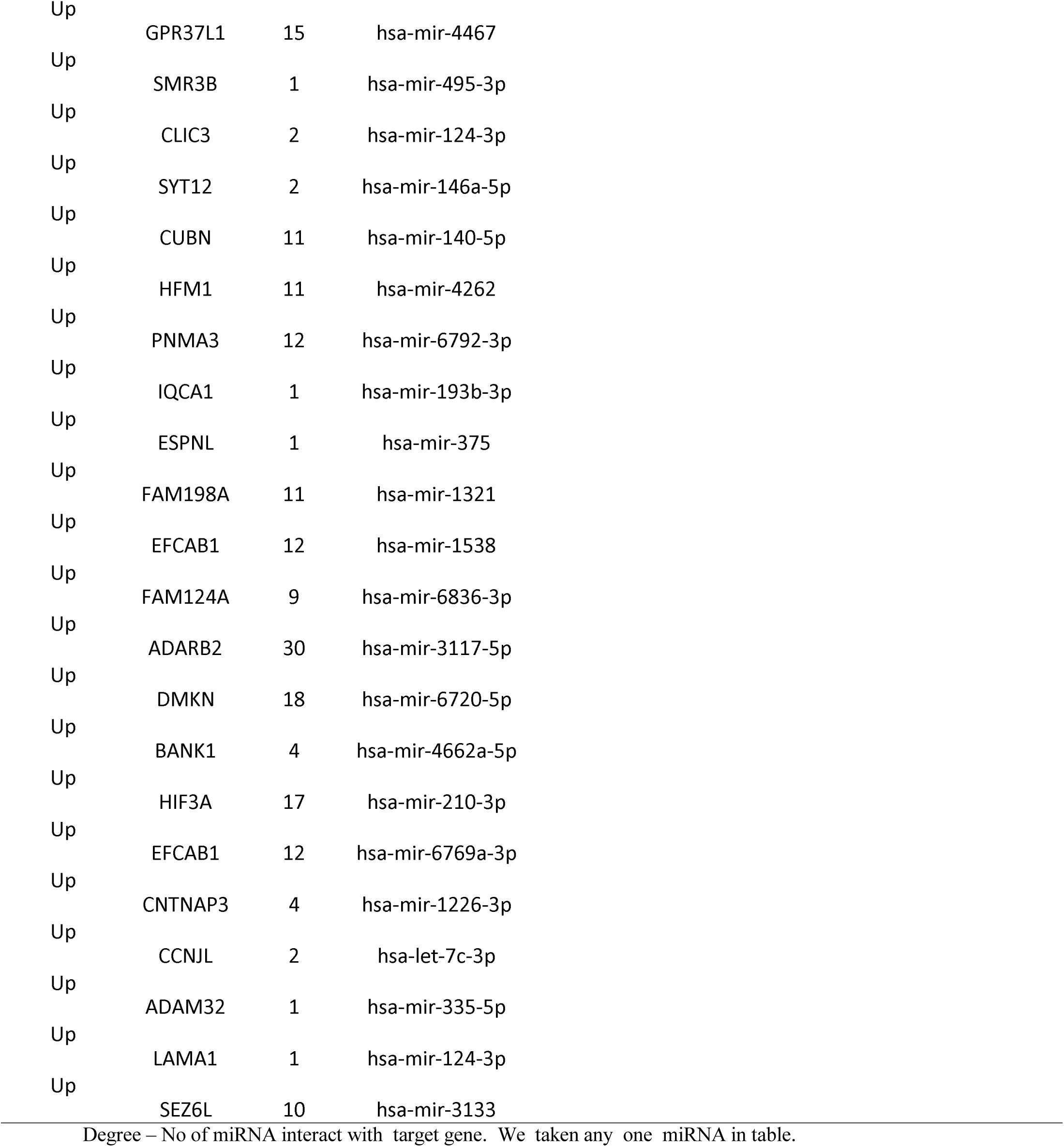
miRNA - target gene interaction table

| Regulation | target Genes | Degree | Top MicroRNA | Regulation | target Genes | Degree | Top MicroRNA |
| --- | --- | --- | --- | --- | --- | --- | --- |
| Up | SALL1 | 53 | hsa-mir-548b-3p | Down | TRIM24 | 21 | hsa-mir-4477a |
| Up | NKX2-2 | 20 | hsa-mir-6829-3p | Down | KLHL13 | 1 | hsa-mir-218-5p |
| Up | PTCHD1 | 60 | hsa-mir-548a-3p | Down | MAGEB2 | 2 | hsa-mir-98-5p |
| Up | BMP7 | 22 | hsa-mir-6890-3p | Down | C3orf80 | 10 | hsa-mir-3153 |
| Up | TMEM154 | 49 | hsa-mir-3187-3p | Down | PPBP | 1 | hsa-mir-335-5p |
| Up | SERTM1 | 13 | hsa-mir-3924 | Down | TTK | 7 | hsa-mir-376a-3p |
| Up | FAM83F | 94 | hsa-mir-612 | Down | TOP2A | 17 | hsa-mir-708-5p |
| Up | SGCD | 41 | hsa-mir-4761-5p | Down | KIF18A | 2 | hsa-mir-192-5p |
| Up | C1orf168 | 2 | hsa-mir-148b-3p | Down | TPX2 | 1 | hsa-mir-193b-3p |
| Up | GRHL2 | 1 | hsa-mir-26b-5p | Down | POC1A | 69 | hsa-mir-3605-5p |
| Up | SMR3B | 1 | hsa-mir-495-3p | Down | ARHGAP29 | 25 | hsa-mir-6814-5p |
| Up | SCRG1 | 33 | hsa-mir-1304-3p | Down | NRG1 | 1 | hsa-mir-124-3p |
| Up | IL20RA | 1 | hsa-mir-26b-5p | Down | CADPS2 | 4 | hsa-mir-30c-5p |
| Up | FCN2 | 8 | hsa-mir-8064 | Down | ACER2 | 46 | hsa-let-7d-5p |
| Up | DPPA4 | 1 | hsa-mir-26b-5p | Down | CCNF | 102 | hsa-mir-324-3p |
| Up | SPTLC3 | 32 | hsa-mir-206 | Down | GTSE1 | 2 | hsa-mir-484 |
| Up | ATP4A | 1 | hsa-mir-1289 | Down | SOX4 | 160 | hsa-mir-519a-3p |
| Up | SDK2 | 33 | hsa-mir-644a | Down | DLEU1 | 31 | hsa-mir-616-5p |
| Up | RSPO3 | 1 | hsa-mir-335-5p | Down | PLK1 | 17 | hsa-mir-3120-3p |
| Up | FGFR2 | 5 | hsa-mir-19b-1-5p | Down | HAUS6 | 14 | hsa-mir-5582-3p |
| Up | IQCA1 | 1 | hsa-mir-193b-3p | Down | BUB1 | 67 | hsa-mir-497-3p |
| Up | MGST1 | 2 | hsa-mir-652-3p | Down | SERINC2 | 1 | hsa-mir-335-5p |
| Up | FABP7 | 7 | hsa-mir-6823-3p | Down | KIF14 | 14 | hsa-mir-3133 |
| Up | PON3 | 1 | hsa-mir-335-5p | Down | VWA1 | 30 | hsa-mir-592 |
| Up | GAL | 8 | hsa-mir-4671-5p | Down | FRMPD3 | 18 | hsa-mir-6511a-3p |
| Up | PAX6 | 23 | hsa-mir-599 | Down | SEL1L | 12 | hsa-mir-548as-3p |
| Up | SYPL2 | 1 | hsa-mir-615-3p | Down | NUSAP1 | 1 | hsa-let-7b-5p |
| Up | SLC15A1 | 19 | hsa-mir-7977 | Down | DOK6 | 23 | hsa-mir-3679-3p |
| Up | TMPRSS3 | 2 | hsa-mir-204-5p | Down | NRG1 | 1 | hsa-mir-124-3p |
| Up | ROR2 | 3 | hsa-mir-451a | Down | DHX15 | 7 | hsa-mir-221-3p |
| Up | DACT2 | 2 | hsa-mir-128-3p | Down | NEK2 | 2 | hsa-mir-92a-3p |
| Up | SMTNL2 | 43 | hsa-mir-3652 | Down | TAS2R7 | 1 | hsa-mir-142-3p |
| Up | FBN2 | 16 | hsa-mir-23c | Down | SPC24 | 26 | hsa-mir-3664-5p |
| Up | NTF4 | 1 | hsa-mir-124-3p | Down | HMMR | 5 | hsa-mir-106b-5p |
| Up | SRL | 2 | hsa-mir-18a-3p | Down | PTP4A1 | 132 | hsa-mir-186-5p |
| Up | SLIT1 | 33 | hsa-mir-5787 | Down | DAZAP2 | 109 | hsa-mir-511-5p |
| Up | BBOX1 | 4 | hsa-mir-4477a | Down | CDCA5 | 4 | hsa-mir-331-3p |
| Up | ART3 | 2 | hsa-mir-122-5p | Down | ANTXR2 | 60 | hsa-mir-3065-5p |
| Up | NRK | 1 | hsa-mir-335-5p | Down | FBXL5 | 31 | hsa-mir-4731-3p |
| Up | ID4 | 91 | hsa-mir-4311 | Down | PLA2G2D | 20 | hsa-mir-548au-3p |
| Up | PLIN4 | 1 | hsa-mir-335-5p | Down | RBPJL | 25 | hsa-mir-6731-5p |
| Up | TRDN | 17 | hsa-mir-501-3p | Down | CENPI | 2 | hsa-mir-192-5p |
| Up | CLIC5 | 54 | hsa-mir-4480 | Down | NLGN1 | 8 | hsa-mir-670-3p |
| Up | BTBD17 | 1 | hsa-mir-335-5p | Down | POU3F2 | 19 | hsa-mir-6781-5p |
| Up | CCKBR | 3 | hsa-mir-148a-3p | Down | PAX4 | 8 | hsa-mir-4783-5p |
| Up | S100A1 | 4 | hsa-mir-138-5p | Down | ASXL2 | 48 | hsa-mir-5002-5p |
| Up | CBLN2 | 6 | hsa-mir-6819-3p | Down | GRIK2 | 1 | hsa-mir-26b-5p |
| Up | PMAIP1 | 128 | hsa-mir-411-3p | Down | NCAPG | 20 | hsa-mir-664b-3p |
| Up | XDH | 1 | hsa-mir-26b-5p | Down | SLC19A1 | 9 | hsa-mir-4422 |
| Up | TRIM29 | 18 | hsa-mir-1268a | Down | EPHB1 | 18 | hsa-mir-548aw |
| Up | UGT2A2 | 9 | hsa-mir-548u | Down | INHBE | 2 | hsa-mir-98-5p |
| Up | SOX5 | 38 | hsa-mir-555 | Down | FIBCD1 | 12 | hsa-mir-6717-5p |
| Up | S100A1 | 4 | hsa-mir-132-3p | Down | RGS5 | 42 | hsa-mir-4255 |
| Up | GALR1 | 8 | hsa-mir-130b-5p | Down | GUCA2B | 1 | hsa-mir-335-5p |
| Up | GAS1 | 33 | hsa-mir-449b-5p | Down | CCNB2 | 6 | hsa-let-7f-5p |
| Up | PPP1R17 | 1 | hsa-mir-335-5p | Down | ADAMTSL1 | 2 | hsa-mir-25-3p |
| Up | IFNE | 16 | hsa-mir-4786-5p | Down | FAM83A | 5 | hsa-mir-5090 |
| Up | CELSR1 | 2 | hsa-mir-92a-3p | Down | LRFN1 | 18 | hsa-mir-6887-3p |
| Up | C4orf19 | 5 | hsa-mir-1273f | Down | SLC25A15 | 20 | hsa-mir-6720-3p |
| Up | POMC | 2 | hsa-mir-488-3p | Down | GABRQ | 1 | hsa-mir-335-5p |
| Up | SLC6A14 | 1 | hsa-mir-26b-5p | Down | PRC1 | 17 | hsa-mir-3977 |
| Up | LRAT | 47 | hsa-mir-4770 | Down | AMIGO2 | 2 | hsa-mir-155-5p |
| Up | SPTLC3 | 32 | hsa-mir-6514-5p | Down | CDC45 | 2 | hsa-mir-575 |
| Up | KLK11 | 1 | hsa-mir-335-5p | Down | PRDM11 | 1 | hsa-mir-335-5p |
| Up | F5 | 4 | hsa-mir-634 | Down | CYP27B1 | 12 | hsa-mir-1178-5p |
| Up | CNTNAP2 | 14 | hsa-mir-571 | Down | KIF11 | 15 | hsa-mir-4495 |
| Up | PI16 | 2 | hsa-mir-1-1 | Down | G2E3 | 54 | hsa-mir-520g-3p |
| Up | ENPP6 | 24 | hsa-mir-6077 | Down | POLQ | 86 | hsa-mir-664a-5p |
| Up | GBP6 | 24 | hsa-mir-548j-3p | Down | GRIK2 | 1 | hsa-mir-26b-5p |
| Up | NEURL3 | 1 | hsa-mir-335-5p | Down | RASSF3 | 3 | hsa-mir-20a-3p |
| Up | CLIC5 | 54 | hsa-mir-4301 | Down | PBK | 4 | hsa-mir-4699-3p |
| Up | RASL11B | 7 | hsa-mir-3137 | Down | CENPA | 23 | hsa-mir-4719 |
| Up | KCNJ12 | 24 | hsa-mir-770-5p | Down | LYNX1 | 1 | hsa-mir-744-5p |
| Up | AQP4 | 1 | hsa-mir-320a | Down | EVX2 | 9 | hsa-mir-33b-3p |
| Up | P2RY2 | 16 | hsa-mir-4433b-3p | Down | KRTAP5-7 | 1 | hsa-mir-26b-5p |
| Up | IL22RA1 | 1 | hsa-mir-26b-5p | Down | DLGAP5 | 21 | hsa-mir-520e |
| Up | RNF207 | 10 | hsa-mir-3190-5p | Down | EDN1 | 57 | hsa-mir-379-5p |
| Up | ADAMTS18 | 16 | hsa-mir-588 | Down | KIF20A | 5 | hsa-mir-23a-3p |
| Up | NGFR | 23 | hsa-mir-6732-5p | Down | CNKSR2 | 10 | hsa-mir-6808-3p |
| Up | KCNJ16 | 1 | hsa-mir-148b-3p | Down | GOLGA6A | 4 | hsa-mir-450b-5p |
| Up | GPHA2 | 24 | hsa-mir-1180-5p | Down | B4GALT6 | 19 | hsa-mir-1248 |
| Up | C6orf132 | 48 | hsa-mir-4638-5p | Down | ARHGAP42 | 22 | hsa-mir-6131 |
| Up | PHACTR2 | 47 | hsa-mir-590-3p | Down | KRT28 | 1 | hsa-mir-1229-3p |
| Up | EVC2 | 18 | hsa-mir-6873-3p | Down | CDKN3 | 18 | hsa-mir-4695-3p |
| Up | FOXL2 | 10 | hsa-mir-1253 | Down | DCAF4L2 | 2 | hsa-mir-494-3p |
| Up | FAM3B | 1 | hsa-mir-335-5p |  |  |  |  |
| Up | DLK1 | 1 | hsa-mir-15a-5p |  |  |  |  |
| Up | BLID | 1 | hsa-mir-335-5p |  |  |  |  |
| Up | UGT2A1 | 10 | hsa-mir-4762-5p |  |  |  |  |
| Up | OGN | 1 | hsa-mir-526b-5p |  |  |  |  |
| Up | TF | 2 | hsa-mir-19a-3p |  |  |  |  |
| Up | SMAD9 | 45 | hsa-mir-4419b |  |  |  |  |
| Up | LINC00598 | 209 | hsa-mir-890 |  |  |  |  |
| Up | SOX2 | 15 | hsa-mir-126-3p |  |  |  |  |
| Up | KLK12 | 2 | hsa-mir-1-3p |  |  |  |  |
| Up | KRT14 | 2 | hsa-mir-124-3p |  |  |  |  |
| Up | DLGAP2 | 8 | hsa-mir-20a-3p |  |  |  |  |
| Up | LRRC2 | 46 | hsa-mir-3914 |  |  |  |  |
| Up | AMOT | 23 | hsa-mir-500a-5p |  |  |  |  |

|  |  |  |  |
| --- | --- | --- | --- |
| Up | LINC00598 | 209 | hsa-mir-518a-5p |
| Up | HSD11B1 | 2 | hsa-mir-26b-5p |
| Up | SGCD | 41 | hsa-mir-4698 |
| Up | PRSS8 | 1 | hsa-mir-335-5p |
| Up | C6orf118 | 2 | hsa-mir-1-1 |
| Up | CRISP2 | 2 | hsa-mir-27a-3p |
| Up | IHH | 22 | hsa-mir-6839-5p |
| Up | MCOLN2 | 18 | hsa-mir-6731-3p |
| Up | SLC39A12 | 1 | hsa-mir-335-5p |
| Up | VIT | 1 | hsa-mir-124-3p |
| Up | PRRX2 | 1 | hsa-mir-124-3p |
| Up | PPP1R17 | 1 | hsa-mir-335-5p |
| Up | TSHB | 1 | hsa-mir-26b-5p |
| Up | AGR2 | 2 | hsa-mir-197-3p |
| Up | KBTBD12 | 39 | hsa-mir-4675 |
| Up | GADD45G | 1 | hsa-mir-181a-5p |
| Up | LIX1 | 3 | hsa-mir-4668-5p |
| Up | PRSS45 | 8 | hsa-mir-765 |
| Up | MFAP4 | 1 | hsa-mir-124-3p |
| Up | ESPNL | 1 | hsa-mir-375 |
| Up | NFIX | 67 | hsa-mir-4488 |
| Up | EVC2 | 18 | hsa-mir-1236-3p |
| Up | COL8A2 | 2 | hsa-mir-124-3p |
| Up | GNRHR | 4 | hsa-mir-1-3p |
| Up | SULT1C2 | 2 | hsa-mir-7-5p |
| Up | CA8 | 54 | hsa-mir-646 |
| Up | CHRD1 | 54 | hsa-mir-583 |
| Up | RPRM | 32 | hsa-mir-939-5p |
| Up | OCA2 | 12 | hsa-mir-1299 |
| Up | MPPED1 | 15 | hsa-mir-516a-3p |
| Up | OGN | 1 | hsa-mir-526b-5p |
| Up | GADD45G | 1 | hsa-mir-181a-5p |
| Up | VSNL1 | 25 | hsa-mir-6873-5p |
| Up | FGFR3 | 9 | hsa-mir-425-5p |
| Up | WNT5A | 29 | hsa-mir-4680-3p |
| Up | SFRP2 | 2 | hsa-mir-218-5p |
| Up | HSPB3 | 1 | hsa-mir-335-5p |
| Up | PAX6 | 23 | hsa-mir-223-3p |
| Up | CYBRD1 | 42 | hsa-mir-4517 |
| Up | PGR | 2 | hsa-mir-181a-5p |
| Up | HSD17B2 | 2 | hsa-mir-124-3p |
| Up | OPCML | 9 | hsa-mir-3149 |
| Up | AGR3 | 1 | hsa-mir-335-5p |
| Up | SLIT1 | 33 | hsa-mir-4726-5p |
| Up | SOX9 | 30 | hsa-mir-1323 |
| Up | CSHL1 | 1 | hsa-mir-16-5p |
| Up | CHST9 | 2 | hsa-mir-92a-3p |
| Up | NRK | 1 | hsa-mir-335-5p |
| Up | SULT1C2 | 2 | hsa-mir-7-5p |
| Up | SULT1C4 | 10 | hsa-mir-6833-3p |
| Up | SNTG2 | 1 | hsa-mir-146a-5p |
| Up | TAC4 | 1 | hsa-mir-335-5p |
| Up | AGMO | 23 | hsa-mir-1976 |
| Up | PTER | 2 | hsa-mir-130b-3p |
| Up | CTRB1 | 1 | hsa-mir-335-5p |
| Up | COL21A1 | 2 | hsa-mir-29c-3p |
| Up | ATF7IP2 | 11 | hsa-mir-548c-3p |
| Up | NEFM | 2 | hsa-let-7a-5p |
| Up | S100A14 | 2 | hsa-mir-16-5p |
| Up | AOX1 | 1 | hsa-mir-26b-5p |
| Up | KLRF1 | 1 | hsa-mir-335-5p |
| Up | CCDC68 | 10 | hsa-mir-6808-5p |
| Up | IRX2 | 9 | hsa-mir-607 |
| Up | CNKSR3 | 138 | hsa-mir-2276-3p |
| Up | MYZAP | 22 | hsa-mir-7153-3p |
| Up | PDE6A | 34 | hsa-mir-4757-5p |
| Up | ABCA8 | 2 | hsa-mir-192-5p |
| Up | INHBA | 40 | hsa-mir-548i |
| Up | LDLRAD2 | 7 | hsa-mir-3974 |
| Up | CNTNAP3 | 4 | hsa-mir-1226-3p |
| Up | C6orf141 | 22 | hsa-mir-1827 |
| Up | RUNX1 | 60 | hsa-mir-1179 |
| Up | TRIM71 | 104 | hsa-mir-3607-3p |
| Up | PNMT | 1 | hsa-mir-26b-5p |
| Up | PRSS50 | 1 | hsa-mir-124-3p |
| Up | SLC47A1 | 52 | hsa-mir-637 |
| Up | IL20RA | 1 | hsa-mir-26b-5p |
| Up | LEFTY2 | 10 | hsa-mir-4267 |
| Up | ITGBL1 | 18 | hsa-mir-579-3p |
| Up | RHBDL3 | 4 | hsa-mir-8485 |
| Up | GH1 | 1 | hsa-mir-7-5p |
| Up | FRAS1 | 1 | hsa-mir-124-3p |
| Up | CT45A1 | 1 | hsa-mir-335-5p |
| Up | FGFR3 | 9 | hsa-mir-4764-5p |
| Up | EYA4 | 9 | hsa-mir-203a-3p |
| Up | AMZ1 | 19 | hsa-mir-1470 |
| Up | CACNA1A | 21 | hsa-mir-4734 |
| Up | FAM196A | 1 | hsa-mir-335-5p |
| Up | CA12 | 24 | hsa-mir-2355-3p |
| Up | GAS7 | 18 | hsa-mir-384 |
| Up | HPGD | 6 | hsa-mir-664a-3p |
| Up | SCGB1D2 | 1 | hsa-mir-26b-5p |
| Up | PLIN5 | 17 | hsa-mir-1184 |
| Up | ALDOB | 2 | hsa-mir-378a-3p |
| Up | GJB7 | 38 | hsa-mir-653-5p |
| Up | PRR15L | 7 | hsa-mir-3679-3p |
| Up | IRX2 | 9 | hsa-mir-6808-3p |
| Up | ALDH1A3 | 31 | hsa-mir-1304-5p |
| Up | EYA2 | 1 | hsa-mir-30a-5p |
| Up | KRT19 | 2 | hsa-mir-193b-3p |
| Up | LY6H | 17 | hsa-mir-3188 |
| Up | MYBPHL | 19 | hsa-mir-4304 |
| Up | RASGRP1 | 13 | hsa-mir-3161 |
| Up | FERMT1 | 4 | hsa-mir-9-5p |
| Up | ASXL3 | 28 | hsa-mir-1297 |
| Up | PDZRN3 | 1 | hsa-mir-197-3p |
| Up | KCNS2 | 19 | hsa-mir-888-3p |
| Up | VANGL2 | 34 | hsa-mir-4733-5p |
| Up | PRDM16 | 6 | hsa-mir-133a-3p |
| Up | SH3BP4 | 42 | hsa-mir-663b |
| Up | COL4A6 | 2 | hsa-mir-197-3p |
| Up | NEUROD1 | 17 | hsa-mir-6738-3p |
| Up | IL6R | 55 | hsa-mir-509-5p |
| Up | FSHB | 7 | hsa-mir-3121-5p |
| Up | CHRD1 | 54 | hsa-mir-4510 |
| Up | NWD1 | 75 | hsa-mir-548ap-3p |
| Up | CPXM2 | 14 | hsa-mir-4539 |
| Up | WWC2 | 39 | hsa-mir-596 |
| Up | GJB7 | 38 | hsa-mir-642a-5p |
| Up | IGSF11 | 4 | hsa-mir-3163 |
| Up | SCN2B | 46 | hsa-mir-4708-5p |
| Up | CCDC158 | 15 | hsa-mir-1261 |
| Up | ASXL3 | 28 | hsa-mir-708-3p |
| Up | DLGAP2 | 8 | hsa-mir-4635 |
| Up | AOX1 | 1 | hsa-mir-26b-5p |
| Up | CH25H | 1 | hsa-mir-26b-5p |
| Up | SLC39A8 | 6 | hsa-mir-331-3p |
| Up | S100B | 2 | hsa-mir-29a-3p |
| Up | SLPI | 26 | hsa-mir-298 |
| Up | PGR | 2 | hsa-mir-126-3p |
| Up | SSTR2 | 35 | hsa-mir-6787-3p |
| Up | C2orf71 | 12 | hsa-mir-3065-3p |
| Up | COL13A1 | 28 | hsa-mir-940 |
| Up | CACNA2D3 | 15 | hsa-mir-6503-3p |
| Up | NR0B1 | 1 | hsa-mir-561-3p |
| Up | CALB2 | 1 | hsa-mir-335-5p |
| Up | SCGB1D1 | 2 | hsa-mir-26b-5p |
| Up | FGF1 | 24 | hsa-mir-589-3p |
| Up | FOXL2 | 10 | hsa-mir-520a-5p |
| Up | MKX | 31 | hsa-mir-548p |
| Up | TXLNB | 27 | hsa-mir-642a-5p |
| Up | ERBB4 | 7 | hsa-mir-302b-3p |
| Up | PCSK2 | 25 | hsa-mir-660-5p |
| Up | DMKN | 18 | hsa-mir-6720-5p |
| Up | TGFBR3 | 67 | hsa-mir-548aq-3p |
| Up | MN1 | 17 | hsa-mir-188-3p |
| Up | SLIT1 | 33 | hsa-mir-629-3p |
| Up | KCNG1 | 1 | hsa-mir-324-5p |
| Up | MEGF11 | 1 | hsa-mir-335-5p |
| Up | PLA2G4A | 35 | hsa-mir-2114-5p |
| Up | GPR37L1 | 15 | hsa-mir-4467 |
| Up | SMR3B | 1 | hsa-mir-495-3p |
| Up | CLIC3 | 2 | hsa-mir-124-3p |
| Up | SYT12 | 2 | hsa-mir-146a-5p |
| Up | CUBN | 11 | hsa-mir-140-5p |
| Up | HFM1 | 11 | hsa-mir-4262 |
| Up | PNMA3 | 12 | hsa-mir-6792-3p |
| Up | IQCA1 | 1 | hsa-mir-193b-3p |
| Up | ESPNL | 1 | hsa-mir-375 |
| Up | FAM198A | 11 | hsa-mir-1321 |
| Up | EFCAB1 | 12 | hsa-mir-1538 |
| Up | FAM124A | 9 | hsa-mir-6836-3p |
| Up | ADARB2 | 30 | hsa-mir-3117-5p |
| Up | DMKN | 18 | hsa-mir-6720-5p |
| Up | BANK1 | 4 | hsa-mir-4662a-5p |
| Up | HIF3A | 17 | hsa-mir-210-3p |
| Up | EFCAB1 | 12 | hsa-mir-6769a-3p |
| Up | CNTNAP3 | 4 | hsa-mir-1226-3p |
| Up | CCNJL | 2 | hsa-let-7c-3p |
| Up | ADAM32 | 1 | hsa-mir-335-5p |
| Up | LAMA1 | 1 | hsa-mir-124-3p |
| Up | SEZ6L | 10 | hsa-mir-3133 |

Degree – No of miRNA interact with target gene. We taken any one miRNA in table.

### Construction of the target gene **-** TF network

The TFs for target up and down regulated genes are shown in Fig. 14 and Fig. 15, respectively. Top 5 up regulated targeted genes such as IRX1 regulated by 73 TFs, CACNA2D3 regulated by 30 TFs, VSNL1 regulated by 29 TFs, BMP7 regulated by 28 TFs and DACT2 regulated by 27 TFs are given in Table 8. Top 5 down regulated targeted genes such as UNC13A regulated by 43 TFs, CCNF regulated by 35 TFs, BDNF regulated by 34 TFs, POU3F4 regulated by 34 TFs and AKAP3 regulated by 31 TFs are given in Table 8.

**Fig. 14.**
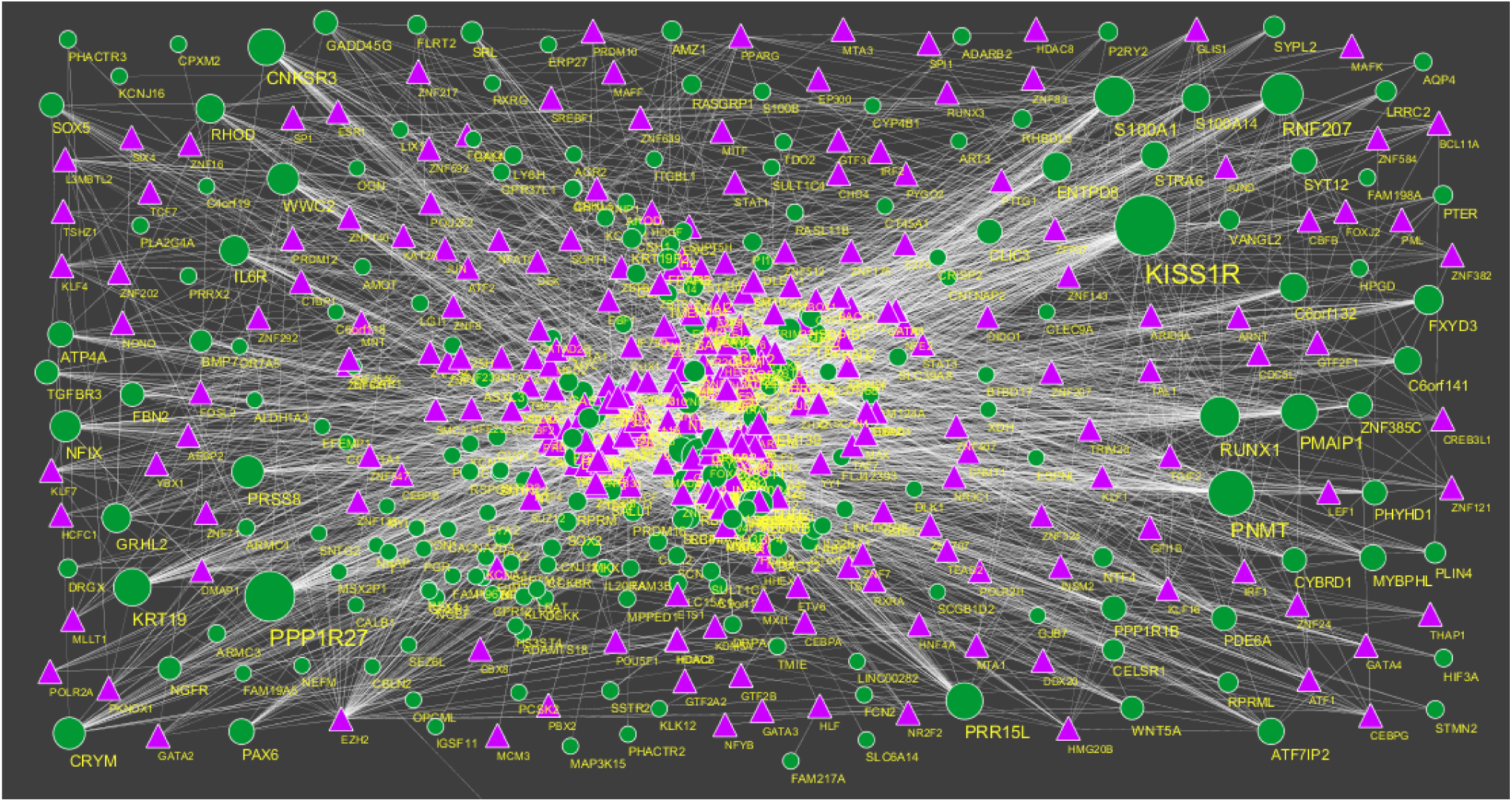
TF - gene network of predicted target up regulated genes. (Purple triangle - TFs and green circles-target up regulated genes)

**Fig. 15.**
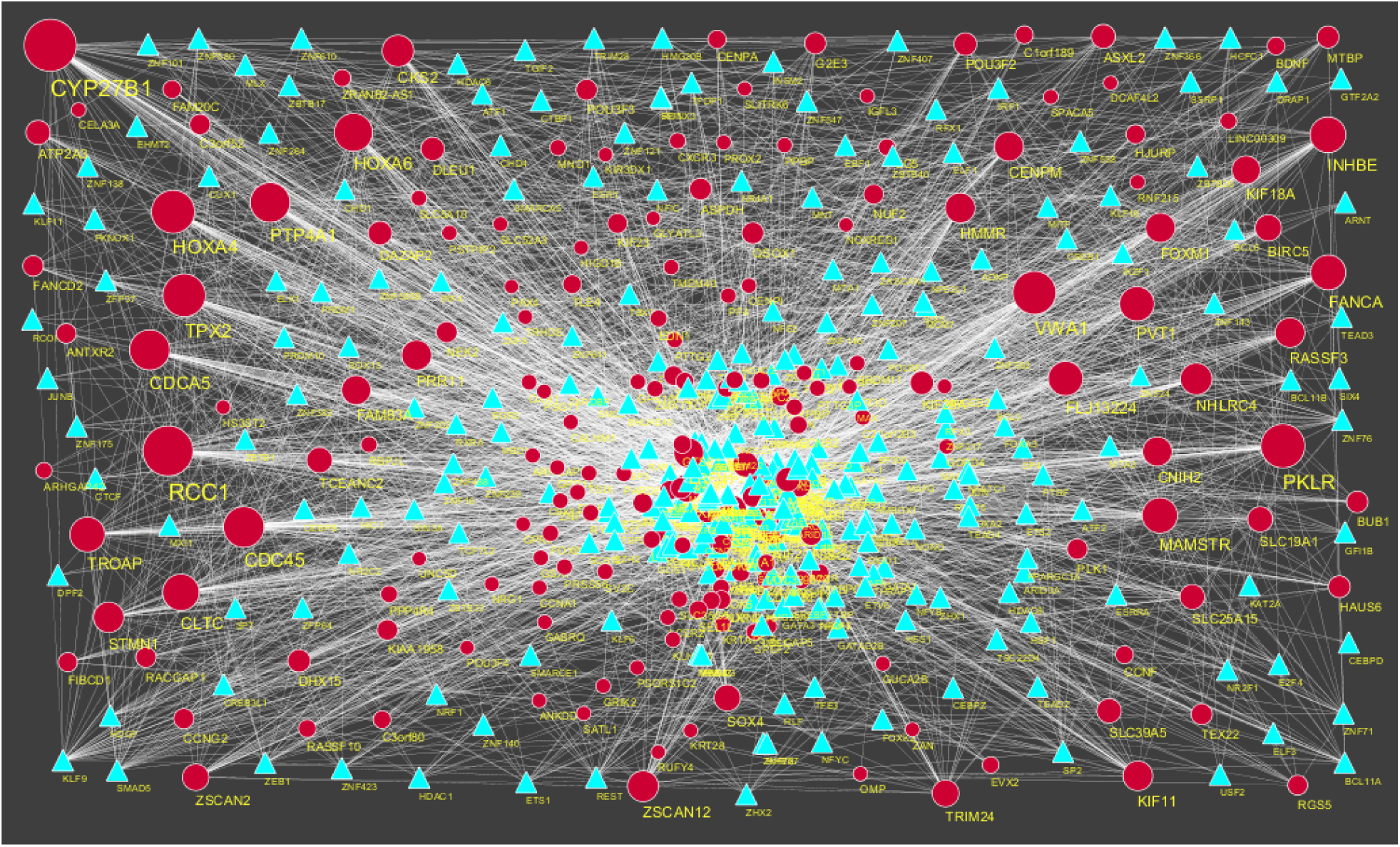
TF-gene network of predicted target up regulated genes. (Sky blue triangle - TFs and red circles-target up regulated genes)

**Table 8.**
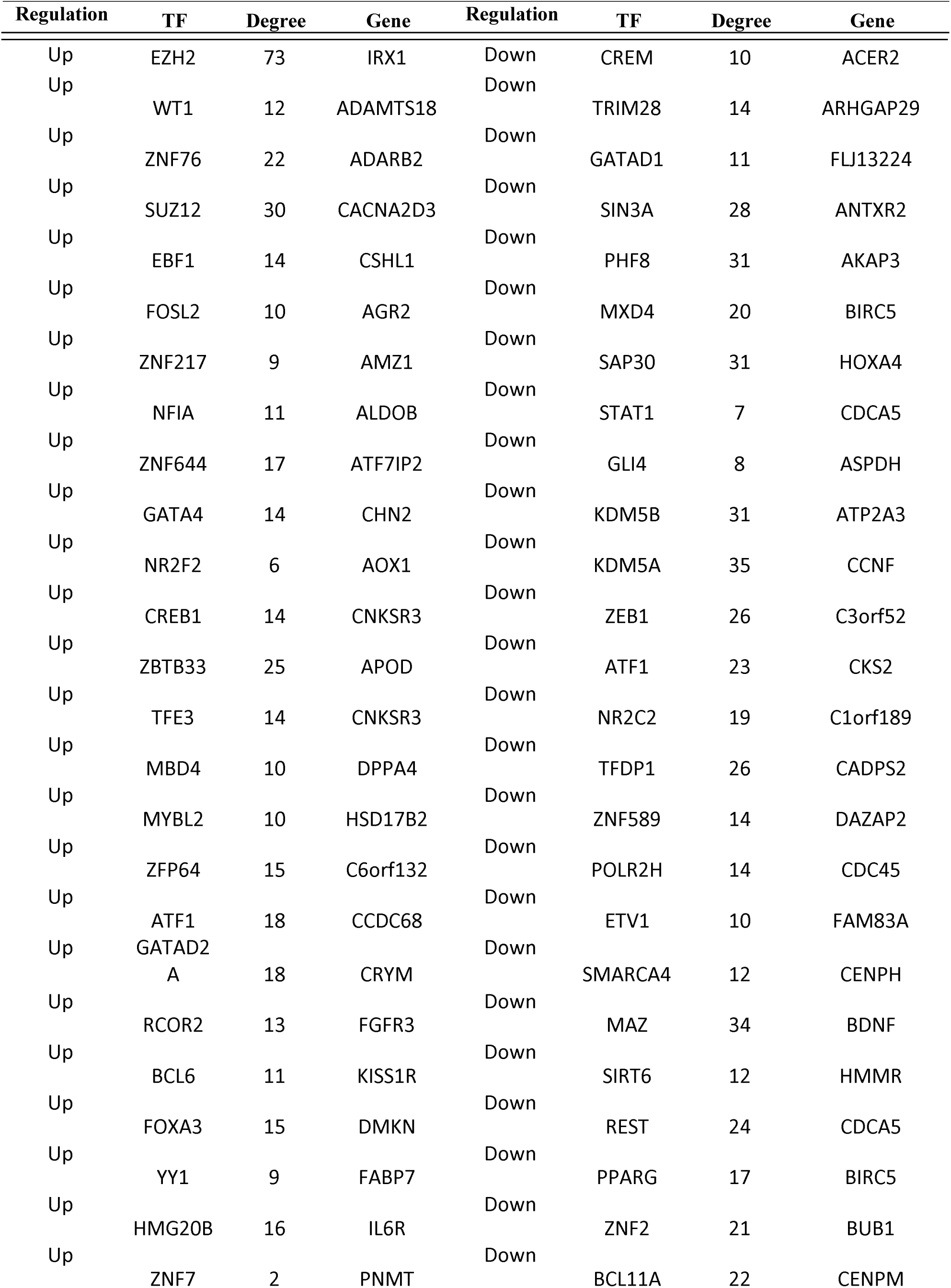
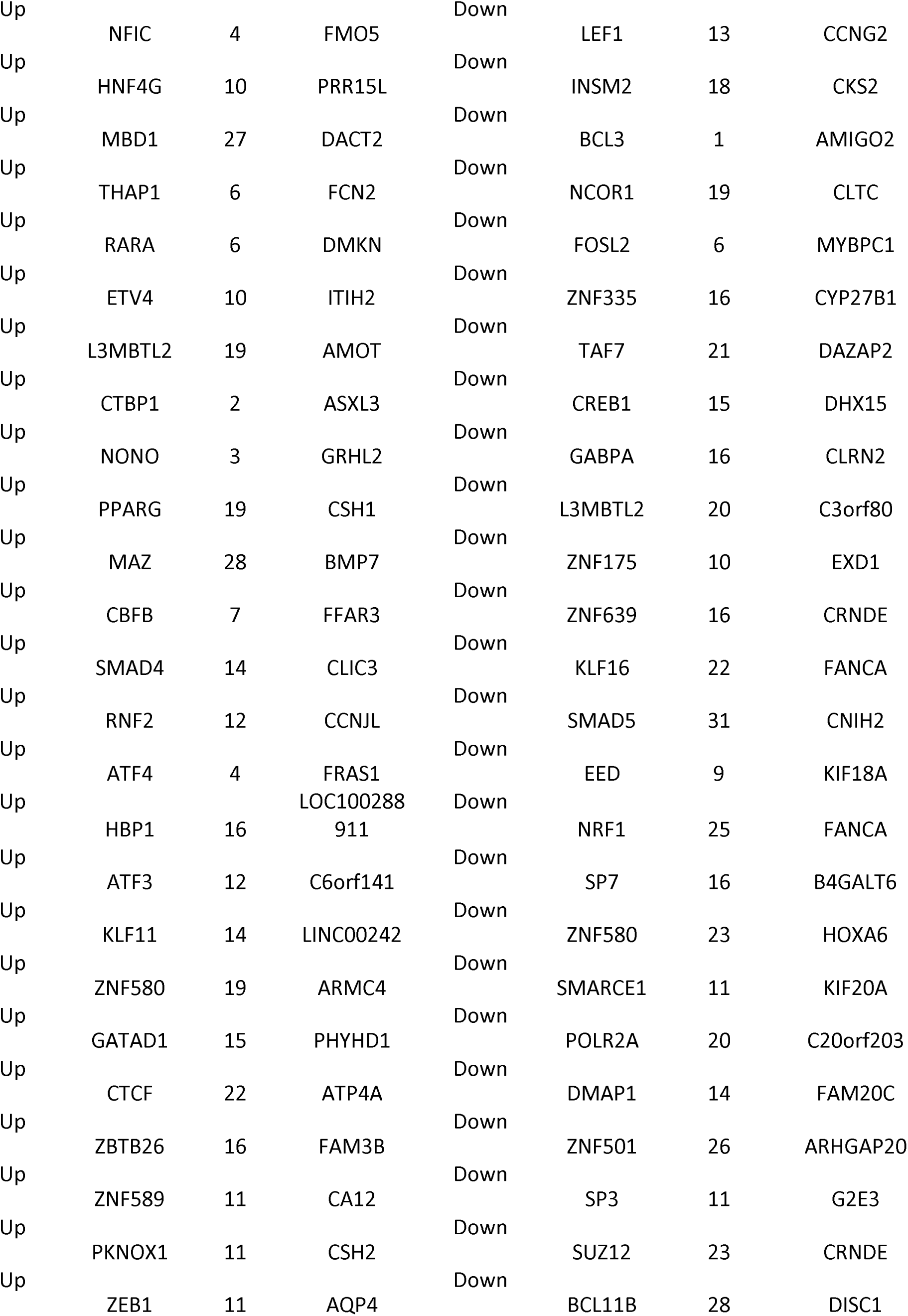
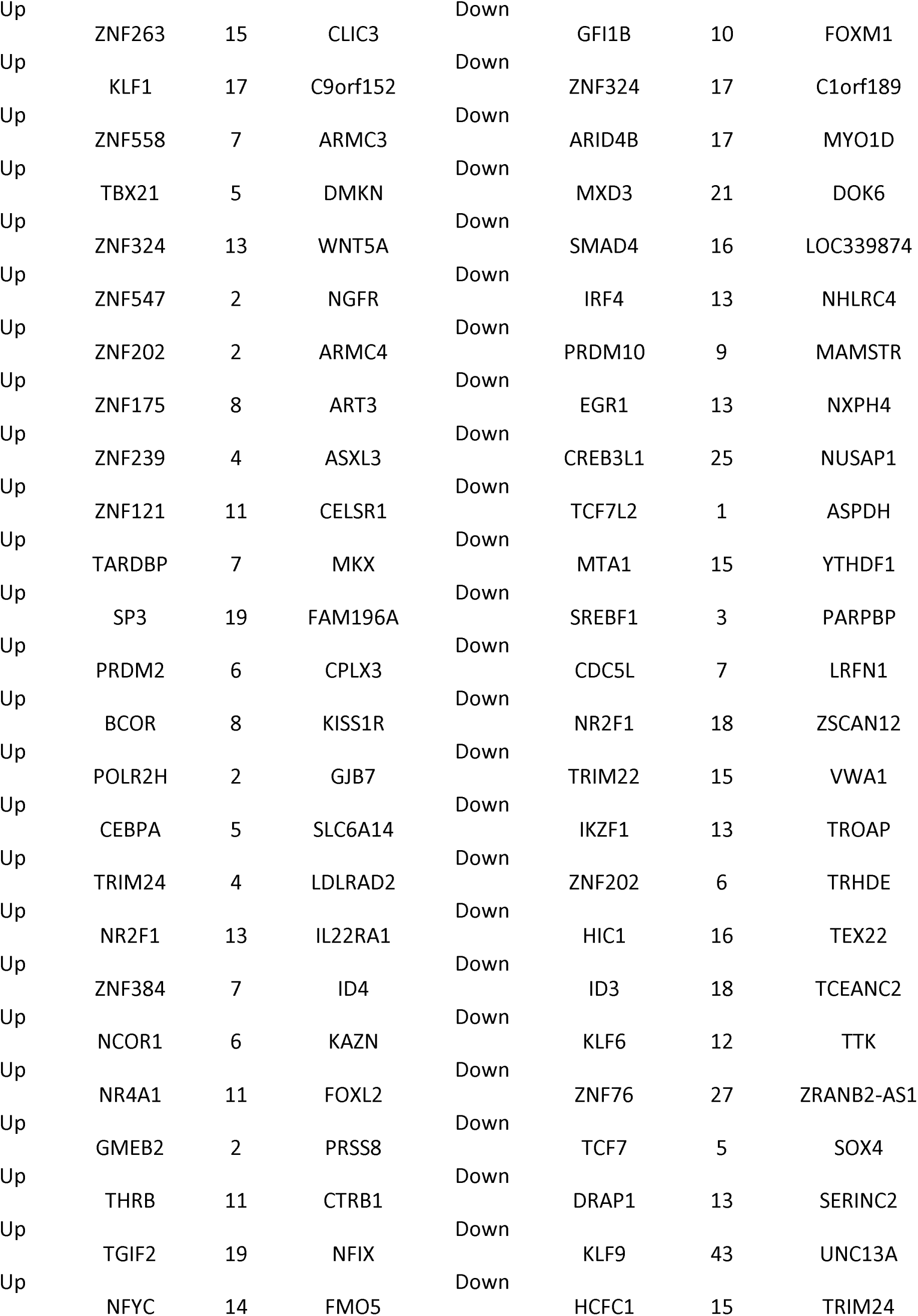
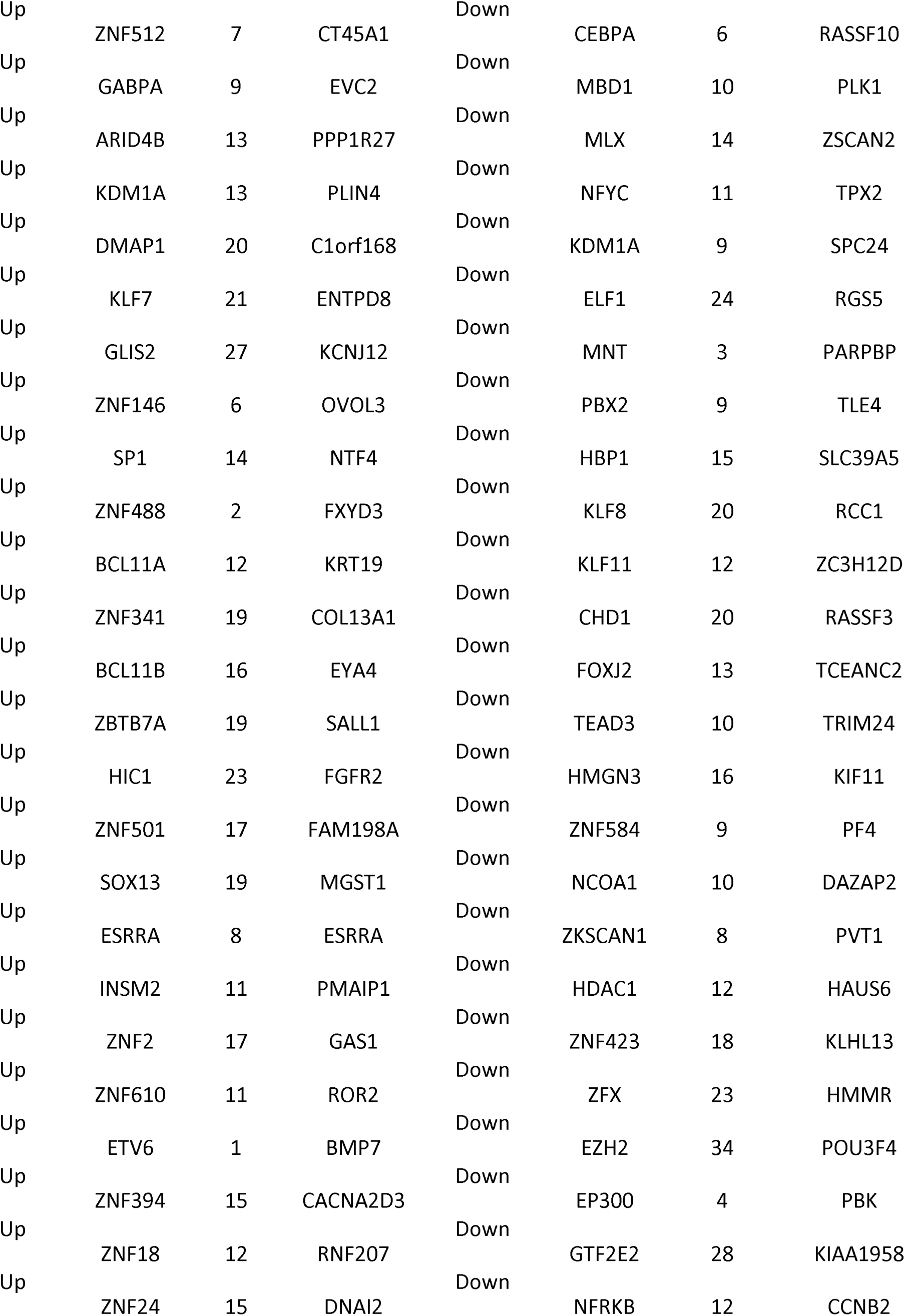
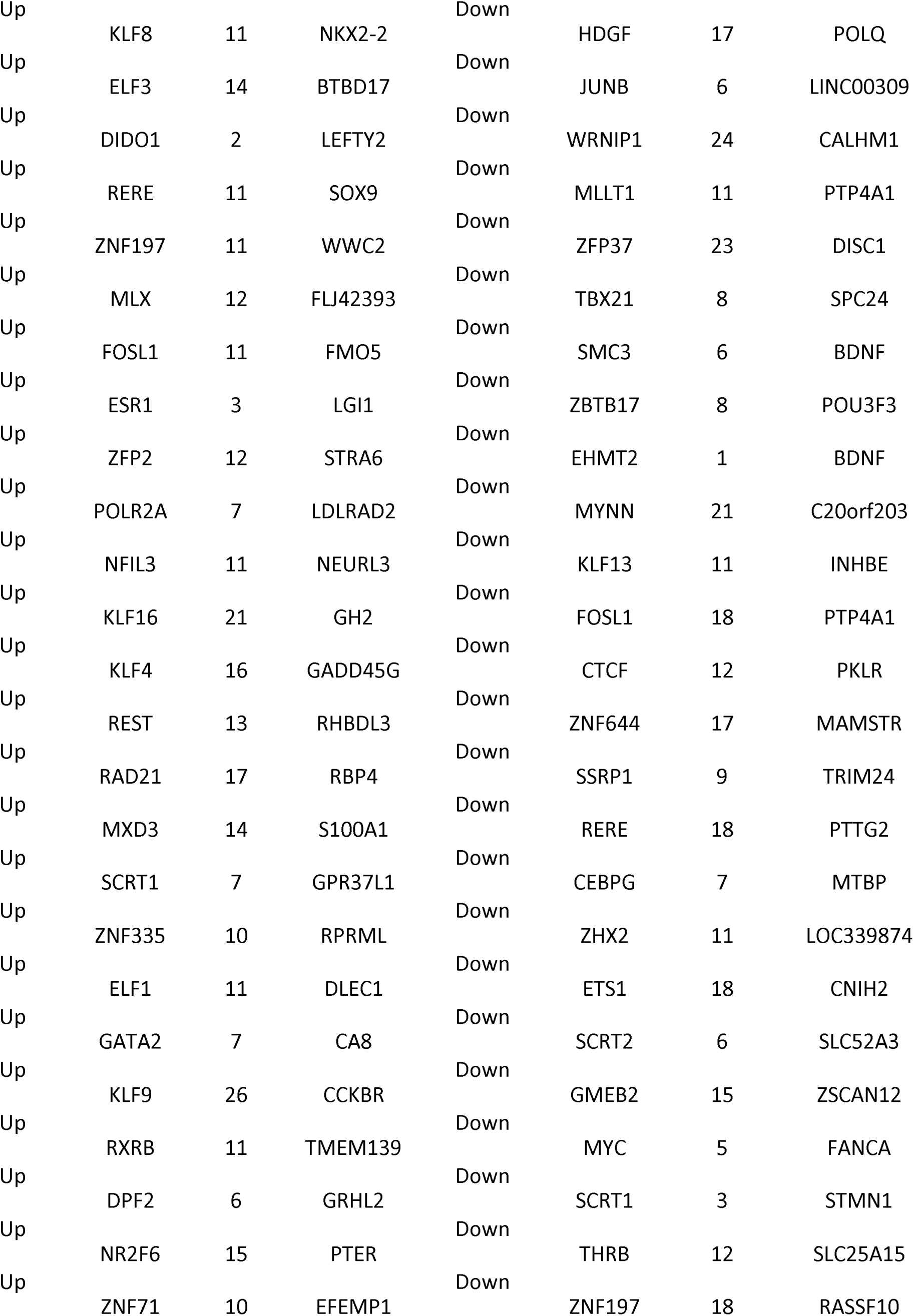
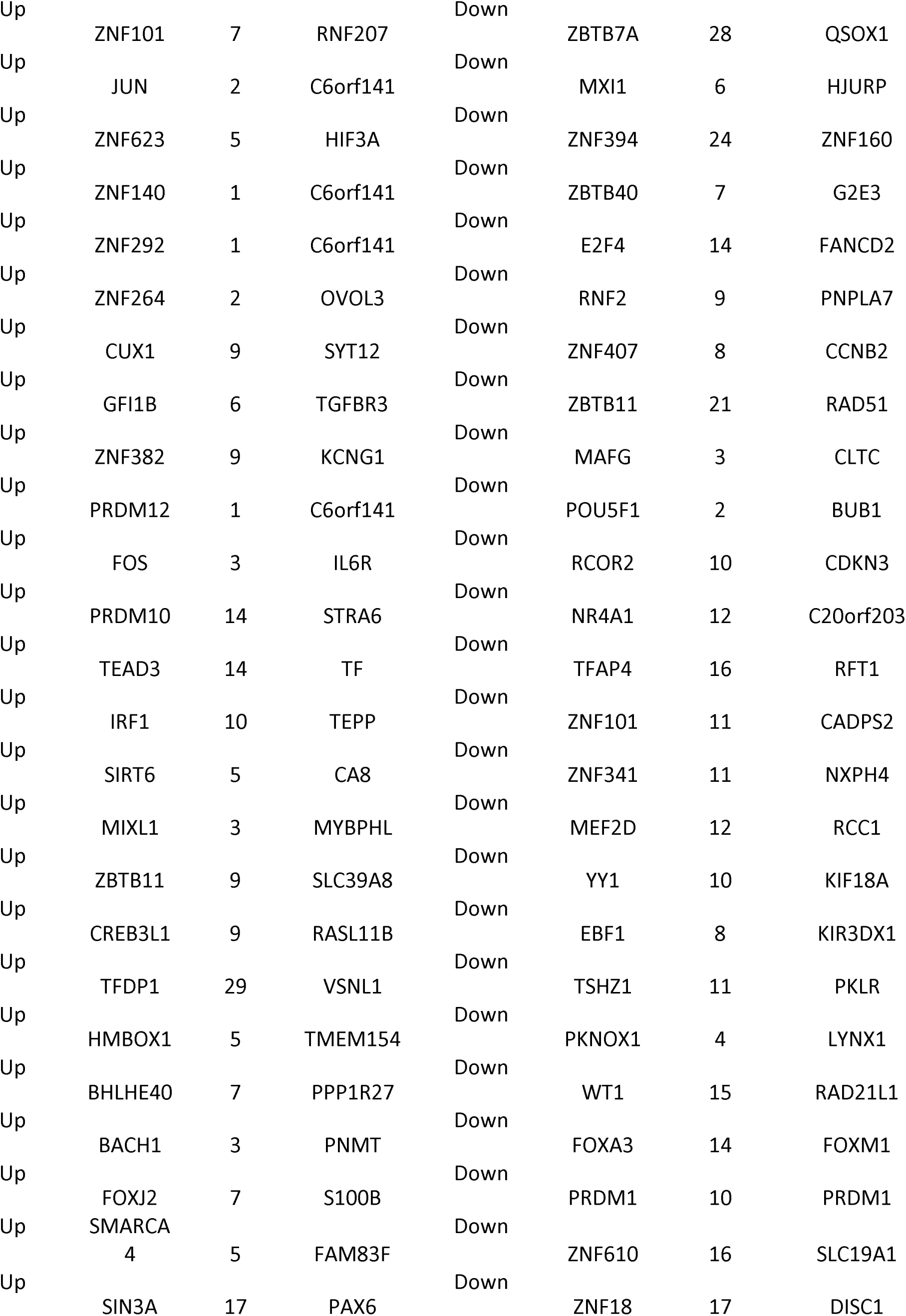
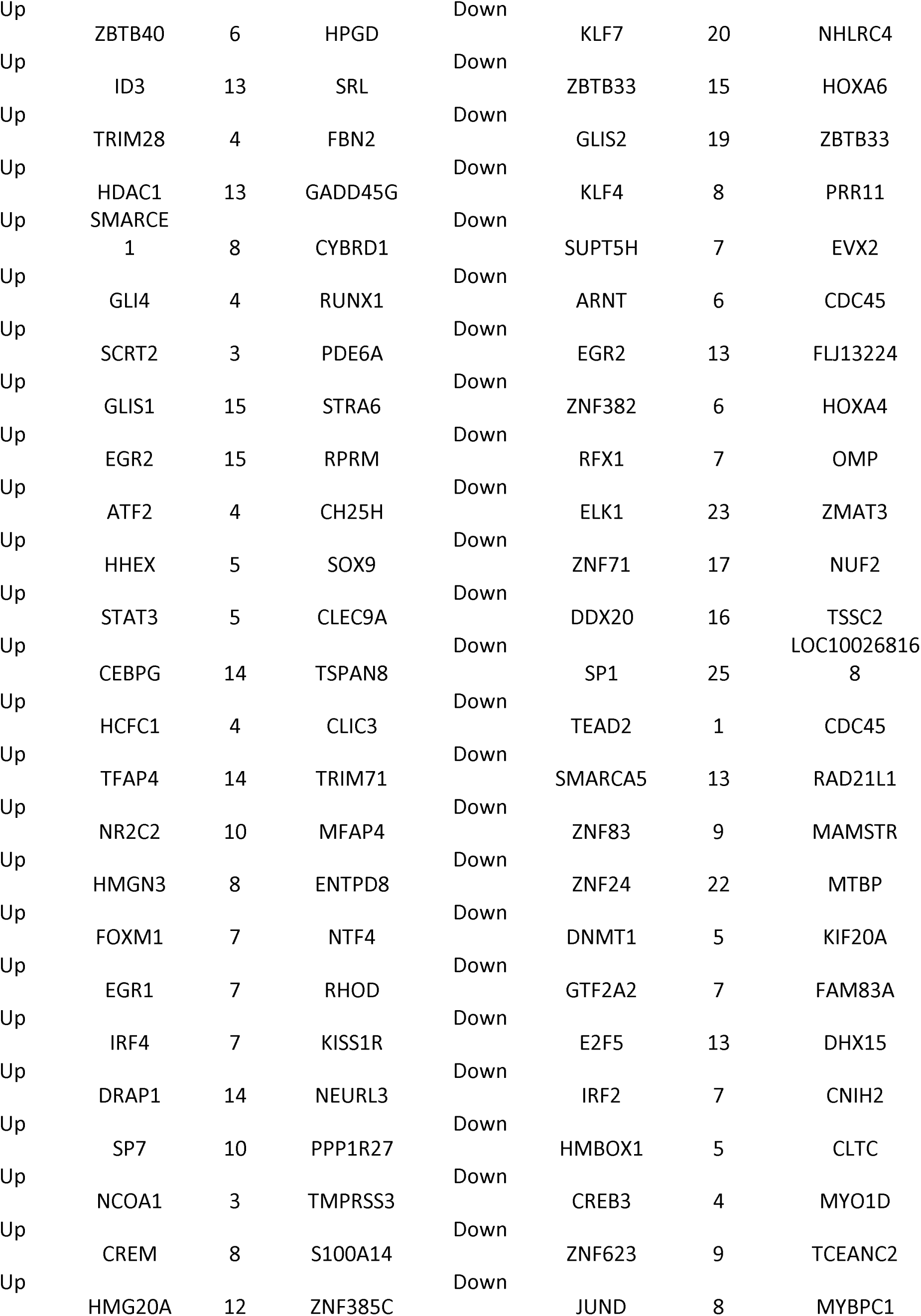
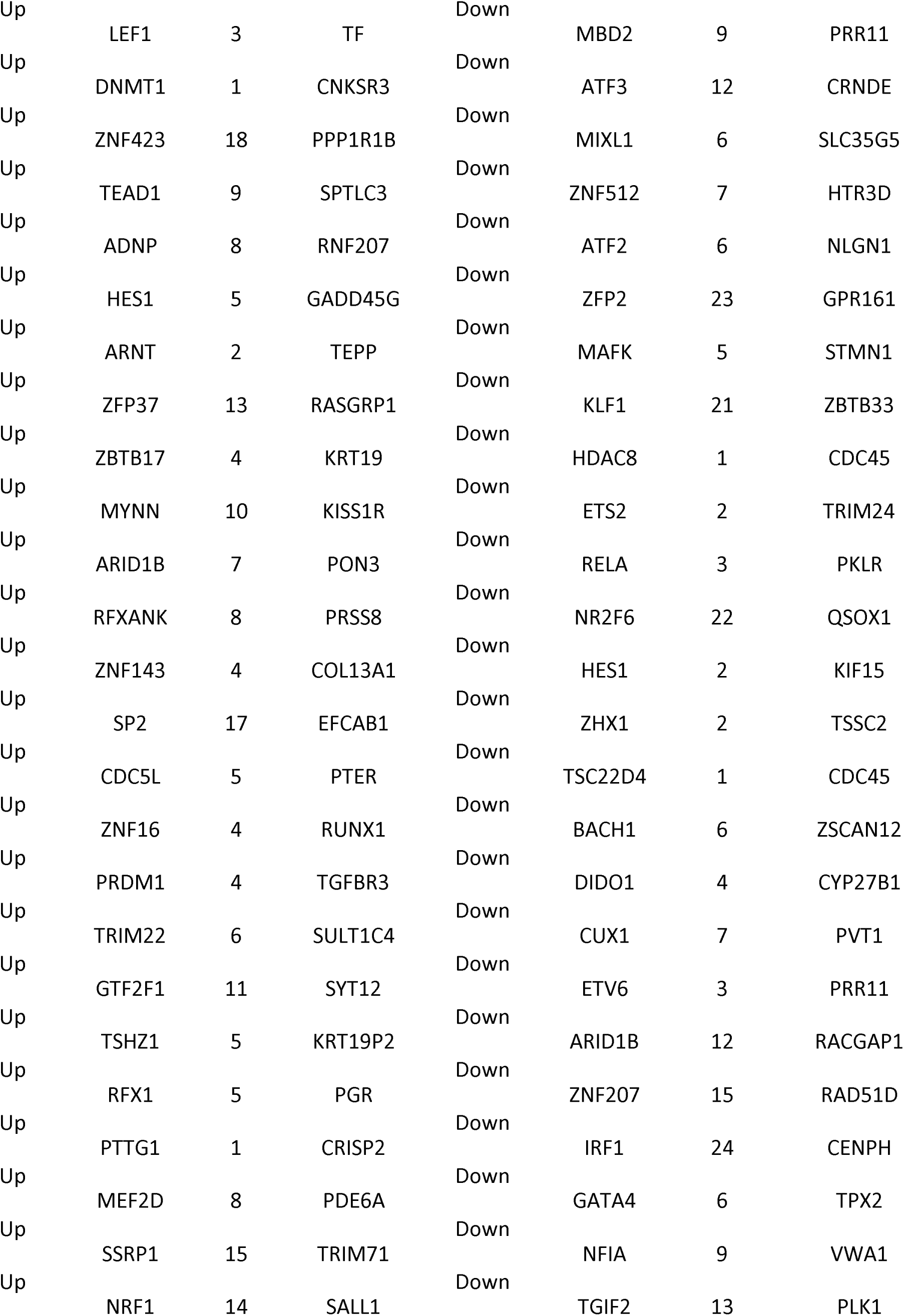
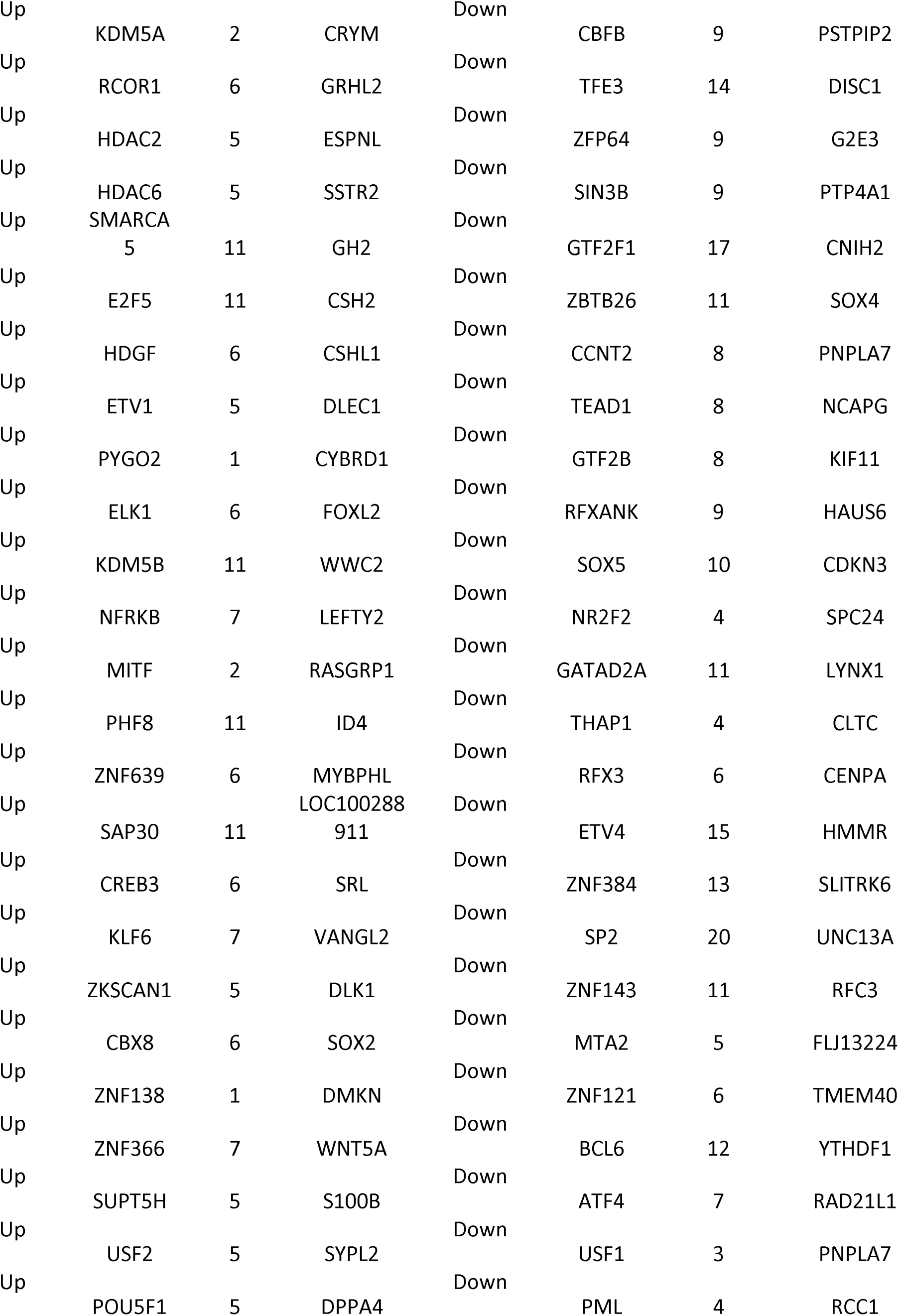
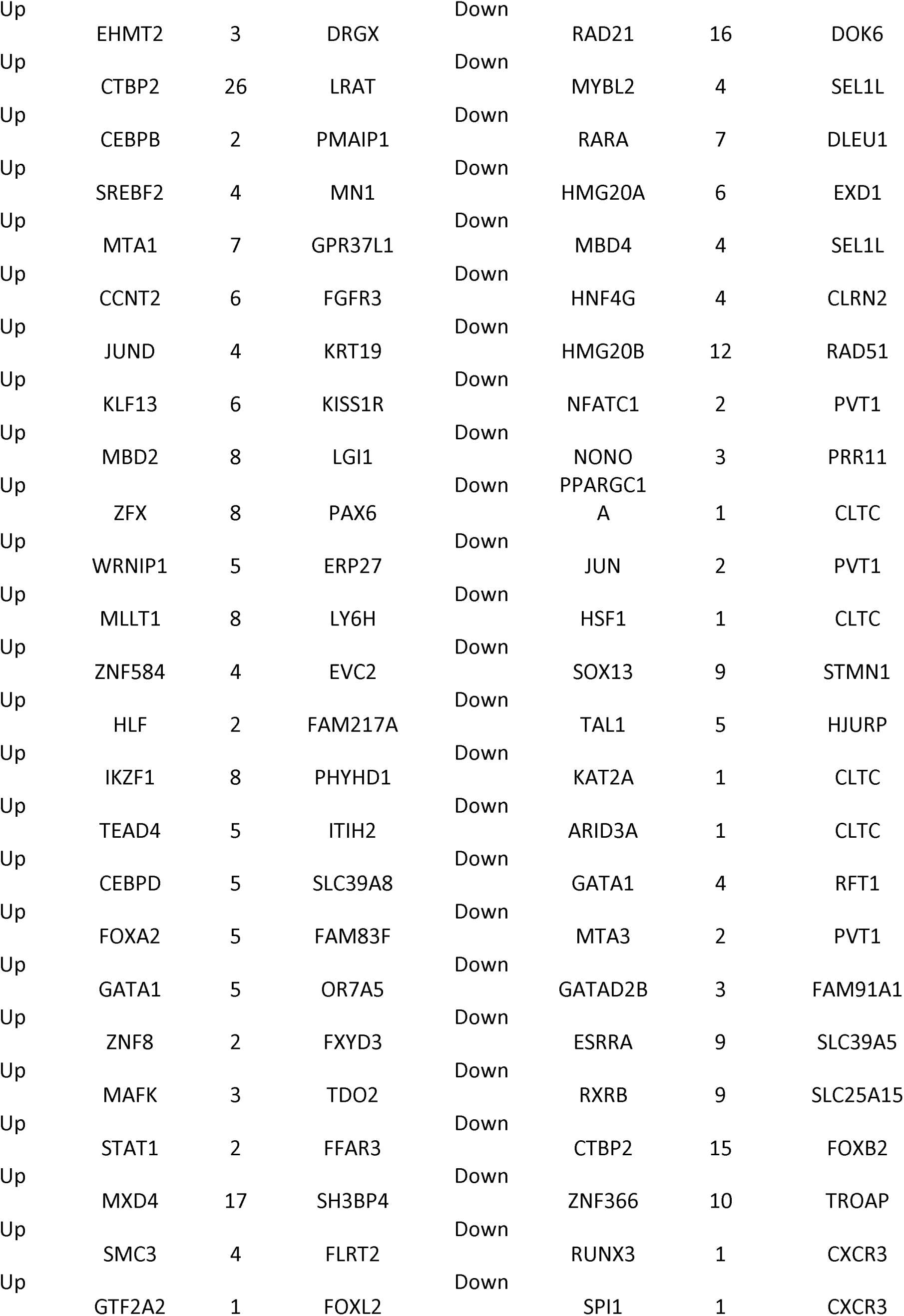
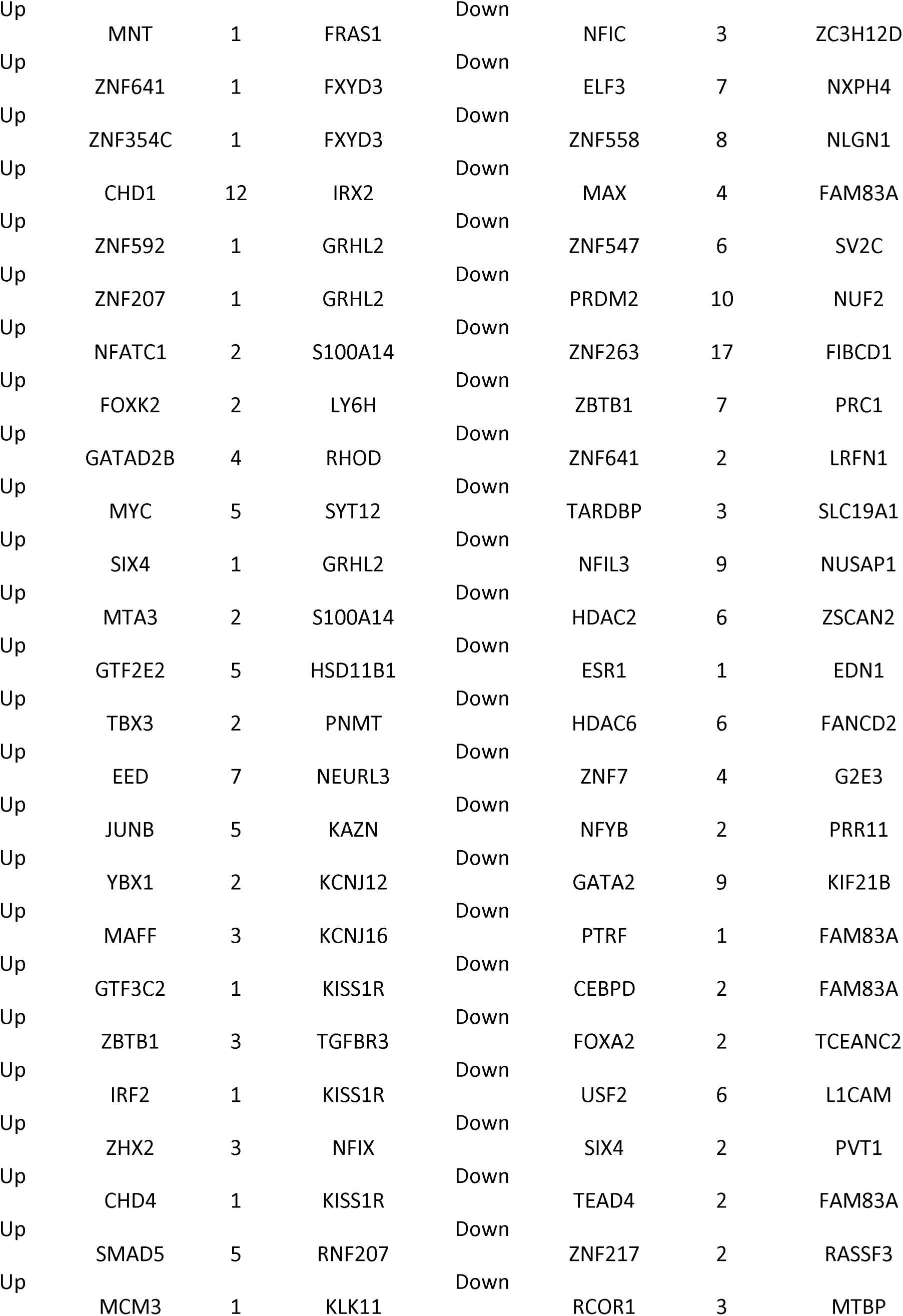
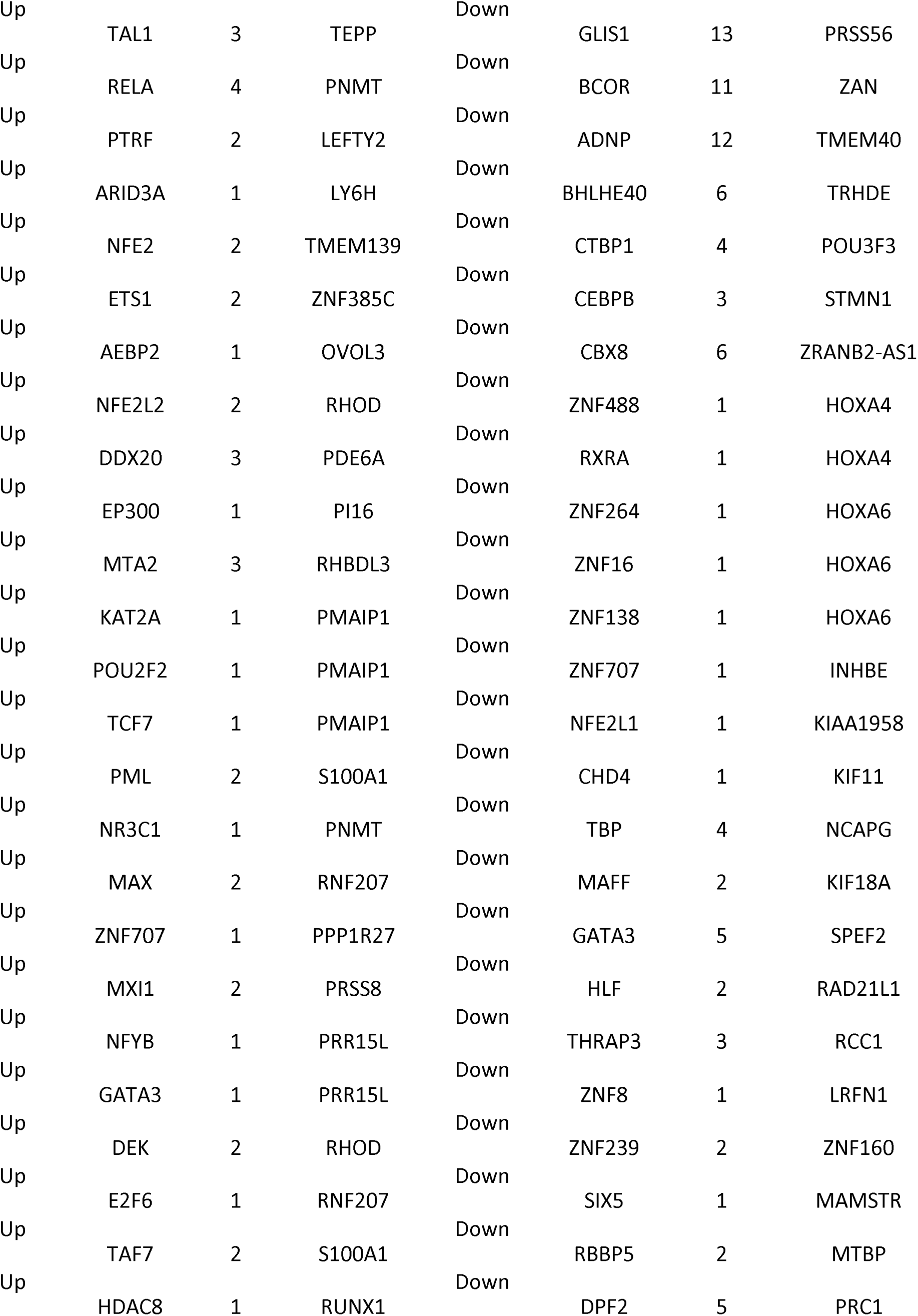
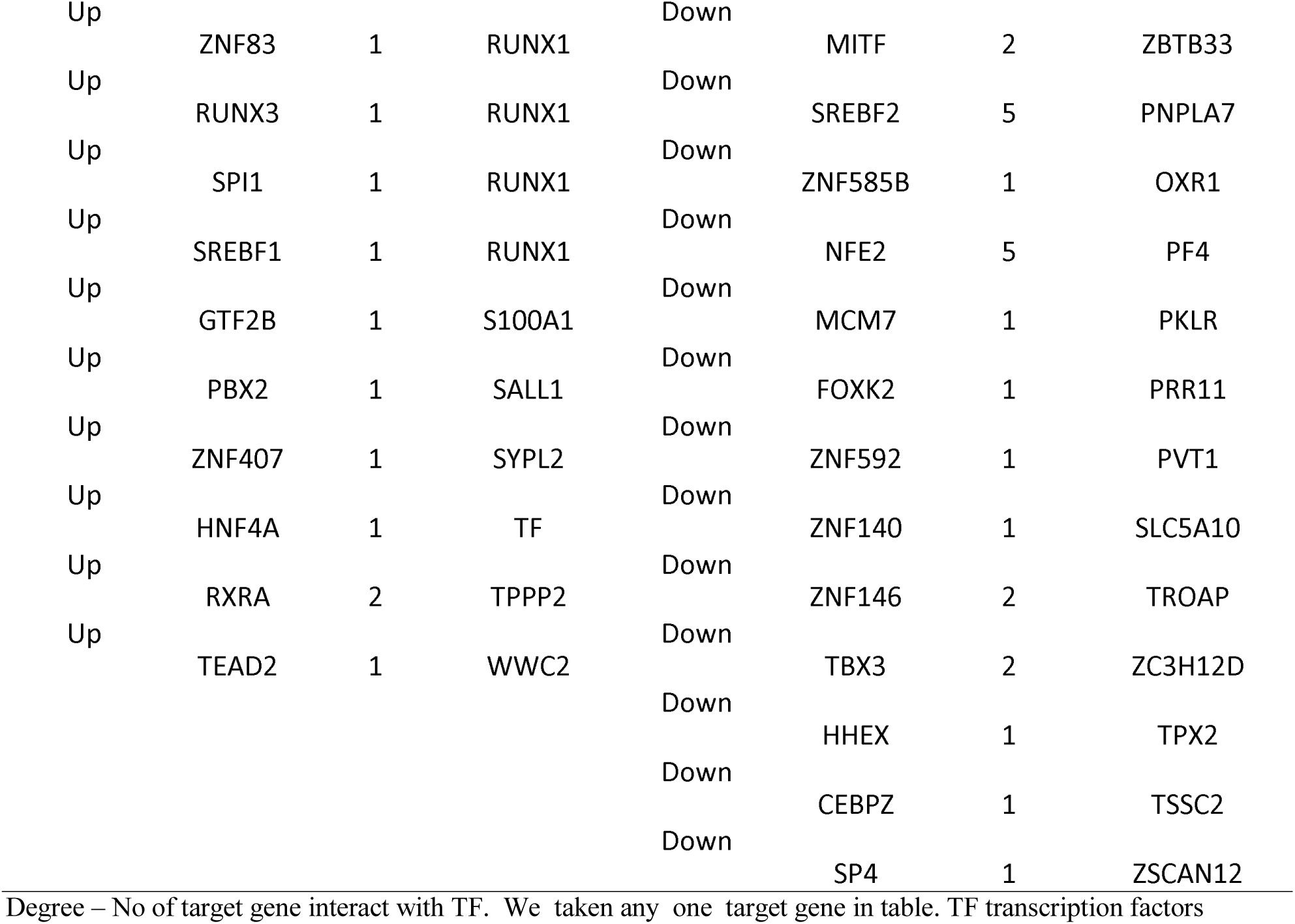
TF - target gene interaction table

| Regulation | TF | Degree | Gene | Regulation | TF | Degree | Gene |
| --- | --- | --- | --- | --- | --- | --- | --- |
| Up | EZH2 | 73 | IRX1 | Down | CREM | 10 | ACER2 |
| Up | WT1 | 12 | ADAMTS18 | Down | TRIM28 | 14 | ARHGAP29 |
| Up | ZNF76 | 22 | ADARB2 | Down | GATAD1 | 11 | FLJ13224 |
| Up | SUZ12 | 30 | CACNA2D3 | Down | SIN3A | 28 | ANTXR2 |
| Up | EBF1 | 14 | CSHL1 | Down | PHF8 | 31 | AKAP3 |
| Up | FOSL2 | 10 | AGR2 | Down | MXD4 | 20 | BIRC5 |
| Up | ZNF217 | 9 | AMZ1 | Down | SAP30 | 31 | HOXA4 |
| Up | NFIA | 11 | ALDOB | Down | STAT1 | 7 | CDCA5 |
| Up | ZNF644 | 17 | ATF7IP2 | Down | GLI4 | 8 | ASPDH |
| Up | GATA4 | 14 | CHN2 | Down | KDM5B | 31 | ATP2A3 |
| Up | NR2F2 | 6 | AOX1 | Down | KDM5A | 35 | CCNF |
| Up | CREB1 | 14 | CNKSR3 | Down | ZEB1 | 26 | C3orf52 |
| Up | ZBTB33 | 25 | APOD | Down | ATF1 | 23 | CKS2 |
| Up | TFE3 | 14 | CNKSR3 | Down | NR2C2 | 19 | C1orf189 |
| Up | MBD4 | 10 | DPPA4 | Down | TFDP1 | 26 | CADPS2 |
| Up | MYBL2 | 10 | HSD17B2 | Down | ZNF589 | 14 | DAZAP2 |
| Up | ZFP64 | 15 | C6orf132 | Down | POLR2H | 14 | CDC45 |
| Up | ATF1 | 18 | CCDC68 | Down | ETV1 | 10 | FAM83A |
| Up | GATAD2<br>A | 18 | CRYM | Down | SMARCA4 | 12 | CENPH |
| Up | RCOR2 | 13 | FGFR3 | Down | MAZ | 34 | BDNF |
| Up | BCL6 | 11 | KISS1R | Down | SIRT6 | 12 | HMMR |
| Up | FOXA3 | 15 | DMKN | Down | REST | 24 | CDCA5 |
| Up | YY1 | 9 | FABP7 | Down | PPARG | 17 | BIRC5 |
| Up | HMG20B | 16 | IL6R | Down | ZNF2 | 21 | BUB1 |
| Up | ZNF7 | 2 | PNMT | Down | BCL11A | 22 | CENPM |
| Up | NFIC | 4 | FMO5 | Down | LEF1 | 13 | CCNG2 |
| Up | HNH4G | 10 | PRR15L | Down | INSM2 | 18 | CKS2 |
| Up | MBD1 | 27 | DACT2 | Down | BCL3 | 1 | AMIGO2 |
| Up | THAP1 | 6 | FCN2 | Down | NCOR1 | 19 | CLTC |
| Up | RARA | 6 | DMKN | Down | FOSL2 | 6 | MYBPC1 |
| Up | ETV4 | 10 | ITIH2 | Down | ZNF335 | 16 | CYP27B1 |
| Up | L3MBTL2 | 19 | AMOT | Down | TAF7 | 21 | DAZAP2 |
| Up | CTBP1 | 2 | ASXL3 | Down | CREB1 | 15 | DHX15 |
| Up | NONO | 3 | GRHL2 | Down | GABPA | 16 | CLRN2 |
| Up | PPARG | 19 | CSH1 | Down | L3MBTL2 | 20 | C3orf80 |
| Up | MAZ | 28 | BMP7 | Down | ZNF175 | 10 | EXD1 |
| Up | CBFB | 7 | FFAR3 | Down | ZNF639 | 16 | CRNDE |
| Up | SMAD4 | 14 | CLIC3 | Down | KLF16 | 22 | FANCA |
| Up | RNF2 | 12 | CCNJL | Down | SMAD5 | 31 | CNIH2 |
| Up | ATF4 | 4 | FRAS1 | Down | EED | 9 | KIF18A |
| Up | HBP1 | 16 | LOC100288911 | Down | NRF1 | 25 | FANCA |
| Up | ATF3 | 12 | C6orf141 | Down | SP7 | 16 | B4GALT6 |
| Up | KLF11 | 14 | LINC00242 | Down | ZNF580 | 23 | HOXA6 |
| Up | ZNF580 | 19 | ARMC4 | Down | SMARCE1 | 11 | KIF20A |
| Up | GATAD1 | 15 | PHYHD1 | Down | POLR2A | 20 | C20orf203 |
| Up | CTCF | 22 | ATP4A | Down | DMAP1 | 14 | FAM20C |
| Up | ZBTB26 | 16 | FAM3B | Down | ZNF501 | 26 | ARHGAP20 |
| Up | ZNF589 | 11 | CA12 | Down | SP3 | 11 | G2E3 |
| Up | PKNOX1 | 11 | CSH2 | Down | SUZ12 | 23 | CRNDE |
| Up | ZEB1 | 11 | AQP4 | Down | BCL11B | 28 | DISC1 |
| Up | ZNF263 | 15 | CLIC3 | Down | GFI1B | 10 | FOXM1 |
| Up | KLF1 | 17 | C9orf152 | Down | ZNF324 | 17 | C1orf189 |
| Up | ZNF558 | 7 | ARMC3 | Down | ARID4B | 17 | MYO1D |
| Up | TBX21 | 5 | DMKN | Down | MXD3 | 21 | DOK6 |
| Up | ZNF324 | 13 | WNT5A | Down | SMAD4 | 16 | LOC339874 |
| Up | ZNF547 | 2 | NGFR | Down | IRF4 | 13 | NHLRC4 |
| Up | ZNF202 | 2 | ARMC4 | Down | PRDM10 | 9 | MAMSTR |
| Up | ZNF175 | 8 | ART3 | Down | EGR1 | 13 | NXPH4 |
| Up | ZNF239 | 4 | ASXL3 | Down | CREB3L1 | 25 | NUSAP1 |
| Up | ZNF121 | 11 | CELSR1 | Down | TCF7L2 | 1 | ASPDH |
| Up | TARDBP | 7 | MKX | Down | MTA1 | 15 | YTHDF1 |
| Up | SP3 | 19 | FAM196A | Down | SREBF1 | 3 | PARPBP |
| Up | PRDM2 | 6 | CPLX3 | Down | CDC5L | 7 | LRFN1 |
| Up | BCOR | 8 | KISS1R | Down | NR2F1 | 18 | ZSCAN12 |
| Up | POLR2H | 2 | GJB7 | Down | TRIM22 | 15 | VWA1 |
| Up | CEBPA | 5 | SLC6A14 | Down | IKZF1 | 13 | TROAP |
| Up | TRIM24 | 4 | LDLRAD2 | Down | ZNF202 | 6 | TRHDE |
| Up | NR2F1 | 13 | IL22RA1 | Down | HIC1 | 16 | TEX22 |
| Up | ZNF384 | 7 | ID4 | Down | ID3 | 18 | TCEANC2 |
| Up | NCOR1 | 6 | KAZN | Down | KLF6 | 12 | TTK |
| Up | NR4A1 | 11 | FOXL2 | Down | ZNF76 | 27 | ZRANB2-AS1 |
| Up | GMEB2 | 2 | PRSS8 | Down | TCF7 | 5 | SOX4 |
| Up | THRB | 11 | CTRB1 | Down | DRAP1 | 13 | SERINC2 |
| Up | TGIF2 | 19 | NFIX | Down | KLF9 | 43 | UNC13A |
| Up | NFYC | 14 | FMO5 | Down | HCFC1 | 15 | TRIM24 |
| Up | ZNF512 | 7 | CT45A1 | Down | CEBPA | 6 | RASSF10 |
| Up | GABPA | 9 | EVC2 | Down | MBD1 | 10 | PLK1 |
| Up | ARID4B | 13 | PPP1R27 | Down | MLX | 14 | ZSCAN2 |
| Up | KDM1A | 13 | PLIN4 | Down | NFYC | 11 | TPX2 |
| Up | DMAP1 | 20 | C1orf168 | Down | KDM1A | 9 | SPC24 |
| Up | KLF7 | 21 | ENTPD8 | Down | ELF1 | 24 | RGS5 |
| Up | GLIS2 | 27 | KCNJ12 | Down | MNT | 3 | PARPBP |
| Up | ZNF146 | 6 | OVOL3 | Down | PBX2 | 9 | TLE4 |
| Up | SP1 | 14 | NTF4 | Down | HBP1 | 15 | SLC39A5 |
| Up | ZNF488 | 2 | FXYD3 | Down | KLF8 | 20 | RCC1 |
| Up | BCL11A | 12 | KRT19 | Down | KLF11 | 12 | ZC3H12D |
| Up | ZNF341 | 19 | COL13A1 | Down | CHD1 | 20 | RASSF3 |
| Up | BCL11B | 16 | EYA4 | Down | FOXJ2 | 13 | TCEANC2 |
| Up | ZBTB7A | 19 | SALL1 | Down | TEAD3 | 10 | TRIM24 |
| Up | HIC1 | 23 | FGFR2 | Down | HMG3 | 16 | KIF11 |
| Up | ZNF501 | 17 | FAM198A | Down | ZNF584 | 9 | PF4 |
| Up | SOX13 | 19 | MGST1 | Down | NCOA1 | 10 | DAZAP2 |
| Up | ESRRA | 8 | ESRRA | Down | ZKSCAN1 | 8 | PVT1 |
| Up | INSM2 | 11 | PMAIP1 | Down | HDAC1 | 12 | HAUS6 |
| Up | ZNF2 | 17 | GAS1 | Down | ZNF423 | 18 | KLHL13 |
| Up | ZNF610 | 11 | ROR2 | Down | ZFX | 23 | HMMR |
| Up | ETV6 | 1 | BMP7 | Down | EZH2 | 34 | POU3F4 |
| Up | ZNF394 | 15 | CACNA2D3 | Down | EP300 | 4 | PBK |
| Up | ZNF18 | 12 | RNF207 | Down | GTF2E2 | 28 | KIAA1958 |
| Up | ZNF24 | 15 | DNAI2 | Down | NFRKB | 12 | CCNB2 |
| Up | KLF8 | 11 | NKX2-2 | Down | HDGF | 17 | POLQ |
| Up | ELF3 | 14 | BTBD17 | Down | JUNB | 6 | LINC00309 |
| Up | DIDO1 | 2 | LEFTY2 | Down | WRNIP1 | 24 | CALHM1 |
| Up | RERE | 11 | SOX9 | Down | MLLT1 | 11 | PTP4A1 |
| Up | ZNF197 | 11 | WWC2 | Down | ZFP37 | 23 | DISC1 |
| Up | MLX | 12 | FLJ42393 | Down | TBX21 | 8 | SPC24 |
| Up | FOSL1 | 11 | FMO5 | Down | SMC3 | 6 | BDNF |
| Up | ESR1 | 3 | LGI1 | Down | ZBTB17 | 8 | POU3F3 |
| Up | ZFP2 | 12 | STRA6 | Down | EHMT2 | 1 | BDNF |
| Up | POLR2A | 7 | LDLRAD2 | Down | MYNN | 21 | C20orf203 |
| Up | NFIL3 | 11 | NEURL3 | Down | KLF13 | 11 | INHBE |
| Up | KLF16 | 21 | GH2 | Down | FOSL1 | 18 | PTP4A1 |
| Up | KLF4 | 16 | GADD45G | Down | CTCF | 12 | PKLR |
| Up | REST | 13 | RHBDL3 | Down | ZNF644 | 17 | MAMSTR |
| Up | RAD21 | 17 | RBP4 | Down | SSRP1 | 9 | TRIM24 |
| Up | MXD3 | 14 | S100A1 | Down | RERE | 18 | PTTG2 |
| Up | SCRT1 | 7 | GPR37L1 | Down | CEBPG | 7 | MTBP |
| Up | ZNF335 | 10 | RPRML | Down | ZHX2 | 11 | LOC339874 |
| Up | ELF1 | 11 | DLEC1 | Down | ETS1 | 18 | CNIH2 |
| Up | GATA2 | 7 | CA8 | Down | SCRT2 | 6 | SLC52A3 |
| Up | KLF9 | 26 | CCKBR | Down | GMEB2 | 15 | ZSCAN12 |
| Up | RXRB | 11 | TMEM139 | Down | MYC | 5 | FANCA |
| Up | DPF2 | 6 | GRHL2 | Down | SCRT1 | 3 | STMN1 |
| Up | NR2F6 | 15 | PTER | Down | THRB | 12 | SLC25A15 |
| Up | ZNF71 | 10 | EFEMP1 | Down | ZNF197 | 18 | RASSF10 |
| Up | ZNF101 | 7 | RNF207 | Down | ZBTB7A | 28 | QSOX1 |
| Up | JUN | 2 | C6orf141 | Down | MXI1 | 6 | HJURP |
| Up | ZNF623 | 5 | HIF3A | Down | ZNF394 | 24 | ZNF160 |
| Up | ZNF140 | 1 | C6orf141 | Down | ZBTB40 | 7 | G2E3 |
| Up | ZNF292 | 1 | C6orf141 | Down | E2F4 | 14 | FANCD2 |
| Up | ZNF264 | 2 | OVOL3 | Down | RNF2 | 9 | PNPLA7 |
| Up | CUX1 | 9 | SYT12 | Down | ZNF407 | 8 | CCNB2 |
| Up | GFI1B | 6 | TGFBR3 | Down | ZBTB11 | 21 | RAD51 |
| Up | ZNF382 | 9 | KCNG1 | Down | MAFG | 3 | CLTC |
| Up | PRDM12 | 1 | C6orf141 | Down | POU5F1 | 2 | BUB1 |
| Up | FOS | 3 | IL6R | Down | RCOR2 | 10 | CDKN3 |
| Up | PRDM10 | 14 | STRA6 | Down | NR4A1 | 12 | C20orf203 |
| Up | TEAD3 | 14 | TF | Down | TFAP4 | 16 | RFT1 |
| Up | IRF1 | 10 | TEPP | Down | ZNF101 | 11 | CADPS2 |
| Up | SIRT6 | 5 | CA8 | Down | ZNF341 | 11 | NXPH4 |
| Up | MIXL1 | 3 | MYBPHL | Down | MEF2D | 12 | RCC1 |
| Up | ZBTB11 | 9 | SLC39A8 | Down | YY1 | 10 | KIF18A |
| Up | CREB3L1 | 9 | RASL11B | Down | EBF1 | 8 | KIR3DX1 |
| Up | TFDP1 | 29 | VSNL1 | Down | TSHZ1 | 11 | PKLR |
| Up | HMBBOX1 | 5 | TMEM154 | Down | PKNOX1 | 4 | LYNX1 |
| Up | BHLHE40 | 7 | PPP1R27 | Down | WT1 | 15 | RAD21L1 |
| Up | BACH1 | 3 | PNMT | Down | FOXA3 | 14 | FOXMI |
| Up | FOXJ2 | 7 | S100B | Down | PRDM1 | 10 | PRDM1 |
| Up | SMARCA4 | 5 | FAM83F | Down | ZNF610 | 16 | SLC19A1 |
| Up | SIN3A | 17 | PAX6 | Down | ZNF18 | 17 | DISC1 |
| Up | ZBTB40 | 6 | HPGD | Down | KLF7 | 20 | NHLRC4 |
| Up | ID3 | 13 | SRL | Down | ZBTB33 | 15 | HOXA6 |
| Up | TRIM28 | 4 | FBN2 | Down | GLIS2 | 19 | ZBTB33 |
| Up | HDAC1 | 13 | GADD45G | Down | KLF4 | 8 | PRR11 |
| Up | SMARCE1 | 8 | CYBRD1 | Down | SUPT5H | 7 | EVX2 |
| Up | GLI4 | 4 | RUNX1 | Down | ARNT | 6 | CDC45 |
| Up | SCRT2 | 3 | PDE6A | Down | EGR2 | 13 | FLJ13224 |
| Up | GLIS1 | 15 | STRA6 | Down | ZNF382 | 6 | HOXA4 |
| Up | EGR2 | 15 | RPRM | Down | RFX1 | 7 | OMP |
| Up | ATF2 | 4 | CH25H | Down | ELK1 | 23 | ZMAT3 |
| Up | HHEX | 5 | SOX9 | Down | ZNF71 | 17 | NUF2 |
| Up | STAT3 | 5 | CLEC9A | Down | DDX20 | 16 | TSSC2 |
| Up | CEBPG | 14 | TSPAN8 | Down | SP1 | 25 | LOC100268168 |
| Up | HCFC1 | 4 | CLIC3 | Down | TEAD2 | 1 | CDC45 |
| Up | TFAP4 | 14 | TRIM71 | Down | SMARCA5 | 13 | RAD21L1 |
| Up | NR2C2 | 10 | MFAP4 | Down | ZNF83 | 9 | MAMSTR |
| Up | HMGN3 | 8 | ENTPD8 | Down | ZNF24 | 22 | MTBP |
| Up | FOXM1 | 7 | NTF4 | Down | DNMT1 | 5 | KIF20A |
| Up | EGR1 | 7 | RHOD | Down | GTF2A2 | 7 | FAM83A |
| Up | IRF4 | 7 | KISS1R | Down | E2F5 | 13 | DHX15 |
| Up | DRAP1 | 14 | NEURL3 | Down | IRF2 | 7 | CNIH2 |
| Up | SP7 | 10 | PPP1R27 | Down | HMBOX1 | 5 | CLTC |
| Up | NCOA1 | 3 | TMPRSS3 | Down | CREB3 | 4 | MYO1D |
| Up | CREM | 8 | S100A14 | Down | ZNF623 | 9 | TCEANC2 |
| Up | HMG20A | 12 | ZNF385C | Down | JUND | 8 | MYBPC1 |
| Up | LEF1 | 3 | TF | Down | MBD2 | 9 | PRR11 |
| Up | DNMT1 | 1 | CNKS3 | Down | ATF3 | 12 | CRNDE |
| Up | ZNF423 | 18 | PPP1R1B | Down | MIXL1 | 6 | SLC35G5 |
| Up | TEAD1 | 9 | SPTLC3 | Down | ZNF512 | 7 | HTR3D |
| Up | ADNP | 8 | RNF207 | Down | ATF2 | 6 | NLGN1 |
| Up | HES1 | 5 | GADD45G | Down | ZFP2 | 23 | GPR161 |
| Up | ARNT | 2 | TEPP | Down | MAFK | 5 | STMN1 |
| Up | ZFP37 | 13 | RASGRP1 | Down | KLF1 | 21 | ZBTB33 |
| Up | ZBTB17 | 4 | KRT19 | Down | HDAC8 | 1 | CDC45 |
| Up | MYNN | 10 | KISS1R | Down | ETS2 | 2 | TRIM24 |
| Up | ARID1B | 7 | PON3 | Down | RELA | 3 | PKLR |
| Up | RFXANK | 8 | PRSS8 | Down | NR2F6 | 22 | QSOX1 |
| Up | ZNF143 | 4 | COL13A1 | Down | HES1 | 2 | KIF15 |
| Up | SP2 | 17 | EFCAB1 | Down | ZHX1 | 2 | TSSC2 |
| Up | CDC5L | 5 | PTER | Down | TSC22D4 | 1 | CDC45 |
| Up | ZNF16 | 4 | RUNX1 | Down | BACH1 | 6 | ZSCAN12 |
| Up | PRDM1 | 4 | TGFBR3 | Down | DIDO1 | 4 | CYP27B1 |
| Up | TRIM22 | 6 | SULT1C4 | Down | CUX1 | 7 | PVT1 |
| Up | GTF2F1 | 11 | SYT12 | Down | ETV6 | 3 | PRR11 |
| Up | TSHZ1 | 5 | KRT19P2 | Down | ARID1B | 12 | RACGAP1 |
| Up | RFX1 | 5 | PGR | Down | ZNF207 | 15 | RAD51D |
| Up | PTTG1 | 1 | CRISP2 | Down | IRF1 | 24 | CENPH |
| Up | MEF2D | 8 | PDE6A | Down | GATA4 | 6 | TPX2 |
| Up | SSRP1 | 15 | TRIM71 | Down | NFIA | 9 | VWA1 |
| Up | NRF1 | 14 | SALL1 | Down | TGIF2 | 13 | PLK1 |
| Up | KDM5A | 2 | CRYM | Down | CBFB | 9 | PSTPIP2 |
| Up | RCOR1 | 6 | GRHL2 | Down | TFE3 | 14 | DISC1 |
| Up | HDAC2 | 5 | ESPNL | Down | ZFP64 | 9 | G2E3 |
| Up | HDAC6 | 5 | SSTR2 | Down | SIN3B | 9 | PTP4A1 |
| Up | SMARCA5 | 11 | GH2 | Down | GTF2F1 | 17 | CNIH2 |
| Up | E2F5 | 11 | CSH2 | Down | ZBTB26 | 11 | SOX4 |
| Up | HDGF | 6 | CSHL1 | Down | CCNT2 | 8 | PNPLA7 |
| Up | ETV1 | 5 | DLEC1 | Down | TEAD1 | 8 | NCAPG |
| Up | PYGO2 | 1 | CYBRD1 | Down | GTF2B | 8 | KIF11 |
| Up | ELK1 | 6 | FOXL2 | Down | RFXANK | 9 | HAUS6 |
| Up | KDM5B | 11 | WWC2 | Down | SOX5 | 10 | CDKN3 |
| Up | NFRKB | 7 | LEFTY2 | Down | NR2F2 | 4 | SPC24 |
| Up | MITF | 2 | RASGRP1 | Down | GATAD2A | 11 | LYNX1 |
| Up | PHF8 | 11 | ID4 | Down | THAP1 | 4 | CLTC |
| Up | ZNF639 | 6 | MYBPHL | Down | RFX3 | 6 | CENPA |
| Up | SAP30 | 11 | LOC100288911 | Down | ETV4 | 15 | HMMR |
| Up | CREB3 | 6 | SRL | Down | ZNF384 | 13 | SLITRK6 |
| Up | KLF6 | 7 | VANGL2 | Down | SP2 | 20 | UNC13A |
| Up | ZKSCAN1 | 5 | DLK1 | Down | ZNF143 | 11 | RFC3 |
| Up | CBX8 | 6 | SOX2 | Down | MTA2 | 5 | FLJ13224 |
| Up | ZNF138 | 1 | DMKN | Down | ZNF121 | 6 | TMEM40 |
| Up | ZNF366 | 7 | WNT5A | Down | BCL6 | 12 | YTHDF1 |
| Up | SUPT5H | 5 | S100B | Down | ATF4 | 7 | RAD21L1 |
| Up | USF2 | 5 | SYPL2 | Down | USF1 | 3 | PNPLA7 |
| Up | POU5F1 | 5 | DPPA4 | Down | PML | 4 | RCC1 |
| Up | EHMT2 | 3 | DRGX | Down | RAD21 | 16 | DOK6 |
| Up | CTBP2 | 26 | LRAT | Down | MYBL2 | 4 | SEL1L |
| Up | CEBPB | 2 | PMAIP1 | Down | RARA | 7 | DLEU1 |
| Up | SREBF2 | 4 | MN1 | Down | HMG20A | 6 | EXD1 |
| Up | MTA1 | 7 | GPR37L1 | Down | MBD4 | 4 | SEL1L |
| Up | CCNT2 | 6 | FGFR3 | Down | HNF4G | 4 | CLRN2 |
| Up | JUND | 4 | KRT19 | Down | HMG20B | 12 | RAD51 |
| Up | KLF13 | 6 | KISS1R | Down | NFATC1 | 2 | PVT1 |
| Up | MBD2 | 8 | LGI1 | Down | NONO | 3 | PRR11 |
| Up | ZFX | 8 | PAX6 | Down | PPARGC1<br>A | 1 | CLTC |
| Up | WRNIP1 | 5 | ERP27 | Down | JUN | 2 | PVT1 |
| Up | MLLT1 | 8 | LY6H | Down | HSF1 | 1 | CLTC |
| Up | ZNF584 | 4 | EVC2 | Down | SOX13 | 9 | STMN1 |
| Up | HLF | 2 | FAM217A | Down | TAL1 | 5 | HJURP |
| Up | IKZF1 | 8 | PHYHD1 | Down | KAT2A | 1 | CLTC |
| Up | TEAD4 | 5 | ITIH2 | Down | ARID3A | 1 | CLTC |
| Up | CEBPD | 5 | SLC39A8 | Down | GATA1 | 4 | RFT1 |
| Up | FOXA2 | 5 | FAM83F | Down | MTA3 | 2 | PVT1 |
| Up | GATA1 | 5 | OR7A5 | Down | GATAD2B | 3 | FAM91A1 |
| Up | ZNF8 | 2 | FXYD3 | Down | ESRRA | 9 | SLC39A5 |
| Up | MAFK | 3 | TDO2 | Down | RXRB | 9 | SLC25A15 |
| Up | STAT1 | 2 | FFAR3 | Down | CTBP2 | 15 | FOXB2 |
| Up | MXD4 | 17 | SH3BP4 | Down | ZNF366 | 10 | TROAP |
| Up | SMC3 | 4 | FLRT2 | Down | RUNX3 | 1 | CXCR3 |
| Up | GTF2A2 | 1 | FOXL2 | Down | SPI1 | 1 | CXCR3 |
| Up | MNT | 1 | FRAS1 | Down | NFIC | 3 | ZC3H12D |
| Up | ZNF641 | 1 | FXYD3 | Down | ELF3 | 7 | NXPH4 |
| Up | ZNF354C | 1 | FXYD3 | Down | ZNF558 | 8 | NLGN1 |
| Up | CHD1 | 12 | IRX2 | Down | MAX | 4 | FAM83A |
| Up | ZNF592 | 1 | GRHL2 | Down | ZNF547 | 6 | SV2C |
| Up | ZNF207 | 1 | GRHL2 | Down | PRDM2 | 10 | NUF2 |
| Up | NFATC1 | 2 | S100A14 | Down | ZNF263 | 17 | FIBCD1 |
| Up | FO XK2 | 2 | LY6H | Down | ZBTB1 | 7 | PRC1 |
| Up | GATAD2B | 4 | RHOD | Down | ZNF641 | 2 | LRFN1 |
| Up | MYC | 5 | SYT12 | Down | TARDBP | 3 | SLC19A1 |
| Up | SIX4 | 1 | GRHL2 | Down | NFIL3 | 9 | NUSAP1 |
| Up | MTA3 | 2 | S100A14 | Down | HDAC2 | 6 | ZSCAN2 |
| Up | GTF2E2 | 5 | HSD11B1 | Down | ESR1 | 1 | EDN1 |
| Up | TBX3 | 2 | PNMT | Down | HDAC6 | 6 | FANCD2 |
| Up | EED | 7 | NEURL3 | Down | ZNF7 | 4 | G2E3 |
| Up | JUNB | 5 | KAZN | Down | NFYB | 2 | PRR11 |
| Up | YBX1 | 2 | KCNJ12 | Down | GATA2 | 9 | KIF21B |
| Up | MAFF | 3 | KCNJ16 | Down | PTRF | 1 | FAM83A |
| Up | GTF3C2 | 1 | KISS1R | Down | CEBPD | 2 | FAM83A |
| Up | ZBTB1 | 3 | TGFBR3 | Down | FOXA2 | 2 | TCEANC2 |
| Up | IRF2 | 1 | KISS1R | Down | USF2 | 6 | L1CAM |
| Up | ZHX2 | 3 | NFIX | Down | SIX4 | 2 | PVT1 |
| Up | CHD4 | 1 | KISS1R | Down | TEAD4 | 2 | FAM83A |
| Up | SMAD5 | 5 | RNF207 | Down | ZNF217 | 2 | RASSF3 |
| Up | MCM3 | 1 | KLK11 | Down | RCOR1 | 3 | MTBP |
| Up | TAL1 | 3 | TEPP | Down | GLIS1 | 13 | PRSS56 |
| Up | RELA | 4 | PNMT | Down | BCOR | 11 | ZAN |
| Up | PTRF | 2 | LEFTY2 | Down | ADNP | 12 | TMEM40 |
| Up | ARID3A | 1 | LY6H | Down | BHLHE40 | 6 | TRHDE |
| Up | NFE2 | 2 | TMEM139 | Down | CTBP1 | 4 | POU3F3 |
| Up | ETS1 | 2 | ZNF385C | Down | CEBPB | 3 | STMN1 |
| Up | AEBP2 | 1 | OVOL3 | Down | CBX8 | 6 | ZRANB2-AS1 |
| Up | NFE2L2 | 2 | RHOD | Down | ZNF488 | 1 | HOXA4 |
| Up | DDX20 | 3 | PDE6A | Down | RXRA | 1 | HOXA4 |
| Up | EP300 | 1 | PI16 | Down | ZNF264 | 1 | HOXA6 |
| Up | MTA2 | 3 | RHBDL3 | Down | ZNF16 | 1 | HOXA6 |
| Up | KAT2A | 1 | PMAIP1 | Down | ZNF138 | 1 | HOXA6 |
| Up | POU2F2 | 1 | PMAIP1 | Down | ZNF707 | 1 | INHBE |
| Up | TCF7 | 1 | PMAIP1 | Down | NFE2L1 | 1 | KIAA1958 |
| Up | PML | 2 | S100A1 | Down | CHD4 | 1 | KIF11 |
| Up | NR3C1 | 1 | PNMT | Down | TBP | 4 | NCAPG |
| Up | MAX | 2 | RNF207 | Down | MAFF | 2 | KIF18A |
| Up | ZNF707 | 1 | PPP1R27 | Down | GATA3 | 5 | SPEF2 |
| Up | MXI1 | 2 | PRSS8 | Down | HLF | 2 | RAD21L1 |
| Up | NFYB | 1 | PRR15L | Down | THRAP3 | 3 | RCC1 |
| Up | GATA3 | 1 | PRR15L | Down | ZNF8 | 1 | LRFN1 |
| Up | DEK | 2 | RHOD | Down | ZNF239 | 2 | ZNF160 |
| Up | E2F6 | 1 | RNF207 | Down | SIX5 | 1 | MAMSTR |
| Up | TAF7 | 2 | S100A1 | Down | RBBP5 | 2 | MTBP |
| Up | HDAC8 | 1 | RUNX1 | Down | DPF2 | 5 | PRC1 |
| Up | ZNF83 | 1 | RUNX1 | Down | MITF | 2 | ZBTB33 |
| Up | RUNX3 | 1 | RUNX1 | Down | SREBF2 | 5 | PNPLA7 |
| Up | SPI1 | 1 | RUNX1 | Down | ZNF585B | 1 | OXR1 |
| Up | SREBF1 | 1 | RUNX1 | Down | NFE2 | 5 | PF4 |
| Up | GTF2B | 1 | S100A1 | Down | MCM7 | 1 | PKLR |
| Up | PBX2 | 1 | SALL1 | Down | FOXK2 | 1 | PRR11 |
| Up | ZNF407 | 1 | SYPL2 | Down | ZNF592 | 1 | PVT1 |
| Up | HNF4A | 1 | TF | Down | ZNF140 | 1 | SLC5A10 |
| Up | RXRA | 2 | TPPP2 | Down | ZNF146 | 2 | TROAP |
| Up | TEAD2 | 1 | WWC2 | Down | TBX3 | 2 | ZC3H12D |
|  |  |  |  | Down | HHEX | 1 | TPX2 |
|  |  |  |  | Down | CEBPZ | 1 | TSSC2 |
|  |  |  |  | Down | SP4 | 1 | ZSCAN12 |

Degree – No of target gene interact with TF. We taken any one target gene in table. TF transcription factors

## Discussion

A molecular understanding of pituitary prolactinoma is particularly essential to eventually improve effective approaches for its control, treatment, and prevention. In the present study, important candidate genes of pituitary prolactinoma were identified by bioinformatics analysis. We extracted the gene expression data from GSE119063 and obtained 461 up regulated and 528 down regulated DEGs in pituitary prolactinoma. SALL1 action as a tumor suppressor and epigenetic inactivation of this gene is responsible for development of cancer [18], but association of this gene still not reported in pituitary prolactinoma. Wang et al [19] showed that expression of homeodomain transcription factor NKX2-2 is linked with neuro endocrine tumor. High expression of pleiotropic signaling molecule BMP7 controls proliferation, migration, and invasion of cancer cells [20], but this gene may be linked with pathogenesis of pituitary prolactinoma. Cipriano et al [21] identified that oncogene FAM83F is associated in stimulating CRAF/MAPK signaling pathway and driving epithelial cell transformation in cancer, but involvement of this gene may responsible for advancement of pituitary prolactinoma. Oncogene GRHL2 is a transcriptional controller of proliferation and differentiation in epithelial cells, both during progression and tumor development, but this gene still not reported in pituitary prolactinoma [22]. TRIM24 play key role in proliferation and invasion of cancer cell [23], but high expression of this gene may be responsible for development of pituitary prolactinoma. Kaistha et al [24] describe that high expression of dual specificity kinase TTK shows proliferative potential of cancer, but this gene may be linked with pathogenesis of pituitary prolactinoma. Over expression of TOP2A is responsible for improvement of cancer [25], but this gene may associate with pituitary prolactinoma. Potential role KIF18A is associated with abnormal cell division, checkpoint activation and tumor formation [26], but this gene may be responsible for advancement pituitary prolactinoma. Over expression of microtubule associated protein TPX2 stimulates the cell cycle kinase aurora A and strongly correlates with chromosomal instability in cancer [27], but this gene may be associated with pathogenesis of pituitary prolactinoma. Genes such as FGFR2 [28], SOX2 [29], POMC [30], FSHB [31], EFEMP1 [32], SFRP2 [33], CSH2 [34], IHH [35], GH1 [36], PTTG2 [37], CCNB2 [38], RACGAP1 [39], CXCR2 [40], CXCR3 [41], FOXL2 [42], RUNX2 [43], TF [44], CCK [45], DPPA4 [46], RUNX1 [43] and BDNF [47] are associated with pituitary prolactinoma.

In present study retinoate biosynthesis II is the most significant BIOCYC pathway for up-regulated genes. XDH and RBP4 are novel biomarkers for pathogenesis of pituitary prolactinoma. Signaling pathways regulating pluripotency of stem cells is the most significant KEGG pathway for up-regulated genes. Alteration in FGFR3 is diagnosed with bladder cancer [48], but this gene may be important for development of pituitary prolactinoma. WNT5A improves cancer cell invasion and proliferation through receptor-mediated endocytosis dependent and independent mechanisms [49], but this gene may be diagnosed with pituitary prolactinoma. ID4 is potential tumor suppressor and epigenetic inactivation of this gene linked with prostate cancer [50], but this gene may be liable for advancement of pituitary prolactinoma. High expression of INHBA improves cell proliferation and is associated with progression of lung adenocarcinoma [51], but this gene may be involve in pathogenesis of pituitary prolactinoma. PAX6 prevents cancer cell proliferation and invasion and epigenetic silencing of this is diagnosed with breast cancer [52], but this gene may be culpable for progression of pituitary prolactinoma. LEFTY2 and SMAD9 are novel biomarkers for pathogenesis of pituitary prolactinoma. ALK2 signaling events are the most significant PID pathway for up regulated genes. Peptide hormone biosynthesis is the most significant REACTOME pathway for up-regulated genes. CGB3 and TSHB are novel biomarkers for pathogenesis of pituitary prolactinoma. Tyrosine metabolism was the most significant GenMAPP pathway for up-regulated genes. Duan et al [53] showed that ALDH1A3 belongs to the member of aldehyde dehydrogenase (ALDH) superfamily and play a regulatory role in the initiation and development of cancer via the clearance of aldehydes and the synthesizes of retinoic acid, but this gene may be linked with pathogenesis of pituitary prolactinoma. PNMT and AOX1 are novel biomarkers for pathogenesis of pituitary prolactinoma. Ensemble of genes encoding extracellular matrix and extracellular matrix-associated proteins is the most significant MSigDB C2 BIOCARTA pathway for up-regulated genes. S100A14 is belongs to a member of the S100 protein family and actions as a modulator of HER2 signaling pathway in breast cancer [54], but this gene may be responsible for development of pituitary prolactinoma. DeRycke et al [55] found that high expression of calcium binding protein S100A1 is identified with ovarian cancer, but this gene may be important for improvement of pituitary prolactinoma. Harpio and Einarsson [56] identified that high level of S100B is answerable for development of melanoma, but this gene may be identified with pituitary prolactinoma. Methylation and silencing of large modular extracellular matrix glycoprotein FBN2 play an essential role in carcinogenesis, invasion and metastasis of non-small cell lung cancer [57], but this gene may be involved in pathogenesis of pituitary prolactinoma. Yoshimura et al [58] reported that FGF1 is an inducer of angiogenesis in breast cancer, but this gene may play a role in the pathogenesis of pituitary prolactinoma. SLIT1 is tumor suppressor gene and epigenetic suppression of this gene is associated with cancer [59], but this silenced gene may be culpable for development pituitary prolactinoma. ADAMTS18 and CHRDL1 are putative tumor suppressive genes and epigenetic silencing of these genes diagnosed with gastric, colorectal and pancreatic cancers [60–61], but inactivation of these genes may be link with pituitary prolactinoma. VIT, MFAP4, FCN2, SBSPON, SPON1, ITIH6, COL4A6, COL8A2, FRAS1, COL13A1, ITLN1, ADAM32, CSH1, CSHL1, SLPI, IFNE, MEGF11, LAMA1, ITIH2, GH2, COL21A1, CLEC9A, NTF4, OGN, RSPO3, MEGF6, CBLN2 and LGI1 are novel biomarkers for pathogenesis of pituitary prolactinoma. Adenine and hypoxanthine salvage pathway is the most significant PantherDB pathway for up-regulated genes. Melanocortin system is the most significant Pathway Ontology for up-regulated genes. PCSK2 is novel biomarkers for pathogenesis of pituitary prolactinoma. Tryptophan metabolism is the most significant PantherDB pathway for up-regulated genes. TDO2 and TPH1 are novel biomarkers for pathogenesis of pituitary prolactinoma. Vitamin D3 biosynthesis is the most significant BIOCYC pathway for down-regulated genes. Polymorphisms of CYP27B1 is modulate vitamin D metabolism in colon cancer [62], but this polymorphic gene may be responsible for improvement of pituitary prolactinoma. Cell cycle is the most significant KEGG pathway for down-regulated genes. High expression of PLK1 is linked with cancer cell proliferation [63], but this may be gene is liable for development pituitary prolactinoma. High expression of mitotic spindle checkpoint gene BUB1 is linked with gastric cancer cell proliferation [64], but this gene may be important in pathogenesis of pituitary prolactinoma. Epigenetic inactivation of CCNA1 is diagnosed with cervical cancer [65], but silencing of this gene may be responsible for advancement of pituitary prolactinoma. CDC45 is novel biomarker for pathogenesis of pituitary prolactinoma. Aurora B signaling is the most significant PID pathway for down-regulated genes. KIF23 plays as both a controller of cytokinesis and a motor enzyme of microtubule and this gene is over expressed in lung cancer [66], but this gene may be play crucial role in pituitary prolactinoma. BIRC5 play key role in cell division and proliferation and by inhibiting apoptosis in hepatocellular carcinoma [67], but this gene may be link with pituitary prolactinoma. NCAPG improves hepatocellular cancer cell proliferation and migration [68], but this gene may be diagnosed with pituitary prolactinoma. Nie et al [69] suggest that STMN1 play essential role in the control of cellular division and proliferation in non-small cell lung cancer, but this gene may be involved in pathogenesis of pituitary prolactinoma. High expression of KIF20A is associated with pancreatic cancer [70], but this gene may be play key role in pituitary prolactinoma. CENPA and KLHL13 are novel biomarkers for pathogenesis of pituitary prolactinoma. Mitotic prometaphase is the most significant REACTOME pathway for down-regulated genes. High expression of SPC24 and SPC25 are associated with genomic instability and disrupted regulation of cell cycle in lung cancer, but this gene may be responsible for progression of pituitary prolactinoma [71]. Over expression cell cycle molecular marker CENPH pay key role in gastric carcinoma cell proliferation and gastric carcinoma cell survival as well as procancer role [72], but this gene may be involved in pituitary prolactinoma. Over expression of NUF2 activates tumor growth and inhibits cell apoptosis [73], but this gene may be liable for pathogenesis of pituitary prolactinoma. CENPM, CENPI, CDCA5 and CENPU are novel biomarkers for pathogenesis of pituitary prolactinoma. Role of Ran in mitotic spindle is the most significant MSigDB C2 BIOCARTA pathway for down-regulated genes. RCC1 and KIF15 are novel biomarkers for pathogenesis of pituitary prolactinoma. Inflammation mediated by chemokine and cytokine signaling pathway is the most significant PantherDB pathway for down-regulated genes. PF4-active platelet accumulation in cancer is crucial because platelets can modulate cancer cells and the cancer microenvironment to stimulate lung Cancer outgrowth [74], but this gene may be answerable for development of pituitary prolactinoma. VWF, CCL3L3, CASK and PLCD4 are novel biomarkers for pathogenesis of pituitary prolactinoma. O-glycans biosynthetic is the most significant Pathway Ontology for down-regulated genes. B4GALT6 is novel biomarker for pathogenesis of pituitary prolactinoma. Eptifibatide pathway is the most significant SMPDB pathway for down-regulated genes. RYR2 is novel biomarker for pathogenesis of pituitary prolactinoma.

In present study, sensory organ morphogenesis is the most significant GO BP term for up-regulated genes. Epigenetic inactivation of tumor suppressor genes EYA4 and GAS1are associated with lung cancer and papillary thyroid carcinoma development, but silencing of these genes may associate with pituitary prolactinoma [75–76]. Expression of SOX9 enhances colorectal cancer cell invasion and migration [77], but this gene may link with pathogenesis of pituitary prolactinoma. ROR2 play a key role as an important mediator of the Wnt signaling pathway in colorectal cancer [78], but this gene may be involves in pathogenesis of pituitary prolactinoma through activation of Wnt signaling pathway. CLIC5, TMIE, CLRN1, PRRX2, SDK2, CELSR1, STRA6, CALB1 and VANGL2 are novel biomarkers for pathogenesis of pituitary prolactinoma. Extracellular matrix is the most significant GO CC term for up-regulated genes. FLRT2, CPXM2, LAD1 and TGFBR3 are novel biomarkers for pathogenesis of pituitary prolactinoma. Hormone activity is the most significant GO MF term for up-regulated genes. CARTPT, GPHA2 and GAL are novel biomarkers for pathogenesis of pituitary prolactinoma. Nuclear division is the most significant GO BP term for down-regulated genes. Elevated expression of NUSAP1 may improve cancer progression by increasing proliferation and invasion of prostate cancer cells [79], but this gene may be responsible for progression of pituitary prolactinoma. High expression of cell cycle regulatory protein CKS2 is culpable for advancement of gastric cancer [80], but this gene may be link with pathogenesis of pituitary prolactinoma. PBK is likely to play a crucial role in cell division and cytokinesis in breast cancer [81], but this gene may be liable for improvement of pituitary prolactinoma. Inappropriate expression of NEK2 interfere with mitotic processes results in breast cancer development [82], but this gene may be play key role in pathogenesis of pituitary prolactinoma. High expression of oncogene KIF14 may play crucial role in cancer development [83], but this gene may be associates with pituitary prolactinoma. Wong et al [84] and Loveday et al [85] demonstrated that modification of normal DNA repair function of RAD51 and RAD51D may lead to genomic instabilities that eventually contribute to breast and ovarian cancer development, but gene may be identify with pituitary prolactinoma. CCNG2 deeply involved in pancreatic cancer cell proliferation, invasion, chemoresistance, and differentiation [86], but this gene may be responsible for important for the pathogenesis of pituitary prolactinoma. FANCA, FANCD2, CLTC, MTBP, HAUS6, DLGAP5, RAD21L1, PTTG3P, KIF11, PRC1, EDN1 and CCNF are novel biomarkers for pathogenesis of pituitary prolactinoma. Condensed chromosome is the most significant GO CC term for down-regulated genes. HJURP is novel biomarker for pathogenesis of pituitary prolactinoma. Microtubule binding is the most significant GO MF term for down-regulated genes. KIF21B and KIF26A are novel biomarkers for pathogenesis of pituitary prolactinoma.

In present study, SOX2, KRT40, SMAD9, AMOT and DPPA4 were identified as hub proteins (up regulated DEGs) in the PPI network. AMOT control of the Hippo/LATS pathway in the processes of cell proliferation, motility, and differentiation in cancer [87], but this may be linked with pituitary prolactinoma. KRT40 is novel biomarkers for pathogenesis of pituitary prolactinoma. SOX2, KRT40, SMAD9, AMOT and TRIM29 are the hub proteins (up regulated DEGs) with highest betweenness centrality in the PPI network. High expression of TRIM29 is important in differentiation, proliferation, and development of gastric cancer [88], but this gene may be responsible for progression of pituitary prolactinoma. SOX2, KRT40, AMOT, C6orf141 and TRIM29 are the hub proteins (up regulated DEGs) with highest stress centrality in the PPI network. C6orf141 is novel biomarkers for pathogenesis of pituitary prolactinoma. SOX2, KRT40, SMAD9, TRIM29 and FGFR2 are the hub proteins (up regulated DEGs) with highest closeness centrality in the PPI network. PRSS45, CARTPT, TAC4, CT45A1 and PON3 are the hub proteins (up regulated DEGs) with lowest clustering coefficient in the PPI network. Shang et al [89] reported that proto-oncogene CT45A1 play key role cancer cell invasion, but this gene may be link with pituitary prolactinoma. PRSS45, TAC4 and PON3 are novel biomarkers for pathogenesis of pituitary prolactinoma. CLTC, PLK1, DHX15, GTSE1 and DISC1 are identified as hub proteins (down regulated DEGs) in the PPI network.

DHX15 play key role in cancer through activating AR activity through Siah2-mediated ubiquitination independent of its ATPase activity [90], but this gene may be liable for development of pituitary prolactinoma. Subhash et al [91] shown that GTSE1 is up-regulated in gastric cancer through DNA damage activated trans-activation of p53 is inhibited, thus affecting p53 activated apoptosis, but this gene may be answerable for development of pituitary prolactinoma. DISC1 is novel biomarker for pathogenesis of pituitary prolactinoma. CLTC, PLK1, DHX15, KIF11 and ATP2B2 are the hub proteins with highest betweenness centrality in the PPI network for down regulated DEGs. ATP2B2 is novel biomarker for pathogenesis of pituitary prolactinoma. CLTC, DHX15, PLK1, ATP2B3 and KIF11 are the hub proteins with highest stress centrality in the PPI network for down regulated DEGs. ATP2B3 is novel biomarker for pathogenesis of pituitary prolactinoma. PLK1, CLTC, ATP2B4, ATP2B8 and DHX15 are the hub proteins with highest closeness centrality in the PPI network for down regulated DEGs. ATP2B4 and ATP2B8 are novel biomarkers for pathogenesis of pituitary prolactinoma. B4GALT6, ZNF160, HIGD1B, CCL3L3 and C20orf203 are the hub proteins with lowest clustering coefficient in the PPI network for down regulated DEGs. ZNF160, HIGD1B and C20orf203 are novel biomarkers for pathogenesis of pituitary prolactinoma.

Modules are extracted from the PPI network for up and down regulated DEGs RUNX1, SOX2, FGFR2, FGF1, FGFR3, S100B, S100A1, SMAD9 and EVC2 are the hub proteins (up regulated DEGs with high degree) in all four modules in the PPI network. EVC2 is novel biomarker for pathogenesis of pituitary prolactinoma. CLTC, GTSE1, NUF2, BUB1, SPC24, SPC25, CKS2, CCNB2, CCNA1, KIF18A, FOXM1 and PRC1 are the hub proteins (down regulated DEGs with high degree) in all four modules in the PPI network. FOXM1 play roles in cancer related processes, such as invasion and metastasis [92], but this may be link with pathogenesis of pituitary prolactinoma.

LINC00598, CNKSR3, PMAIP1, TRIM71 and FAM83F are identified as up regulated target genes with high degree of connectivity in target gene – miRNA-regulatory network. LINC00598, CNKSR3, PMAIP1 and TRIM71 are novel biomarkers for pathogenesis of pituitary prolactinoma. SOX4, ZMAT3, PTP4A1, RAD51 and DAZAP2 are identified as down regulated target genes with high degree of connectivity in target gene-miRNA regulatory network. Oncogene SOX4 plays an essential role in the stimulation of and response to developmental pathways, such as Wnt, Notch, Hedgehog, and TGFβ in prostate cancer [93], but this gene may be associates with pituitary prolactinoma. PTP4A1 play key role in cancer cell growth and invasion of breast cancer cells [94], but this gene may be responsible for improvement of pituitary prolactinoma. ZMAT3 and DAZAP2 are novel biomarkers for pathogenesis of pituitary prolactinoma.

IRX1, CACNA2D3, VSNL1, BMP7 and DACT2 are identified as up regulated target gene with high degree of connectivity in TFs-target gene regulatory network. CACNA2D3 and DACT2 are tumor suppressor genes and epigenetic inactivation of this gene is linked with gastric and lung cancer [95–96], but suppression of these genes may be identify with pituitary prolactinoma. IRX1 and VSNL1 are novel biomarkers for pathogenesis of pituitary prolactinoma. UNC13A, CCNF, BDNF, POU3F4 and AKAP3 are identified as down regulated target genes with high degree of connectivity in TFs-target gene regulatory network. UNC13A, POU3F4 and AKAP3 are novel biomarkers for pathogenesis of pituitary prolactinoma.

## Methods

### Microarray Data

We chose a gene expression profile of GSE119063 from GEO database. GSE119063 was based on the Agilent GPL13607 platform (Agilent-028004 SurePrint G3 Human GE 8×60K Microarray (Feature Number version)). The GSE119063 dataset included 9 samples, containing 5 pituitary prolactinoma samples and 4 normal pituitaries samples. Besides, we downloaded the Series Matrix File of GSE119063 from GEO database.

### Data preprocessing

The raw data used in this study were downloaded. The raw GSE119063 data was preprocessed by the Limma package (http://www.bioconductor.org/packages/release/bioc/html/limma.html) [97] in Bioconductor. The data preprocessing included background correction and quantile normalization. Probe identities (IDs) are mapped to gene IDs using the corresponding platform files.

### Identification of DEGs

The Limma package was subsequently used for identifying DEGs. p < 0.05 and absolute fold change ≥ 0.93 for up regulated gene and fold change ≥ -0.29 for down regulated gene [98] were considered as the cutoff values for DEG screening using the Benjamini & Hochberg procedure. R software was used to produce heat maps of common significant differentially expressed genes between pituitary prolactinoma samples and normal pituitaries samples. Genes are ordered according to the fold change in the expression values. This information was presented as a heat map and a volcano plot.

### Pathway enrichment analysis of DEGs

BIOCYC (https://biocyc.org/) [99], Kyoto Encyclopedia of Genes and Genomes (KEGG) (http://www.genome.jp/kegg/pathway.html) [100], Pathway Interaction Database (PID) (https://wiki.nci.nih.gov/pages/viewpage.action?pageId=315491760) [101], REACTOME (https://reactome.org/) [102], GenMAPP (http://www.genmapp.org/) [103], MSigDB C2 BIOCARTA (http://software.broadinstitute.org/gsea/msigdb/collections.jsp) [104], PantherDB (http://www.pantherdb.org/) [105], Pathway Ontology (http://www.obofoundry.org/ontology/pw.html) [106] and Small Molecule Pathway Database (SMPDB) (http://smpdb.ca/) [107] were a collection of databases which helps to handle genomes, biological pathways, diseases, chemical substances, and drugs. ToppGene (https://toppgene.cchmc.org/enrichment.jsp) is a web-based online bioinformatics resource that aims to provide tools for the functional interpretation of large lists of genes or proteins [108]. p value < 0.05 is regarded as the cutoff criterion. We could visualize the pathways among those DEGs using ToppGene.

### Gene Ontology (GO) enrichment analysis of DEGs

Gene ontology (GO) (http://www.geneontology.org/) enrichment analysis served as a useful approach to annotate genes and gene products and also analyze characteristic biological attributing to high-throughput genome or transcriptome data [109]. ToppGene (https://toppgene.cchmc.org/enrichment.jsp) is a web-based online bioinformatics resource that aims to provide tools for the functional interpretation of large lists of genes or proteins [108]. P value < 0.05 is regarded as the cutoff criterion. We could visualize the core biological process (BP), molecular function (MF) and cellular component (CC) among those DEGs using ToppGene.

### PPI network construction and topology analysis

The Human Integrated Protein-Protein Interaction rEference (HIPPIE) (http://cbdm.uni-mainz.de/hippie/) is an online tool providing experimental and predicted PPI information [110] through interfacing different data bases such as IntAct Molecular Interaction Database (https://www.ebi.ac.uk/intact/) [111], Biological General Repository for Interaction Datasets (BioGRID) (https://thebiogrid.org/) [112], The Human Protein Reference Database (HPRD) (http://www.hprd.org/) [113], the Molecular INTeraction database (MINT) (https://mint.bio.uniroma2.it/) [114], The Biomolecular Interaction Network Database (BIND) (http://baderlab.org/BINDTranslation) [115], MIPS (http://mips.helmholtz-muenchen.de/proj/ppi/) [116] and DIP (http://dip.doe-mbi.ucla.edu/dip/Main.cgi) [117]. In this study, the HIPPIE [110] was used to analyze the PPIs among the proteins encoded by the DEGs, then the PPI networks for the up-regulated and the down-regulated genes are separately visualized by Cytoscape version 3.5.1 software (http://www.cytoscape.org/) [118]. The degree of a gene in a PPI network is equal to the number of edges containing that node [119]. Betweenness centrality of a gene which is located on the shortest path between two other genes has most influence over the “information transfer” between them [120]. Stress centrality is number of genes in the shortest path between two other genes [121]. Closeness centrality is an inverse of the average length of the shortest paths to/from all the other genes in the PPI network [122]. Cluster coefficient measures the density of interactions in the network neighborhood of a gene [123].

### Module analysis

In PPI networks, genes in the same module typically show the same or similar function and work together to implement their biological function. To visualize the network and identify the modules in the network, PEWCC1 java plug-in [124] on the Cytospace software (www.cytoscape.org/) [118] was used. The parameters were set as follows: Degree cutoff ≥ 10 (degrees of each node in module were at least larger than 2), K-core ≥ 2 (subgraphs of each node in module were at least 2 and more than 2).

### Construction of the target gene **-** miRNA network

The NetworkAnalyst (http://www.networkanalyst.ca/) is a online tool available comprehensive resource containing the predicted and the experimentally validated target gene-miRNA interaction pairs [125]. The DEGs -associated predicted miRNA were selected when they were included two TarBase (http://diana.imis.athena-innovation.gr/DianaTools/index.php?r=tarbase/index) [126] and miRTarBase (http://mirtarbase.mbc.nctu.edu.tw/php/download.php) [127]. Subsequently, the overlapping target genes were identified and the gene-miRNA pair was selected. The gene-miRNA network was formed and visualized using the Cytoscape version 3.5.1 software (http://www.cytoscape.org/) [118].

### Construction of the target gene **-** TF network

The DEGs and transcription factors (TFs) that potentially regulated the DEGs are predicted using Overrepresentation Enrichment Analysis (ORA) in NetworkAnalyst (http://www.networkanalyst.ca/) [125]. The DEGs-associated predicted TF were selected when they were included database such as ENCODE (http://cistrome.org/BETA/) [128]. Then gene-TF-network are also visualized using version 3.5.1 software (http://www.cytoscape.org/) [118].

## Conclusion

We use bioinformatics analysis of pituitary prolactinoma to investigate the biological and clinical value genes. Finally, using a series of particular conditions we screened crucial genes from DEGs. These findings may improve our understanding of the etiology, pathology, and the potential molecular mechanisms and gene targets of pituitary prolactinoma, which may be beneficial for the identification of diagnostic biomarkers and treatment methods for pituitary prolactinoma. Nevertheless, lacking of experimental verification is a limitation of this study. Further molecular biological experiments *in vivo* and *in vitro* are required to confirm the function of the identified genes in pituitary prolactinoma.

## Abbreviations

BioGRID: Biological General Repository for Interaction Datasets
BP: biological process
CC: cellular component
DEGs: Differentially Expressed Genes
GEO: Gene Expression Omnibus
GO: Gene ontology
HIPPIE: Human Integrated Protein-Protein Interaction rEference
KEGG: Kyoto Encyclopedia of Genes and Genomes
MF: molecular function
SMPDB: Small Molecule Pathway Database

## Acknowledgement

I thank Ni Li, Institute of Health Sciences, Shanghai Institute for Biological Sciences, Chinese Academy of Sciences, China, very much, the author who deposited their microarray dataset, GSE119063, into the public GEO database.

## Ethical approval

This article does not contain any studies with human participants or animals performed by any of the authors.

## Informed consent

No informed consent because this study does not contain human or animals participants.

## Author Contributions

Vikrant Ghatnatti - Methodology and validation Basavaraj Vastrad - Writing original draft, investigation, and review and editing Swetha Patil **-** Formal analysis and validation Chanabasayya Vastrad - Software and investigation Iranna Kotturshetti - Supervision and resources

## Availability of data and materials

The datasets supporting the conclusions of this article are available in the GEO (Gene Expression Omnibus) (https://www.ncbi.nlm.nih.gov/geo/) repository. [(GSE119063) (https://www.ncbi.nlm.nih.gov/geo/query/acc.cgi?acc=GSE119063)]

## Consent for publication

Not applicable.

## References

1. Thorner, M.O.; Martin, W.H.; Rogol, A.D.; Morris, J.L.; Perryman, R.L.; Conway, B.P.; Howards, S.S.; Wolfman, M.G.; MacLeod, R.M. Rapid regression of pituitary prolactinomas during bromocriptine treatment. J Clin Endocrinol Metab. 1980,51,438–445. doi:10.1210/jcem-51-3-438

2. Doumith, R.; Gennes, J.L.; Cabane, J.P.; Zygelman, N. Pituitary prolactinoma, adrenal aldosterone-producing adenomas, gastric schwannoma and colonic polyadenomas: a possible variant of multiple endocrine neoplasia (MEN) type I. Acta Endocrinol (Copenh). 1982,100,189–195.

3. Oruçkaptan, H.H.; Senmevsim, O.; Ozcan, O.E.; Ozgen, T. Pituitary adenomas: results of 684 surgically treated patients and review of the literature. Surg Neurol. 2000,53,211–219.

4. Cho, D.Y.; Liau, W.R. Comparison of endonasal endoscopic surgery and sublabial microsurgery for prolactinomas. Surg Neurol. 2002;58,371–375

5. Murakami, M.; Mizutani, A.; Asano, S.; Katakami, H.; Ozawa, Y.; Yamazaki, K.; Ishida, Y.; Takano, K.; Okinaga, H.; Matsuno, A. A mechanism of acquiring temozolomide resistance during transformation of atypical prolactinoma into prolactin-producing pituitary carcinoma: case report. Neurosurgery. 2011,68,E1761–E1767. doi:10.1227/NEU.0b013e318217161a

6. Tsang, R.W.; Laperriere, N.J.; Simpson, W.J.; Brierley, J.; Panzarella, T.; Smyth, H.S. Glioma arising after radiation therapy for pituitary adenoma. A report of four patients and estimation of risk. Cancer. 1993;72,2227–2233.

7. Friedman, E.; Adams, E.F.; Höög, A,; Gejman, P.V.; Carson, E.; Larsson, C.; De Marco, L.; Werner, S.; Fahlbusch, R.; Nordenskjöld, M. Normal structural dopamine type 2 receptor gene in prolactin-secreting and other pituitary tumors. J Clin Endocrinol Metab. 1994,78,568–574. doi:10.1210/jcem.78.3.7907340

8. Fedele, M.; Pentimalli, F.; Baldassarre, G.; Battista, S.; Klein-Szanto, A.J.; Kenyon, L.; Visone, R.; De Martino, I.; Ciarmiello, A.; Arra, C.;, et al. Transgenic mice overexpressing the wild-type form of the HMGA1 gene develop mixed growth hormone/prolactin cell pituitary adenomas and natural killer cell lymphomas. Oncogene. 2005,24,3427–3435. doi:10.1038/sj.onc.1208501

9. Finelli, P.; Pierantoni, G.M.; Giardino, D.; Losa, M.; Rodeschini, O.; Fedele, M.; Valtorta, E.; Mortini, P.; Croce, C.M.; Larizza, L.;, et al. The High Mobility Group A2 gene is amplified and overexpressed in human prolactinomas. Cancer Res. 2002,62,2398–2405.

10. Paez-Pereda, M.; Giacomini, D.; Refojo, D.; Nagashima, A.C.; Hopfner, U.; Grubler, Y.; Chervin, A.; Goldberg, V.; Goya, R.; Hentges, S.T.;, et al. Involvement of bone morphogenetic protein 4 (BMP-4) in pituitary prolactinoma pathogenesis through a Smad/estrogen receptor crosstalk. Proc Natl Acad Sci U S A. 2003,100,1034–1039. doi:10.1073/pnas.0237312100

11. Shimon, I.; Hinton, D.R.; Weiss, M.H.; Melmed, S. Prolactinomas express human heparin-binding secretory transforming gene (hst) protein product: marker of tumour invasiveness. Clin Endocrinol (Oxf). 1998;48(1):23–29.

12. Lania, A.G.; Ferrero, S.; Pivonello, R.; Mantovani, G.; Peverelli, E.; Di Sarno, A.; Beck-Peccoz, P.; Spada, A.; Colao, A. Evolution of an aggressive prolactinoma into a growth hormone secreting pituitary tumor coincident with GNAS gene mutation. J Clin Endocrinol Metab. 2010,95,13–17. doi:10.1210/jc.2009-1360

13. Dworakowska, D.; Wlodek, E.; Leontiou, C.A.; Igreja, S.; Cakir, M.; Teng, M.; Prodromou, N.; Góth, M.I.; Grozinsky-Glasberg, S.; Gueorguiev, M.;, et al. Activation of RAF/MEK/ERK and PI3K/AKT/mTOR pathways in pituitary adenomas and their effects on downstream effectors. Endocr Relat Cancer. 2009,16,1329–1338. doi:10.1677/ERC-09-0101

14. Semba, S.; Han, S.Y.; Ikeda, H.; Horii, A. Frequent nuclear accumulation of beta-catenin in pituitary adenoma. Cancer. 2001,91,42–48.

15. Seemann, N.; Kuhn, D.; Wrocklage, C.; Keyvani, K.; Hackl, W.; Buchfelder, M.; Fahlbusch, R.; Paulus, W. CDKN2A/p16 inactivation is related to pituitary adenoma type and size. J Pathol. 2001,193,491–497. doi:10.1002/path.833

16. Ozfirat, Z.; Korbonits, M. AIP gene and familial isolated pituitary adenomas. Mol Cell Endocrinol. 2010,326,71–79. doi:10.1016/j.mce.2010.05.001

17. Lock, C.; Hermans, G.; Pedotti, R.; Brendolan, A.; Schadt, E.; Garren, H.; Langer-Gould, A.; Strober, S.; Cannella, B.; Allard, J.;, et al. Gene-microarray analysis of multiple sclerosis lesions yields new targets validated in autoimmune encephalomyelitis. Nat Med. 2002,8,500–508. doi:10.1038/nm0502-500

18. Ma, C.; Wang, F.; Han, B.; Zhong, X.; Si, F.; Ye, J.; Hsueh, E.C.; Robbins, L.; Kiefer, S.M.; Zhang, Y.;, et al. SALL1 functions as a tumor suppressor in breast cancer by regulating cancer cell senescence and metastasis through the NuRD complex. Mol Cancer. 2018,17,78. doi:10.1186/s12943-018-0824-y

19. Wang, Y.C.; Gallego-Arteche, E.; Iezza, G.; Yuan, X.; Matli, M.R.; Choo, S.P.; Zuraek, M.B.; Gogia, R.; Lynn, F.C.; German, M.S.;, et al. Homeodomain transcription factor NKX2.2 functions in immature cells to control enteroendocrine differentiation and is expressed in gastrointestinal neuroendocrine tumors. Endocr Relat Cancer. 2009,16,267–279. doi:10.1677/ERC-08-0127

20. Alarmo, E.L. Pärssinen, J. Ketolainen, J.M. Savinainen, K. Karhu R. Kallioniemi, A. BMP7 influences proliferation, migration, and invasion of breast cancer cells. Cancer Lett. 2009,275,35–43. doi:10.1016/j.canlet.2008.09.028

21. Cipriano, R.; Miskimen, K.L.; Bryson, B.L.; Foy, C.R.; Bartel, C.A.; Jackson, M.W. Conserved oncogenic behavior of the FAM83 family regulates MAPK signaling in human cancer. Mol Cancer Res. 2014,12,1156–1165. doi:10.1158/1541-7786.MCR-13-0289

22. Quan, Y.; Xu, M.; Cui, P.; Ye, M.; Zhuang, B.; Min, Z. Grainyhead-like 2 Promotes Tumor Growth and is Associated with Poor Prognosis in Colorectal Cancer. J Cancer. 2015;6,342–350. doi:10.7150/jca.10969

23. Li, H. Sun, L. Tang, Z. Fu, L. Xu, Y. Li, Z. Luo, W. Qiu, X. Wang, E. Overexpression of TRIM24 correlates with tumor progression in non-small cell lung cancer. PLoS One. 2012,7,e37657. doi:10.1371/journal.pone.0037657

24. Kaistha, B.P.; Honstein, T.; Müller, V.; Bielak, S.; Sauer, M.; Kreider, R.; Fassan, M.; Scarpa, A.; Schmees, C.; Volkmer, H.;, et al. Key role of dual specificity kinase TTK in proliferation and survival of pancreatic cancer cells. Br J Cancer. 2014;111,1780–1787. doi:10.1038/bjc.2014.460

25. Di Leo, A.; Desmedt, C.; Bartlett, J.M.; Piette, F.; Ejlertsen, B.; Pritchard, K.I.; Larsimont, D.; Poole, C.; Isola, J.; Earl, H.;, et al. HER2 and TOP2A as predictive markers for anthracycline-containing chemotherapy regimens as adjuvant treatment of breast cancer: a meta-analysis of individual patient data. Lancet Oncol. 2011,12,1134–1142. doi:10.1016/S1470-2045(11)70231-5

26. Zhang, C.; Zhu, C;, Chen, H.; Li, L.; Guo, L.; Jiang, W.; Lu, S.H. Kif18A is involved in human breast carcinogenesis. Carcinogenesis. 2010,31,1676–1684. doi:10.1093/carcin/bgq134

27. Aguirre-Portolés C, Bird AW, Hyman A, Cañamero M, Pérez de Castro I, Malumbres M. Tpx2 controls spindle integrity, genome stability, and tumor development. Cancer Res. 2012,72,1518–1528. doi:10.1158/0008-5472.CAN-11-1971

28. Zhu, X.; Lee, K.; Asa, S.L.; Ezzat, S. Epigenetic silencing through DNA and histone methylation of fibroblast growth factor receptor 2 in neoplastic pituitary cells. Am J Pathol. 2007,170,1618–1628. doi:10.2353/ajpath.2007.061111

29. Alatzoglou, K.S. Andoniadou, C.L. Kelberman, D. Buchanan, C.R. Crolla, J. Arriazu, M.C. Roubicek, M. Moncet, D. Martinez-Barbera, J.P. Dattani, M.T. SOX2 haploinsufficiency is associated with slow progressing hypothalamo-pituitary tumours. Hum Mutat. 2011,32,1376–1380. doi:10.1002/humu.21606

30. Heinrichs, M. Baumgärtner, W. Capen, C.C. Immunocytochemical demonstration of proopiomelanocortin-derived peptides in pituitary adenomas of the pars intermedia in horses. Vet Pathol. 1990,27,419–425. doi:10.1177/030098589902700606

31. Katznelson, L. Alexander, J.M. Bikkal, H.A. Jameson, J.L. Hsu, D.W. Klibanski, A. Imbalanced follicle-stimulating hormone beta-subunit hormone biosynthesis in human pituitary adenomas. J Clin Endocrinol Metab. 1992;74,1343–1351. doi:10.1210/jcem.74.6.1375599

32. Duong, C.V. Yacqub-Usman, K. Emes, R.D. Clayton, R.N. Farrell, W.E. The EFEMP1 gene: a frequent target for epigenetic silencing in multiple human pituitary adenoma subtypes. Neuroendocrinology. 2013;98,200–211. doi:10.1159/000355624

33. Wu, Y, Bai, J, Hong, L, Yu, S, Yu, G, Zhang, Y. Low expression of secreted frizzled-related protein 2 and nuclear accumulation of β-catenin in aggressive nonfunctioning pituitary adenoma. Oncol Lett. 2016;12,199–206. doi:10.3892/ol.2016.4560

34. Lekva, T.; Berg, J.P.; Lyle, R.; Heck, A.; Bollerslev, J.; Ueland, T. Alternative splicing of placental lactogen (CSH2) in somatotroph pituitary adenomas. Neuro Endocrinol Lett. 2015,36,136–142.

35. Yavropoulou, M.P.; Maladaki, A.; Topouridou, K.; Kotoula, V.; Poulios, C.; Daskalaki, E.; Foroglou, N.; Karkavelas, G.; Yovos, J.G. Expression pattern of the Hedgehog signaling pathway in pituitary adenomas. Neurosci Lett. 2016,611,94–100. doi:10.1016/j.neulet.2015.10.076

36. Giustina, A.; Bonfanti, C.; Licini, M.; De Rango, C.; Milani, G. Inhibitory effect of galanin on growth hormone release from rat pituitary tumor cells (GH1) in culture. Life Sci. 1994,55,1845–1851.

37. Hunter, J.A.; Skelly, R.H.; Aylwin, S.J.; Geddes, J.F.; Evanson, J.; Besser, G.M.; Monson, J.P.; Burrin, J.M. The relationship between pituitary tumour transforming gene (PTTG) expression and in vitro hormone and vascular endothelial growth factor (VEGF) secretion from human pituitary adenomas. Eur J Endocrinol. 2003,148,203–211.

38. De Martino, I.; Visone, R.; Wierinckx, A.; Palmieri, D.; Ferraro, A.; Cappabianca, P.; Chiappetta, G.; Forzati, F.; Lombardi, G.; Colao, A.;, et al. HMGA proteins up-regulate CCNB2 gene in mouse and human pituitary adenomas. Cancer Res. 2009,69,1844–1850. doi:10.1158/0008-5472.CAN-08-4133

39. Wierinckx, A.; Auger, C.; Devauchelle, P.; Reynaud, A.; Chevallier, P.; Jan, M.; Perrin, G.; Fèvre-Montange, M.; Rey, C.; Figarella-Branger, D.;, et al. A diagnostic marker set for invasion, proliferation, and aggressiveness of prolactin pituitary tumors. Endocr Relat Cancer. 2007;14,887–900. doi:10.1677/ERC-07-0062

40. Tecimer, T. Dlott, J. Chuntharapai, A. Martin, A.W. Peiper, S.C. Expression of the chemokine receptor CXCR2 in normal and neoplastic neuroendocrine cells. Arch Pathol Lab Med. 2000;124,520–525.doi:10.1043/0003-9985(2000)124<0520:EOTCRC>2.0.CO;2

41. Grizzi, F. Borroni, E.M. Vacchini, A. Qehajaj, D. Liguori, M. Stifter, S. Chiriva-Internati, M. Di Ieva, A. Pituitary Adenoma and the Chemokine Network: A Systemic View. Front Endocrinol (Lausanne). 2015,6,141. doi:10.3389/fendo.2015.00141

42. Egashira, N.; Takekoshi, S.; Takei, M.; Teramoto, A.; Osamura, R.Y. Expression of FOXL2 in human normal pituitaries and pituitary adenomas. Mod Pathol. 2011,24,765–773. doi:10.1038/modpathol.2010.169

43. Zhang, H.Y. Jin, L. Stilling, G.A. Ruebel, K.H. Coonse, K. Tanizaki, Y. Raz, A. Lloyd, R.V. RUNX1 and RUNX2 upregulate Galectin-3 expression in human pituitary tumors. Endocrine. 2009,35,101–111. doi:10.1007/s12020-008-9129-z

44. Tampanaru-Sarmesiu, A, Stefaneanu, L, Thapar, K, Kontogeorgos, G, Sumi, T, Kovacs, K. Transferrin and transferrin receptor in human hypophysis and pituitary adenomas. Am J Pathol. 1998;152,413–422.

45. Rehfeld, J.F.; Lindholm, J.; Andersen, B.N.; Bardram, L.; Cantor, P.; Fenger, M.; Lüdecke, D.K. Pituitary tumors containing cholecystokinin. N Engl J Med. 1987;316,1244–1247. doi:10.1056/NEJM198705143162004

46. Shaima, J.; William, B.; Omkaram, G.;, et al. Developmental pluripotency associated 4: A novel putative predictor for prognosis of aggressive prolactin secreting tumors in the pituitary. Cancer Research. 2017,77.

47. Missale, C. Losa, M. Sigala, S. Balsari, A. Giovanelli, M. Spano, P.F. Nerve growth factor controls proliferation and progression of human prolactinoma cell lines through an autocrine mechanism. Mol Endocrinol. 1996,10,272–285. doi:10.1210/mend.10.3.8833656

48. Tomlinson, D.C. Baldo, O. Harnden, P. Knowles, M.A. FGFR3 protein expression and its relationship to mutation status and prognostic variables in bladder cancer. J Pathol. 2007,213,91–98. doi:10.1002/path.2207

49. Shojima, K.; Sato, A.; Hanaki, H.; Tsujimoto, I.; Nakamura, M.; Hattori, K.; Sato, Y.; Dohi, K.; Hirata, M.; Yamamoto, H.;, et al. Wnt5a promotes cancer cell invasion and proliferation by receptor-mediated endocytosis-dependent and -independent mechanisms, respectively. Sci Rep. 2015;5:8042. doi:10.1038/srep08042

50. Sharma P, Chinaranagari S, Patel D, et al. Epigenetic inactivation of inhibitor of differentiation 4 (Id4) correlates with prostate cancer. Cancer Med. 2012,1,176–186. doi:10.1002/cam4.16

51. Seder, C.W.; Hartojo, W.; Lin, L.; Carey, J.; Chaudhary, J. Upregulated INHBA expression may promote cell proliferation and is associated with poor survival in lung adenocarcinoma. Neoplasia. 2009;11(4):388–396.

52. Salem, C.E.; Markl, I.D.; Bender, C.M.; Gonzales, F.A.; Jones, P.A.; Liang, G. PAX6 methylation and ectopic expression in human tumor cells. Int J Cancer. 2000,87,179–185.

53. Duan, J.J.; Cai, J.; Guo, Y.F.; Bian, X.W.; Yu, S.C.; ALDH1A3, a metabolic target for cancer diagnosis and therapy. Int J Cancer. 2016,139,965–975. doi:10.1002/ijc.30091

54. Xu, C.; Chen, H.; Wang, X.; Gao, J.; Che, Y.; Li, Y.; Ding, F.; Luo, A.; Zhang, S.; Liu, Z. S100A14, a member of the EF-hand calcium-binding proteins, is overexpressed in breast cancer and acts as a modulator of HER2 signaling. J Biol Chem. 2014,289,827–837. doi:10.1074/jbc.M113.469718

55. DeRycke, M.S.; Andersen, J.D.; Harrington, K.M.; Pambuccian, S.E.; Kalloger, S.E.; Boylan, K.L.; Argenta, P.A.; Skubitz, A.P.; S100A1 expression in ovarian and endometrial endometrioid carcinomas is a prognostic indicator of relapse-free survival. Am J Clin Pathol. 2009,132,846–856. doi:10.1309/AJCPTK87EMMIKPFS

56. Harpio, R.; Einarsson, R. S100 proteins as cancer biomarkers with focus on S100B in malignant melanoma. Clin Biochem. 2004,37,512–528. doi:10.1016/j.clinbiochem.2004.05.012

57. Chen, H.; Suzuki, M.; Nakamura, Y.; Ohira, M.; Ando, S.; Iida, T.; Nakajima, T.; Nakagawara, A.; Kimura, H. Aberrant methylation of FBN2 in human non-small cell lung cancer. Lung Cancer. 2005,50,43–49. doi:10.1016/j.lungcan.2005.04.013

58. Yoshimura, N.; Sano, H.; Hashiramoto, A.; Yamada, R.; Nakajima, H.; Kondo, M.; Oka, T. The expression and localization of fibroblast growth factor-1 (FGF-1) and FGF receptor-1 (FGFR-1) in human breast cancer. Clin Immunol Immunopathol. 1998,89,28–34.

59. Dickinson, R.E. Dallol, A. Bieche, I. Krex, D. Morton, D. Maher, E.R. Latif, F. Epigenetic inactivation of SLIT3 and SLIT1 genes in human cancers. Br J Cancer. 2004,91,2071–2078. doi:10.1038/sj.bjc.6602222

60. Li, Z.; Zhang, W.; Shao, Y.; Zhang, C.; Wu, Q.; Yang, H.; Wan, X.; Zhang, J.; Guan, M.; Wan, J.;, et al. High-resolution melting analysis of ADAMTS18 methylation levels in gastric, colorectal and pancreatic cancers. Med Oncol. 2010;27,998–1004. doi:10.1007/s12032-009-9323-8

61. Pei Y.F.; Zhang Y.J.; Lei Y.; Wu D.W.; Ma T.H.; Liu X.Q. Hypermethylation of the CHRDL1 promoter induces proliferation and metastasis by activating Akt and Erk in gastric cancer. Oncotarget. 2017,8,23155–23166. doi:10.18632/oncotarget.15513

62. Jacobs E.T.; Van Pelt C.; Forster R.E.; Zaidi W.; Hibler E.A.; Galligan M.A.; Haussler M.R.; Jurutka P.W. CYP24A1 and CYP27B1 polymorphisms modulate vitamin D metabolism in colon cancer cells. Cancer Res. 2013,73,2563–2573. doi:10.1158/0008-5472.CAN-12-4134

63. Liu, X.; Erikson, R. Polo-like kinase 1 in the life and death of cancer cells. Cell Cycle. 2003,2,424–425.

64. Grabsch, H, Takeno, S, Parsons, WJ, Pomjanski, N, Boecking, A, Gabbert, HE, Mueller, W. Overexpression of the mitotic checkpoint genes BUB1, BUBR1, and BUB3 in gastric cancer--association with tumour cell proliferation. J Pathol. 2003,200,16–22. doi:10.1002/path.1324

65. Kitkumthorn, N.; Yanatatsanajit, P.; Kiatpongsan, S.; Phokaew, C.; Triratanachat, S.; Trivijitsilp, P.; Termrungruanglert, W.; Tresukosol, D.; Niruthisard, S.; Mutirangura, A. Cyclin A1 promoter hypermethylation in human papillomavirus-associated cervical cancer. BMC Cancer. 2006,6,55. doi:10.1186/1471-2407-6-55

66. Kato, T.; Wada, H.; Patel, P.; Hu, H.P.; Lee, D.; Ujiie, H.; Hirohashi, K.; Nakajima, T.; Sato, M.; Kaji, M.;, et al. Overexpression of KIF23 predicts clinical outcome in primary lung cancer patients. Lung Cancer. 2016,92:53–61. doi:10.1016/j.lungcan.2015.11.018

67. Cao, L.; Li, C.; Shen, S.; Yan, Y.; Ji, W.; Wang, J.; Qian, H.; Jiang, X.; Li, Z.; Wu, M.;, et al. OCT4 increases BIRC5 and CCND1 expression and promotes cancer progression in hepatocellular carcinoma. BMC Cancer. 2013,13,82. doi:10.1186/1471-2407-13-82

68. Zhang, Q.;Su, R.; Shan, C.; Gao, C.; Wu, P. Non-SMC Condensin I Complex, Subunit G (NCAPG) is a Novel Mitotic Gene Required for Hepatocellular Cancer Cell Proliferation and Migration. Oncol Res. 2018;26,269–276. doi:10.3727/096504017X15075967560980

69. Nie, W, Xu, MD, Gan, L, Huang, H, Xiu, Q, Li, B. Overexpression of stathmin 1 is a poor prognostic biomarker in non-small cell lung cancer. Lab Invest. 2015;95,56–64. doi:10.1038/labinvest.2014.124

70. Imai, K.; Hirata, S.; Irie, A.; Senju, S.; Ikuta, Y.; Yokomine, K.; Harao, M.; Inoue, M.; Tomita, Y.; Tsunoda, T.;, et al. Identification of HLA-A2-restricted CTL epitopes of a novel tumour-associated antigen, KIF20A, overexpressed in pancreatic cancer. Br J Cancer. 2011,104,300–307. doi:10.1038/sj.bjc.6606052

71. Zhou, J.; Yu, Y.; Pei, Y.; Cao, C.; Ding, C.; Wang, D.; Sun, L.; Niu, G. A potential prognostic biomarker SPC24 promotes tumorigenesis and metastasis in lung cancer. Oncotarget. 2017,8,65469–65480. doi:10.18632/oncotarget.18971

72. He, W.L.; Li, Y.H.; Yang, D.J.; Song, W.; Chen, X.L.; Liu, F.K.; Wang, Z.; Li, W.; Chen, W.; Chen, C.Y.;, et al. Combined evaluation of centromere protein H and Ki-67 as prognostic biomarker for patients with gastric carcinoma. Eur J Surg Oncol. 2013,39,141–149. doi:10.1016/j.ejso.2012.08.023

73. Hu, P.; Shangguan, J.; Zhang, L. Down regulation of NUF2 inhibits tumor growth and induces apoptosis by regulating lncRNA AF339813. Int J Clin Exp Pathol. 2015,8,2638–2648.

74. Pucci, F.; Rickelt, S,; Newton, A.P.; Garris, C.; Nunes, E.; Evavold, C.; Pfirschke, C.; Engblom, C.; Mino-Kenudson, M,;, Hynes, R.O.;, et al. PF4 Promotes Platelet Production and Lung Cancer Growth. Cell Rep. 2016;17,1764–1772. doi:10.1016/j.celrep.2016.10.031

75. Wilson, I.M.; Vucic, E.A.; Enfield, K.S.; Thu, K.L.; Zhang, Y.A.; Chari, R.; Lockwood, W.W.; Radulovich, N.; Starczynowski, D.T.; Banáth, J.P.;, et al. EYA4 is inactivated biallelically at a high frequency in sporadic lung cancer and is associated with familial lung cancer risk. Oncogene. 2014,33,4464–4473. doi:10.1038/onc.2013.396

76. Ma, Y.; Qin, H.; Cui, Y. MiR-34a targets GAS1 to promote cell proliferation and inhibit apoptosis in papillary thyroid carcinoma via PI3K/Akt/Bad pathway. Biochem Biophys Res Commun. 2013;441,958–963. doi:10.1016/j.bbrc.2013.11.010

77. Xu, Y.; Zhang, X.; Hu, X.; Zhou, W.; Zhang, P.; Zhang, J.; Yang, S.; Liu, Y. The effects of lncRNA MALAT1 on proliferation, invasion and migration in colorectal cancer through regulating SOX9. Mol Med. 2018;24,52. doi:10.1186/s10020-018-0050-5

78. Mei, H.; Lian, S.; Zhang, S.; Wang, W.; Mao, Q.; Wang, H. High expression of ROR2 in cancer cell correlates with unfavorable prognosis in colorectal cancer. Biochem Biophys Res Commun. 2014,453,703–709. doi:10.1016/j.bbrc.2014.09.141

79. Gordon, C.A.; Gulzar, Z.G.; Brooks, J.D. NUSAP1 expression is upregulated by loss of RB1 in prostate cancer cells. Prostate. 2015,75,517–526. doi:10.1002/pros.22938

80. Kang, M.A.; Kim, J.T.; Kim, J.H.; Kim, S.Y.; Kim, Y.H.;, Yeom, Y.I.; Lee, Y.; Lee, H.G. Upregulation of the cycline kinase subunit CKS2 increases cell proliferation rate in gastric cancer. J Cancer Res Clin Oncol. 2009;135,761–769. doi:10.1007/s00432-008-0510-3

81. Park, J.H.; Lin, M.L.; Nishidate, T.; Nakamura, Y.; Katagiri, T. PDZ-binding kinase/T-LAK cell-originated protein kinase, a putative cancer/testis antigen with an oncogenic activity in breast cancer. Cancer Res. 2006,66,9186–9195. doi:10.1158/0008-5472.CAN-06-1601

82. Hayward, D.G.; Clarke, R.B.; Faragher, A.J.; Pillai, M.R.; Hagan, I.M.; Fry, A.M. The centrosomal kinase Nek2 displays elevated levels of protein expression in human breast cancer. Cancer Res. 2004,64,7370–7376. doi:10.1158/0008-5472.CAN-04-0960

83. Corson, T.W.; Huang, A.; Tsao, M.S.; Gallie, B.L. KIF14 is a candidate oncogene in the 1q minimal region of genomic gain in multiple cancers. Oncogene. 2005,24,4741–4753. doi:10.1038/sj.onc.1208641

84. Wong, A.K.; Pero, R.; Ormonde, P.A.; Tavtigian, S.V.; Bartel, P.L. RAD51 interacts with the evolutionarily conserved BRC motifs in the human breast cancer susceptibility gene brca2. J Biol Chem. 1997,272,31941–31944.

85. Loveday, C.; Turnbull, C.; Ramsay, E.; Hughes, D.; Ruark, E.; Frankum, J.R.; Bowden, G.; Kalmyrzaev, B.; Warren-Perry, M.; Snape, K.;, et al. Germline mutations in RAD51D confer susceptibility to ovarian cancer. Nat Genet. 2011,43,879–882. doi:10.1038/ng.893

86. Hasegawa, S.; Eguchi, H.; Nagano, H.; Konno, M.; Tomimaru, Y.; Wada, H.; Hama, N.; Kawamoto, K.; Kobayashi, S.; Nishida, N.;, et al. MicroRNA-1246 expression associated with CCNG2-mediated chemoresistance and stemness in pancreatic cancer. Br J Cancer. 2014,111,1572–1580. doi:10.1038/bjc.2014.454

87. Paramasivam M, Sarkeshik A, Yates JR et al Angiomotin family proteins are novel activators of the LATS2 kinase tumor suppressor. Mol Biol Cell. 2011;22(19):3725–3733. doi:10.1091/mbc.E11-04-0300

88. Kosaka, Y.; Inoue, H.; Ohmachi, T.; Fernandes, M.J.; McCollum, D. Tripartite motif-containing 29 (TRIM29) is a novel marker for lymph node metastasis in gastric cancer. Ann Surg Oncol. 2007;14,2543–2549. doi:10.1245/s10434-007-9461-1

89. Shang, B.; Gao, A.; Pan, Y.; Zhang, G.; Tu, J.; Zhou, Y.; Yang, P.; Cao, Z.; Wei, Q.; Ding, Y.;, et al. CT45A1 acts as a new proto-oncogene to trigger tumorigenesis and cancer metastasis. Cell Death Dis. 2014,5,e1285. doi:10.1038/cddis.2014.244

90. Jing, Y.; Nguyen, M.M.; Wang, D.; Pascal, L.E.; Guo,W.; Xu, Y.; Ai, J.; Deng, F.M.; Masoodi, K.Z.; Yu, X.;, et al. DHX15 promotes prostate cancer progression by stimulating Siah2-mediated ubiquitination of androgen receptor. Oncogene. 2018;37,638–650. doi:10.1038/onc.2017.371

91. Subhash, V.V.; Tan, S.H.; Tan, W.L.; Yeo, M.S.; Xie, C.; Wong, F.Y.; Kiat, Z.Y.; Lim, R.; Yong, W.P. GTSE1 expression represses apoptotic signaling and confers cisplatin resistance in gastric cancer cells. BMC Cancer. 2015,15,550. doi:10.1186/s12885-015-1550-0

92. Alvarez-Fernández, M.; Medema, R.H. Novel functions of FoxM1: from molecular mechanisms to cancer therapy. Front Oncol. 2013,3,30. doi:10.3389/fonc.2013.00030

93. Scharer, C.D.; McCabe, C.D.; Ali-Seyed, M.; Berger, M.F.; Bulyk, M.L.; Moreno, C.S. Genome-wide promoter analysis of the SOX4 transcriptional network in prostate cancer cells. Cancer Res. 2009;69,709–717. doi:10.1158/0008-5472.CAN-08-3415

94. Hu, J.Y.; Yi, W.; Wei, X.; Zhang, M.Y.; Xu, R.; Zeng, L.S.; Huang, Z.J.; Chen, J.S. miR-601 is a prognostic marker and suppresses cell growth and invasion by targeting PTP4A1 in breast cancer. Biomed Pharmacother. 2016,79,247–253. doi:10.1016/j.biopha.2016.02.014

95. Wanajo, A.; Sasaki, A.; Nagasaki, H.; Shimada, S.; Otsubo, T.; Owaki, S.; Shimizu, Y.; Eishi, Y.; Kojima, K.; Nakajima, Y.;, et al. Methylation of the calcium channel-related gene, CACNA2D3, is frequent and a poor prognostic factor in gastric cancer. Gastroenterology. 2008,135,580–590. doi:10.1053/j.gastro.2008.05.041

96. Jia, Y.; Yang, Y.; Brock, M.V.; Zhan, Q.; Herman, J.G.; Guo, M. Epigenetic regulation of DACT2, a key component of the Wnt signalling pathway in human lung cancer. J Pathol. 2013;230,194–204. doi:10.1002/path.4073

97. Diboun, I.; Wernisch, L.; Orengo, C.A.; Koltzenburg, M. Microarray analysis after RNA amplification can detect pronounced differences in gene expression using limma. BMC Genomics. 2006,7,252. doi:10.1186/1471-2164-7-252

98. Reiner-Benaim, A. FDR control by the BH procedure for two-sided correlated tests with implications to gene expression data analysis. Biom J. 2007,49,107–126.

99. Caspi, R.; Billington, R.; Ferrer, L.; Foerster, H.; Fulcher, C.A.; Keseler, I.M.; Kothari, A.; Krummenacker, M.; Latendresse, M.; Mueller, L.A.;, et al. The MetaCyc database of metabolic pathways and enzymes and the BioCyc collection of pathway/genome databases. Nucleic Acids Res. 2016,44,D471–D480. doi:10.1093/nar/gkv1164

100. Kanehisa, M.; Sato, Y.; Furumichi, M.; Morishima, K.; Tanabe, M. New approach for understanding genome variations in KEGG. Nucleic Acids Res. 2019,47,D590–D595. doi:10.1093/nar/gky962

101. Schaefer, C.F.; Anthony, K.; Krupa, S.; Buchoff, J.; Day, M.; Hannay, T.; Buetow, K.H. PID: the Pathway Interaction Database. Nucleic Acids Res 2009,37,D674–D679. doi:10.1093/nar/gkn653

102. Fabregat, A.; Jupe, S.; Matthews, L.; Sidiropoulos, K.; Gillespie, M.; Garapati, P.; Haw, R.; Jassal, B.; Korninger, F.; May, B.;, et al. The Reactome Pathway Knowledgebase. Nucleic Acids Res. 2018,46,D649–D655. doi:10.1093/nar/gkx1132

103. Dahlquist, K.D.; Salomonis, N.; Vranizan, K.; Lawlor, S.C.; Conklin, B.R. GenMAPP, a new tool for viewing and analyzing microarray data on biological pathways. Nat Genet. 2002;31,19–20. doi:10.1038/ng0502-19

104. Subramanian, A, Tamayo, P, Mootha, VK, Mukherjee, S.; Ebert, B.L.; Gillette, M.A.; Paulovich, A.; Pomeroy, S.L.; Golub, T.R.; Lander, E.S.;, et al. Gene set enrichment analysis: a knowledge-based approach for interpreting genome-wide expression profiles. Proc Natl Acad Sci. U. S. A. 2005,102,15545–15550. doi:10.1073/pnas.0506580102

105. Mi, H.; Huang, X.; Muruganujan, A.; Tang, H.; Mills, C.; Kang, D.; Thomas, P.D. PANTHER version 11: expanded annotation data from Gene Ontology and Reactome pathways, and data analysis tool enhancements. Nucleic Acids Res. 2017;45,D183–D189. doi:10.1093/nar/gkw1138

106. Petri, V.; Jayaraman, P.; Tutaj, M.; Hayman, G.T.; Smith, J.R.; De Pons, J.; Laulederkind, S.J.; Lowry, T.F.; Nigam, R.; Wang, S.J.;, et al. The pathway ontology - updates and applications. J Biomed Semantics. 2014,5,7. doi:10.1186/2041-1480-5-7

107. Jewison, T.; Su, Y.; Disfany, F.M.; Liang, Y.; Knox, C.; Maciejewski, A.; Poelzer, J.; Huynh, J.; Zhou, Y.; Arndt, D.;, et al. SMPDB 2.0: big improvements to the Small Molecule Pathway Database. Nucleic Acids Res. 2014,42,D478–D484. doi:10.1093/nar/gkt1067

108. Chen, J.;Bardes, E.E.; Aronow, B.J.; Jegga, A.G. ToppGene Suite for gene list enrichment analysis and candidate gene prioritization. Nucleic Acids Res. 2009,37,W305–W311. doi:10.1093/nar/gkp427

109. Ashburner, M.; Ball, C.A.; Blake, J.A.; Botstein, D.;Butler, H.; Cherry, J.M.; Davis, A.P.; Dolinski, K.; Dwight, S.S.; Eppig, J.T.;, et al. Gene ontology: tool for the unification of biology. The Gene Ontology Consortium. Nat Genet. 2000,25,25–29. doi:10.1038/75556

110. Alanis-Lobato, G.; Andrade-Navarro, M.A.; Schaefer, M.H. HIPPIE v2.0: enhancing meaningfulness and reliability of protein-protein interaction networks. Nucleic Acids Res. 2017,45,D408–D414. doi:10.1093/nar/gkw985

111. Orchard, S.; Ammari, M.; Aranda, B.; Breuza, L.; Briganti, L.; Broackes-Carter, F.;Campbell, N.H.; Chavali, G.; Chen, C.; del-Toro, N.;, et al. The MIntAct project--IntAct as a common curation platform for 11 molecular interaction databases. Nucleic Acids Res. 2014;42,D358–D363. doi:10.1093/nar/gkt1115

112. Chatr-Aryamontri, A, Oughtred, R, Boucher, L, Rust, J, Chang, C, Kolas, NK, O’Donnell, L, Oster, S, Theesfeld, C, Sellam, A et al. The BioGRID interaction database: 2017 update. Nucleic Acids Res. 2017,45,D369–D379. doi:10.1093/nar/gkw1102

113. Keshava Prasad, TS, Goel, R, Kandasamy, K, Keerthikumar, S, Kumar, S, Mathivanan, S, Telikicherla, D, Raju, R, Shafreen, B, Venugopal, A, et al. Human Protein Reference Database--2009 update. Nucleic Acids Res. 2009,37,D767–D772. doi:10.1093/nar/gkn892

114. Licata, L.; Briganti, L.; Peluso, D.; Perfetto, L.; Iannuccelli, M.; Galeota, E.; Sacco, F.; Palma, A.; Nardozza, A.P.; Santonico, E.;, et al. MINT, the molecular interaction database: 2012 update. Nucleic Acids Res. 2012;40,D857–D861. doi:10.1093/nar/gkr930

115. Isserlin, R, El-Badrawi, RA, Bader, GD. The Biomolecular Interaction Network Database in PSI-MI 2.5. Database (Oxford). 2011,2011,baq037. doi:10.1093/database/baq037

116. Pagel, P.; Kovac, S.; Oesterheld, M.; Dunger-Kaltenbach, I.; Frishman, G.; Montrone, C.; Mark, P.; Stümpflen, V.; Mewes, H.W.; Ruepp, A.;, et al. The MIPS mammalian protein-protein interaction database. Bioinformatics. 2005;21,832–834. doi:10.1093/bioinformatics/bti115

117. Salwinski, L, Miller, CS, Smith, AJ, Pettit, FK, Bowie, JU, Eisenberg, D. The Database of Interacting Proteins: 2004 update. Nucleic Acids Res. 2004,32,D449–D451. doi:10.1093/nar/gkh086

118. Shannon, P, Markiel, A, Ozier, O, Baliga, NS, Wang, JT, Ramage, D, Amin, N, Schwikowski, B, Ideker, T. Cytoscape: a software environment for integrated models of biomolecular interaction networks. Genome Res. 2003,13,2498–2504. doi:10.1101/gr.1239303

119. Przulj, N, Wigle, DA, Jurisica, I. Functional topology in a network of protein interactions. Bioinformatics. 2004,20,340–348. doi:10.1093/bioinformatics/btg415

120. Joy, M.P.; Brock, A.; Ingber, D.E.; Huang, S. High-betweenness proteins in the yeast protein interaction network. J Biomed Biotechnol. 2005,2005,96–103. doi:10.1155/JBB.2005.96

121. Lehtinen, S, Marsellach, FX, Codlin, S, Schmidt, A, Clément-Ziza, M, Beyer, A, Bähler, J, Orengo, C, Pancaldi, V. Stress induces remodelling of yeast interaction and co-expression networks. Mol Biosyst. 2013,9,1697–1707. doi:10.1039/c3mb25548d

122. Hsu, C.W.; Juan, H.F.; Huang, H.C. Characterization of microRNA-regulated protein-protein interaction network. Proteomics. 2008,8,1975–1979. doi:10.1002/pmic.200701004

123. Stelzl, U.; Worm, U,; Lalowski, M,; Haenig, C.; Brembeck, F.H.; Goehler, H.; Stroedicke, M.; Zenkner, M.; Schoenherr, A.; Koeppen, S.;, et al. A human protein-protein interaction network: a resource for annotating the proteome. Cell. 2005,122,957–968. doi:10.1016/j.cell.2005.08.029

124. Zaki, N, Efimov, D, Berengueres, J. Protein complex detection using interaction reliability assessment and weighted clustering coefficient. BMC Bioinformatics. 2013,14,163. doi:10.1186/1471-2105-14-163

125. Xia, J.; Benner, M.J.; Hancock, R.E. NetworkAnalyst--integrative approaches for protein-protein interaction network analysis and visual exploration. Nucleic Acids Res. 2014,42,W167–w174. doi:10.1093/nar/gku443

126. Vlachos, I.S.; Paraskevopoulou, M.D.; Karagkouni, D.; Georgakilas, G.; Vergoulis, T.; Kanellos, I.; Anastasopoulos, I.L.; Maniou, S.; Karathanou, K.; Kalfakakou, D.;, et al. DIANA-TarBase v7.0: indexing more than half a million experimentally supported miRNA:mRNA interactions. Nucleic Acids Res. 2015,43,D153–D159. doi:10.1093/nar/gku1215

127. Chou, C.H.; Shrestha, S.; Yang, C.D.; Chang, N.W.; Lin, Y.L.; Liao, K.W.; Huang, W.C.; Sun, T.H.; Tu, S.J.; Lee, W.H.;, et al. miRTarBase update 2018: a resource for experimentally validated microRNA-target interactions. Nucleic Acids Res. 2018,46,D296–D302. doi:10.1093/nar/gkx1067

128. Wang, S.; Sun, H.; Ma, J.; Zang, C.; Wang, C.; Wang, J.; Tang, Q.; Meyer, C.A.; Zhang, Y.; Liu, X.S. Target analysis by integration of transcriptome and ChIP-seq data with BETA. Nat Protoc. 2013,8,2502–2515. doi:10.1038/nprot.2013.150

